# A general design of caging-group free photoactivatable fluorophores for live-cell nanoscopy

**DOI:** 10.1101/2021.11.15.468659

**Authors:** Richard Lincoln, Mariano L. Bossi, Michael Remmel, Elisa D’Este, Alexey N. Butkevich, Stefan W. Hell

## Abstract

The controlled switching of fluorophores between non-fluorescent and fluorescent states is central to every superresolution fluorescence microscopy (nanoscopy) technique, and the exploration of radically new switching mechanisms remains critical to boosting the performance of established, as well as emerging superresolution methods. Photoactivatable dyes offer significant improvements to many of these techniques, but often rely on photolabile protecting groups that limit their applications. Here we describe a general method to transform 3,6-diaminoxanthones into caging-group free photoactivatable fluorophores. These photoactivatable xanthones (PaX) assemble rapidly and cleanly into highly fluorescent, photo- and chemically stable pyronine dyes upon irradiation with light. The strategy is extendable to carbon- and silicon-bridged xanthone analogs, yielding a new family of photoactivatable labels spanning much of the visible spectrum. Our results demonstrate the versatility and utility of PaX dyes in fixed and live-cell fluorescence microscopy, and both coordinate-targeted stimulated emission depletion (STED) and coordinate-stochastic single-molecule localization superresolution microscopy (SMLM).

## MAIN

Fluorescence nanoscopy has revolutionized our ability to visualize (living) cells by extending the limits of optical imaging to single-digit nanometer resolution, and by enabling minimally-invasive observation of the internal nanoscale structures and dynamics of biological samples with molecular specificity^1–3^. Central to these techniques are chemically-specific fluorescent labels and the intrinsic control between fluorescent (on) and non-fluorescent (off) states of the fluorophores. This sequential off-on transition is key to separating adjacent fluorophores at molecule-scale proximities. Photoactivatable or caged dyes—in which the off-on transition is irreversible and triggered by light—render these nanoscopy techniques very powerful, because they eliminate the need for specific imaging buffers and high intensities of UV light. Such needs are prevalent in single molecule based microscopy, such as those called PALM or STORM, to drive commonly used fluorophores (e.g. cyanines) between non-fluorescent and fluorescent states^4–6^, as well as enabling high-density single particle tracking^7–9^. Most recently, photoactivatable dyes have been used to reduce fluorescence background in DNA-PAINT^10^, and to increase the number of cellular structures that may be simultaneously imaged in stimulated emission depletion (STED) microscopy through channel duplexing^11^.

Rhodamine dyes have emerged as some of the most widely employed fluorophores in fluorescence microscopy and nanoscopy, due to the remarkable tunability of their optical and chemical properties^12–14^, cell membrane permeability^15^, photostability^16^, and brightness^17^. In particular, silicon rhodamines^18^ are often favored for their intrinsic red-shifted emission, fluorogenic behaviour^19^, and live-cell compatibility^20–22^. However, the reported caging strategies for rhodamines rely on ‘locking’ the dyes in a non-fluorescent form, either through installation of photolabile protecting groups on the nitrogen atoms (such as with nitroveratryloxycarbonyl^6,23,24^ or nitroso^25^ groups) or by synthetic transformation of the lactone ring into the corresponding cyclic α-diazoketones^8,26^. The former strategy restricts the attainable substitution patterns, reduces water solubility, and yields stoichiometric amounts of potentially toxic byproducts upon photoactivation. The latter strategy instead suffers from varying uncaging efficiencies, and the concomitant formation of non-fluorescent side-products, whose abundance depends on the medium and substitution pattern^26,27^.

Therefore, caging-group free, compact photoactivatable and biocompatible fluorophores would be highly desired in fluorescence microscopy and nanoscopy applications, enabling lower molecular weight labels, provided that the photoactivation is rapid, complete and byproduct-free. Most recently, Frei *et al.* demonstrated the photoactivation of a Si-pyronine analogue, where the fluorophore was initially masked with an exocyclic double bond at the 9-position of the Si-xanthene scaffold^28^. Upon UV irradiation in aqueous solution, protonation of the exocyclic double bond yielded the fluorescent 9-alkyl-Si-pyronine. The resulting cationic fluorophore, however, was a reactive species susceptible to nucleophilic addition of thiols and water, with a rapid equilibrium in favor of the non-fluorescent addition products, thus limiting its applicability.

Inspired by the long-established radical photochemistry of benzophenone and other diarylketones, we have now designed and report herein a new class of functionalized xanthones, which upon one or two-photon excitation, convert efficiently and cleanly into the corresponding dihydropyran-fused pyronine dyes. These photoactivatable xanthone (PaX) dyes can be prepared from readily available starting materials via a straightforward and efficient three-step synthetic route, also compatible with carbon- and silicon-bridged analogs, to yield a family of fluorophores spanning much of the visible spectrum. In particular, PaX-derived Si-pyronine dyes display good live-cell compatibility, resilience to nucleophiles, and an unprecedented photostability for orange-emitting (TAMRA-like) fluorophores. We highlight the utility of PaX dyes and labels in optical microscopy and nanoscopy techniques, in fixed and living cells, including STED and PALM.

## RESULTS AND DISCUSSION

### Synthetic design and proposed mechanism of photoactivation

In our search for minimalistic photoactivatable fluorophores, we reasoned that the concept of employing photochemical reactions to assemble or ‘lock’ fluorophores, rather than ‘unlocking’ photocleavable caging elements, provided an improved alternative to caged rhodamine dyes (Fig. 1a). Diaryl ketones are known photoinitiators of radical reactions^29^ due to their high inherent rate of intersystem crossing (via spin-orbit coupling) and triplet states with diradical character^30,31^. We hypothesized that their photochemistry would be extendable to 3,6-diaminoxanthones, utilized as precursors in the synthesis of rhodamines^32–34^. With introduction of a suitable intramolecular radical trap onto the xanthone scaffold, a juxtaposition of a radical source (diaryl ketone) and a radical trap (styrene) could be exploited to photoassemble 9-alkoxypyronine fluorophores through a light-triggered cascade (Fig. 1b)^35^.

**Fig. 1.**
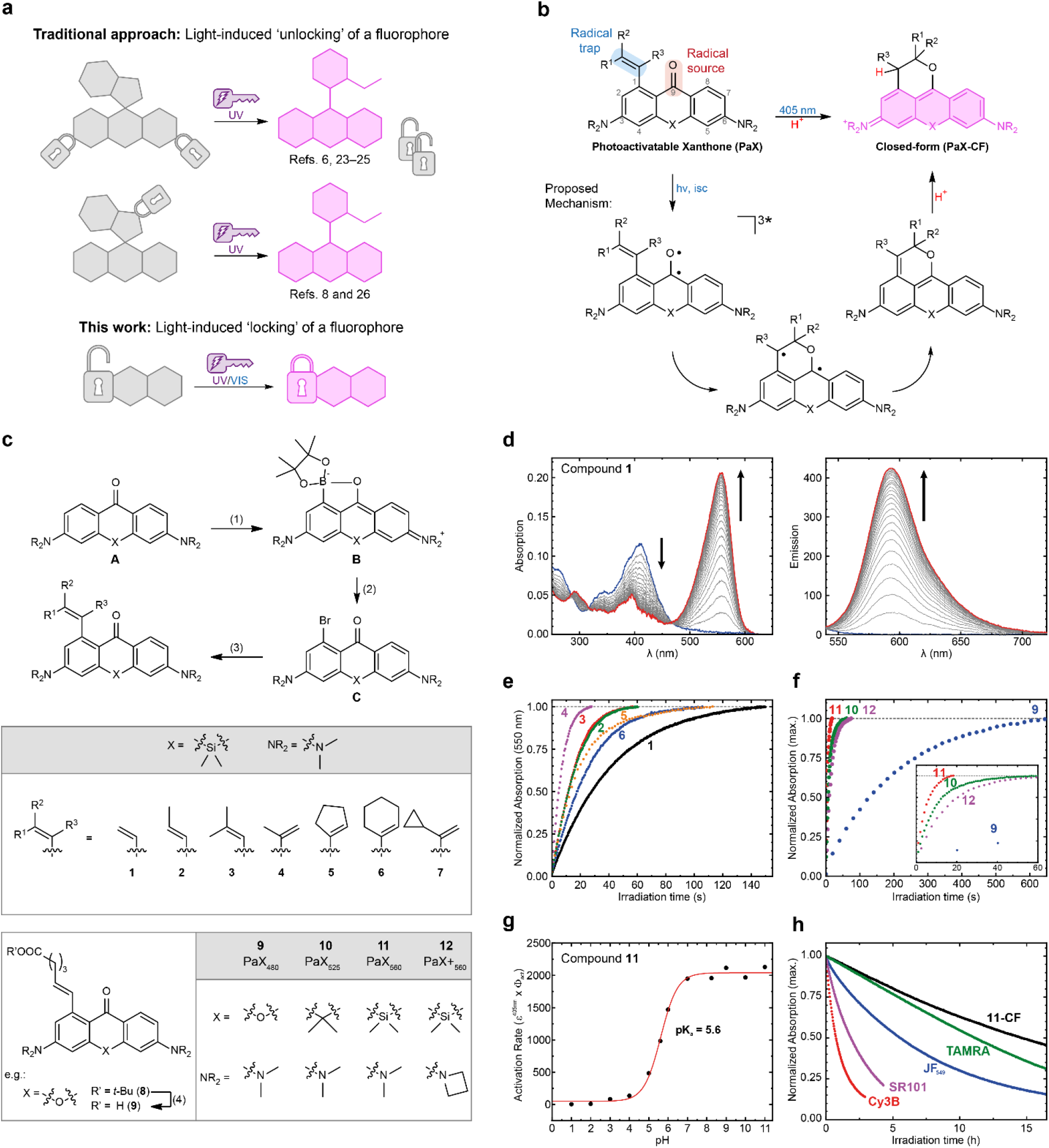
Design, synthesis, and characterization of photoactivatable xanthone (PaX) dyes. **a**, Whereas traditional strategies for photoactivatable dyes for nanoscopy rely on the release (‘unlocking’) of caging groups, our approach relies on the light-induced assembly (‘locking’) of a fluorophore. **b**, General structure of a PaX with 1-alkenyl radical trap and its 9-alkoxypyronine photoproduct (closed-form, **CF**), and proposed photoactivation mechanism. **c**, Synthetic route for the preparation of PaX. (1) B_2_pin_2_, [Ir(cod)(OMe)]_2_, AsPh_3_, *n*-octane, 120°C, 22 h; (2) CuBr_2_, KF, pyridine, DMSO/H_2_O, 80°C, 30 min; (3) RB(OH)_2_, RBpin or RBF_3_K (R = alkenyl), Pd(dppf)Cl_2_, K_2_CO_3_, dioxane/H_2_O, 80°C, 3–18 h; (4). CH_2_Cl_2_/TFA 3:1, r.t., 1 h. **d**, Temporal evolution of absorption and fluorescence spectra of compound **1** irradiated in phosphate buffer (100 mM, pH = 7; λ_act_ = 405 nm). **e**, Comparative photoactivation kinetics of Si-bridged PaX **1**–**6**, same conditions as d. **f**, Comparative photoactivation kinetics of PaX dyes **9**–**12**, same conditions as d. **g**, Comparative photoactivation kinetics of **11** in phosphate buffer (100 mM) at different pH values (λ_act_ = 405 nm). **h**, Photo-fatigue resistance of compound **11-CF** and established commercial fluorophores, with similar spectral properties, measured in phosphate buffer (λ_exc_ = 530 nm).

To investigate the effects of substitution of the radical acceptor, we first synthesized a series of photoactivatable Si-xanthones (**1**–**7**, Fig. 1c). The target compounds were prepared by an Ir-catalyzed, chelation-assisted, *ortho*-selective C–H borylation of the diaryl ketone (**A)**^36^. Conversion of the resulting boronate ester (**B**) into the corresponding aryl bromide (**C**) was done with a CuBr2–pyridine system37 in the presence of KF (for details, see the Supplementary Information). A series of alkene substituents were then installed using standard Suzuki-Miyaura cross-coupling reaction conditions. Compounds **1**–**6** showed a strong absorption band (ε ~ 10^4^ M^−1^cm^−1^) at around 400 nm, characteristic of Michler’s ketone and its analogues (see Supplementary Table S1 for photophysical characterization). Upon irradiation in protic media, for example in dilute aqueous solutions (phosphate buffer, 100 mM, pH = 7), compounds **1**–**6** underwent rapid and complete conversion to give highly fluorescent ‘closed-form’ (CF) products with TMR-like spectral properties (**1-CF**–**6-CF**; Fig. 1d and Supplementary Fig. 1). LC-MS analysis of the reaction mixtures revealed no formation of byproducts for most samples. The measured quantum yields of photoactivation (Φ_PA_) ranged from 1×10^−2^ to 6×10^−2^ (Supplementary Table 1). The rate of photoactivation was slowest for vinyl-substituted compound **1**, increased with additional substitution of the alkene and was highest for compound **4**, possibly due to a favorable orientation of the alkene induced by the methyl substituent (Fig. 1e). To confirm the formation of predicted 9-alkoxypyronine product **1-CF**, a solution of compound **1** in methanol was irradiated with a 405 nm LED in a batch photoreactor (see Supplementary Information for details), and the resulting product was isolated and fully-characterized by NMR and HR-MS analysis (Supplementary Fig. 2), confirming the expected dihydropyran ring fusion. A solvent-dependent protonation step was confirmed by conducting photolysis in methanol-*d*_4_ (resulting in deuterium incorporation at the benzylic position) and by the absence of efficient photoactivation in aprotic solvents such as 1,4-dioxane. Deoxygenating the solvent increased the rate of photoactivation, confirming the role of xanthone triplet state. Photolysis of **1** (4.8 μM) in the presence of millimolar concentrations of the radical trap 4-hydroxy-TEMPO resulted in the formation of a PaX-TEMPO adduct (Supplementary Fig. 3), however the radical clock probe **7** showed no evidence of cyclopropane ring-opening upon photoactivation (Supplementary Fig. 4).

To render the PaX dyes suitable for bioconjugation, xanthone **9** (PaX_480_), anthrone **10** (PaX_525_), and Si-xanthone **11** (PaX_560_), along with its bis-azetidine analog **12** (PaX+_560_), have been prepared (Fig. 1c) bearing alkenyl substituents with a short carboxylate-terminated spacer. The keto forms of **9**, **10**, and **11** showed initial absorbance maxima at 399, 408 and 414 nm, respectively (Supplementary Fig. 5 and Supplementary Table 2). The rate of photoactivation yielding the pyronine dyes (with absorption/emission maxima at 480/514 nm for **9-CF**, 524/564 nm for **10-CF,** and 558/596 nm for **11-CF**) decreased in the order **11 > 10 > 9** (Fig. 1f). As we expected, the azetidine auxochromic groups had little impact on the spectral properties of both the Si-xanthone (**12**) and Si-pyronine (**12-CF**) forms, but instead reduced the rate of photoactivation compared to the bis(*N*,*N*-dimethylamino) analogue (**11**). The fluorophore **12-CF** demonstrated remarkably improved quantum efficiency of emission (QY 0.92 vs. 0.48 for **11-CF**), which can be attributed to the suppression of transfer into a twisted internal charge transfer (TICT) state upon excitation^17^.

Screening the photoactivation properties of **11** over a range of biologically-relevant pH values (Fig. 1g and Supplementary Fig. 6a) revealed a 6-fold decrease in the photoactivation rate in acidic media (pH 4.3) as compared to neutral, and only little rate changes at basic pH values (up to 9.0). The low pH-dependence on the activation rate is similar to previous observations on the protonation of the benzophenone triplet excited state^38,39^, supporting the assumed involvement of this diradical in the activation mechanism. At high pH values hydrolysis of **11-CF** was observed (Supplementary Fig. 6b), however, there was no observed difference in the absorption and emission spectra of **11-CF** and no change in product composition as measured by LC-MS up to pH 8.5 (Supplementary Fig. 6c), indicating little observable pH-sensitivity for this dye across the biologically relevant pH range. Furthermore, photoactivation of **11** proceeded normally in buffered solutions (pH 7) containing 10 mM cysteamine, anticipating the lack of unwanted radical or electrophilic reactivity towards biomolecules, and a potential orthogonality with SMLM blinking buffers used for cyanine dyes^4^. Finally, we assessed the photostability of **11-CF**, benchmarking it against a series of commercially available dyes with similar spectral properties (Fig. 1h), and found that **11-CF** outperformed all of the tested fluorophores.

### Caging-group-free photoactivatable labels for fluorescence nanoscopy

Encouraged by the versatility of the PaX mechanism, we proceeded to construct targeted labels for fluorescence microscopy and nanoscopy. For indirect immunolabelling (with secondary antibodies or nanobodies), an amino-reactive NHS ester (**13**) and a thiol-reactive maleimide (**14**) derivative of PaX_560_ were prepared (Fig. 2a), along with the NHS esters of PaX_480_, PaX_525_, and PaX+_560_ (Supplementary Fig. 7a, **16**-**18**). For actin labeling in fixed cells, a phalloidin derivative (**15**) of PaX_560_ was assembled (Fig. 2a).

**Fig. 2.**
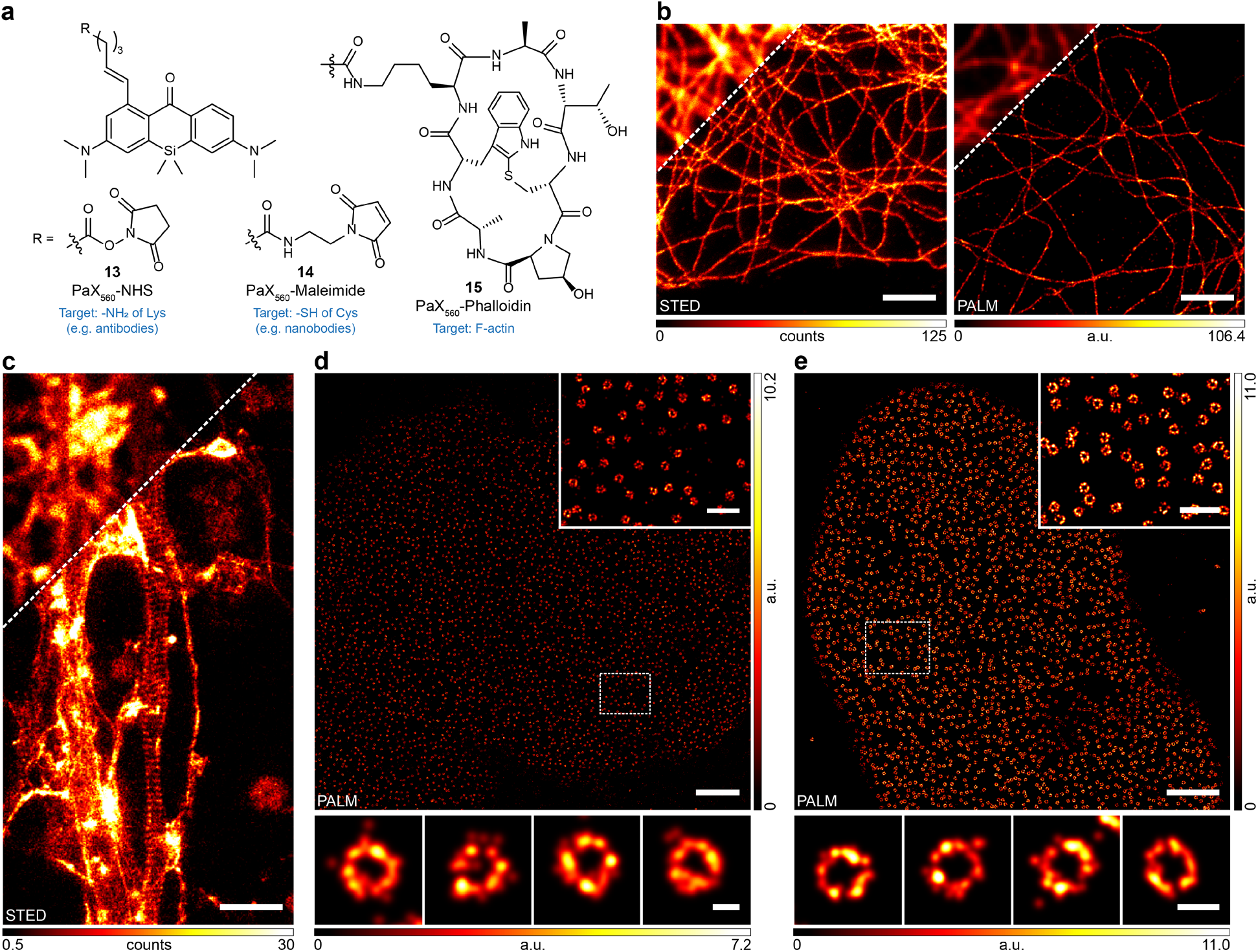
Photoactivatable labels for optical nanoscopy. **a**, Structures of PaX_560_ derivatives for bioconjugation (**13, 14**) and actin labelling (**15**). **b**, STED (left) and PALM (right) images of microtubules in COS-7 cells labelled via indirect immunofluorescence with a secondary antibody bearing **13**. Preactivation to **13-CF** for STED imaging was achieved with widefield illumination (AHF, DAPI filter set F46-816). **c**, Actin structures of the periodic membrane cytoskeleton in the axon of fixed primary hippocampal neuron cultures labelled with **15** and mounted in Mowiol. Preactivation to **15-CF** for STED imaging was achieved with widefield illumination (AHF, EGFP filter set F46-002) followed by a 518 nm laser. Image data were smoothed with a 1-pixel low pass Gaussian filter. **d**, PALM image of NPCs in COS-7 cells labelled via indirect immunofluorescence with an anti-NUP98 primary antibody and a secondary nanobody labelled with **14**. Inset: Magnified region marked in overview image. Bottom row: Individual NPCs. **e**, PALM image of NPCs in HK cells expressing NUP107-mEGFP anti-NUP98 labelled with anti-GFP nanobodies conjugated to **14**. Inset: Magnified region marked in overview image. Bottom row: Individual NPCs. Scale bars: 2 μm (b-e), 500 nm (d,e insets), 50 nm (d, bottom row), 100 nm (e, bottom row).

Thanks to their remarkable photo-fatigue resistance, we reasoned that PaX dyes would be strong candidates for STED imaging. We tested their performance in indirect immunofluorescence labelling of microtubules in fixed samples. The closed fluorescent form of the dye was generated *in situ* upon photoactivation before STED imaging using a 561 nm excitation and a 660 nm STED beam for de-excitation. Super-resolved images of microtubules were successfully acquired for antibody conjugates of PaX_560_ (**13**, Fig. 2b) as well as of PaX_525_ and PaX+_560_ (**17**,**18**, Supplementary Fig. 7b) demonstrating their compatibility with STED nanoscopy. The specificity of PaX_560_-phalloidin (**15**) for the actin structure was validated in fixed neuron cultures, in which the membrane cytoskeleton of the axon was visualized by STED (Fig. 2c).

Next, we tested the performance of our photoactivatable labels in single molecule localization microscopy (SMLM)^40,41^. To this aim, PALM imaging was carried out on indirectly-immunolabelled microtubules, and super-resolved images could be obtained for antibody conjugates bearing **13** (Fig. 2b) and **16**-**18** (Supplementary Fig. 7c). Thanks to the efficient photoactivation mechanism, very low powers (<100 μW) of the activation light were required. Importantly, all samples were imaged in PBS or in Mowiol, without the need for special blinking buffers or photostabilizing agents.

To further benchmark the utility of PaX labels for PALM imaging, indirect immunofluorescent labelling of nuclear pore complexes (NPCs) was conducted with a primary anti-NUP-98 antibody and secondary anti-rabbit nanobodies bearing **14** (Supplementary Fig. 8a,b). The PALM images of NPCs (Fig. 2d) were comparable in quality to those acquired through more mechanically- and temporally- demanding methods (e.g. qPAINT^42^). Alternatively, the large-sized (~150 kDa) primary antibodies can be avoided, as previously reported to improve labelling precision,^43^ in cell lines expressing an mEGFP^44^ (~27 kDa) fusion to NUP107 when combined with anti-GFP nanobodies labelled with **14** (Fig. 2e).

Given the difference in photoactivation rates for the PaX dyes, we surmised that two complimentary labels could be used for multiplexing purposes by subsequently applying a lower and a higher doses of activation light. We explored this possibility by indirect immunofluorescence of clathrin-coated pits and microtubules with the oxygen- and silicon-bridged PaX_480_ (**16**) and PaX_560_ (**13**), respectively. Confocal imaging performed before activation (Supplementary Fig. 9a), after a low photoactivation dose (selectively activating compound **13**, Supplementary Fig. 9b), and after a higher dose of light (activating **16**, Supplementary Fig. 9c), enabled the visualization and discrimination of the structures of interest without need of image post-processing (Supplementary Fig. 9d). We then expanded this strategy to oxygen- and carbon-bridged PaX_480_ (**16**) and PaX_525_ (**17**), which can be differentiated in an analogous fashion (Supplementary Fig. 9e). Similarly, sequential two-color (with a single detector) PALM images could be acquired for PaX_480_ (**16**) and PaX_560_ (**13**). This was achieved by first using a sufficiently low photoactivation dose to selectively activate compound **13** and imaging until photoactivation events were exhausted. Compound **16** was then imaged by applying a higher photoactivation light dose. In this case, the two PaXs were detected in their respective imaging channels using appropriate excitation laser and filter combinations (Supplementary Fig. 9f).

### Targeted labels for live-cell imaging

To evaluate the compatibility of the PaX photoactivation mechanism for live imaging, we first prepared PaX_560_ constructs (Fig. 3a) containing mitochondria-targeting triphenylphosphonium (**19**) and lysosome-targeting Pepstatin A (**20**) moieties, as these two selected organelles represent the extreme pH values found within the cell (pH 7.8 for mitochondrial matrix and down to 4.5 in lysosomal lumen). COS-7 cells were co-incubated with **19** and MitoTracker Deep Red and imaged with confocal microscopy before and after photoactivation with 365 nm light (Fig. 3b). The resulting fluorescence of **19-CF** co-localized strongly with the MitoTracker signal (Pearson correlation coefficient *r* = 0.94). Similarly, COS-7 cells concurrently labelled with the Pepstatin A conjugate **20** and the lysosome-targeting fluorophore SiR-lysosome,^19^ demonstrated colocalization after photoactivation with *r* = 0.84. These results confirmed that the photoactivation mechanism is suitable for live-cell imaging in both high and low pH cellular environments.

**Fig. 3.**
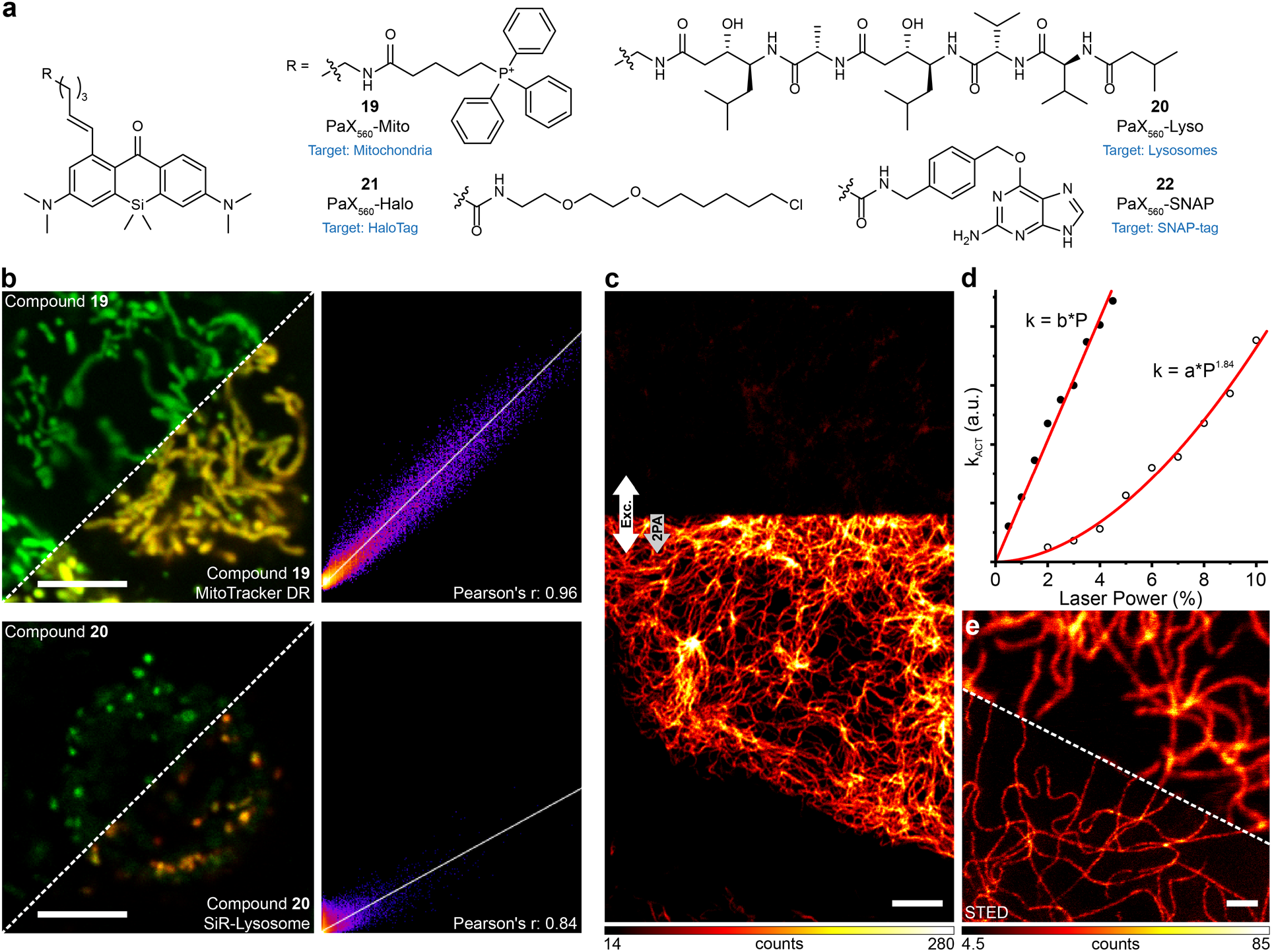
Imaging with photoactivatable PaX labels in living cells. **a**, Structures of PaX_560_ derivatives (**19**–**22**) for live-cell imaging. **b**, Confocal images and corresponding Pearson correlation analysis of COS-7 cells co-incubated with **19** (200 nM) and MitoTracker Deep Red (50 nM, top row) or **20** (20 nM) and SiR-lysosome (200 nM, bottom row). Conversion to **19-CF** and **20-CF** was achieved by a 365 nm laser. **c**, Confocal image of vimentin filaments labelled with **21** (200 nM) in U2OS cells before activation (upper portion) and with 2-photon activation (lower portion, indicated by the arrows). **d,** Activation rate vs laser power plot for a one-photon (365 nm) or two-photon activation laser (810 nm). The lines represent fittings to a linear or a quadratic function for one- or two-photon activation, respectively. **e.** Confocal and STED images of the same sample following activation by a 405 nm laser. Scale bars: 5 μm (b, c), 1 μm (e).

Self-labelling protein tags, such as HaloTag and SNAP-tag, are well-established tools for targeting synthetic fluorophores to specific proteins in live-cell imaging^45^. Seeking to exploit this targeting strategy, we prepared the HaloTag-specific chloroalkane derivative (**21**) and the SNAP-tag specific *O*^6^-benzylguanine derivative (**22**) of PaX_560_ (Fig. 3a). Chloroalkane derivatives of PaX_480_, PaX_525_, and PaX+_560_ were additionally prepared (Supplementary Fig. 10, **23**-**25**).

Upon covalent linking of PaX_560_-Halo (**21**) with the HaloTag protein, we observed a 7.8-fold increase in the photoactivation rate of the dye (Supplementary Fig. 11a–d). Complete reaction of **21** with HaloTag7 was confirmed by mass spectroscopy, even with only a mild excess (~1.1 eqs.) of the protein (Supplementary Fig. 11b,d). No major fluorescence intensity changes were observed for **21-CF** covalently bound to HaloTag vs. free **21-CF** in buffered solution (Supplementary Fig. 11e,f). However, similar labelling efficiency and a greater fluorogenic response was observed upon binding of PaX_560_-SNAP (**22**) to SNAP-tag with an 11-fold increase of photoactivation rate (Supplementary Fig. 12a–d) and a 3.3-fold fluorescence intensity increase of SNAP-tag bound **22-CF** vs free **22-CF** (Supplementary Fig. 12e,f).

We then assessed the feasibility of two-photon activation of PaX_560_-Halo (**21**) with 810 nm near-infrared (NIR) light, as shifting the excitation wavelength from UV to NIR-range advantageously reduces phototoxicity and increases imaging depth in tissues. U2OS cells stably expressing a Vimentin-HaloTag fusion construct^46^ were labelled with compound **21** and imaged with a confocal microscope equipped with a subpicosecond pulsed laser (Fig. 3c). The activation rate constant was determined for selected areas of the same sample by mono-exponential fitting of the activation rates measured with variable powers of a UV laser for one-photon activation (365 nm) or a subpicosecond pulsed laser for two-photon activation in the NIR (810 nm). Two photon activation was confirmed by the nearly-quadratic (1.84) dependence on the power of the excitation light (Fig. 3d). A pre-activated region of the same sample was further imaged using STED (at 660 nm) to resolve vimentin filaments with sub-diffraction resolution, confirming that live-cell STED was readily possible with compound **21** (Fig. 3e).

Photoactivatable fluorophores can also be utilized together with regular ‘always-on’ fluorescent dyes having similar spectral properties for color duplexing within a single excitation / detection channel, effectively doubling the number of available imaging channels in a confocal or STED system^11^. To demonstrate such possibility with PaX labels, U2OS cells stably expressing a vimentin-HaloTag fusion protein were concurrently labelled with PaX_560_-Halo (**21**) and abberior LIVE 560-Tubulin (AL-560) probe and imaged by confocal microscopy using a single detection channel (Fig. 4a–e). Initially, the AL-560-labelled tubulin filaments were visualized (Fig. 4a), followed by AL-560-photobleaching with high-intensity 560 nm excitation light (Fig. 4b); then, compound **21** was in turn photoactivated with a 405 nm laser to reveal the **21-CF**-labelled vimentin structure (Fig. 4c).

**Fig. 4.**
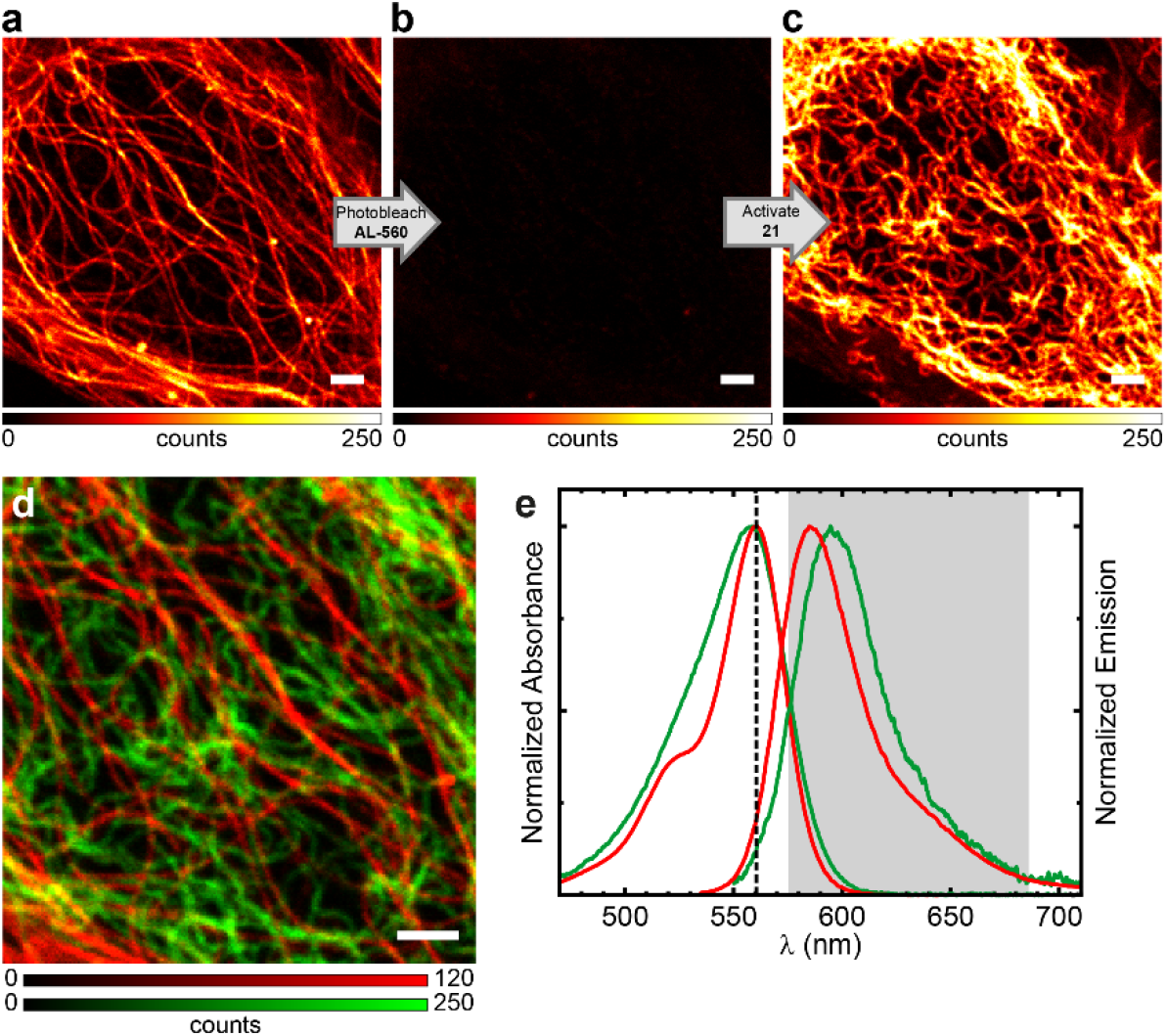
Channel duplexing with PaX labels. Confocal imaging of U2OS labelled with abberior LIVE 560 tubulin (AL-560, 500 nM) and vimentin filaments labelled with compound **21** (100 nM) before (**a**) and after (**b**) photobleaching of AL-560 and after (**c**) photoactivation by a 405 nm laser of **21**. **d**, Combined pseudo two-color image showing tubulin (red) and vimentin (green) filaments obtained by sequential imaging (**a**-**c**). **e**, Absorption and emission spectra of AL-560 (red) and **21-CF** (green), the excitation laser (dashed line) and detection window (grey) used. Scale bars: 2 μm (a-d).

Furthermore, two color imaging of two spectrally-distinct PaX dyes by selective activation was performed in living cells (i.e. discrimination by photoactivation efficiency). We demonstrated this experiment firstly with PaX_560_-Mito (**19**, specific for mitochondria) and the HaloTag-targeting derivative of PaX_480_-Halo (**23**, Supplementary Fig. 13a) and secondly with PaX_560_-Lyso (**20**, specific for lysosomes) and the HaloTag-targeting derivative of PaX_480_-Halo (**23**, Supplementary Fig. 13b).

Finally, we sought to explore the utility of the self-labeling protein tag substrates **21** and **22** for live-cell SMLM. U2OS cells stably expressing a SNAP-tag fusion with NUP10747 were labelled with PaX_560_-SNAP (**22**) and imaged with PALM (Fig. 5a). The reconstructed image shows largely complete circumferential labelling, which is remarkable given the one-to-one dye to protein ratio, highlighting the efficient labelling and efficient detection of activated PaX. Likewise, U2OS cells stably-expressing HaloTag fusion proteins with NUP9643 (another NPC protein) were labelled with PaX_560_-Halo (**21**). The reconstructed image (Fig. 5b) resolved the structural elements of the NPCs with even greater efficiency. Fixation of live-labelled samples (with HaloTag and SNAP-tag fusion proteins) also allowed PALM imaging with similar contrast. Thus the established fixation and permeabilization treatments used to preserve NUP structures^43^ for super-resolution imaging do not affect the PaX labels, or their photoactivation.

**Fig. 5.**
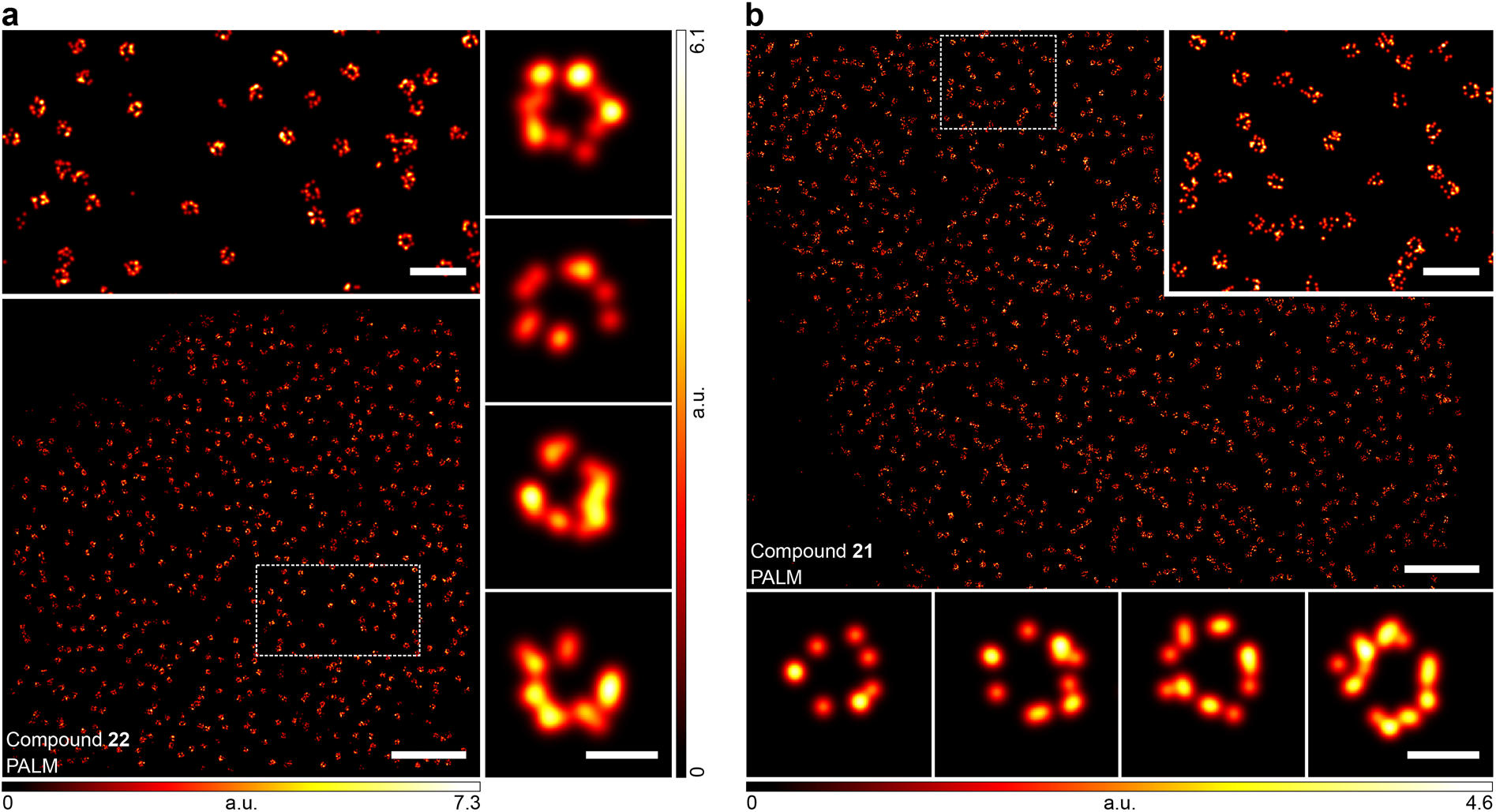
PALM imaging of NPCs in living cells using self-labelling PaX_560_ substrates. **a**, PALM image of U2OS stably expressing a NUP107-SNAP-tag construct labelled with **22**. Top: Magnified region marked in overview image. Right column: Magnified individual NPCs. **b**, PALM image of U2OS cells stably expressing a NUP96-HaloTag construct labelled with **21**. Inset: Magnified region marked in overview image. Bottom row: Magnified individual NPCs. Scale bars: 2 μm (a,b), 500 nm (a,b insets), 100 nm (a, right column; b, bottom row).

## CONCLUSION

We have introduced a general design strategy for caging-group free, bright and live-cell compatible photoactivatable dyes, suitable for a wide range of optical microscopy and nanoscopy techniques, including PALM and STED. The unique structural feature of these PaX dyes is the combination of a light-responsive 3,6-diaminoxanthone core functionalized with a intramolecular alkene radical trap, to give a highly-compact and intrinsically-uncharged, intact cell membrane-permeant label. Under one or two-photon activation, these compounds rapidly assemble into highly photostable fluorescent pyronine dyes. By changing the substitution pattern of PaX dyes, the kinetics of the photoactivation as well as the spectral properties can be tuned, allowing for both multiplexed pseudocolor as well as conventional multicolor imaging. The utility and versatility of PaX dyes was illustrated with a diverse range of target-specific probes and labelling strategies, for fixed and live-cell (superresolution) fluorescence microscopy experiments. We expect that our methodology will further stimulate the development of new photoactivatable probes and sensors for biological imaging and material science.

## METHODS

Detailed procedures for the synthesis of all compounds and their characterizations, as well as methods for cell culture, sample preparation, live and fixed cell labelling for microscopy and nanoscopy are given in the Supplementary Information. Image acquisition conditions for confocal and STED (Supplementary Table 3), and PALM (Supplementary Table 4) as well as detailed procedures for image processing and rendering are provided in the Supplementary Information.

## Supporting information

Supplementary compressed archive

## ACKNOWLEDGEMENTS

R.L. is grateful to the Max Planck Society for a Nobel Laureate Fellowship. We thank Prof. Stefan Jakobs (MPI-BPC, Göttingen) for providing the U2OS-Vim-Halo and U2OS-Vim-SNAP cells and the EMBL (Heidelberg) for providing the U-2 OS-ZFN-SNAP-NUP107, U-2 OS-CRISPR-NUP96-Halo, and HK-2xZFN-mEGFP-Nup107 cells. We thank the following people at the MPI for Medical Research (Heidelberg): Dr. Sebastian Fabritz (Mass Spectrometry Core Facility) for recording mass spectra of proteins and small molecules, Matthias Steigleder (Electronics Workshop) for building the custom 405 nm light source for Penn m1 photoreactor, Prof. Kai Johnsson (Department of Chemical Biology) for SNAP-tag protein, Dr. Miroslaw Tarnawski (Protein Expression and Characterization Facility) for HaloTag7 protein; and our colleagues at the Department of Optical Nanoscopy: Jasmine Hubrich for supporting the cell culture, Ayse Aktalay for assistance with labelling of nanobody samples, Dr. Jessica Matthias for critical reading of the manuscript.

## AUTHOR CONTRIBUTIONS

R.L. was responsible for the project’s conception and wrote the manuscript with input from A.N.B, M.L.B, and S.W.H; A.N.B. designed and validated the synthetic routes; A.N.B. and R.L. performed the chemical synthesis; M.L.B. and R.L. performed dye characterization and mechanistic studies; M.L.B. and M.R. performed labelling, microscopy, and data analysis; E.D. performed the culture, labelling, microscopy, and data analysis of the primary neurons; S.W.H. directed and supervised the investigations; all the authors discussed the results and commented on the manuscript.

