## Supplementary information for "A general design of caging-group free photoactivatable fluorophores for live-cell nanoscopy"


***This section includes:***


- Supplementary Tables — Page 2
- Supplementary Figures — Page 6
- Supplementary Methods — Page 23
- Synthesis and Characterization — Page 34
- References — Page 78
- NMR Spectra — Page 80

### SUPPLEMENTARY TABLES

#### Supplementary Table 1. Photophysical properties of PaX 1–6.

| Compound | $\Phi_{Act}$[a] | $\Phi_{Act} \times \varepsilon^{405nm}$ |
|---|---|---|
| | ***PaX → Closed-Form.*** | |
| **1** | 9.9 ×10$^{-3}$ | 234 |
| **2** | 2.5 ×10$^{-2}$ | 591 |
| **3** | 2.5 ×10$^{-2}$ | 556 |
| **4** | 5.6 ×10$^{-2}$ | 1410 |
| **5** | 2.1 ×10$^{-2}$ | 464 |
| **6** | 1.7 ×10$^{-2}$ | 383 |

[a] All measurements performed in phosphate buffer (100 mM, pH = 7.0) irradiation at 405 nm

#### Supplementary Table 2. Photophysical properties of PaX 9–12.

| Compound | $\Phi_{Act}$[a] | $\Phi_{Act} \times \varepsilon^{405nm}$ | $\lambda_{Abs}^{max}$ (nm) / $\varepsilon^{max}$ (M$^{-1}$cm$^{-1}$) | | $\lambda_{Em}^{max}$ | $\Phi_{Fluo}$ | $\tau_{Fluo}$ (ns) |
|---|---|---|---|---|---|---|---|
| | ***PaX → Closed-Form*** | | ***PaX*** | ***Closed-Form*** | | | |
| **9** (PaX$_{480}$) | 1.0 ×10$^{-3}$ | 42 | 399 / 3.2 ×10$^{4}$ | 480 / 4.5 ×10$^{4}$ | 514 | 0.22 | 0.86 |
| **10** (PaX$_{525}$) | 3.7 ×10$^{-2}$ | 1030 | 408 / 2.9 ×10$^{4}$ | 524 / 6.1 ×10$^{4}$ | 564 | 0.35 | 1.84 |
| **11** (PaX$_{560}$) | 7.9 ×10$^{-2}$ | 1930 | 414 / 2.6 ×10$^{4}$ | 558 / 7.0 ×10$^{4}$ | 596 | 0.48 | 2.10 |
| **12** (PaX+$_{560}$) | 3.1 ×10$^{-2}$ | 647 | 405 / 2.1 ×10$^{4}$ | 560 / 6.6 ×10$^{4}$ | 598 | 0.92 | 3.55 |

[a] All measurements performed in phosphate buffer (100 mM) pH = 7.0; a) irradiation at 405 nm

**Supplementary Table 3. Confocal and STED imaging parameters.**

| Figure | Micro-scope | Compound | Photoactivation | Excitation | | Depletion | | Detection windows (nm) | Pixel size (nm) | Dwell time (μs) | Line reps. |
|---|---|---|---|---|---|---|---|---|---|---|---|
| | | | | Laser (nm) | Power (μW) | Laser (nm) | Power (mW) | | | | |
| Fig. 2b | 2 | PaX **13** (Confocal) | WF (AHF, DAPI) | 561 | ~4.5 | - | - | 571–650 | 20 | 20 | 1x1 |
| | | PaX **13** (STED) | WF (AHF, DAPI) | 561 | ~6.1 | 655 | ~17 | 571–650 | 20 | 20 | 1x2 |
| Fig. 2c | 2 | STAR GREEN (Confocal) | - | 518 | ~6.5 | - | - | 535-560 | 30 | 8 | 1x2 |
| | | PaX **15** (Confocal) | WF (AHF, EGPF); 518 nm (~6.5 μW) | 561 | ~0.3 | - | - | 580–602 | 30 | 8 | 1x2 |
| | | PaX **15** (STED) | WF (AHF, EGPF); 518 nm (~6.5 μW) | 561 | ~2.8 | 655 | ~29 | 580–602 | 30 | 8 | 8x2 |
| Fig. 3b (Mito) | 1 | PaX **19** (Confocal) | 405 nm (13 μW) | 561 | ~4.9 | - | - | 571–630 | 70 | 20 | 1x2 |
| | | MitoTracker DR (Confocal) | - | 640 | ~0.38 | - | - | 650–757 | 70 | 20 | 1x2 |
| Fig. 3b (Lyso) | 1 | PaX **20** (Confocal) | 405 nm (13 μW) | 561 | ~11 | - | - | 571–630 | 70 | 20 | 1x2 |
| | | LysoTracker DR | - | 640 | ~1.8 | - | - | 650–757 | 70 | 20 | 1x2 |
| Fig. 3c | 2 | PaX **20** (Confocal) | 810 nm (2PA) | 518 | ~6.6 | - | - | 575–700 | 70 | 20 | 2x1 |
| Fig. 3e | 2 | PaX **20** (Confocal) | WF (AHF, DAPI) | 561 | ~2.4 | - | - | 571–650 | 20 | 20 | 2x1 |
| | | PaX **20** (STED) | WF (AHF, DAPI) | 561 | ~3.2 | 655 | ~14 | 571–650 | 20 | 20 | 2x3 |
| Fig. 4a–d | 1 | PaX **21** (Confocal) | 405 nm (13 μW) | 561 | ~4.9 | - | - | 571–750 | 70 | 40 | 1x2 |
| | | AL-560 (Confocal) | - | 561 | ~4.9 | - | - | 571–750 | 70 | 40 | 1x2 |
| Supp. Fig. 7b | 2 | PaX **17** (Confocal) | WF (AHF, DAPI) | 518 | ~7.6 | - | - | 540–650 | 20 | 20 | 2x1 |
| | | PaX **17** (STED) | WF (AHF, DAPI) | 518 | ~7.6 | 655 | ~18 | 540–650 | 20 | 20 | 2x3 |
| Supp. Fig. 7b | 2 | PaX **18** (Confocal) | WF (AHF, DAPI) | 561 | ~6.6 | - | - | 571–650 | 20 | 20 | 2x1 |
| | | PaX **18** (STED) | WF (AHF, DAPI) | 561 | ~15 | 655 | ~27 | 571–650 | 20 | 20 | 2x3 |
| Supp. Fig. 9a–d | 1 | PaX **16** (Confocal) | 405 nm (45 μW) | 485 | ~5.7 | - | - | 500–551 | 70 | 15 | 2x1 |
| | | PaX **13** (Confocal) | 405 nm (13 μW) | 561 | ~4.9 | - | - | 571–691 | 70 | 15 | 2x1 |

| | | | | | | | | | | | |
|---|---|---|---|---|---|---|---|---|---|---|---|
| Supp.<br>Fig. 9e | 1 | PaX **16** (Confocal)<br>PaX **17** (Confocal) | 405 nm (13 μW)<br>355 nm (0.14 μW) | 485<br>561 | ~5.7<br>~11 | -<br>- | -<br>- | 495–515<br>580–650 | 70<br>70 | 40<br>40 | 4x1<br>4x1 |
| Supp.<br>Fig. 13a | 1 | PaX **23** (Confocal)<br>PaX **19** (Confocal)<br>Hoechst (Confocal) | 405 nm (45 μW)<br>405 nm (13 μW)<br>- | 485<br>561<br>355 | ~5.7<br>~4.9<br>~0.29 | -<br>-<br>- | -<br>-<br>- | 495–551<br>571–691<br>400–475 | 70<br>70<br>70 | 15<br>15<br>15 | 2x1<br>2x1<br>2x1 |
| Supp.<br>Fig. 13b | 1 | PaX **23** (Confocal)<br>PaX **19** (Confocal)<br>Hoechst (Confocal) | 405 nm (45 μW)<br>405 nm (13 μW)<br>- | 485<br>561<br>355 | ~8.9<br>~8.2<br>~0.29 | -<br>-<br>- | -<br>-<br>- | 495–551<br>571–691<br>400–475 | 70<br>70<br>70 | 20<br>20<br>20 | 2x1<br>2x1<br>2x1 |

**Supplementary Table 4. PALM imaging parameters.**

| Figure | Compound | EM gain | Exposure Time (ms) | Excitation | | Mean Uncertainty (nm) | Total Frames | Mean $N_{photons}$ Per Localization |
|---|---|---|---|---|---|---|---|---|
| | | | | Laser (nm) | Power (mW) | | | |
| Fig. 2b | PaX **13** | 200 | 15 | 560 | 120 | 17 | 23k | 1300 |
| Fig. 2d | PaX **14** | 50 | 20 | 560 | 150 | 9 | 25k | 3300 |
| Fig. 2e | PaX **14** | 200 | 20 | 560 | 150 | 9 | 30k | 3500 |
| Fig. 5a | PaX **22** | 50 | 20 | 560 | 300 | 12 | 4k | 2500 |
| Fig. 5b | PaX **21** | 50 | 20 | 560 | 300 | 10 | 2k | 3700 |
| Supp. Fig. 7c | PaX **16** | 50 | 20 | 473 | 120 | 15 | 5k | 1300 |
| Supp. Fig. 7c | PaX **17** | 50 | 20 | 532 | 300 | 14 | 10k | 1900 |
| Supp. Fig. 7c | PaX **18** | 50 | 20 | 560 | 300 | 10 | 20k | 5500 |
| Supp. Fig. 9f | PaX **13** | 50 | 20 | 560 | 300 | 11 | 7k | 5400 |
| Supp. Fig. 9f | PaX **16** | 50 | 20 | 473 | 150 | 19 | 4k | 1100 |
| Supp. Fig. 14a | PaX **22** | 200 | 20 | 560 | 150 | 14 | 18k | 5200 |
| Supp. Fig. 14b | PaX **21** | 200 | 20 | 560 | 60 | 12 | 42k | 3500 |

### SUPPLEMENTARY FIGURES

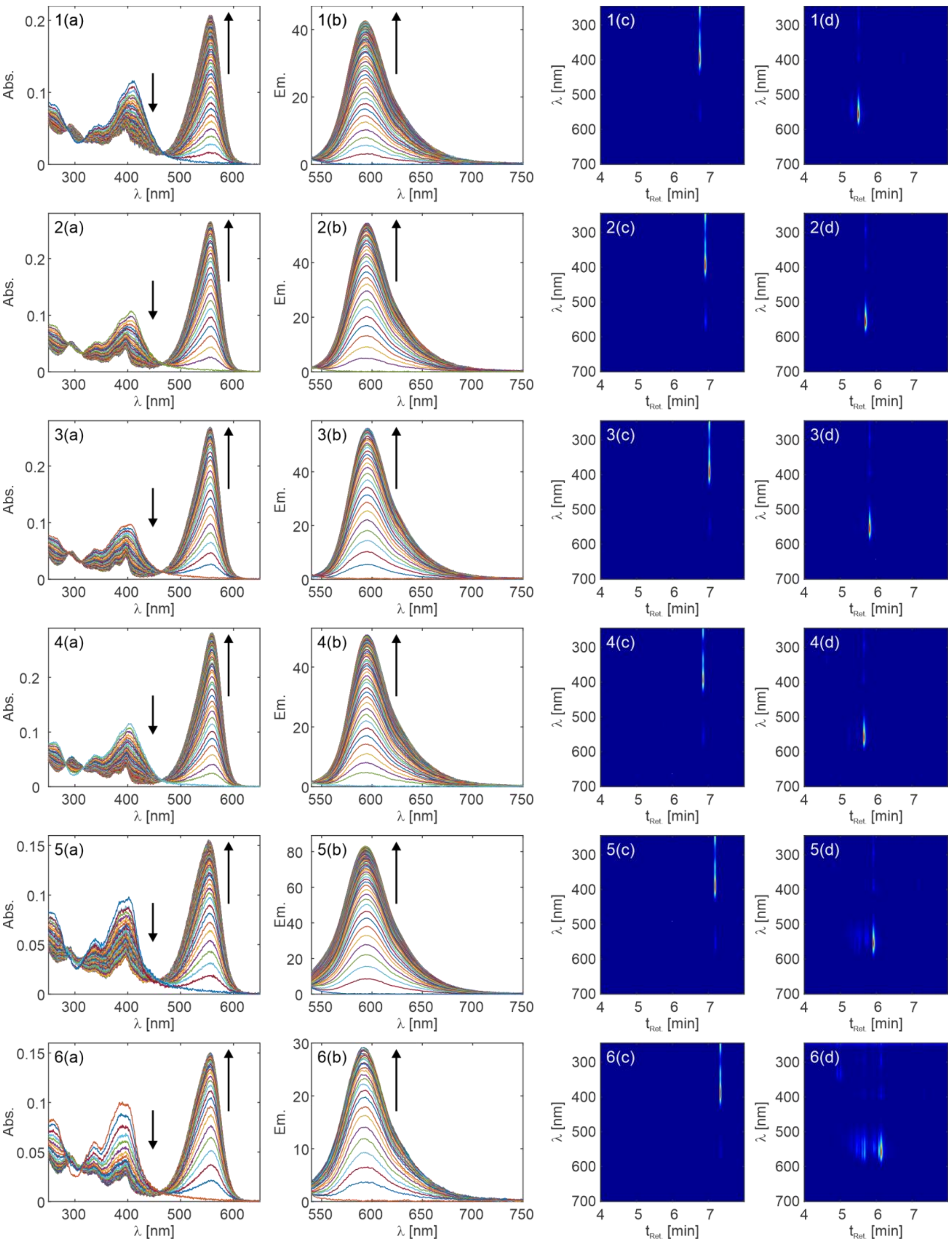

**Supplementary Fig. 1 | Effect of alkene substitution on photoactivation of Si-bridged PaX.** Temporal evolution of absorption (a) and fluorescence (b) spectra of compound **1**–**6** (top to bottom) irradiated in phosphate buffer (100 mM, pH = 7; $\lambda_{act}$ = 405 nm). The HPLC of the starting solutions (c) and irradiated solutions (d) are shown on the right.

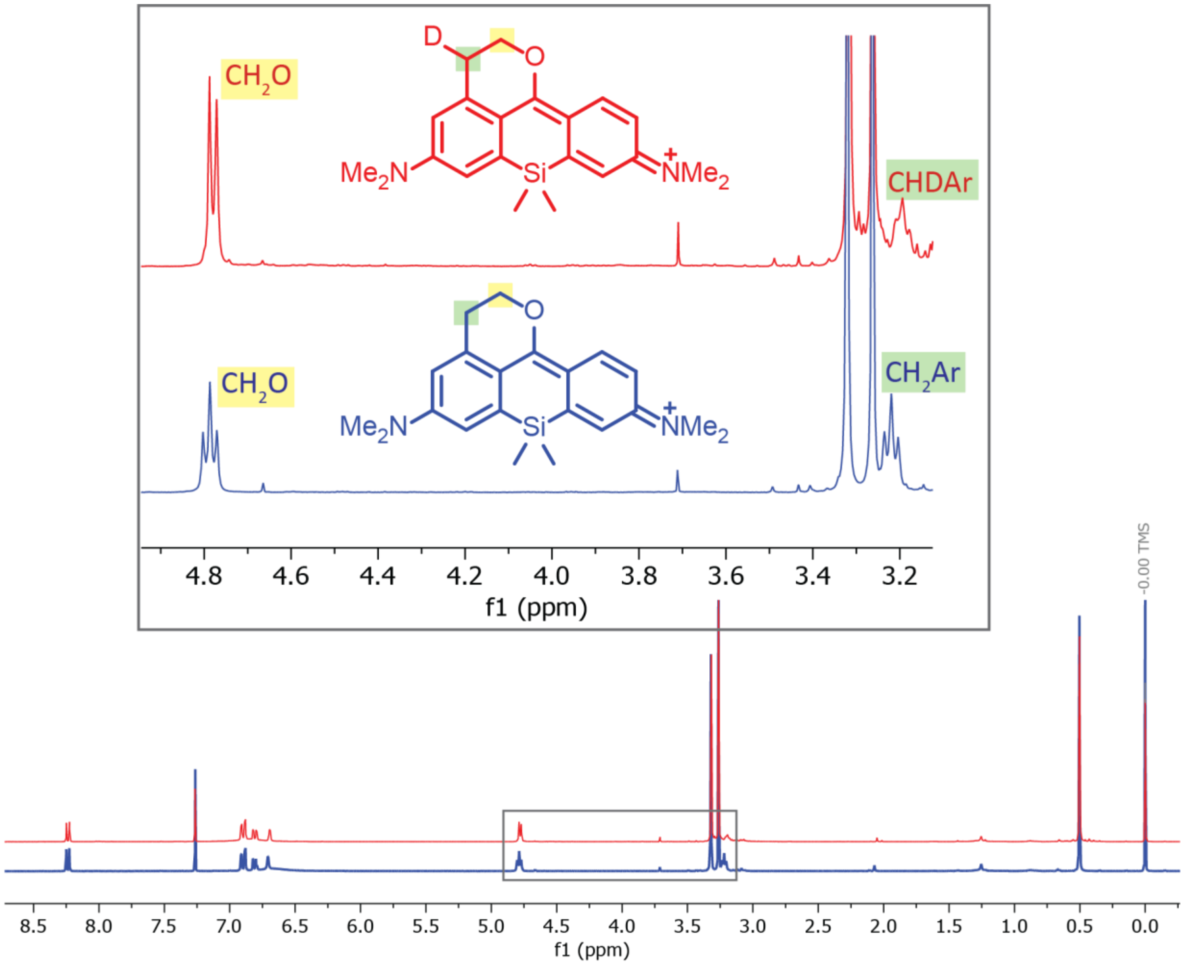


**Supplementary Fig. 2 | $^{1}$H NMR spectra of Compound 1-CF.** The compound **1** was irradiated with 405 nm LED in methanol-$d_4$ (red) and methanol (blue). All spectra were recorded in $CDCl_3$

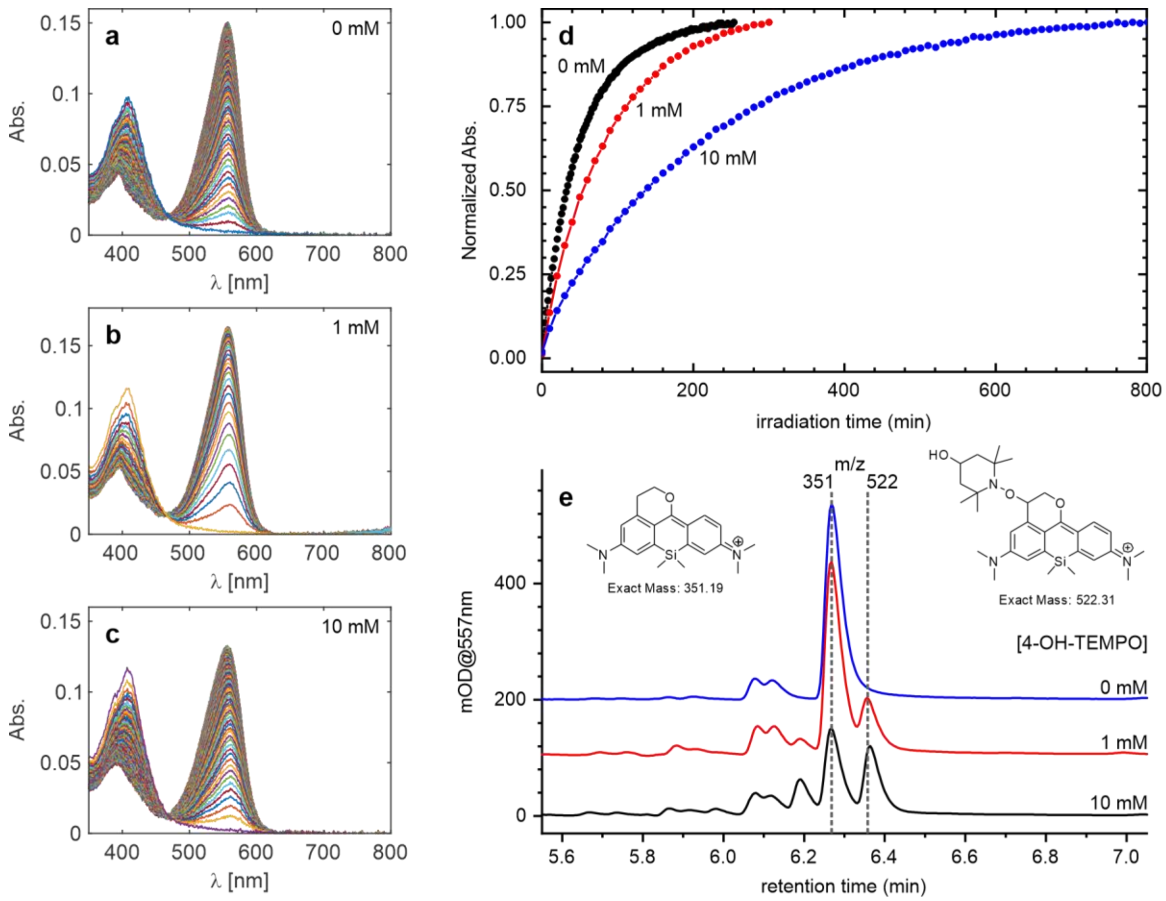


**Supplementary Fig. 3 | Effect of 4-hydroxy-TEMPO on the photoactivation rate of compound 1.** Temporal evolution of absorption spectra changes during photoactivation of **1** (4.8 µM) in the presence of 0 mM (**a**), 1 mM (**b**), and 10 mM (**c**) 4-hydroxy-TEMPO (methanol, $\lambda_{act}$ = 405 nm). **d**, Comparative photoactivation kinetics of **1** in methanol with different concentrations of 4-hydroxy-TEMPO ($\lambda_{act}$ = 405 nm). **e**, LC-MS chromatogram at 557 nm of the reaction mixtures of **1** photoactivated in methanol with different concentrations of 4-hydroxy-TEMPO, showing the formation of TEMPO-adducted pyronine dye. The anticipated product structure is shown.

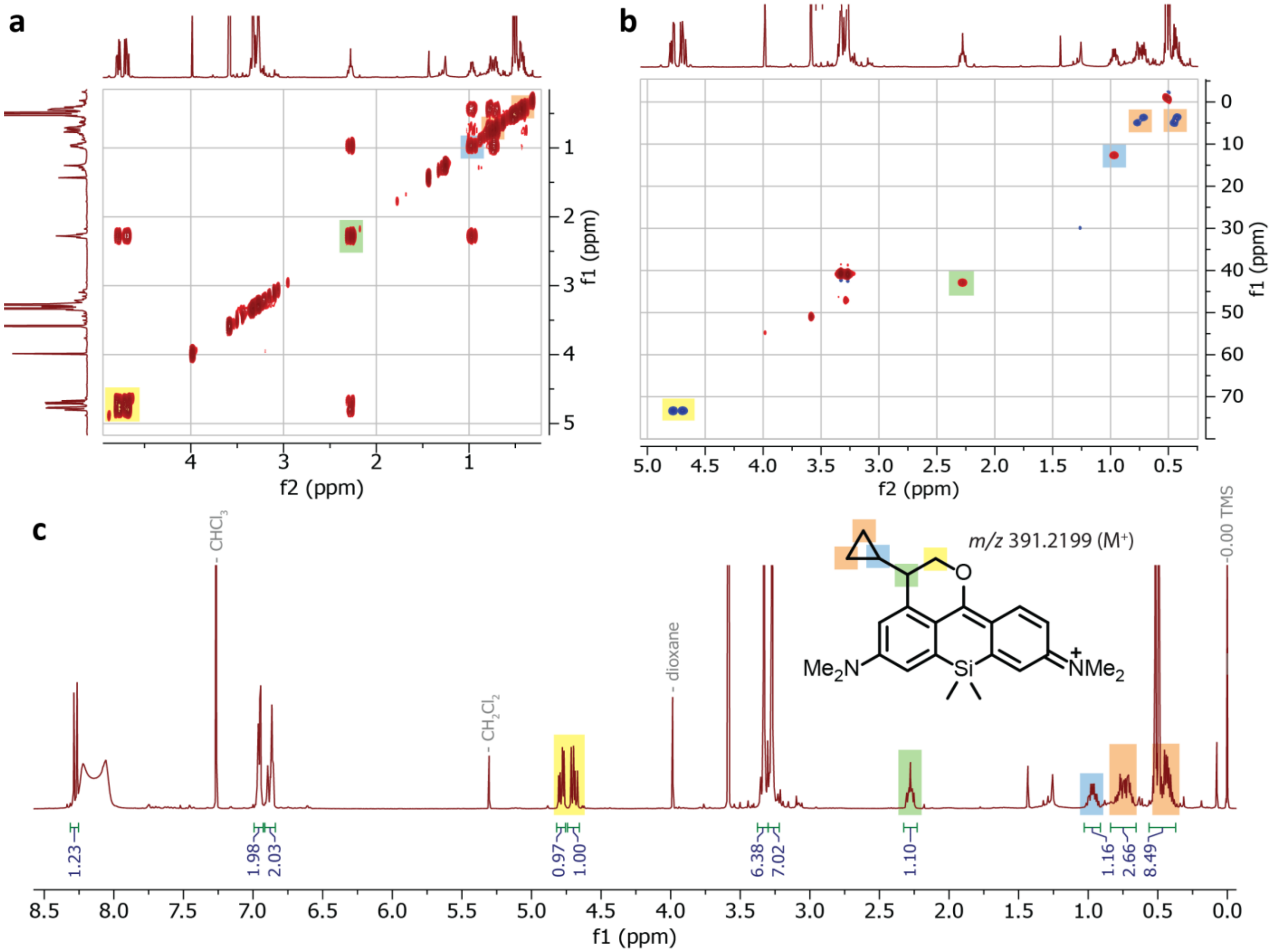


**Supplementary Fig. 4 | NMR spectra of Compound 7 upon photoactivation.** H-H COSY (**a**), HSQC (**b**) and $^1$H NMR (**c**) of **7-CF** prepared by photoactivation of **7** in methanol solution (crude reaction mixture). All spectra were recorded in $CDCl_3$ + 1% (v/v) TFA-*d*.

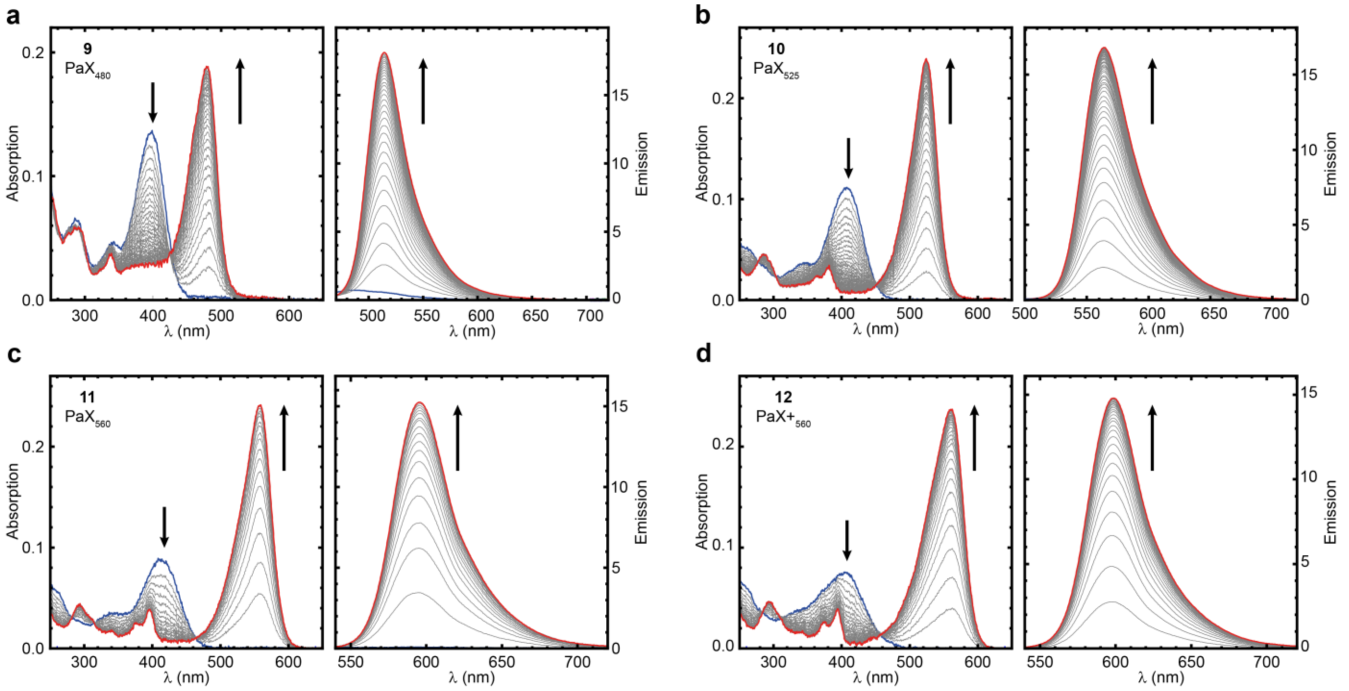


**Supplementary Fig. 5 | Absorption and emission spectrum of PaX derivatives. a**, Temporal evolution of absorption and fluorescence spectra changes during photoactivation of $PaX_{480}$ **9** (**a**), $PaX_{525}$ **10** (**b**), $PaX_{560}$ **11** (**c**), and $PaX+_{560}$ **12** (**d**) in phosphate buffer (100 mM, pH = 7; $\lambda_{act}$ = 405 nm). Initial spectra are shown in blue, and final spectra are shown in red.

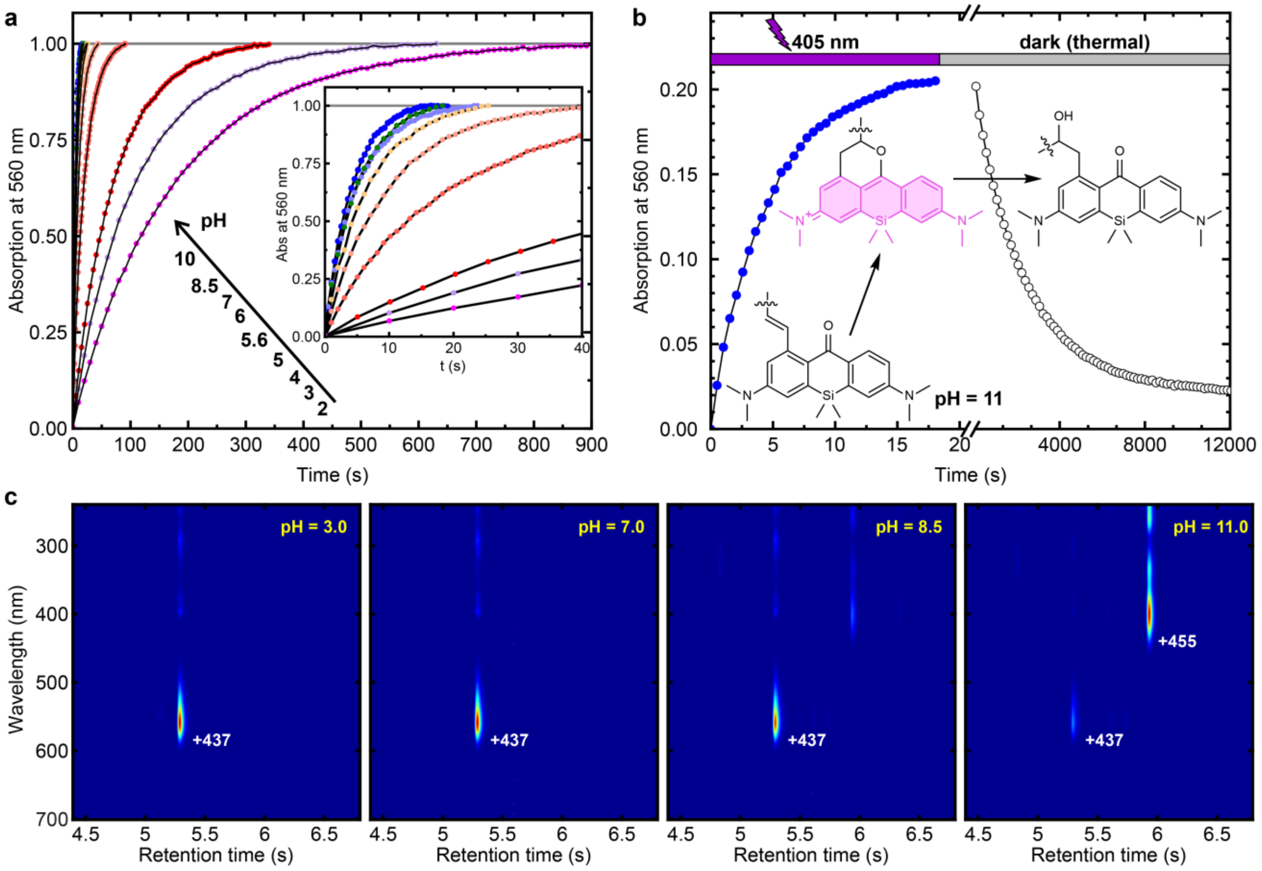


**Supplementary Fig. 6 | Effect of pH on the photoactivation rate of PaX 11. a**, Comparative photoactivation kinetics of PaX **11** in phosphate buffered saline at different pH, measured as the absorption change at 560 nm; photoactivation at 405 nm. **b**, Photoactivation of PaX **11** ($\lambda_{act}$ = 405 nm) and thermal hydrolysis of resulting **11-CF** (dark) in phosphate buffered saline at pH = 11. **c**, LC-MS chromatograms of the reaction mixtures of PaX **11** photoactivated at different pH, showing the formation of the hydrolysis product at pH 8.5 and above.

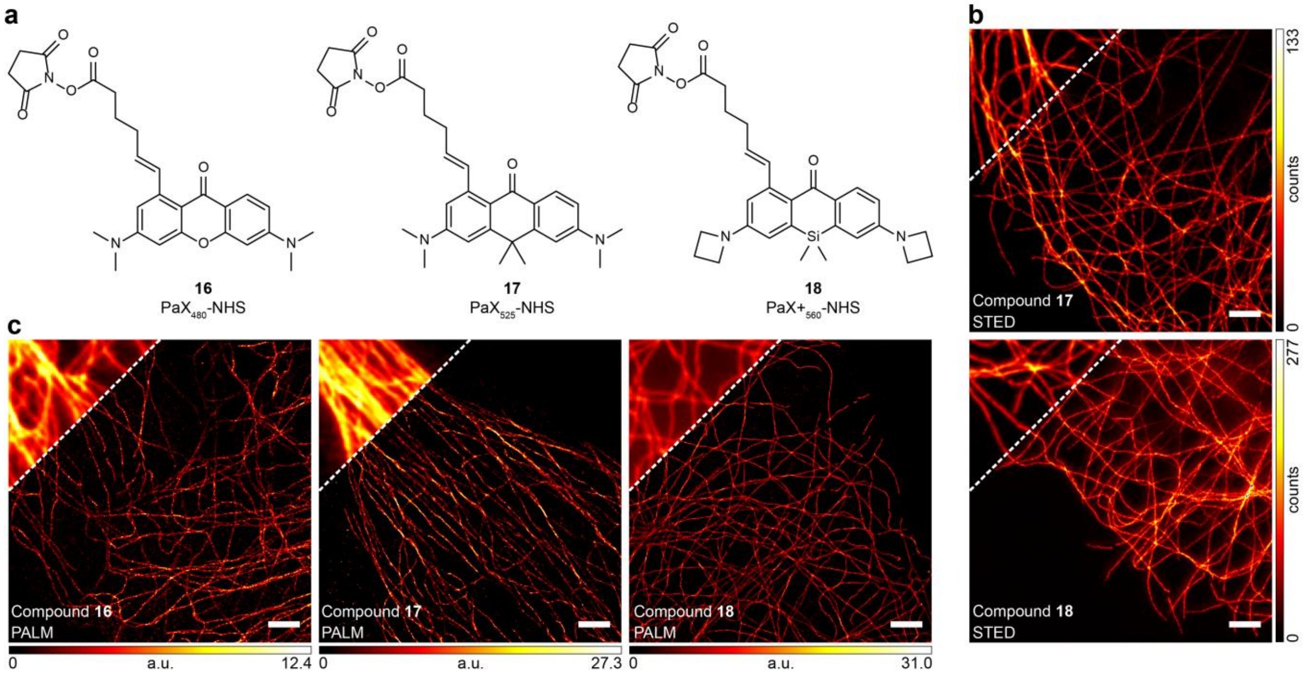


**Supplementary Fig. 7 | PaX derivatives for bioconjugation to amines. a**, Chemical structures of PaX derivatives for bioconjugation to amines. **b**, STED image of microtubules in COS-7 cells labelled by indirect immunofluorescence with a secondary antibody bearing **17** and **18**. Preactivation of PaX for STED imaging was achieved with widefield illumination (AHF, DAPI filter set F46-816). **c**, PALM images of microtubules in COS-7 cells labelled by indirect immunofluorescence with a secondary antibody bearing compounds **16**–**18.** Scale bars: 2 µm.

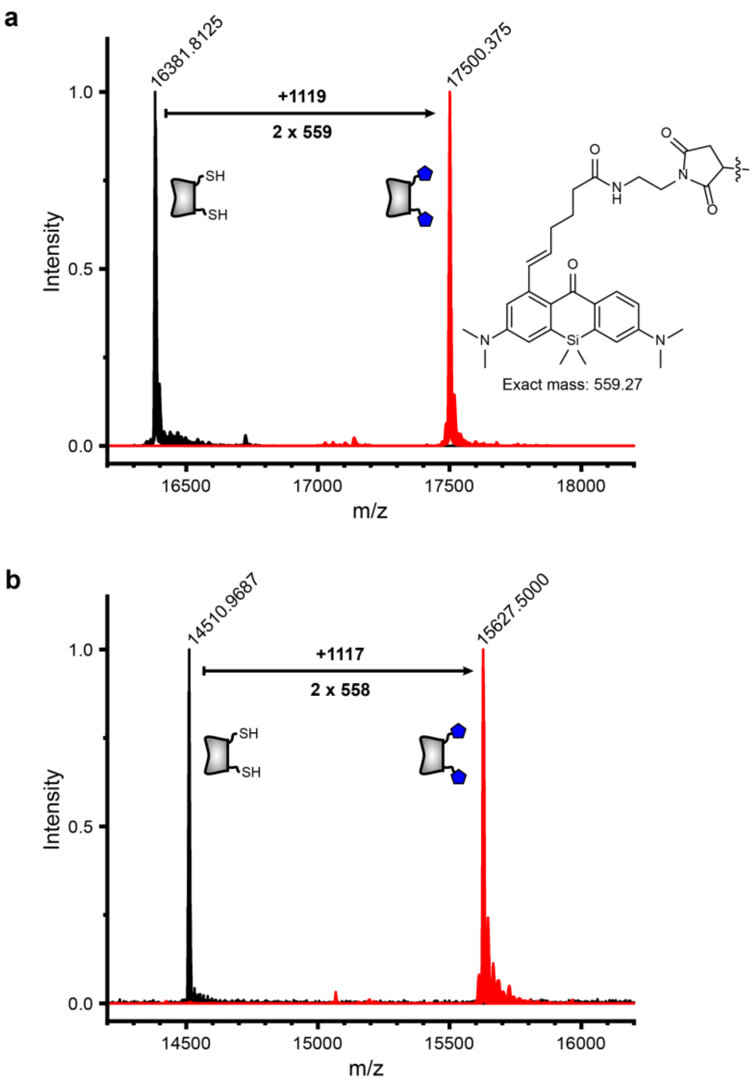


**Supplementary Fig. 8 | Mass spec of nanobody-dye conjugates.** Electrospray ionization mass spectrometry of FluoTag-X2 anti-Rabbit IgG nanobody clones 8H10 (**a**) and 10E10 (**b**), each containing two cysteine residues. The spectra of the unlabeled nanobody (black) and the labeled one (red) are presented, and the molecular mass of the dye-adduct (**14**) is indicated.

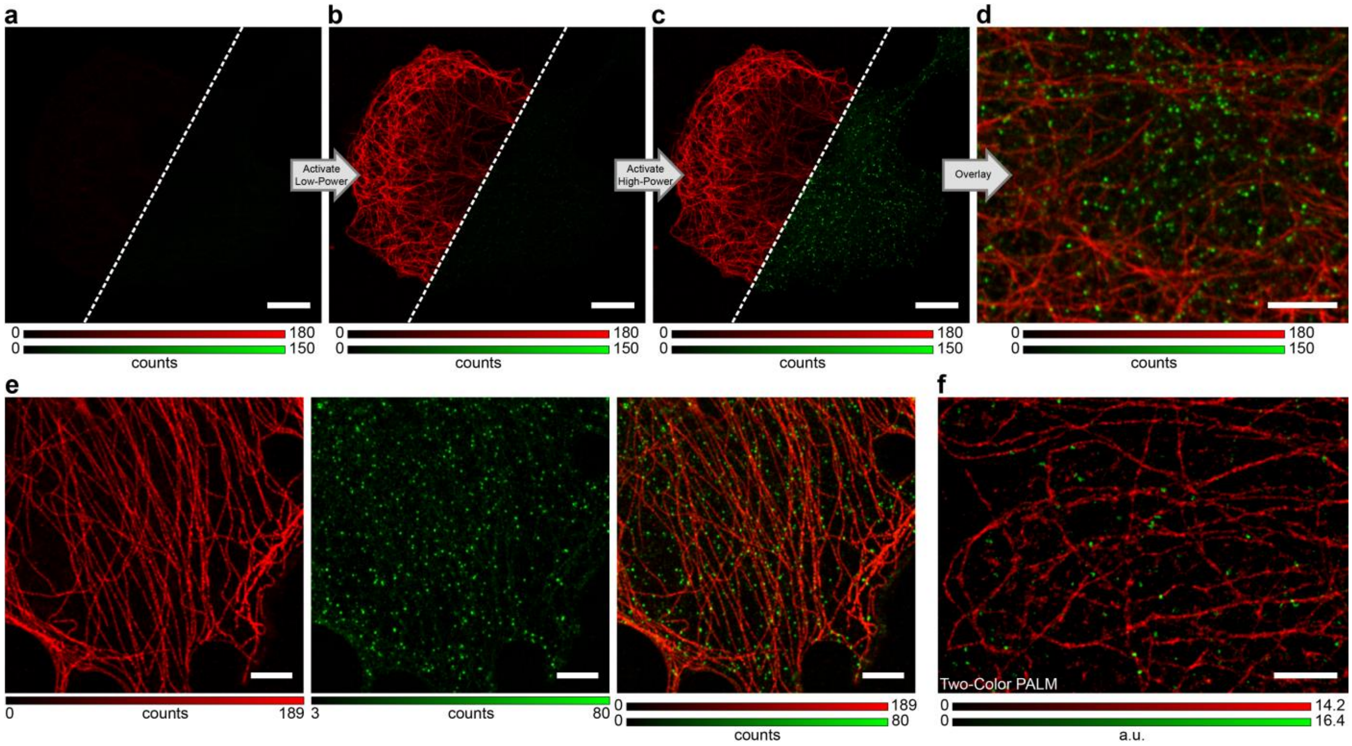


**Supplementary Fig. 9 | Selective activation of PaX by attenuation of light-dose. a–d**, Confocal images of microtubules (red) and clathrin-coated pits (green) in COS-7 cells labelled with **13** and **16**, respectively. Images were acquired before (**a**) and after a low photoactivation dose sufficient to activate **13** (**b**), and after high-power photoactivation dose to activate **16** (**c**). **d**, magnified region of the same cell. Photoactivation was carried out with 405 nm light. **e**, Confocal images of microtubules (red) and clathrin-coated pits (green) in COS-7 cells labelled cells labelled with **17** and **16**, respectively, after selective photoactivation (with 405 nm light) and imaging. **f**, Two-color PALM image of microtubules (red) and clathrin-coated pits (green) in COS-7 cells labelled with **13** and **16**, respectively. Scale bars, 10 µm (a, b, c), 5 µm (d, e), and 2 µm (f).

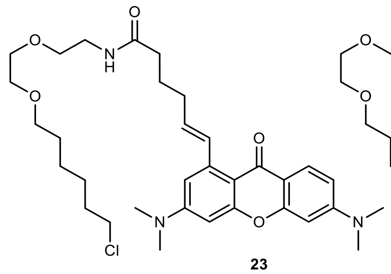

**23**
$PaX_{480}$-Halo

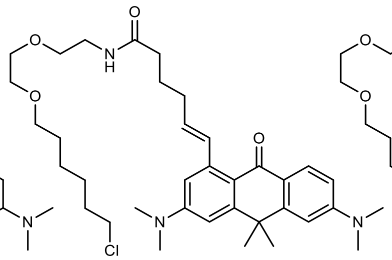

**24**
$PaX_{525}$-Halo

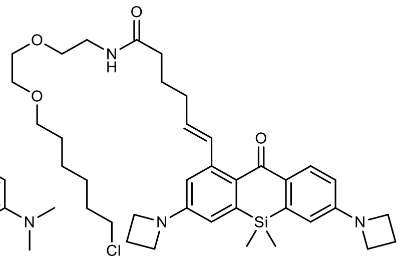

**25**
$PaX+_{560}$-Halo

**Supplementary Fig. 10 | Chemical structures of PaX-HaloTag ligand derivatives (23-25).**

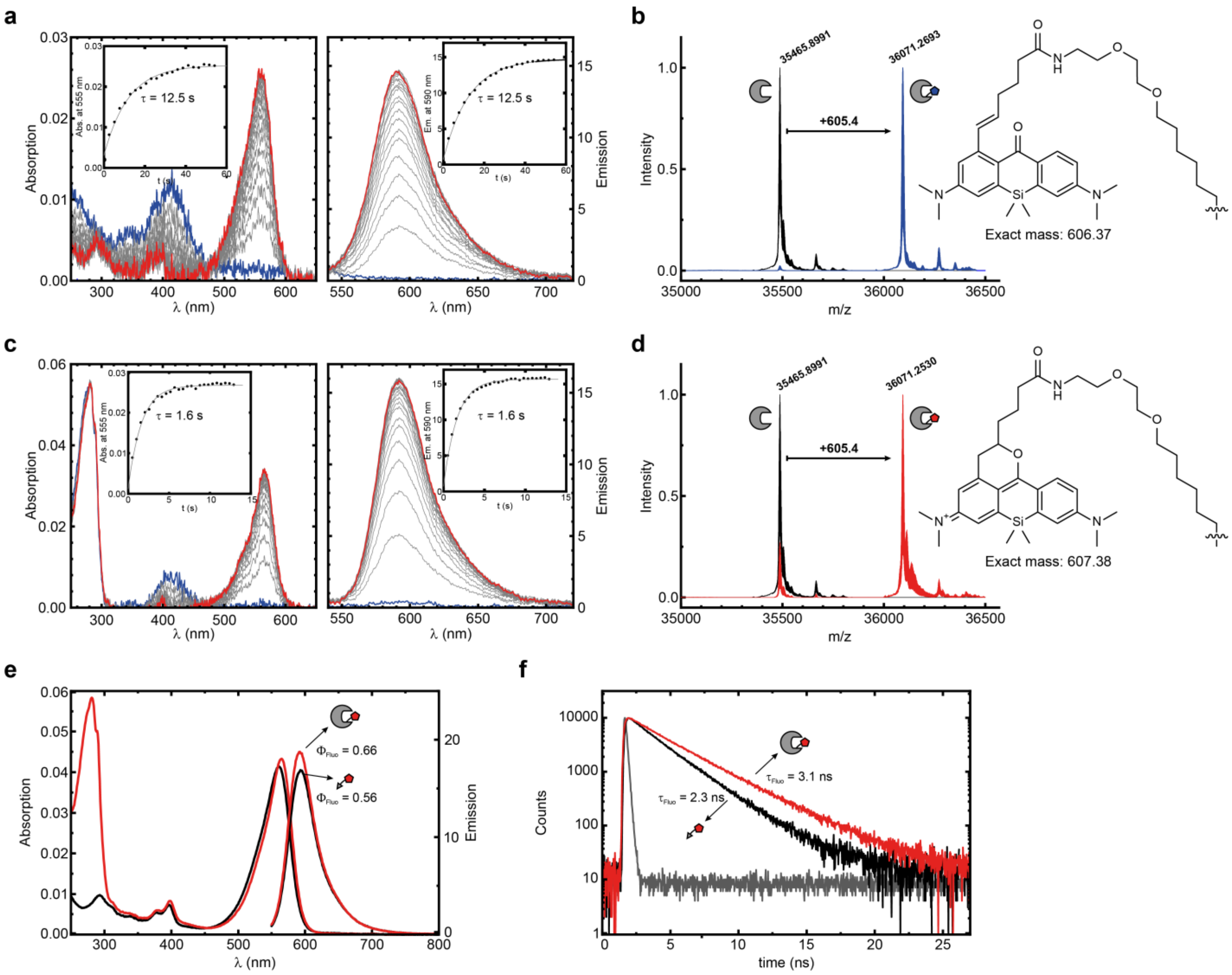


**Supplementary Fig. 11 | Characterization of PaX$_{560}$-Halo (21) with HaloTag protein. a**, Temporal evolution of absorption and fluorescence spectra during photoactivation of PaX$_{560}$-Halo (**21**) in phosphate buffer (100 mM, pH = 7; $\lambda_{act}$ = 405 nm). Initial spectra shown in blue, and final spectra are shown in red. **b**, Electrospray ionization mass spectrometry of HaloTag7 labelled with compound **21**. The spectra of the unlabeled HaloTag (black) and the labeled one (blue) are presented, and the molecular mass of the PaX-adduct (**21**) is indicated. **c**, Temporal evolution of absorption and fluorescence spectra during photoactivation of HaloTag7 protein labelled with compound **21**, in phosphate buffer (100 mM, pH = 7; $\lambda_{act}$ = 405 nm). Initial spectra shown in blue, and final spectra are shown in red. **d**, Electrospray ionization mass spectrometry of HaloTag7 labelled with compound **21** after photoactivation to **21-CF**. The spectra of the unlabeled HaloTag7 (black) and the labeled one (red) are presented, and the molecular mass of the dye-adduct (**21-CF**) is indicated. **e**, Absorption and emission spectra of **21-CF** before (black) and after (red) binding to HaloTag7. The quantum yield of fluorescent is indicated. **f**, Fluorescence lifetime of **21-CF** before (black) and after (red) binding to HaloTag7.

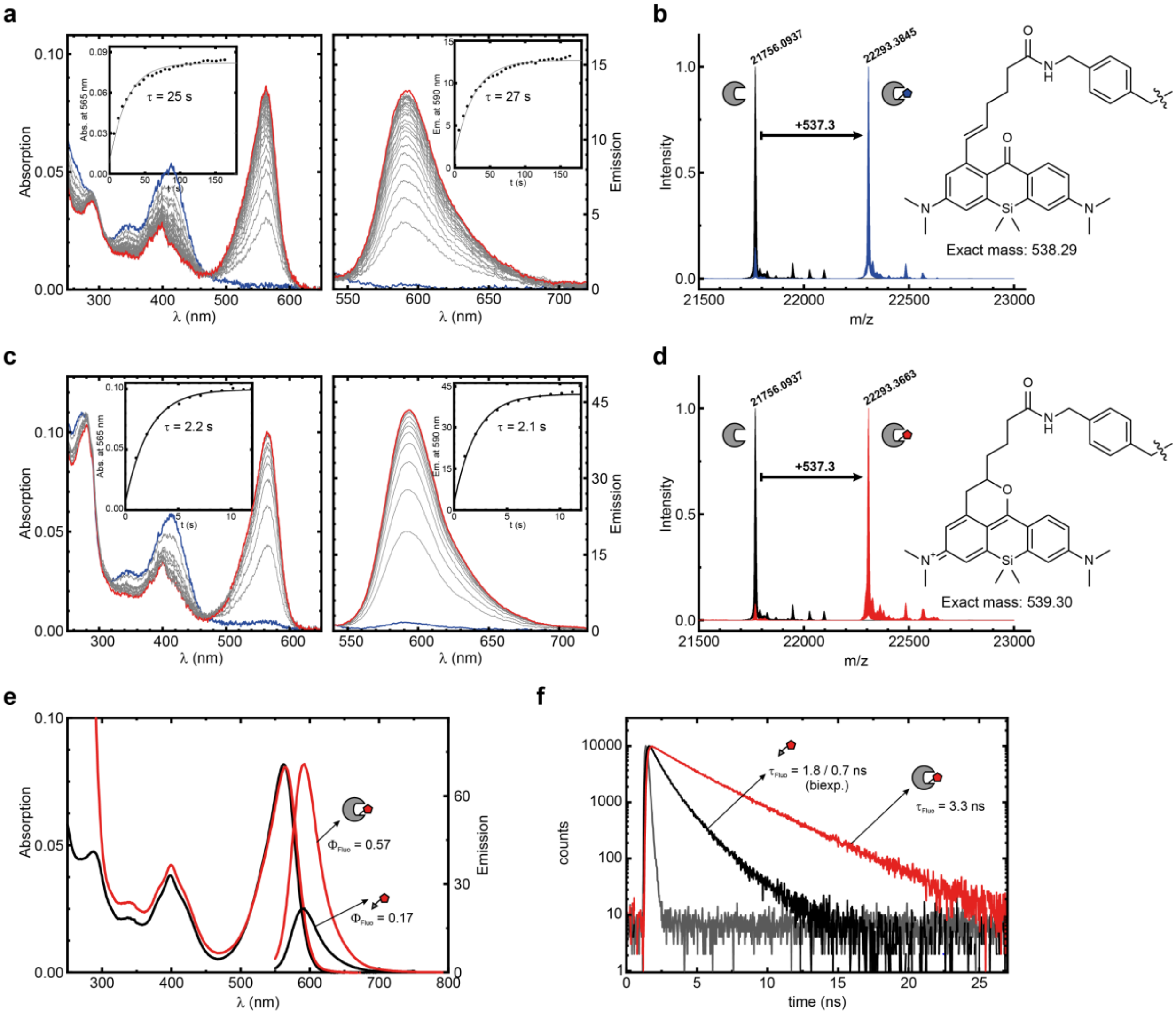


**Supplementary Fig. 12 | Characterization of PaX$_{560}$-SNAP (22) with SNAP-tag protein. a**, Temporal evolution of absorption and fluorescence spectra of **22** during photoactivation of compound **22** in phosphate buffer (100 mM, pH = 7; $\lambda_{act}$ = 405 nm). Initial spectra shown in blue, and final spectra are shown in red. **b**, Electrospray ionization mass spectrometry of SNAP-tag labelled with compound **22**. The spectra of the unlabelled SNAP-tag (black) and the labeled one (blue) are presented, and the molecular mass of the PaX-adduct (**22**) is indicated. **c**, Temporal evolution of absorption and fluorescence spectra during photoactivation of SNAP-tag protein labelled with compound **22**, in phosphate buffer (100 mM, pH = 7; $\lambda_{act}$ = 405 nm). Initial spectra shown in blue, and final spectra are shown in red. **d**, Electrospray ionization mass spectrometry of SNAP-tag labelled with compound **22** after photoactivation to **22-CF**. The spectra of the unlabeled SNAP-tag (black) and the labeled one (red) are presented, and the molecular mass of the dye-adduct (**22-CF**) is indicated. **e**, Absorption and emission spectra of **22-CF** before (black) and after (red) binding to SNAP-tag. The quantum yield of fluorescent is indicated. **f**, Fluorescence lifetime of **22-CF** before (black) and after (red) binding to SNAP-tag.

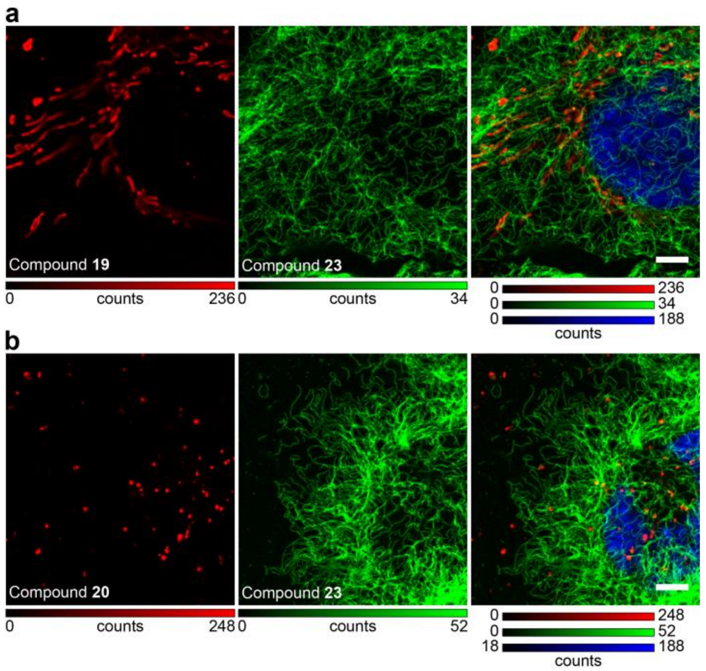


**Supplementary Fig. 13 | Selective activation of PaX dyes in living cells. a**, Confocal images of COS-7 cells stably expressing a vimentin-HaloTag construct co-labelled with compound **19** (red, specific for mitochondria), compound **23** (green, specific for HaloTag) and Hoechst (blue). Selective activation of PaX was conducted by first low-power photoactivation selective for **19** and subsequent, higher-power photoactivation to activation compound **23**. **b**, Confocal images of COS-7 cells stably expressing a vimentin-HaloTag construct, co-labelled with compound **20** (red, specific for lysosomes), compound **20** (green, specific for HaloTag) and Hoechst (blue). Selective activation of PaX was conducted by first low-power photoactivation selective for **20** and subsequent, higher-power photoactivation to activation compound **23**. Scale bars, 5 µm.

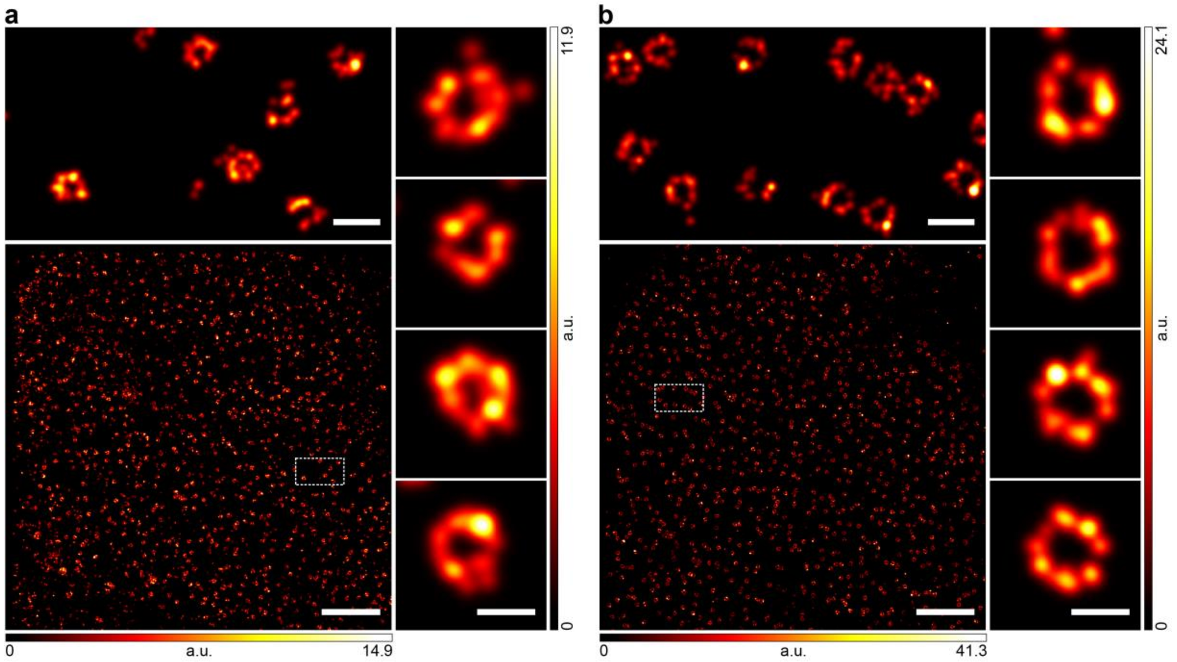


**Supplementary Fig. 14 | PALM image of NPCs labelled with live-cell labels imaged after fixation. a,** PALM image of U2OS stably expressing a NUP107-SNAP-tag construct labelled with **22** followed by fixation. Top: Magnified region marked in overview image. Right column: Magnified individual NPCs. **b**, PALM image of U2OS cells stably expressing a NUP96-HaloTag construct labelled with **21** followed by fixation. Top: Magnified region marked in overview image. Right column: Magnified individual NPCs. Scale bars: 2 µm (a,b), 200 nm (a,b insets), 100 nm (a,b right columns).

### SUPPLEMENTARY METHODS

#### General experimental information and synthesis

All chemical reagents (TCI, Sigma-Aldrich, Alfa Aesar) and dry solvents for synthesis (over molecular sieves, AcroSeal package, Acros Organics) were purchased from reputable suppliers and were used as received without further purification. The products were lyophilized from a suitable solvent system using Alpha 2-4 LDplus freeze-dryer (Martin Christ Gefriertrocknungsanlagen GmbH).

#### Thin Layer Chromatography

Normal phase TLC was performed on silica gel 60 $F_{254}$ (Merck Millipore, Germany). For TLC on reversed phase silica gel 60 RP-18 $F_{254}$s (Merck Millipore) was used. Compounds were detected by exposing TLC plates to UV-light (254 or 366 nm) or heating with vanillin stain (6 g vanillin and 1.5 mL conc. $H_2SO_4$ in 100 mL ethanol), unless indicated otherwise.

#### Flash Chromatography

Preparative flash chromatography was performed with an automated Isolera One system with Spektra package (Biotage AG) using commercially available cartridges of suitable size as indicated (RediSep Rf series from Teledyne ISCO, Puriflash Silica HP 30µm series from Interchim).

#### Nuclear Magnetic Resonance (NMR)

NMR spectra were recorded on a Bruker DPX 400 spectrometer. All spectra are referenced to tetramethylsilane as an internal standard (δ = 0.00 ppm). Multiplicities of the signals are described as follows: s = singlet, d = doublet, t = triplet, q = quartet, m = multiplet or overlap of non-equivalent resonances; br = broad signal. Coupling constants $^{n}J_{X\text{-}Y}$ are given in Hz, where n is the number of bonds between the coupled nuclei X and Y ($J_{H\text{-}H}$ are always listed as *J* without indices).

**Mass-Spectrometry (MS)**

Low resolution mass spectra (100 - 1500 *m/z*) with electro-spray ionization (ESI) were obtained on a Shimadzu LC-MS system described below. High resolution mass spectra (HRMS) were obtained on a maXis II ETD (Bruker) with electrospray ionization (ESI) at the Mass Spectrometry Core facility of the Max-Planck Institute for Medical Research (Heidelberg, Germany).

**High-Performance Liquid Chromatography (HPLC)**

Analytical liquid chromatography-mass spectrometry was performed on an LC-MS system (Shimadzu): 2x LC-20AD HPLC pumps with DGU-20A3R solvent degassing unit, SIL-20ACHT autosampler, CTO-20AC column oven, SPD-M30A diode array detector and CBM-20A communication bus module, integrated with CAMAG TLC-MS interface 2 and LCMS-2020 spectrometer with electrospray ionization (ESI, 100 – 1500 *m/z*). Analytical column: Hypersil GOLD 50×2.1 mm 1.9µm, standard conditions: sample volume 1-2 µL, solvent flow rate 0.5 mL/min, column temperature 30 °C. General method: isocratic 95:5 A:B over 2 min, then gradient 95:5 – 0:100 A:B over 5 min, then isocratic 0:100 A:B over 2 min; solvent A = water + 0.1% v/v $HCO_2H$, solvent B = acetonitrile + 0.1% v/v $HCO_2H$.

Preparative high-performance liquid chromatography was performed on a Büchi Reveleris Prep system using the suitable preparative columns and conditions as indicated for individual preparations. Method scouting was performed on a HPLC system (Shimadzu): 2x LC-20AD HPLC pumps with DGU-20A3R solvent degassing unit, CTO-20AC column oven equipped with a manual injector with a 20 µL sample loop, SPD-M20A diode array detector, RF-20A fluorescence detector and CBM-20A communication bus module; or on a Dionex Ultimate 3000 UPLC system: LPG-3400SD pump, WPS-3000SL autosampler, TCC-3000SD column compartment with 2× 7-port 6-position valves and DAD-3000RS diode array detector. The test runs were performed on analytical columns with matching phases (HPLC: Interchim 250×4.6 mm 10 µm C18HQ, Interchim 250×4.6 mm 5 µm PhC4, solvent flow rate 1.2 mL/min; UPLC: Interchim C18HQ or PhC4 75×2.1 mm 2.2 µm, ThermoFisher Hypersil GOLD 100×2.1 mm 1.9 µm, solvent flow rate 0.5 mL/min).

### Description of the photoreactor

For photoactivation of PaX dyes on a semipreparative scale, Photoreactor m1 (Penn PhD, USA) was used together with a custom home-built 405 nm LED unit. The LED unit consists of a high-power LZC-C0UB00-00U7 400-405 nm LED module (LED Engin, OSRAM Group, Germany), with two series of LEDs connected in parallel (optional parallel jumpers soldered on), mounted on a SA-LED-113E aluminium heat sink (Ohmite, USA) and driven with an LDD-1500H LED constant current power supply (MEAN WELL, Taiwan), set in a 3D-printed (PETG, signal white) enclosure. For a detailed description (bill of materials for electronic parts, schematic, board outline and production files, 3D models of enclosure and flask holders), see the attached Supplementary Compressed Archive File (.zip).

### Optical spectroscopy

Fluorescence quantum yield measurements were measured according to a relative method, using Fluorescein in NaOH (0.1 M), Rhodamine6G (Ethanol), and Sulforhodamine 101 (Ethanol) as references[3]. Absorption spectra for fluorescence quantum yield measurements were recorded with a Varian Cary 4000 UV-Vis double-beam spectrophotometer (Agilent Technologies). The emission spectra for fluorescence quantum yield measurements were recorded with a Varian Cary Eclipse fluorescence spectrophotometer (Agilent Technologies). The absorption and emission spectra were recorded in 3 mL quartz cells (optical path length 1 cm, model 119F-10-40 or model 111-10-40, Hellma Analytics). Fluorescence lifetimes were measured with a FluoTime 300 fluorescence lifetime spectrometer (PicoQuant). All measurements were performed in air-saturated solvents at ambient temperature.

### Confocal and STED (stimulated emission depletion) microscopy.

Confocal and STED images were acquired using two Abberior Expert Line (Abberior Instruments GmbH, Göttingen, Germany) fluorescence microscopes built on a motorized inverted microscope IX83 (Olympus, Tokyo, Japan). Microscope 1 is equipped with pulsed STED lasers at 595 nm and 775 nm shaped by Spatial Light Modulators (SLMs), and with 355 nm, 405 nm,

485 nm, 561 nm, and 640 nm excitation lasers, and a 100x/1.40 oil immersion objective lenses (Olympus). Microscope 2 is equipped with pulsed STED lasers at 655 nm and 775 nm, and with 520 nm, 561 nm, 640 nm, and multiphoton (Chameleon Vision II, Coherent, Santa Clara, USA) excitation lasers, and a 60x/1.42 oil immersion objective lens (Olympus). The multiphoton laser is tuneable in the 680 nm – 1080 nm range. Spectral detection is performed in both cases with avalanche photodiodes at spectral windows adjusted for each particular fluorophore.

Imaging and image processing was done with ImSpector software, and all images are displayed as raw data unless otherwise noted.

Particular imaging conditions are given in Supplementary Table 3.

**Superresolution single molecule localization microscopy (SMLM) / Photoactivated localization microscopy (PALM)**

Images were acquired on a custom-built setup[1], equipped with a 473 nm (500 mW), a 532 nm (1 W) and a 560 nm (1 W) laser for excitation, a 405 nm (300 mW) laser for activation, a back illuminated EMCCD camera (Andor iXon 897 / 512×512 sensor), and a Leica HCX PL APO CS 100x/1.46 oil lens. Emission light was separated from the excitation and activation light with proper combination of a dichroic mirror and a suppression filter for each excitation laser. A movable mirror was used to switch between wide field, highly inclined and laminated optical sheet (HILO) and total internal reflection fluorescence (TIRF) illumination modes. Images were acquired with a 10-20 ms exposure time, and actual excitation laser powers in the back focal plane of ca. 50-400 mW, depending on the dye and sample properties. The 405 nm activation laser was incorporated as 100-500 µs pulses, in between frames, with a power of 0.001-1 mW in the back focal plane.

Particular imaging conditions are given in Supplementary Table 4.

All images were analyzed and processed using the ThunderSTORM plugin[2] on ImageJ (version 1.52p). In brief, images were filtered with a wavelet filer (B-spline order 3, scale 2.0), approximate localization of the molecules was performed with a local maximum method (peak intensity threshold of ca. 2-3 standard deviations; connectivity 8-neighbourhood), and sub-pixel localization of the molecules was performed with a maximum likelihood fitting method (PSF integrated method, fitting radius 3 px, initial sigma 1.3–1.6). Post-processing was performed with the same plug-in. Data was drift-corrected based on the cross-correlation method, merged, and a density filter was applied (distance radius ca. 50 nm, minimum neighbors in the radius ca. 2-5). Filtering and rendering of localizations was done using custom-built MatLab (version R2007a) routines. Final images were produced using an amplitude-normalized Gaussian rendering method with a fixed sigma value corresponding to the average localization uncertainty (Supplementary Table 4) and a pixel-size of 2 nm for the rendered image.

**Photolysis of PaX compounds, free and protein-bound HaloTag and SNAP-tag PaX ligands, and chemometric analysis of the photoactivation and photobleaching reaction kinetics**

Solutions in phosphate buffer (100 mM, pH = 7.0; 1.66 µg $mL^{-1}$) or methanol were irradiated in a previously described[4] home-built setup with a 405 nm LED source (M405L3, Thorlabs Inc.) in combination with a bandpass (10 nm) filter (FB405-10, Thorlabs Inc.). During the irradiation, samples were maintained at 20 °C and continuously stirred with a Peltier-based temperature-controlled cuvette holder (Luma 40, Quantum Northwest, Inc.). The absorption and emission of irradiated solutions was monitored at desired irradiation intervals with a fiber-based spectrometer (Flame-S-UV-Vis-ES, Ocean Insight). For absorption measurements, a deuterium and tungsten halogen source was used for illumination (DH-2000-BAL, Ocean Insight), and for fluorescence excitation was performed in a 90° configuration with an LED source (Thorlabs Inc.) in combination with an appropriate bandpass (10 nm) filter emitting at a wavelength suitable for each compound. Data collection and analysis was performed with custom-made routines in Matlab. Samples for LCMS or ESI-MS analysis were taken before and after photolysis was performed.

To prepare protein-bound HaloTag and SNAP-tag PaX ligands, the solutions of PaX incubated with protein (~1.1 eqs.) for one hour to ensure complete labelling, before photolysis experiments.

### Conjugation of antibodies via amino groups

Amino-reactive NHS-esters of PaX dyes were coupled to secondary antibodies (Jackson ImmunoResearch Europe Ltd. product nos.: 111-005-003 or 115-005-003) using a standard coupling protocol. In brief, the reactive dye (**13**, **16-18**) was dissolved in anhydrous DMSO (ca. 2 mg $ml^{-1}$), and mixed with 0.5–1 mg antibody in a proportion of 5–10 equivalents (dye/protein). The pH of the solution was adjusted to ≈8.4 with carbonate buffer (1 M), and stirred for 1 h in the dark. The mixture was purified using a size exclusion column (PD 10, GE Healthcare). UV-Vis measurements in a small volume spectrometer (DS-11+, DeNovix) were performed to determine the fractions containing the protein, and for the determination of the degree of labeling (DOL).

### Conjugation of nanobodies via thiol groups

Unconjugated nanobodies (NanoTag Biotechnologies product nos.: N1202, N2402, N0302 or N0303) with site-specific cysteine residues) were reconstituted and labelled according to manufacturers protocols. In brief, then the pH of the solution was adjusted by the addition of 1/10 volume of 1 M Tris/HCl buffer (pH 8.0), and 1.5- to 3-fold molar excess (in respect to the number of cysteines per nanobody) of maleimide-PaX derivative was immediately added. The reaction mixture was vortexed, overlaid with argon and incubated in the dark, on ice for 1.5 h. The unreacted excess dye was removed, and the buffer was exchanged to PBS using a desalting column (7K MWCO Zeba Spin Desalting Column, Thermo Scientific). The DOL value of the resulting conjugate was determined by ESI-MS.

### Cell culture

COS-7, HeLa, U2OS-Vim-Halo, U2OS-Vim-SNAP[5,6] and HK-2xZFN-mEGFP-Nup107[7] were cultured in Dulbecco's Modified Eagle Medium (DMEM, 4.5 g/L glucose) containing GlutaMAX and sodium pyruvate (ThermoFisher 31966), supplemented with 10 % (v/v) fetal bovine serum (FBS, ThermoFisher 10500064) and 1% Pen Strep (GIBCO 15140122) in a humidified 5 % CO2 incubator at 37 °C. Cells were split every 2 – 4 days or at confluency and regularly tested for mycoplasma contamination.

U2OS-ZFN-SNAP-Nup107[8] and U2OS-NUP96-Halo[9] cells were cultured in McCoy's 5a (modified) medium (GIBCO 26600023) containing L-glutamine and sodium pyruvate, supplemented with 10 % (v/v) FBS and 1% Pen Strep (GIBCO 15140122) in a humidified 5 % $CO_2$ incubator at 37 °C. Cells were split every 2 – 4 days or at confluency and regularly tested for mycoplasma contamination.

**Indirect Immunofluorescence Labelling with Antibody and Nanobody conjugates**

Cells were grown for 12-72 h on glass coverslips and washed twice with PBS (pH 7.4), and fixed with either methanol (MeOH) or preheated (37°C) paraformaldehyde (PFA)—depending on the favored method for the chosen antibodies or the imaging structures.

For MeOH fixation, the samples were treated with MeOH previously cooled to -20°C for 5 min, and finally washed twice with PBS.

PFA fixation was performed with a 4% formaldehyde solution in PBS at room temperature for 25 min, washed once with a quenching solution (QS, 0.1 M $NH_4Cl$ and 0.1 M Glycine in PBS) and then incubated with QS for 10 min at room temperature then washed with PBS.

To reduce unspecific binding blocking buffer (2% BSA in PBS) was added and incubated for 30-60 min at room temperature. For samples fixed with PFA, Triton 100-X was added to the blocking buffer to a concentration of 0.1%.The coverslips were overlaid with the primary antibody solution in blocking buffer and incubated in a humid chamber for 1 h at room temperature, or overnight at 4 °C, and then washed with blocking buffer (3×5 min). The coverslips were then incubated with the secondary antibody or nanobody in blocking buffer, in a humid chamber for 45-60 min at room temperature, and then washed with blocking buffer (1×5 min), and with PBS (2×5 min). After labeling with PaX conjugates, the samples were protected from the ambient light. For nanobody conjugates, postfixation for preservation of nuclear pore complexes, was conducted with PFA (4%) for 10 min and washed with PBS (2×5 min).

Samples were mounted with PBS or Mowiol, and sealed with nail polish or a two-component silicone resin (Picodent Twinsil, Picodent Dental-Produktions- und Vertriebs-GmbH).

In the case of NPC labelling in HeLa Kyoto mEGFP-NUP107 with anti-GFP nanobodies, fixation and permeabilization was conducted according to previously reported protocols[9], after labelling with PaX nanobody conjugates (60 min at rt), samples were post-fixed with PFA (4%) for 10 min and washed with PBS (2×5 min).

#### Neuronal culture preparation and labeling

Cultures of dissociated rat hippocampal primary neurons were prepared from postnatal P0-P1 Wistar rats of either sex and cultured on glass coverslips coated with 100 µg $mL^{-1}$ poly-ornithine (Merck KGaA) and 1 µg $mL^{-1}$ laminin (BD Biosciences). Procedures were performed in accordance with the Animal Welfare Act of the Federal Republic of Germany (Tierschutzgesetz der Bundesrepublik Deutschland, TierSchG) and the Animal Welfare Laboratory Animal Regulations (Tierschutzversuchsverordnung). According to the TierSchG and the Tierschutzversuchsverordnung no ethical approval from the ethics committee is required for the procedure of sacrificing rodents for subsequent extraction of tissues. The procedure for sacrificing P0-P1 rats performed in this study was supervised by animal welfare officers of the Max Planck Institute for Medical Research (MPImF) and conducted and documented according to the guidelines of the TierSchG (permit number assigned by the MPImF: MPI/T-35/18).

Cells were grown in presence of 1-β-D-Arabinofuranosyl-cytosin (Merck KGaA) at 37°C and 5% $CO_2$. Cultures were fixed at 27 days *in vitro* in 4% PFA in PBS, pH 7.4 for 20 min, and quenched for 5 min in PBS supplemented with 100 mM glycine and 100 mM ammonium chloride. Cells were permeabilised for 5 min in 0.1% Triton X-100, blocked with 1% BSA for 30 min and incubated with 1 µM 15 diluted in PBS. After extensive washing in PBS, samples were mounted in Mowiol supplemented with DABCO. The identification of axons was facilitated by the staining of the axon initial segment by means of an anti-neurofascin primary antibody (NeuroMab, cat. 75-172) and an anti-mouse STAR GREEN (Abberior, cat STGREEN-1001) secondary antibody.

#### Optical microscopy and nanoscopy of cells with organelle-specific PaX derivatives

Stocks solutions of $PaX_{560}$-Mito (**19**) or $PaX_{560}$-Lyso (**20**) were prepared in DMSO (ca. 1 mM). HeLa cells were grown for 12-72 h on glass coverslips. Cells were incubated in the dark for 30 min to overnight (depending on the dye and experiment) with the respective fluorescent ligands diluted from DMSO stock solutions with culture medium (without phenol red) to a final concentration of 10 - 500 nM. After labeling with PaX dyes, the samples were protected from the ambient light. Cells were washed with cell culture medium for ca. 15-30 minutes; then the medium was changed for fresh media for imaging. For colocalization or two-color experiments, the cells were co-stained with the always-on dyes MitoTracker™ Deep Red FM (ThermoFischer Scientific) or SiR Lysosome (SpiroChrome).

Colocalization analysis (Pearson r) was performed in Coloc 2 plugin on ImageJ (version 1.52p).

**Optical microscopy and nanoscopy of cells with self-labelling enzymes**

Stocks solutions of SNAP-tag ligand (**22**) or HaloTag ligand derivatives of the PaX dyes (**21**, **23**-**25**) were prepared in DMSO (ca. 1 mM). U2OS cells that stably expressed Vimentin-HaloTag, Vimentin-SNAP[5,6], NUP96-HaloTag, or NUP107-SNAP were grown for 12-72 h on glass coverslips. Cells were incubated in the dark for 30 min to overnight (depending on the dye and experiment) with the respective fluorescent ligands diluted from DMSO stock solutions with culture medium (without phenol red) to a final concentration of 10 - 500 nM. After labeling with PaX dyes, the samples were protected from the ambient light. Cells were washed with cell culture medium for ca. 15-30 minutes; then the medium was changed for fresh media for live-cell imaging. For two-color experiments, the cells were co-stained with the always-on dyes Abberior Live 560-Tubulin, Hoechst (Hoechst 33342 Solution, ThermoFischer Scientific), or an organelle-specific PaX as described above.

Fixation after live labelling, for preservation of nuclear pore complexes, was performed with 4% PFA as described above, followed by washing with QS (1×) followed by 10 min incubation with QS. Samples were then washed with PBS (2×5 min).

Samples were mounted in PBS, and imaged by confocal, STED, or single molecule localization microscopy.

#### Single detection channel two-color multiplexing by photoactivation

U2OS cells that stably expressed vimentin-HaloTag were labelled with $PaX_{560}$-Halo (**21**) and Abberior Live 560-Tubulin (AL-560) as described above. Sequential imaging was performed on a confocal setup before photoactivation, after photobleaching of Abberior Live 560-Tubulin with high power of 561 nm light, and after photoactivation of **20-Halo** with 405 nm activation. All images were acquired with the same excitation laser, 561 nm, and detection channel, 574 nm–626 nm range. A single-channel, pseudo two-color (multiplexed) image of two different targets may be obtained from overlaying the images.

#### Color multiplexing of PaX by photoactivation kinetics

COS-7 cells were fixed with MeOH and immunolabeled as described above with a mixture of anti-clathrin heavy chain rabbit antibody (ab21679, abcam) and anti-alpha tubulin mouse antibody (302 211, Synaptic Systems), and then with a mixture of anti-rabbit secondary antibody labelled with $PaX_{480}$-NHS (**16**) and anti-mouse secondary antibody labelled with $PaX_{560}$-NHS (**13**)**.** Two-color imaging was performed simultaneously (line-by-line) in two channels with excitation at 485 nm and detection in a 500-551 nm window and excitation at 561 nm and detection in a 571-691 nm window for compounds $PaX_{480}$ and $PaX_{560}$, respectively. Sequential imaging was performed on a confocal setup before photoactivation, after photoactivation of $PaX_{560}$ with low activation laser dose (405 nm), and after photoactivation of $PaX_{480}$ with higher activation laser dose (405 nm) to obtain two-color image of two photoactivatable dyes multiplexed by photoactivation kinetics.

The analogous experiment was conducted with a mixture of anti-rabbit secondary antibody labelled with $PaX_{480}$-NHS (**16**) and anti-mouse secondary antibody labelled with $PaX_{525}$-NHS (**17**). Two-color imaging was performed simultaneously (line-by-line) in two channels with excitation at 485 nm and detection in a 500-551 nm window and excitation at 561 nm and detection in a 580-650 nm window for compounds $PaX_{480}$ and $PaX_{525}$, respectively.

**Two-Color PALM of PaX by photoactivation kinetics**

COS-7 cells were fixed with MeOH and immunolabeled as described above with a mixture of anti-clathrin primary antibody (from rabbit) and anti-alpha tubulin primary antibody (from mouse), and then with a mixture of anti-rabbit secondary antibody labelled with $PaX_{480}$-NHS (**16**) and anti-mouse secondary antibody labelled with $PaX_{560}$-NHS (**13**).

Two-color PALM imaging was done by first imaging compound **13** using a minimal amount of photoactivation light (1 μW), and the appropriate excitation laser and filter combination until photoactivation events were exhausted. Compound **16** was then imaged, using appropriate excitation laser and filter combination, by applying an increasing photoactivation light dose.

### SYNTHESIS AND CHARACTERIZATION

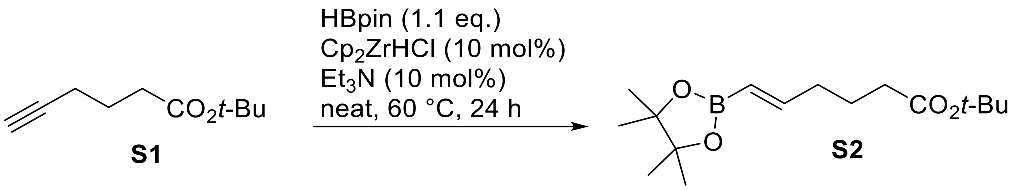


**Compound S2** was prepared according to the method reported in [10]. In a dried 10 mL crimp-top tube (a 2-5 mL Biotage microwave vial was used), compound **S1** (1 g, 6 mmol; prepared according to the literature procedure[11]) and Schwartz's reagent (zirconocene chloride hydride; 155 mg, 0.6 mmol, 10 mol%) were placed, and the contents of the vial were degassed on a Schlenk line. Pinacolborane (HBpin; 950 µL, 6.55 mmol, ~1.1 equiv) was then injected followed by triethylamine (84 µL, 0.6 mmol, 10 mol%), the vial was placed in a 60 °C oil bath and the reaction mixture was stirred for 24 h. The mixture was then cooled down to rt, diluted with diethyl ether and filtered through a 2 cm plug of silica, washing with diethyl ether (50 mL). The filtrate was evaporated and the product was isolated by flash chromatography on Biotage Isolera system (40 g RediSep Rf cartridge, gradient 0% to 20% ethyl acetate/hexane) to give colorless oil, yield 1.28 g (72%).

$^1$H NMR (400 MHz, $CDCl_3$): δ 6.60 (dt, *J* = 18.0, 6.4 Hz, 1H), 5.45 (dt, *J* = 18.0, 1.6 Hz, 1H), 2.26 – 2.14 (m, 4H), 1.72 (p, *J* = 7.6 Hz, 2H), 1.44 (s, 9H), 1.27 (s, 12H).
$^{13}$C NMR (101 MHz, $CDCl_3$): δ 173.0, 153.4, 119.6 (br.), 83.2, 80.2, 35.2, 35.1, 28.2, 24.9, 23.7.
HRMS (ESI) m/z: $[M+H]^+$ Calcd for $C_{16}H_{29}BO_4$ 297.2235; Found 297.2231.

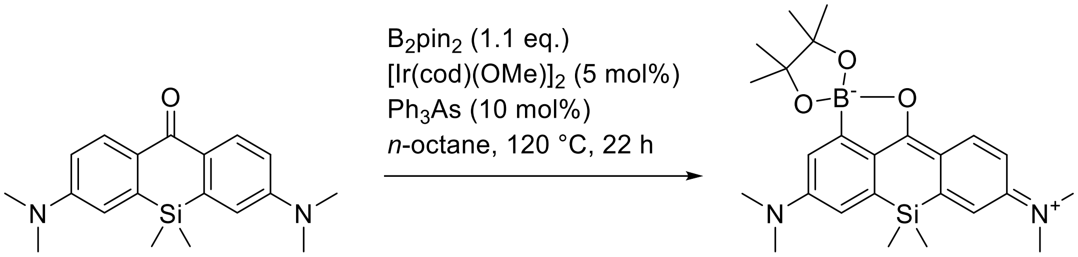


**A1** **B1**

**Compound B1.** In a 25 mL round-bottom flask, the mixture of compound **A1** (324 mg, 1 mmol; known compound [12]), bis(pinacolato)diboron (280 mg, 1.1 mmol, 1.1 equiv), $[Ir(cod)(OMe)]_2$ (33 mg, 0.05 mmol, 5 mol%) and triphenylarsine (31 mg, 0.01 mmol, 10 mol%) in dry *n*-octane (10 mL) was degassed on a Schlenk line and stirred at 120 °C for 22 h. On cooling, the reaction mixture was evaporated on Celite and the product was isolated by flash chromatography on Biotage Isolera system (10 g Biotage Sfär Duo cartridge, gradient 0% to 100% A:B, A = 20% ethyl acetate in dichloromethane, B = dichloromethane) and freeze-dried from dioxane to give 322 mg (72%) of **B1** as an orange solid.

$^1$H NMR (400 MHz, $CDCl_3$): δ 8.38 (d, *J* = 9.0 Hz, 1H), 7.02 (d, *J* = 2.5 Hz, 1H), 6.80 (d, *J* = 2.7 Hz, 1H), 6.75 (dd, *J* = 9.1, 2.7 Hz, 1H), 6.72 (br.s, 1H), 3.14 (s, 12H), 1.43 (s, 12H), 0.39 (s, 6H). $^{13}$C NMR (101 MHz, $CDCl_3$): δ 187.5, 153.4, 153.3, 145.9, 139.2, 133.8, 122.4, 115.6, 115.1, 114.7, 112.3, 80.5, 40.5, 40.1, 25.7, -1.2.

HRMS (ESI) m/z: $[M]^+$ Calcd for $C_{19}H_{24}BN_2O_2Si$ 351.1698; Found 351.1693 – corresponds to the pinacol ester hydrolysis product (**B1a**):

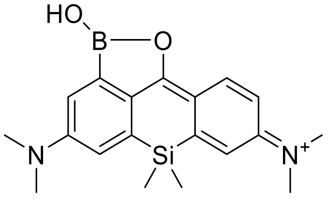

**B1a**

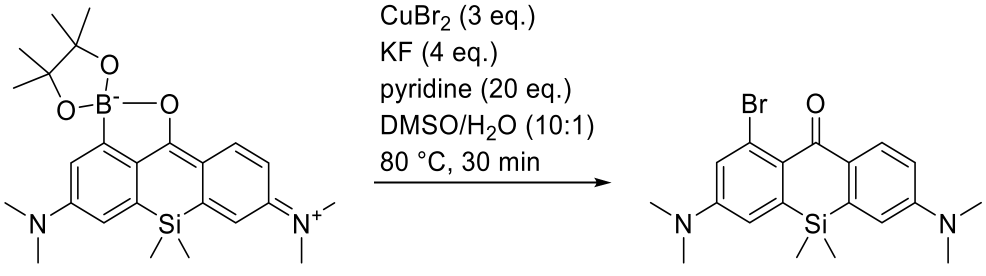


**B1** **C1**

**Compound C1.** In a 25 mL round-bottom flask, compound **B1** (259 mg, 0.58 mmol), potassium fluoride (135 mg, 2.32 mmol, 4 equiv), copper(II) bromide (388 mg, 1.74 mmol, 3 equiv) were placed, followed by addition of DMSO (5 mL), water (500 µL) and pyridine (940 µL, 11.6 mmol, 20 equiv). The reaction mixture was stirred at 80 °C for 30 min. Upon cooling to rt, the mixture was diluted with ethyl acetate and poured into water (80 mL). The product was extracted with ethyl acetate (4 × 30 mL), the combined extracts were washed with brine (50 mL) and dried over $Na_2SO_4$. The product was isolated by flash chromatography on Biotage Isolera system (25 g Interchim SiHP 30 µm cartridge, gradient 0% to 100% A/B, A: 10% ethyl acetate in dichloromethane, B: dichloromethane) and freeze-dried from dioxane to give 212 mg (91%) of **C1** as yellow solid.

$^1$H NMR (400 MHz, $CDCl_3$): δ 8.20 (d, *J* = 8.9 Hz, 1H), 7.06 (d, *J* = 2.7 Hz, 1H), 6.83 (dd, *J* = 9.0, 2.8 Hz, 1H), 6.76 (br.d, *J* = 2.8 Hz, 1H), 6.74 (d, *J* = 2.7 Hz, 1H), 3.07 (s, 6H), 3.05 (s, 6H), 0.46 (s, 6H).

$^{13}$C NMR (101 MHz, $CDCl_3$): δ 186.3, 151.2, 150.9, 142.3, 138.4, 131.5, 127.9, 125.3, 120.1, 114.1, 113.8, 113.6, 40.3, 40.0, -1.0.

HRMS (ESI) m/z: $[M+H]^+$ Calcd for $C_{19}H_{23}BrN_2OSi$ 403.0836; Found 403.0830.

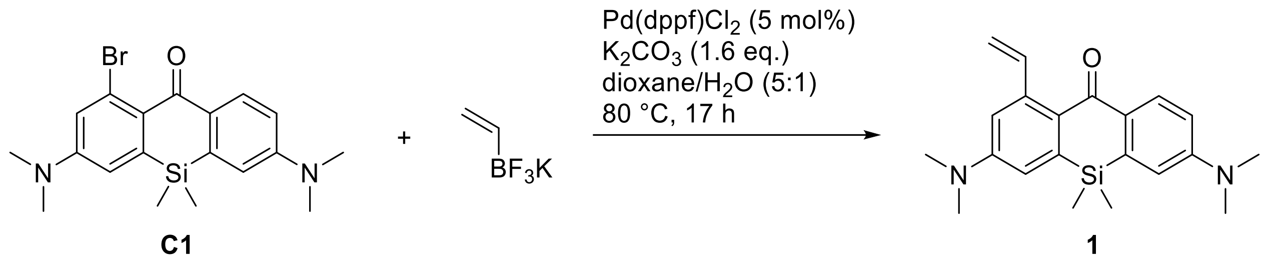

**Compound 1.** In a 10 mL tube, compound **C1** (40 mg, 0.10 mmol), potassium vinyltrifluoroborate (15 mg, 0.11 mmol, 1.1 equiv), $K_2CO_3$ (22 mg, 0.16 mmol, 1.6 equiv) and Pd(dppf)$Cl_2$·$CH_2Cl_2$ (4.1 mg, 5.0 µmol, 5 mol%) were loaded. Dioxane (1.0 mL) and water (0.2 mL) were added. The mixture was sparged with argon for 30 min, and the tube sealed and stirred at 80 °C for 17 h. Upon cooling, the reaction mixture was diluted with ethyl acetate (10 mL) and washed with sat. aq. $NH_4Cl$ (10 mL) and brine (10 mL). The organics were dried over $Na_2SO_4$, filtered, evaporated. The product was isolated by flash chromatography on Biotage Isolera system (12g Interchim SiHP 30 µm cartridge, gradient 0% to 30% ethyl acetate/hexane) and freeze-dried from 1,4-dioxane to yield 31 mg (87%) of **1** as a yellow solid.

$^1$H NMR (400 MHz, $CDCl_3$): δ 8.25 (d, *J* = 8.9 Hz, 1H), 7.58 (dd, *J* = 17.2, 10.7 Hz, 1H), 6.82 (dd, *J* = 9.0, 2.8 Hz, 1H), 6.80 – 6.75 (m, 3H), 5.44 (dd, *J* = 17.2, 1.8 Hz, 1H), 5.24 (dd, *J* = 10.8, 1.8 Hz, 1H), 3.10 (s, 6H), 3.07 (s, 6H), 0.46 (s, 6H).

$^{13}$C NMR (101 MHz, $CDCl_3$): δ 187.8, 151.3, 150.9, 144.2, 142.0, 141.4, 139.1, 132.0, 131.5, 128.5, 114.6, 114.1, 113.8, 113.4, 112.6, 40.2, 40.1, -0.8.

HRMS (ESI) m/z: $[M+H]^+$ Calcd for $C_{21}H_{27}N_2OSi$: 351.1887, found: 351.1888.

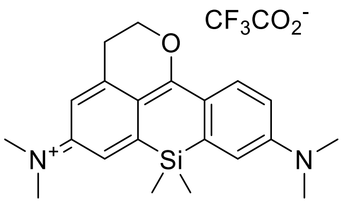

**1-CF**

**Compound 1-CF.** Compound **1** (8 mg, 22.8 µmol) was placed in a 5 mL pear-shaped flask equipped with a stirring bar, dissolved in degassed methanol (3 mL) and irradiated in a Penn OC Photoreactor m1 using a custom 405 nm LED light source (20% power, 1 min; see General experimental information for details). The resulting red solution was evaporated, the product was isolated by preparative HPLC (Interchim Uptisphere Strategy PhC4 250×21.2 mm 5 µm, solvent flow rate 18 mL/min, gradient 35% to 70% A:B, A – acetonitrile + 0.1% (v/v) trifluoroacetic acid, B – water + 0.1% (v/v) trifluoroacetic acid) and freeze-dried from dioxane to give 6 mg (57%) of **1-CF** as dark purple solid.

$^{1}$H NMR (400 MHz, $CDCl_3$): δ 8.24 (d, $J$ = 9.5 Hz, 1H), 6.91 (d, $J$ = 2.5 Hz, 1H), 6.88 (d, $J$ = 2.8 Hz, 1H), 6.81 (dd, $J$ = 9.5, 2.8 Hz, 1H), 6.71 (d, $J$ = 2.5 Hz, 1H), 4.79 (t, $J$ = 6.4 Hz, 2H), 3.32 (s, 6H), 3.26 (s, 6H), 3.22 (t, $J$ = 6.4 Hz, 2H), 0.50 (s, 5H).

$^{13}$C NMR (101 MHz, $CDCl_3$): δ 134.1, 118.3, 117.1, 113.6, 112.5, 69.3, 40.6, 40.4, 28.5, -1.3 (indirect detection from a gHSQC experiment, only H-coupled carbons are resolved).

HRMS (ESI) m/z: $[M]^+$ Calcd for $C_{21}H_{27}N_2OSi$: 351.1887, found: 351.1883.

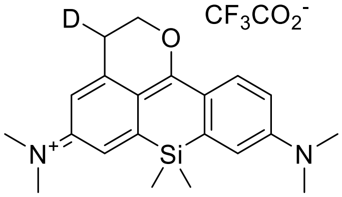


**1-CF-*d***

**Compound 1-CF-*d*.** Compound **1** (5.5 mg, 15.7 µmol) was placed in a 5 mL pear-shaped flask equipped with a stirring bar, dissolved in degassed methanol-$d_4$ (2 mL) and irradiated in a Penn OC Photoreactor m1 using a custom 405 nm LED light source (20% power, 1 min; see General experimental information for details). The resulting red solution was evaporated, the product was isolated by preparative HPLC (Interchim Uptisphere Strategy PhC4 250×21.2 mm 5 µm, solvent flow rate 18 mL/min, gradient 30% to 70% A:B, A – acetonitrile + 0.1% (v/v) trifluoroacetic acid, B – water + 0.1% (v/v) trifluoroacetic acid) and freeze-dried from dioxane to give 4.5 mg (62%) of **1-CF-*d*** as dark purple solid.

$^1$H NMR (400 MHz, $CDCl_3$): δ 8.24 (d, *J* = 9.5 Hz, 1H), 6.91 (d, *J* = 2.5 Hz, 1H), 6.88 (d, *J* = 2.8 Hz, 1H), 6.81 (dd, *J* = 9.5, 2.8 Hz, 1H), 6.69 (d, *J* = 2.5 Hz, 1H), 4.78 (d, *J* = 6.4 Hz, 2H), 3.32 (s, 6H), 3.26 (s, 6H), 3.19 (t, *J* = 6.4 Hz, 1H), 0.50 (s, 5H).

$^{13}$C NMR (101 MHz, $CDCl_3$): δ 134.1, 118.3, 117.2, 113.6, 112.4, 69.3, 40.6, 40.4, 28.2, -1.3 (indirect detection from a gHSQC experiment, only H-coupled carbons are resolved).

HRMS (ESI) m/z: $[M]^+$ Calcd for $C_{21}H_{26}DN_2OSi$: 352.1950, found: 352.1949.

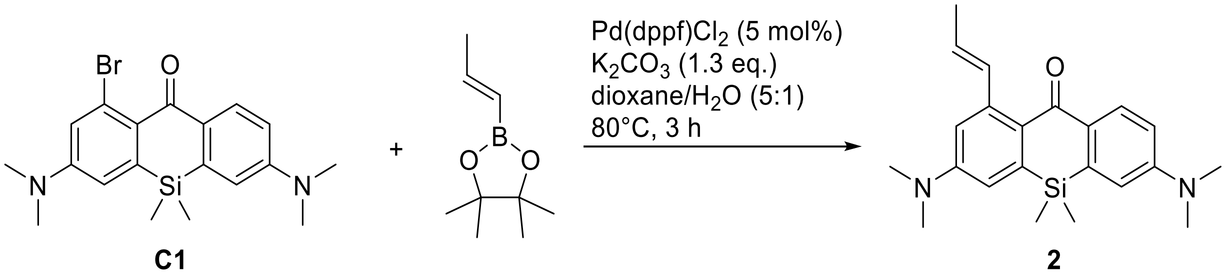

**Compound 2.** In a 10 mL tube, compound **C1** (40 mg, 0.10 mmol), *trans*-1-propenylboronic acid pinacol ester (18 mg, 0.11 mmol, 1.1 equiv), $K_2CO_3$ (18 mg, 0.13 mmol, 1.3 equiv) and $Pd(dppf)Cl_2{\cdot}CH_2Cl_2$ (4.1 mg, 5.0 µmol, 5 mol%) were loaded. Dioxane (1.0 mL) and water (0.2 mL) were added. The mixture was sparged with argon for 30 min, the tube was sealed and the mixture was stirred at 80°C for 3 h. Upon cooling, the reaction mixture was diluted with ethyl acetate (10 mL) and washed with sat. aq. $NH_4Cl$ (10 mL) and brine (10 mL). The organics were dried over $Na_2SO_4$, filtered, evaporated. The product was isolated by flash chromatography on Biotage Isolera system (12 g Interchim SiHP 30 µm cartridge, gradient 0% to 30% ethyl acetate/hexane) and freeze-dried from dioxane to yield 16 mg (43%) of **2** as a yellow solid.

$^1$H NMR (400 MHz, $CDCl_3$): δ 8.25 (d, *J* = 8.9 Hz, 1H), 7.33 – 7.26 (m, 1H), 6.82 (dd, *J* = 9.0, 2.8 Hz, 1H), 6.75 (dt, *J* = 5.4, 2.8 Hz, 3H), 5.93 (dq, *J* = 15.4, 6.6 Hz, 1H), 3.09 (s, 6H), 3.07 (s, 6H), 1.95 (dd, *J* = 6.6, 1.7 Hz, 3H), 0.45 (s, 6H).

$^{13}$C NMR (101 MHz, $CDCl_3$): δ 188.1, 151.2, 150.8, 144.0, 141.3, 139.0, 135.4, 132.4, 131.4, 128.2, 124.7, 114.2, 114.0, 113.8, 113.4, 40.2, 40.1, 18.8, -0.8.

HRMS (ESI) m/z: $[M+H]^+$ Calcd for $C_{22}H_{28}N_2OSi$: 365.2044, found: 365.2044.

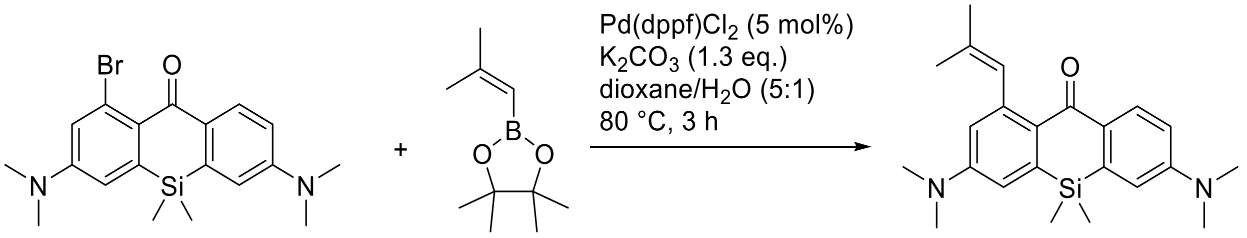


**Compound 3.** In a 10 mL tube, compound **C1** (40 mg, 0.10 mmol), 2-methyl-1-propenylboronic acid pinacol ester (20 mg, 0.11 mmol, 1.1 equiv), $K_2CO_3$ (18 mg, 0.13 mmol, 1.3 equiv) and Pd(dppf)$Cl_2$·$CH_2Cl_2$ (4.1 mg, 5.0 µmol, 5 mol %) were loaded. Dioxane (1.0 mL) and water (0.2 mL) were added. The mixture was sparged with argon for 30 min, the tube was sealed and the mixture was stirred at 80 °C for 3 h. Upon cooling, the reaction mixture was diluted with ethyl acetate (10 mL) and washed with sat. aq. $NH_4Cl$ (10 mL) and brine (10 mL). The organics were dried over $Na_2SO_4$, filtered, evaporated. The product was isolated by flash chromatography on Biotage Isolera system (12g Interchim SiHP 30 µm cartridge, gradient 0% to 30% ethyl acetate/hexane) and freeze-dried from dioxane to yield 28 mg (74%) of **3** as a yellow solid.

$^1$H NMR (400 MHz, $CDCl_3$): δ 8.26 (d, *J* = 9.0 Hz, 1H), 6.87 (s, 1H), 6.81 (dd, *J* = 8.9, 2.8 Hz, 1H), 6.75 (dd, *J* = 10.3, 2.8 Hz, 2H), 6.60 (dd, *J* = 2.9, 0.8 Hz, 1H), 3.07 (d, *J* = 0.9 Hz, 12H), 1.97 (d, *J* = 1.4 Hz, 3H), 1.74 (d, *J* = 1.5 Hz, 3H), 0.46 (s, 6H).

$^{13}$C NMR (101 MHz, $CDCl_3$): δ 187.3, 151.0, 150.1, 143.4, 141.3, 139.0, 132.0, 131.4, 129.7, 129.6, 128.7, 117.0, 113.7, 113.5, 113.2, 40.1, 40.0, 26.3, 19.5, -0.9.

HRMS (ESI) m/z: $[M+H]^+$ Calcd for $C_{23}H_{31}N_2OSi$: 379.2200, found: 379.2200.

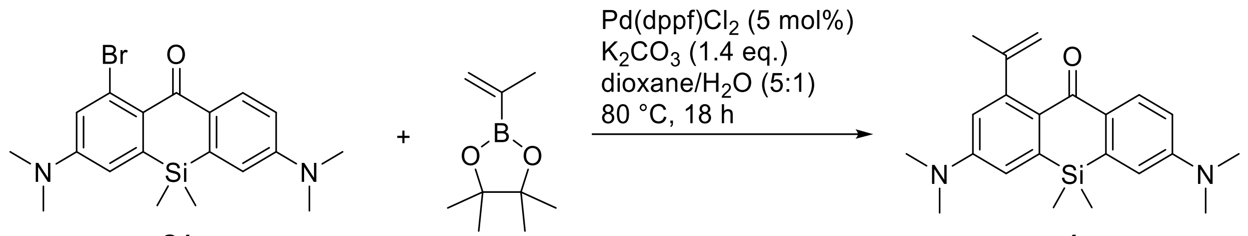


**Compound 4.** In a 10 mL tube, compound **C1** (40 mg, 0.10 mmol), isopropenylboronic acid pinacol ester (18 mg, 0.11 mmol, 1.1 equiv), $K_2CO_3$ (20 mg, 0.14 mmol, 1.4 equiv) and Pd(dppf)$Cl_2$·$CH_2Cl_2$ (4.1 mg, 5.0µmol, 5 mol%) were loaded. Dioxane (1.0 mL) and water (0.2 mL) were added. The mixture was sparged with argon for 30 min, and the tube was sealed and the mixture was stirred at 80 °C for 18 h. Upon cooling, the reaction mixture was diluted with ethyl acetate (10 mL) and washed with sat. aq. $NH_4Cl$ (10 mL) and brine (10 mL). The organics were dried over $Na_2SO_4$, filtered, evaporated. The product was isolated by flash chromatography on Biotage Isolera system (12g Interchim SiHP 30 µm cartridge, gradient 0% to 30% ethyl acetate/hexane) and freeze-dried from dioxane to yield 29 mg (81%) of **4** as a light orange solid.

$^1$H NMR (400 MHz, $CDCl_3$): δ 8.26 (d, *J* = 8.9 Hz, 1H), 6.82 (dd, *J* = 8.9, 2.8 Hz, 1H), 6.76 (dd, *J* = 8.3, 2.8 Hz, 2H), 6.57 (d, *J* = 2.8 Hz, 1H), 5.01 (dq, *J* = 2.9, 1.4 Hz, 1H), 4.80 (dd, *J* = 2.2, 0.9 Hz, 1H), 3.08 (s, 6H), 3.07 (s, 6H), 2.12 (dd, *J* = 1.4, 0.8 Hz, 3H), 0.47 (s, 6H).

$^{13}$C NMR (101 MHz, $CDCl_3$): δ 186.7, 152.1, 151.3, 150.7, 149.5, 141.5, 139.2, 131.9, 131.4, 128.1, 115.6, 113.9, 113.9, 113.4, 109.6, 40.2, 40.1, 24.7, -0.8.

HRMS (ESI) m/z: $[M+H]^+$ Calcd for $C_{22}H_{28}N_2OSi$: 365.2044, found: 365.2045.

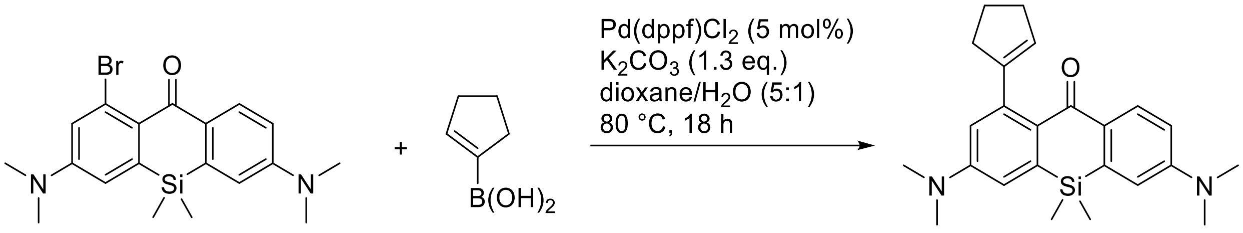

**C1** **5**

**Compound 5.** In a 10 mL tube, compound **C1** (40 mg, 0.10 mmol), 1-cyclopentenylboronic acid (21 mg, 0.11 mmol, 1.1 equiv), $K_2CO_3$ (18 mg, 0.13 mmol, 1.3 equiv) and Pd(dppf)$Cl_2$·$CH_2Cl_2$ (4.1 mg, 5.0 µmol, 5 mol%) were loaded. Dioxane (1.0 mL) and water (0.2 mL) were added. The mixture was sparged with argon for 30 min, and the tube was sealed and the mixture was stirred at 80 °C for 18 h. Upon cooling, the reaction mixture was diluted with ethyl acetate (10 mL) and washed with sat. aq. $NH_4Cl$ (10 mL) and brine (10 mL). The organics were dried over $Na_2SO_4$, filtered, evaporated. The product was isolated by flash chromatography on Biotage Isolera system (12g Interchim SiHP 30 µm cartridge, gradient 0% to 30% ethyl acetate/hexane) and freeze-dried from dioxane to yield 23 mg (59%) of **5** as yellow solid.

$^1$H NMR (400 MHz, $CDCl_3$): δ 8.20 (d, *J* = 8.8 Hz, 1H), 6.81 (dd, *J* = 8.9, 2.8 Hz, 1H), 6.76 (dd, *J* = 13.2, 2.8 Hz, 2H), 6.60 (d, *J* = 2.8 Hz, 1H), 5.58 (p, *J* = 2.0 Hz, 1H), 3.07 (s, 6H), 3.07 (s, 6H), 2.60 – 2.50 (m, 4H), 2.08 (tt, *J* = 8.0, 6.7 Hz, 2H), 0.46 (s, 6H).
$^{13}$C NMR (101 MHz, $CDCl_3$): δ 187.2, 151.2, 150.7, 149.9, 144.3, 141.1, 139.0, 132.3, 131.2, 129.1, 123.8, 115.9, 113.9, 113.9, 113.4, 40.2, 40.1, 36.7, 33.3, 24.8, -0.8.
HRMS (ESI) m/z: $[M+H]^+$ Calcd for $C_{24}H_{30}N_2OSi$: 391.2227, found: 392.2225.

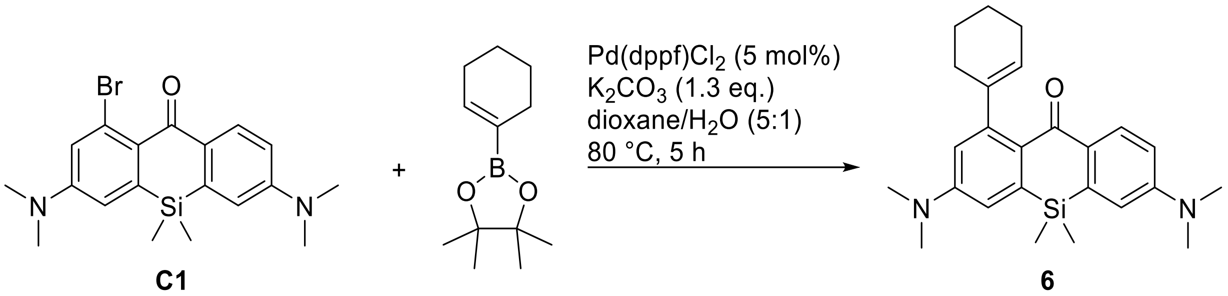

**Compound 6.** In a 10 mL tube, compound **C1** (40 mg, 0.10 mmol), 1-cyclohexen-1-ylboronic acid pinacol ester (23 mg, 0.11 mmol, 1.1 equiv), $K_2CO_3$ (18 mg, 0.13 mmol, 1.3 equiv) and $Pd(dppf)Cl_2 \cdot CH_2Cl_2$ (4.1 mg, 5.0 µmol, 5 mol%) were loaded. Dioxane (1.0 mL) and water (0.2 mL) were added. The mixture was sparged with argon for 30 min, and the tube was sealed and the mixture was stirred at 80°C for 18 h. The reaction mixture was diluted with ethyl acetate (10 mL) and washed with sat. aq. $NH_4Cl$ (10 mL) and brine (10 mL). The organics were dried over $Na_2SO_4$, filtered, evaporated. The product was isolated by flash chromatography on Biotage Isolera system (12g Interchim SiHP 30 µm cartridge, gradient 0% to 40% ethyl acetate/hexane) and freeze-dried from dioxane to yield 17 mg (42%) of **6** as a light yellow solid.

$^1$H NMR (400 MHz, $CDCl_3$): δ 8.21 (d, *J* = 8.9 Hz, 1H), 6.81 (dd, *J* = 8.9, 2.8 Hz, 1H), 6.77 (d, *J* = 2.7 Hz, 1H), 6.73 (d, *J* = 2.9 Hz, 1H), 6.53 (d, *J* = 2.9 Hz, 1H), 5.50 (tt, *J* = 3.7, 1.5 Hz, 1H), 3.08 (s, 6H), 3.07 (s, 6H), 2.26 – 2.13 (m, 4H), 1.88 – 1.79 (m, 2H), 1.78 – 1.68 (m, 2H), 0.46 (s, 6H).

$^{13}$C NMR (101 MHz, $CDCl_3$): δ 187.1, 151.2, 150.7, 149.9, 144.6, 141.2, 139.0, 132.4, 131.3, 128.6, 120.4, 116.1, 113.9, 113.7, 113.3, 40.2, 40.2, 30.6, 25.7, 23.6, 22.6, -0.8.

HRMS (ESI) m/z: $[M+H]^+$ Calcd for $C_{25}H_{32}N_2OSi$: 405.2357, found: 405.2356.

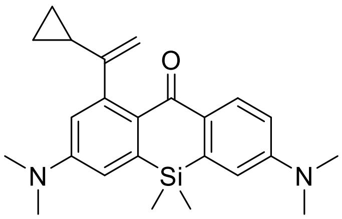

**7**

**Compound 7.** In a 10 mL tube, compound **C1** (40 mg, 0.10 mmol), 1-cyclopropylvinylboronic acid pinacol ester (25 mg, 0.13 mmol, 1.3 equiv; prepared according to the literature procedure[13], $K_2CO_3$ (41 mg, 0.3 mmol, 3 equiv) and $Pd(dppf)Cl_2 \cdot CH_2Cl_2$ (4.2 mg, 5.0 µmol, 5 mol%) were loaded. Dioxane (1.0 mL) and water (0.2 mL) were added. The mixture was degassed on a Schlenk line, the tube was sealed and the reaction mixture was stirred at 80 °C for 6 h. Upon cooling, the reaction mixture was diluted with ethyl acetate (10 mL), poured into brine (50 mL) and extracted with ethyl acetate (3×30 mL). The combined organic layers were dried over $Na_2SO_4$, filtered and evaporated on Celite. The product was isolated by flash chromatography on Biotage Isolera system (25g Interchim SiHP 30 µm cartridge, gradient 10% to 50% ethyl acetate/hexane) and freeze-dried from dioxane to yield 31 mg (79%) of **7** as orange-yellow solid.

$^1$H NMR (400 MHz, $CDCl_3$): δ 8.28 (d, *J* = 8.9 Hz, 1H), 6.81 (dd, *J* = 8.9, 2.8 Hz, 1H), 6.77 (d, *J* = 2.8 Hz, 1H), 6.76 (d, *J* = 2.8 Hz, 1H), 6.45 (d, *J* = 2.8 Hz, 1H), 4.97 (dd, *J* = 1.7, 0.9 Hz, 1H), 4.74 (d, *J* = 1.7 Hz, 1H), 3.08 (s, 6H), 3.07 (s, 6H), 1.83 (ttd, *J* = 8.4, 5.3, 0.9 Hz, 1H), 0.65 (dd, *J* = 8.4, 1.9 Hz, 2H), 0.50 – 0.44 (m, 8H).
$^{13}$C NMR (101 MHz, $CDCl_3$): δ 186.5, 157.2, 151.2, 150.3, 147.2, 141.3, 139.1, 131.9, 131.5, 128.5, 116.6, 114.0, 113.8, 113.4, 106.0, 40.2, 40.1, 17.9, 6.8, -0.8.
HRMS (ESI) m/z: $[M+H]^+$ Calcd for $C_{24}H_{30}N_2OSi$: 391.2200, found: 391.2199.

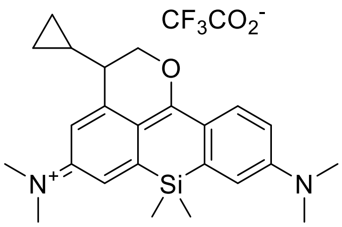

**7-CF**

**Compound 7-CF.** Compound **7** (2.0 mg, 5.12 µmol) was placed in a 5 mL pear-shaped flask equipped with a stirring bar, dissolved in degassed methanol (0.8 mL) and irradiated in a Penn OC Photoreactor m1 using a custom 405 nm LED light source (20% power, 1 min; see General experimental information for details). The resulting red solution was evaporated, chased with $CCl_4$ (2×1 mL), dissolved in 0.6 mL $CDCl_3$ with addition of TFA-*d* (6 µL, 1% v/v), and the photoproduct was characterized by NMR and (HR-)MS analysis of the crude reaction mixture (see Figure S4).

$^1$H NMR (400 MHz, $CDCl_3$ + 1% TFA-*d*): δ 8.28 (d, *J* = 9.4 Hz, 1H), 6.98 – 6.94 (m, 2H), 6.91 – 6.85 (m, 2H), 4.79 (dd, *J* = 11.0, 4.4 Hz, 1H), 4.69 (dd, *J* = 11.0, 6.8 Hz, 1H), 3.33 (s, 6H), 3.27 (s, 6H), 2.32 – 2.24 (m, 1H), 1.02 – 0.92 (m, 1H), 0.81 – 0.67 (m, 2H), 0.55 – 0.37 (m, 2H), 0.52 (s, 3H), 0.49 (s, 3H).
$^{13}$C NMR (101 MHz, $CDCl_3$ + 1% TFA-*d*): δ 134.4, 118.2 (2C), 114.6, 111.4, 73.3, 42.9, 40.9, 40.8, 12.7, 5.0, 3.7, -1.0 (2C) (indirect detection from a gHSQC experiment, only H-coupled carbons are resolved).

HRMS (ESI) m/z: $[M]^+$ Calcd for $C_{24}H_{31}N_2OSi$: 391.2200, found: 391.2199.

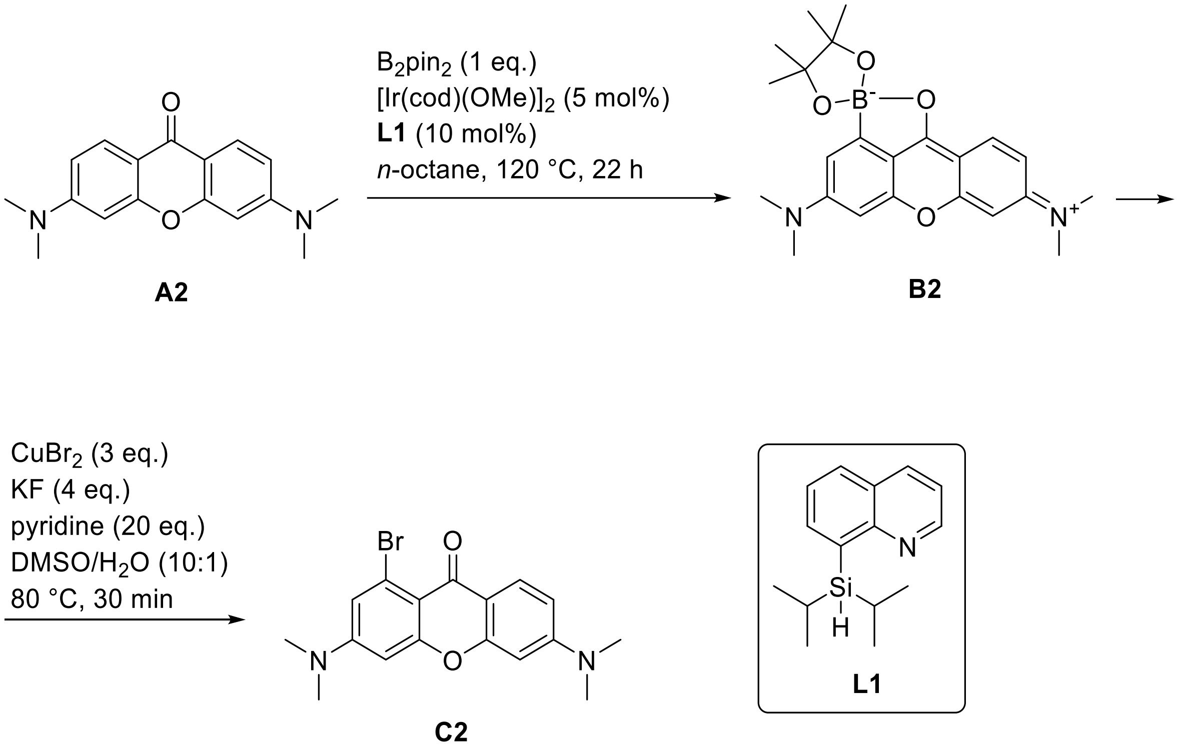


**Compound C2.** In a 25 mL round-bottom flask, the mixture of compound **A2** (382 mg, 1.35 mmol; known compound[14]), bis(pinacolato)diboron (344 mg, 1.35 mmol, 1 equiv), $[Ir(cod)(OMe)]_2$ (45 mg, 0.068 mmol, 5 mol%) and ligand **L1** (8-(diisopropylsilyl)quinoline; known compound [15]) (33 mg, 0.135 mmol, 10 mol%) in dry *n*-octane (13 mL) was degassed on a Schlenk line and stirred at 120 °C for 22 h. On cooling, the reaction mixture was diluted with dichloromethane and filtered through a plug of Celite, washing with dichloromethane. The filtrate was evaporated to dryness in a 50 mL round-bottom flask and the obtained crude **2** was used directly in the next step.

To the residue of crude compound **B2**, potassium fluoride (313 mg, 5.40 mmol, 4 equiv), copper(II) bromide (903 mg, 4.05 mmol, 3 equiv) were added, followed by DMSO (12 mL), water (1.2 mL) and pyridine (2.2 mL, 27 mmol, 20 equiv). The reaction mixture was stirred at 80 °C for 30 min. Upon cooling to rt, the reaction mixture was diluted with ethyl acetate and poured into water (100 mL). The product was extracted with ethyl acetate (3 × 50 mL), the combined extracts were washed with brine (50 mL) and dried over $Na_2SO_4$. The product was isolated by flash chromatography on Biotage Isolera system (40 g RediSep Rf cartridge, gradient 0% to 20% ethyl acetate/dichloromethane) to give 327 mg (67%) of **C2** as yellow solid.

$^{1}H$ NMR (400 MHz, $CDCl_3$): δ 8.08 (d, *J* = 9.0 Hz, 1H), 6.86 (d, *J* = 2.6 Hz, 1H), 6.64 (dd, *J* = 9.0, 2.5 Hz, 1H), 6.40 (d, *J* = 2.6 Hz, 1H), 6.35 (d, *J* = 2.5 Hz, 1H), 3.06 (s, 6H), 3.03 (s, 6H).
$^{13}C$ NMR (101 MHz, $CDCl_3$): δ 174.1, 159.0, 157.0, 154.4, 152.9, 128.1, 122.4, 115.6, 112.2, 109.5, 109.2, 97.7, 96.5, 40.3, 40.1.
HRMS (ESI) m/z: $[M+H]^+$ Calcd for $C_{17}H_{17}BrN_2O_2$ 361.0546; Found 361.0546.

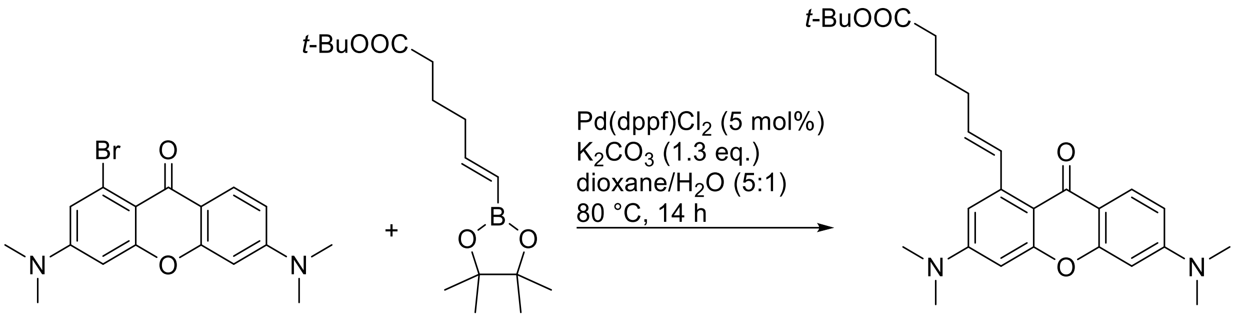


**C2** **S2** **8**

**Compound 8.** In a 10 mL tube, compound **C2** (36 mg, 0.10 mmol), compound **S2** (44 mg, 0.15 mmol), $K_2CO_3$ (18 mg, 0.13 mmol) and $Pd(dppf)Cl_2 \cdot CH_2Cl_2$ (4.1 mg, 5 µmol, 5 mol%) were loaded. Dioxane (1.0 mL) and water (0.2 mL) were added. The mixture was sparged with argon for 30 min, and the tube sealed and stirred at 80 °C for 14 h. Upon cooling, the reaction mixture was diluted with ethyl acetate (10 mL) and washed with sat. aq. $NH_4Cl$ (10 mL), and brine (10 ml). The organics were dried over $Na_2SO_4$, filtered, evaporated. The product was isolated by flash chromatography on Biotage Isolera system (12g Interchim SiHP 30 µm cartridge, gradient 0% to 50% ethyl acetate/hexane) and freeze-dried from dioxane to yield 40 mg (89%) of **8** as a pale yellow solid.

$^1$H NMR (400 MHz, $CDCl_3$): δ 8.08 (d, $J$ = 9.0 Hz, 1H), 7.91 (d, $J$ = 15.7 Hz, 1H), 6.66 (dd, $J$ = 9.0, 2.4 Hz, 1H), 6.63 (dd, $J$ = 2.6, 0.6 Hz, 1H), 6.42 (dd, $J$ = 5.2, 2.5 Hz, 2H), 6.01 (dt, $J$ = 15.5, 6.9 Hz, 1H), 3.09 (s, 6H), 3.08 (s, 6H), 2.39 – 2.28 (m, 4H), 1.90 – 1.78 (m, 2H), 1.45 (s, 9H).
$^{13}$C NMR (101 MHz, $CDCl_3$): δ 176.7, 173.4, 159.3, 157.3, 154.3, 153.1, 142.4, 132.6, 131.4, 127.8, 113.1, 109.5, 109.1, 107.9, 97.1, 96.6, 80.1, 40.3, 40.2, 35.4, 32.7, 28.3, 25.1.
HRMS (ESI) m/z: $[M+H]^+$ Calcd for $C_{27}H_{34}N_2O_4$: 451.2591, found: 451.2590.

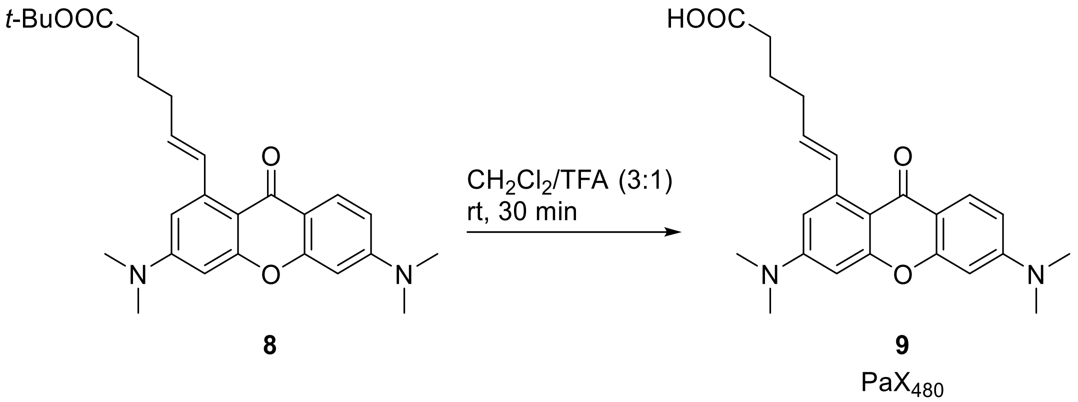

**Compound 9 ($PaX_{480}$).** To a solution of compound **8** (34 mg; 0.075 mmol) in $CH_2Cl_2$ (600 µL) trifluoroacetic acid (200 µL) was added dropwise. The resulting reaction mixture was stirred for 30 minutes at rt, protected from light. The volatiles were removed *in vacuo* by coevaporation with toluene (3 × 10 ml). The product was freeze-dried from dioxane to give 30 mg (~100%, remainder dioxane) of **9** as a pale yellow solid.

$^1$H NMR (400 MHz, DMSO-$d_6$): δ 12.02 (s, 1H), 7.85 (d, *J* = 3.7 Hz, 1H), 7.82 (d, *J* = 3.0 Hz, 1H), 6.75 (dd, *J* = 9.0, 2.4 Hz, 1H), 6.68 (d, *J* = 2.5 Hz, 1H), 6.49 – 6.45 (m, 2H), 6.10 (dt, *J* = 15.7, 6.7 Hz, 1H), 3.06 (s, 6H), 3.05 (s, 6H), 2.32 (t, *J* = 7.4 Hz, 2H), 2.23 (q, *J* = 6.8 Hz, 2H), 1.73 (p, *J* = 7.4 Hz, 2H).

$^{13}$C NMR (101 MHz, DMSO-$d_6$): δ 174.9, 174.5, 158.6, 156.5, 154.1, 152.8, 141.0, 131.4, 131.1, 127.0, 111.8, 109.2, 107.9, 107.0, 96.6, 96.0, 39.6, 33.2, 32.0, 24.1.

HRMS (ESI) m/z: $[M+H]^+$ Calcd for $C_{23}H_{26}N_2O_4$: 395.1965, found: 395.1961.

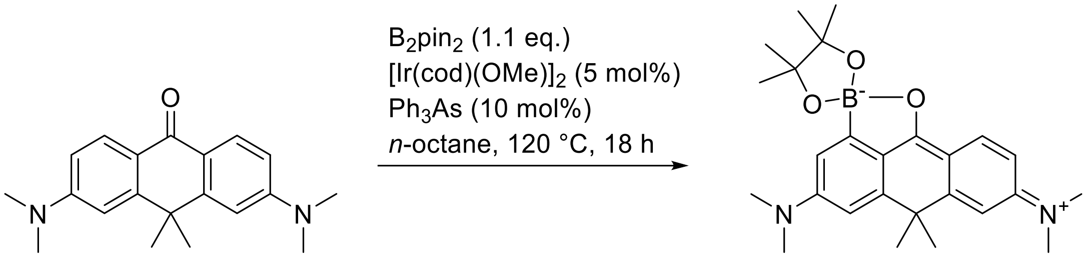


**A3** **B3**

**Compound B3.** In a 10 mL tube, the mixture of compound **A3** (92 mg, 0.30 mmol; known compound[16]), bis(pinacolato)diboron (85 mg, 0.33 mmol, 1.1 equiv), $[Ir(cod)(OMe)]_2$ (12 mg, 0.018 mmol, 5 mol%), triphenylarsine (10 mg, 0.033 mmol, 10 mol%) in dry *n*-octane (3 mL) was degassed on a Schlenk line and stirred at 120 °C for 18 h. On cooling, the reaction mixture was diluted with dichloromethane and filtered through a plug of Celite, washing with dichloromethane. The filtrate was evaporated to dryness and the product was isolated by flash chromatography on a Biotage Isolera system (12g Interchim SiHP 30 µm cartridge, gradient 0% to 10% ethyl acetate/dichloromethane) and freeze-dried from dioxane to 59 mg (45%) of **B3** as an orange solid.

$^1$H NMR (400 MHz, $CDCl_3$): δ 8.20 (d, *J* = 8.9 Hz, 1H), 6.85 (d, *J* = 2.2 Hz, 1H), 6.76 (d, *J* = 2.5 Hz, 1H), 6.70 (dd, *J* = 8.9, 2.5 Hz, 1H), 6.61 (d, *J* = 2.2 Hz, 1H), 3.13 (s, 6H), 3.12 (s, 6H), 1.62 (s, 6H), 1.47 (s, 12H).

$^{13}$C NMR (101 MHz, $CDCl_3$): δ 182.7, 155.4, 154.8, 154.0, 150.8, 130.4, 123.5, 116.1, 113.5, 110.5, 108.4, 107.5, 81.9, 40.6, 40.3, 39.1, 33.3, 25.5.

HRMS (ESI) m/z: $[M]^+$ Calcd for $C_{20}H_{24}BN_2O_2$: 335.1929, found: 335.1928 – corresponds to the pinacol ester hydrolysis product (**B3a**).

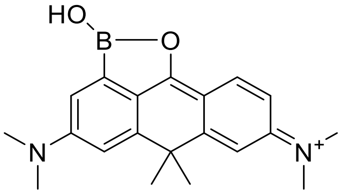

**B3a**

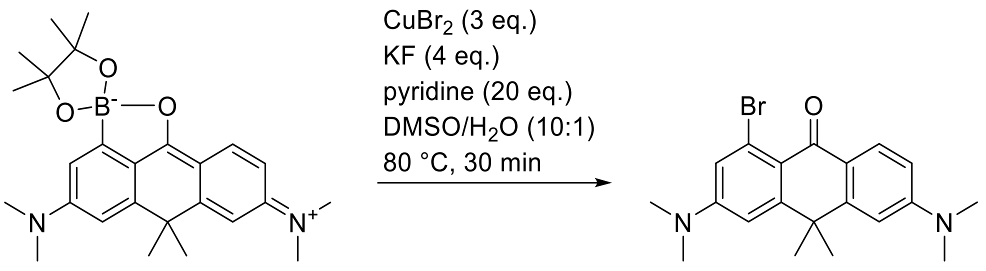


**B3** **C3**

**Compound C3.** In a 10 mL round-bottom flask, compound **B3** (59 mg, 0.14 mmol), potassium fluoride (33 mg, 0.56 mmol, 4 equiv), copper(II) bromide (94 mg, 0.42 mmol, 3 equiv) were placed, followed by addition of DMSO (1.2 mL), water (120 µL) and pyridine (220 µL, 2.7 mmol, 20 equiv). The reaction mixture was stirred at 80 °C for 30 min. Upon cooling to rt, the mixture was diluted with ethyl acetate and poured into water (10 mL). The product was extracted with ethyl acetate (3 × 10 mL), the combined extracts were washed with brine (50 mL) and dried over $Na_2SO_4$. The product was isolated by flash chromatography on Biotage Isolera system (25 g Interchim SiHP 30 µm cartridge, gradient 0% to 10% ethyl acetate/dichloromethane and freeze-dried from 1,4-dioxane to give 39 mg (72%) of **C3** as yellow solid.

$^1$H NMR (400 MHz, $CDCl_3$): δ 8.22 (d, *J* = 8.8 Hz, 1H), 7.00 (d, *J* = 2.6 Hz, 1H), 6.79 (d, *J* = 2.6 Hz, 1H), 6.75 (dd, *J* = 8.9, 2.5 Hz, 1H), 6.70 (d, *J* = 2.5 Hz, 1H), 3.09 (s, 6H), 3.08 (s, 6H), 1.70 (s, 6H).

$^{13}$C NMR (101 MHz, $CDCl_3$): δ 180.6, 154.5, 153.1, 152.0, 150.5, 129.7, 124.1, 121.0, 118.1, 117.6, 111.2, 108.3, 107.0, 40.3, 40.2, 39.1, 34.2.

HRMS (ESI) m/z: $[M+H]^+$ Calcd for $C_{20}H_{23}BrN_2O$: 387.1067, found: 387.1066.

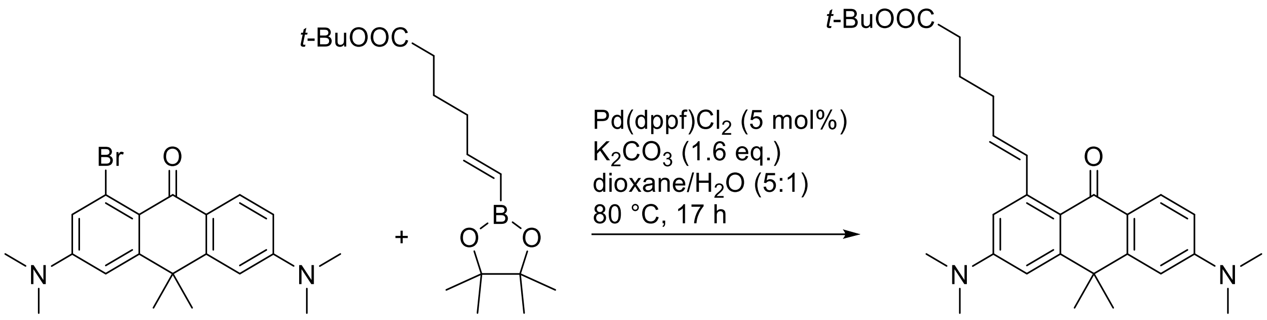


**C3** **S1** **10-COO*t*-Bu**

**Compound 10-COO*t*-Bu.** In a 10 mL tube, compound **7** (33 mg, 0.085 mmol), compound **C3** (37 mg, 0.13 mmol, 1.5 equiv), $K_2CO_3$ (20 mg, 0.14 mmol, 1.6 equiv) and Pd(dppf)$Cl_2$·$CH_2Cl_2$ (3.4 mg, 4.2 µmol, 5 mol%) were loaded. Dioxane (1.0 mL) and water (0.2 mL) were added. The mixture was sparged with argon for 30 min, the tube was sealed and the reaction mixture stirred at 80 °C for 17 h. Upon cooling, the reaction mixture was diluted with ethyl acetate (10 mL) and washed with sat. aq. $NH_4Cl$ (10 mL) and brine (10 ml). The organics were dried over $Na_2SO_4$, filtered, evaporated. The product was isolated by flash chromatography on a Biotage Isolera system (12g Interchim SiHP 30 µm cartridge, gradient 0% to 50% ethyl acetate/hexane) and freeze-dried from dioxane to yield 23 mg (58%) of **10-COO*t*-Bu** as a yellow solid.

$^1$H NMR (400 MHz, $CDCl_3$): δ 8.19 (d, *J* = 8.5 Hz, 1H), 7.64 (d, *J* = 15.6 Hz, 1H), 6.79 – 6.71 (m, 3H), 6.67 (dd, *J* = 2.7, 0.6 Hz, 1H), 5.90 (dt, *J* = 15.4, 6.9 Hz, 1H), 3.11 (s, 6H), 3.09 (s, 6H), 2.38 – 2.30 (m, 4H), 1.85 (p, *J* = 7.6 Hz, 2H), 1.71 (s, 6H), 1.45 (s, 9H).
$^{13}$C NMR (101 MHz, $CDCl_3$): δ 183.0, 173.5, 153.7, 152.9, 152.1, 151.2, 143.2, 134.9, 129.5, 129.3, 121.7, 118.0, 111.3, 111.1, 108.1, 107.3, 80.1, 40.4, 40.3, 38.9, 35.4, 34.3, 32.6, 28.3, 25.3.
HRMS (ESI) m/z: $[M+H]^+$ Calcd for $C_{30}H_{40}N_2O_3$: 477.3112, found: 477.3112.

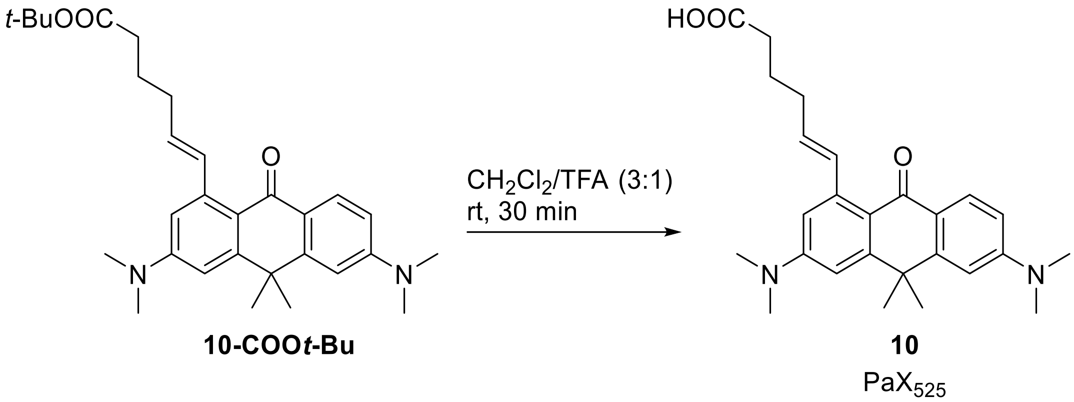

**Compound 10 ($PaX_{525}$).** To a solution of compound **10-COO*t*-Bu** (17.5 mg, 0.075 mmol) in $CH_2Cl_2$ (300 µL), trifluoroacetic acid (100 µL) was added dropwise. The resulting reaction mixture was stirred for 60 minutes at rt, protected from light. The volatiles were removed *in vacuo* by coevaporation with toluene (3 × 10 ml). The product was isolated by flash chromatography on a Biotage Isolera system (12g Interchim SiHP 30 µm cartridge, gradient 20% to 100% ethyl acetate/hexane) and freeze-dried from dioxane to yield 16.5 mg (~100%, remainder dioxane) of **10** as a yellow solid.

$^1$H NMR (400 MHz, DMSO-*d*$_6$) δ 12.01 (s, 1H), 7.92 (d, *J* = 8.9 Hz, 1H), 7.54 (dt, *J* = 15.6, 1.6 Hz, 1H), 6.85 (d, *J* = 2.6 Hz, 1H), 6.82 (d, *J* = 2.5 Hz, 1H), 6.76 (dd, *J* = 8.9, 2.5 Hz, 1H), 6.64 (d, *J* = 2.5 Hz, 1H), 5.89 (dt, *J* = 15.5, 6.7 Hz, 1H), 3.07 (s, 10H), 3.05 (s, 11H), 2.33 (t, *J* = 7.4 Hz, 2H), 2.22 (q, *J* = 7.2 Hz, 2H), 1.72 (p, *J* = 7.6 Hz, 2H), 1.66 (s, 6H).

$^{13}$C NMR (101 MHz, DMSO-*d*$_6$) δ 181.5, 174.5, 153.3, 152.7, 151.8, 150.8, 141.7, 133.9, 129.0, 128.2, 120.5, 116.5, 110.8, 110.3, 108.2, 107.4, 39.8, 39.6, 38.4, 33.8, 33.2, 31.9, 24.3.
HRMS (ESI) m/z: $[M+H]^+$ Calcd for $C_{26}H_{32}N_2O_3$: 421.2486, found: 421.2486.

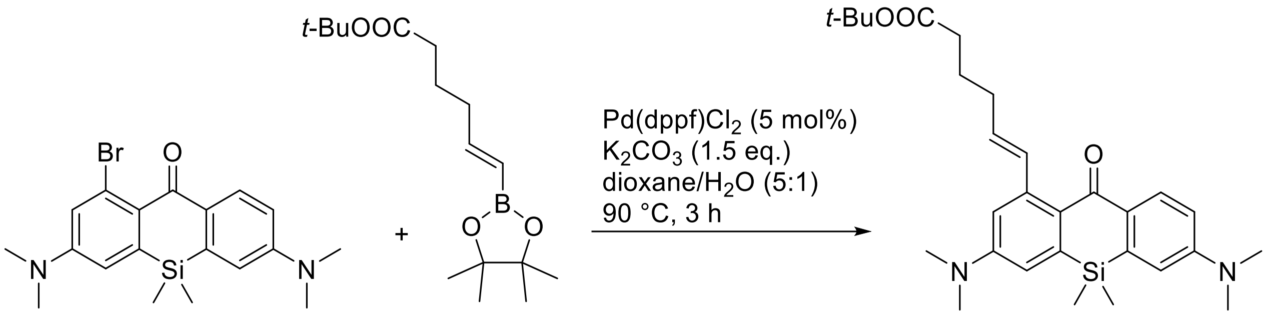


**Compound 11-COO*t*-Bu.** In a 100 mL round bottom flask, compound **C1** (200 mg, 0.50 mmol), compound **S2** (220 mg, 0.75 mmol, 1.5 equiv), $K_2CO_3$ (104 mg, 0.75 mmol, 1.5 equiv) and Pd(dppf)$Cl_2$·$CH_2Cl_2$ (20 mg, 25 µmol, 5 mol%) were loaded. Dioxane (10 mL) and water (2 mL) were added. The mixture was sparged with argon for 30 min and then stirred at 90°C for 3 h. Upon cooling, the reaction mixture was diluted with ethyl acetate (50 mL) and washed with sat. aq. $NH_4Cl$ (50 mL) and brine (50 mL). The organics were dried over $Na_2SO_4$, filtered, evaporated. The product was isolated by flash chromatography on Biotage Isolera system (25 g Interchim SiHP 30 µm cartridge, gradient 0% to 40% ethyl acetate/hexane) and freeze-dried from dioxane to yield 195 mg (79%) of **11-COO*t*-Bu** as a yellow solid.

$^1$H NMR (400 MHz, $CDCl_3$): δ 8.23 (d, *J* = 8.9 Hz, 1H), 7.31 – 7.26 (m, 1H), 6.81 (dd, *J* = 9.0, 2.8 Hz, 1H), 6.77 – 6.72 (m, 3H), 5.89 (dt, *J* = 15.4, 6.9 Hz, 1H), 3.09 (s, 6H), 3.07 (s, 6H), 2.37 – 2.28 (m, 4H), 1.83 (p, *J* = 7.6 Hz, 2H), 1.45 (s, 9H), 0.45 (s, 6H).
$^{13}$C NMR (101 MHz, $CDCl_3$): δ 188.1, 173.5, 151.2, 150.8, 143.6, 141.3, 139.0, 135.0, 132.4, 131.4, 128.9, 128.4, 114.3, 113.9, 113.8, 113.4, 80.1, 40.2, 40.1, 35.4, 32.6, 28.3, 25.3, -0.8.
HRMS (ESI) m/z: $[M+H]^+$ Calcd for $C_{29}H_{42}N_2O_3Si$: 495.3037, found: 495.3032.

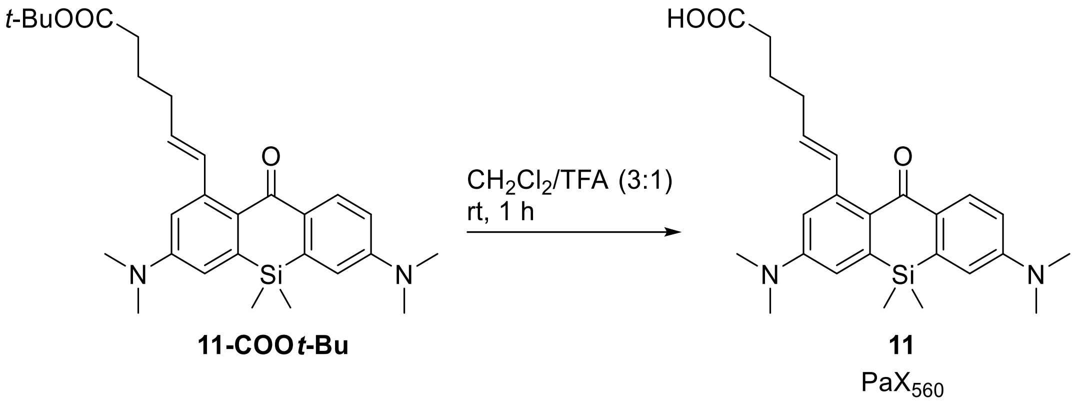


**Compound 11 (PaX$_{560}$).** To a solution of compound **11-COO*t*-Bu** (195 mg, 0.40 mmol) in $CH_2Cl_2$ (3.0 mL), trifluoroacetic acid (1.0 mL) was added dropwise. The resulting reaction mixture was stirred for 60 minutes at rt, protected from light. The volatiles were removed in vacuo by coevaporation with toluene (3×10 ml). The product was isolated by flash chromatography on a Biotage Isolera system (25 g Interchim SiHP 30 µm cartridge, gradient 10% to 50% ethyl acetate/hexane) and freeze-dried from 1,4-dioxane to yield 172 mg (99%) of **11** as an orange solid.

$^1$H NMR (400 MHz, DMSO-$d_6$): δ 12.03 (s, 1H), 7.98 (d, $J$ = 8.9 Hz, 1H), 7.16 (dt, $J$ = 15.5, 1.5 Hz, 1H), 6.89 – 6.80 (m, 3H), 6.73 (d, $J$ = 2.8 Hz, 1H), 5.88 (dt, $J$ = 15.5, 6.8 Hz, 1H), 3.06 (s, 6H), 3.03 (s, 6H), 2.33 (t, $J$ = 7.4 Hz, 2H), 2.21 (q, $J$ = 6.8 Hz, 2H), 1.71 (p, $J$ = 7.4 Hz, 2H), 0.43 (s, 6H).

$^{13}$C NMR (101 MHz, DMSO-$d_6$): δ 186.6, 174.6, 151.0, 150.5, 142.5, 140.8, 138.4, 134.1, 131.1, 130.5, 128.5, 127.0, 114.4, 113.9, 113.1, 112.8, 39.6, 33.2, 31.9, 24.3, -1.1.

HRMS (ESI) m/z: $[M+H]^+$ Calcd for $C_{25}H_{32}N_2O_3Si$: 437.2255, found: 437.2257.

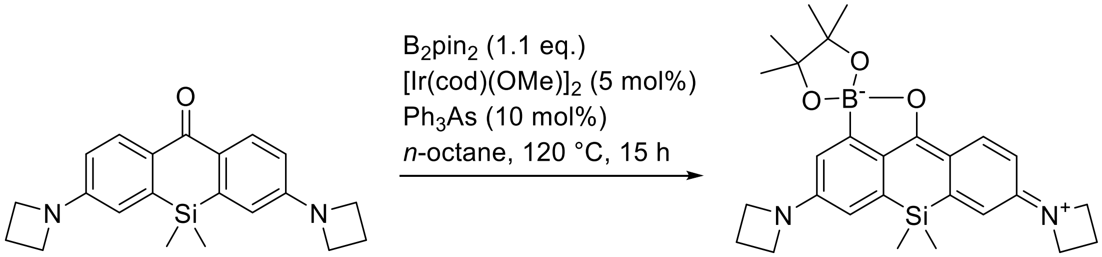


**A4** **B4**

**Compound B4.** In a 10 mL tube, compound **A4** [17] (86 mg, 0.25 mmol), bis(pinacolato)diboron (70 mg, 0.28 mmol, 1.1 equiv), (1,5-cyclooctadiene)(methoxy)iridium(I) dimer (8.2 mg, 0.012 mmol, 5 mol%), triphenylarsine (7.7 mg, 0.025 mmol, 10 mol%) in dry *n*-octane (2.5 mL) was degassed on a Schlenk line and stirred at 120 °C for 15 h. The reaction mixture was diluted with dichloromethane and evaporated onto Celite. The product was isolated by flash chromatography on a Biotage Isolera system (25 g Interchim SiHP 30 µm cartridge, gradient 0% to 20% ethyl acetate/dichloromethane) and freeze-dried from 1,4-dioxane to yield 70 mg (80%) of **B4** as a yellow solid.

$^1$H NMR (400 MHz, $CDCl_3$): δ 8.35 (d, *J* = 8.8 Hz, 1H), 6.66 (d, *J* = 2.2 Hz, 1H), 6.45 (d, *J* = 2.4 Hz, 1H), 6.41 (dd, *J* = 8.7, 2.5 Hz, 1H), 6.33 (d, *J* = 2.2 Hz, 1H), 4.18 – 4.03 (m, 8H), 2.53 – 2.36 (m, 4H), 1.41 (s, 12H), 0.35 (s, 6H).

$^{13}$C NMR (101 MHz, $CDCl_3$): δ 187.6, 153.9, 153.6, 145.8, 139.2, 133.9, 131.6, 122.7, 114.1, 113.4, 113.1, 110.8, 80.4, 51.4, 51.3, 25.7, 16.6, 16.5, -1.3.

HRMS (ESI) m/z: $[M]^+$ Calcd for $C_{21}H_{24}BN_2O_2Si$: 375.1699, found: 375.1701– corresponds to the pinacol ester hydrolysis product (**B4a**):

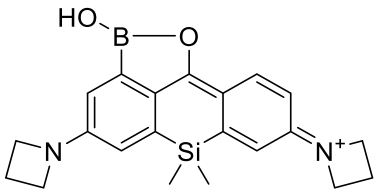

**B4a**

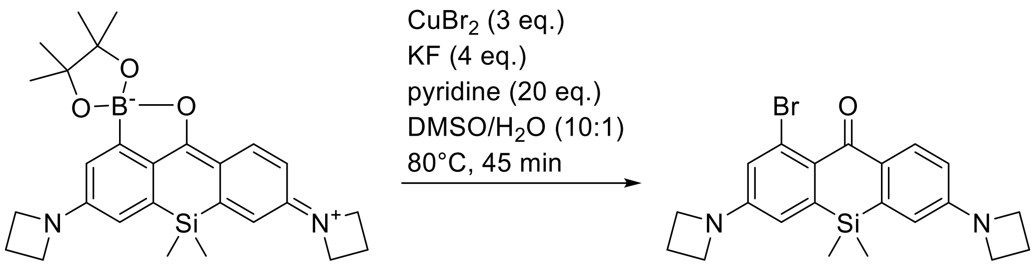


**B4** **C4**

**Compound C4.** In a 25 ml flask, compound **B4** (64 mg, 0.13 mmol), copper(II) bromide (90 mg, 0.40 mmol, 3 equiv) and potassium fluoride (31 mg, 0.54 mmol, 4 equiv) were loaded. DMSO (5 mL), water (500 µL) and pyridine (210 mg, 2.7 mmol, 20 equiv) were added, and the reaction mixture was stirred at 80 °C for 45 minutes. The reaction mixture was then quenched with water (20 mL) and extracted with ethyl acetate (3 × 20 mL). The combined organics were washed with brine (75 mL), dried over $Na_2SO_4$, filtered, and evaporated. The product was isolated by flash chromatography on Biotage Isolera system (12g Interchim SiHP 30 µm cartridge, gradient 0% to 10% ethyl acetate/dichloromethane) and freeze-dried from dioxane to yield 46 mg (83%) of **C4** as a yellow solid.

$^1$H NMR (400 MHz, $CDCl_3$): δ 8.15 (d, *J* = 8.6 Hz, 1H), 6.75 (d, *J* = 2.4 Hz, 1H), 6.51 (dd, *J* = 8.7, 2.5 Hz, 1H), 6.46 – 6.40 (m, 2H), 4.01 (td, *J* = 7.3, 2.0 Hz, 8H), 2.50 – 2.35 (m, 4H), 0.43 (s, 6H).

$^{13}$C NMR (101 MHz, $CDCl_3$): δ 186.7, 152.5, 151.9, 142.3, 138.2, 132.9, 131.2, 128.7, 125.0, 119.0, 112.9, 112.6, 112.4, 51.9, 51.8, 16.9, 16.8, -1.2.

HRMS (ESI) m/z: $[M+H]^+$ Calcd for $C_{21}H_{23}BrN_2OSi$: 427.0836, found: 427.0833.

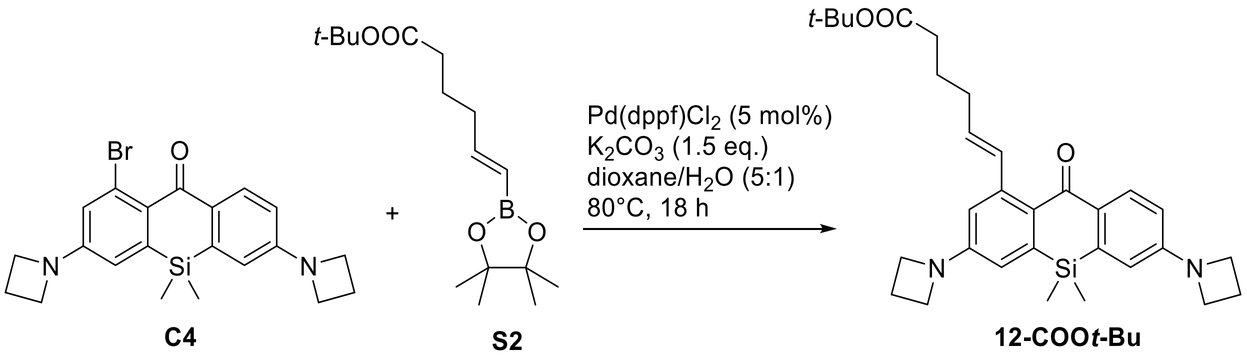

**Compound 12-COO*t*-Bu.** In a 10 mL tube, compound **C4** (43 mg, 0.10 mmol), compound **S2** (44 mg, 0.15 mmol), $K_2CO_3$ (21 mg, 0.15 mmol) and Pd(dppf)$Cl_2$·$CH_2Cl_2$ (4.1 mg, 5.0 µmol, 5 mol %) were loaded. Dioxane (1.0 mL) and water (0.2 mL) were added. The mixture was sparged with argon for 30 min, and the tube was sealed and the mixture was stirred at 80 °C for 18 h. The reaction mixture was quenched with sat. aq. $NH_4Cl$ (10 mL), and extracted with ethyl acetate (3 × 10 mL). The combined organics were washed with brine (50 mL), dried over $Na_2SO_4$, filtered, and evaporated. The product was isolated by flash chromatography on Biotage Isolera system (12g Interchim SiHP 30 µm cartridge, gradient 0% to 30% ethyl acetate/hexane) and freeze-dried from dioxane to yield 37 mg (72%) of **12-COO*t*-Bu** as yellow solid.

$^1$H NMR (400 MHz, $CDCl_3$): δ 8.17 (d, *J* = 8.7 Hz, 1H), 7.21 (dt, *J* = 15.6, 1.5 Hz, 1H), 6.50 (dd, *J* = 8.7, 2.6 Hz, 1H), 6.48 – 6.41 (m, 3H), 5.88 (dt, *J* = 15.4, 6.9 Hz, 1H), 4.06 – 3.96 (m, 8H), 2.49 – 2.36 (m, 4H), 2.36 – 2.25 (m, 4H), 1.82 (p, *J* = 7.6 Hz, 2H), 1.45 (s, 9H), 0.42 (s, 6H).

$^{13}$C NMR (101 MHz, $CDCl_3$): δ 188.4, 173.5, 152.4, 152.0, 143.4, 141.1, 138.8, 134.4, 133.3, 131.1, 129.2, 129.1, 113.2, 112.7, 112.5, 112.3, 80.1, 52.0, 51.9, 35.3, 32.6, 28.3, 25.2, 16.9, -1.0.

HRMS (ESI) m/z: $[M+H]^+$ Calcd for $C_{31}H_{40}N_2O_3Si$: 517.2881, found: 517.2879.

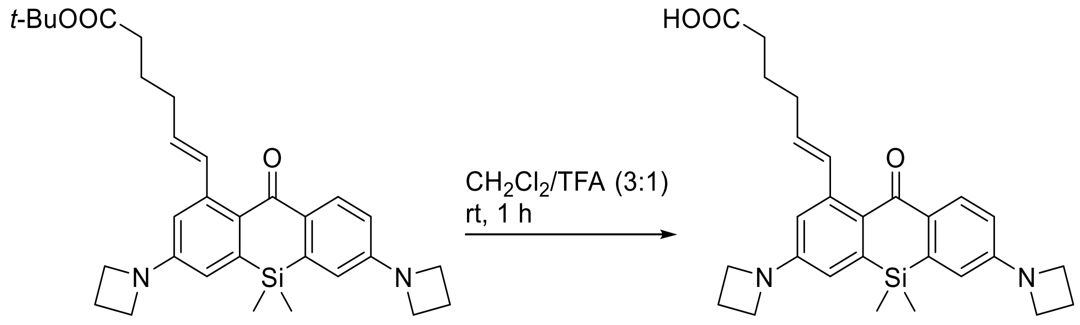


**12-COOt-Bu** → **12** PaX+$_{560}$

**Compound 12 (PaX+$_{560}$).** To a solution of compound **12-COO*t*-Bu** (32 mg; 0.062 mmol) in $CH_2Cl_2$ (600 µL) trifluoroacetic acid (200 µL) was added dropwise. The resulting reaction mixture was stirred for 60 minutes at rt, protected from light. The volatiles were removed in vacuo by coevaporation with toluene (3 × 10 ml). The product was isolated by flash chromatography on Biotage Isolera system (12g Interchim SiHP 30 µm cartridge, gradient 20% to 80% ethyl acetate/hexane) and freeze-dried from 1,4-dioxane to yield 19 mg (67%) of **12** as yellow solid.

$^1$H NMR (400 MHz, DMSO-*d*$_6$): δ 12.01 (s, 1H), 7.95 (d, *J* = 8.7 Hz, 1H), 7.09 (d, *J* = 15.5 Hz, 1H), 6.55 (d, *J* = 2.5 Hz, 1H), 6.53 (d, *J* = 2.5 Hz, 1H), 6.50 (dd, *J* = 8.7, 2.5 Hz, 1H), 6.42 (d, *J* = 2.5 Hz, 1H), 5.85 (dt, *J* = 15.5, 6.8 Hz, 1H), 4.02 – 3.90 (m, 8H), 2.41 – 2.27 (m, 6H), 2.19 (q, *J* = 6.7 Hz, 2H), 1.70 (p, *J* = 7.4 Hz, 2H), 0.40 (s, 6H).

$^{13}$C NMR (101 MHz, DMSO-*d*$_6$): δ 186.9, 174.5, 152.2, 151.8, 142.3, 140.5, 138.2, 133.6, 132.0, 130.4, 128.7, 127.8, 113.3, 112.8, 112.0, 111.5, 51.5, 51.4, 33.1, 31.9, 24.3, 16.2, -1.3.

HRMS (ESI) m/z: $[M+H]^+$ Calcd for $C_{27}H_{32}N_2O_3Si$: 461.2255, found: 461.2253.

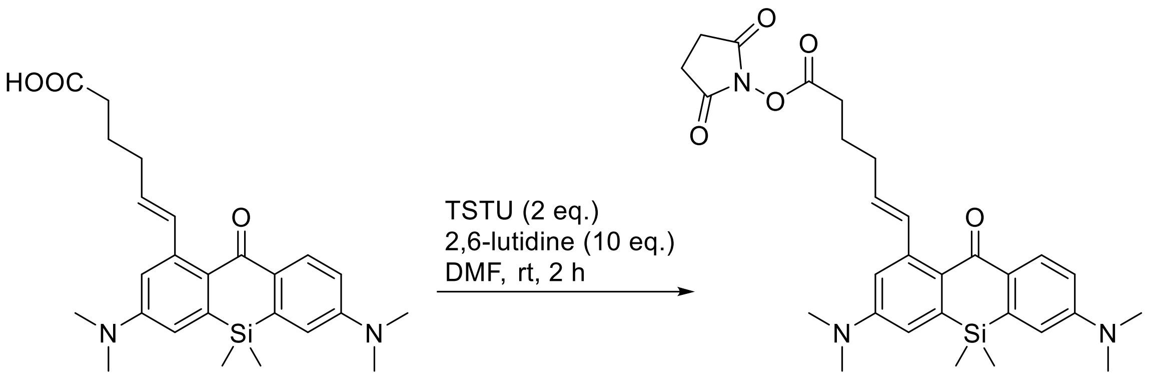


**9** → **13**
$PaX_{560}$-NHS

**Compound 13 ($PaX_{560}$-NHS).** In an amber vial, compound **9** (8.0 mg, 0.018 mmol) and TSTU (11 mg, 0.036 mmol, 2 equiv) were dissolved in DMF (200 µL). 2,6-Lutidine (19 mg, 0.18 mmol) was added, and the reaction mixture was stirred for 3 h at rt. The volatiles were removed *in vacuo*. The product was isolated by flash chromatography on a Biotage Isolera system (12g Interchim SiHP 30 µm cartridge, gradient 20% to 80% ethyl acetate/hexane) and freeze-dried from dioxane to yield 7.5 mg (78%) of **13** as a beige solid.

$^1$H NMR (400 MHz, DMSO-$d_6$): δ 7.99 (d, *J* = 8.9 Hz, 1H), 7.21 – 7.12 (m, 1H), 6.88 – 6.80 (m, 3H), 6.73 (d, *J* = 2.8 Hz, 1H), 5.88 (dt, *J* = 15.5, 6.8 Hz, 1H), 3.06 (s, 6H), 3.03 (s, 6H), 2.85 – 2.78 (m, 6H), 2.30 (q, *J* = 7.2, 6.5 Hz, 2H), 1.85 (p, *J* = 7.4 Hz, 2H), 0.43 (s, 6H).
$^{13}$C NMR (101 MHz, DMSO-$d_6$) δ 186.6, 170.3, 169.1, 151.0, 150.6, 142.4, 140.8, 138.5, 134.8, 131.0, 130.6, 127.6, 127.0, 114.5, 113.9, 113.1, 112.9, 39.6, 31.3, 29.7, 25.5, 24.2, -1.1.
HRMS (ESI) m/z: $[M+H]^+$ Calcd for $C_{29}H_{35}N_3O_5Si$: 534.2419, found: 534.2415.

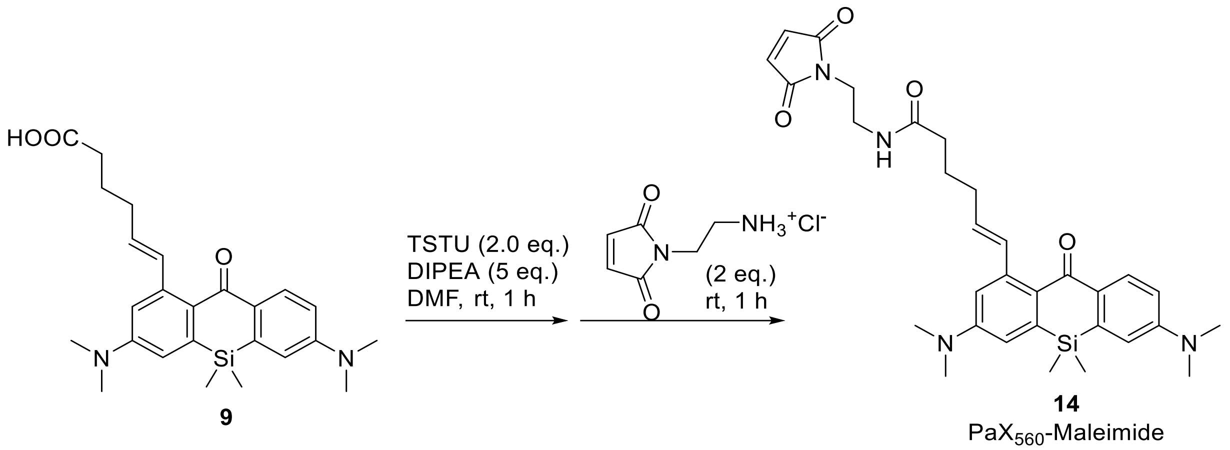

**Compound 14 ($PaX_{560}$-Maleimide).** In an amber vial, compound **9** (22 mg, 0.050 mmol) and TSTU (30 mg, 0.10 mmol) were dissolved in DMF (500 µL) and DIPEA (64 mg, 0.25 mmol) was added and stirred for 60 min. at rt. 1-(1-Aminoethyl)maleimide hydrochloride (18 mg, 0.10 mmol) was added and the reaction was stirred at rt for 60 min. The volatiles were removed *in vacuo*. The product was isolated by flash chromatography on a Biotage Isolera system (12g Interchim SiHP 30 µm cartridge, gradient 50% to 100% ethyl acetate/hexane) and freeze-dried from dioxane to yield 18.8 mg (67%) of **14** as a yellow solid.

$^{1}H$ NMR (400 MHz, $CDCl_3$): δ 8.09 (t, J = 6.3 Hz, 1H), 8.06 (d, J = 9.0 Hz, 1H), 7.03 (d, J = 15.5 Hz, 1H), 6.85 (dd, J = 9.0, 2.8 Hz, 1H), 6.76 (d, J = 2.8 Hz, 1H), 6.75 (d, J = 2.9 Hz, 1H), 6.71 (d, J = 2.8 Hz, 1H), 6.49 (s, 2H), 5.68 (dt, J = 15.4, 7.2 Hz, 1H), 3.81 – 3.75 (m, 2H), 3.68 – 3.58 (m, 2H), 3.10 (s, 12H), 2.39 – 2.32 (m, 2H), 2.27 (q, J = 7.0 Hz, 2H), 1.94 – 1.81 (m, 2H), 0.46 (s, 6H.

$^{13}C$ NMR (101 MHz, $CDCl_3$): δ 188.0, 174.7, 171.0, 151.4, 151.0, 144.2, 141.8, 136.7, 134.0, 131.6, 131.3, 128.7, 114.3, 114.1, 113.9, 113.5, 40.2, 40.1, 38.3, 38.2, 34.4, 31.2, 24.8, -0.8.

HRMS (ESI) m/z: $[M+H]^+$ Calcd for $C_{31}H_{38}N_4O_4Si$: 559.2735, found: 559.2735.

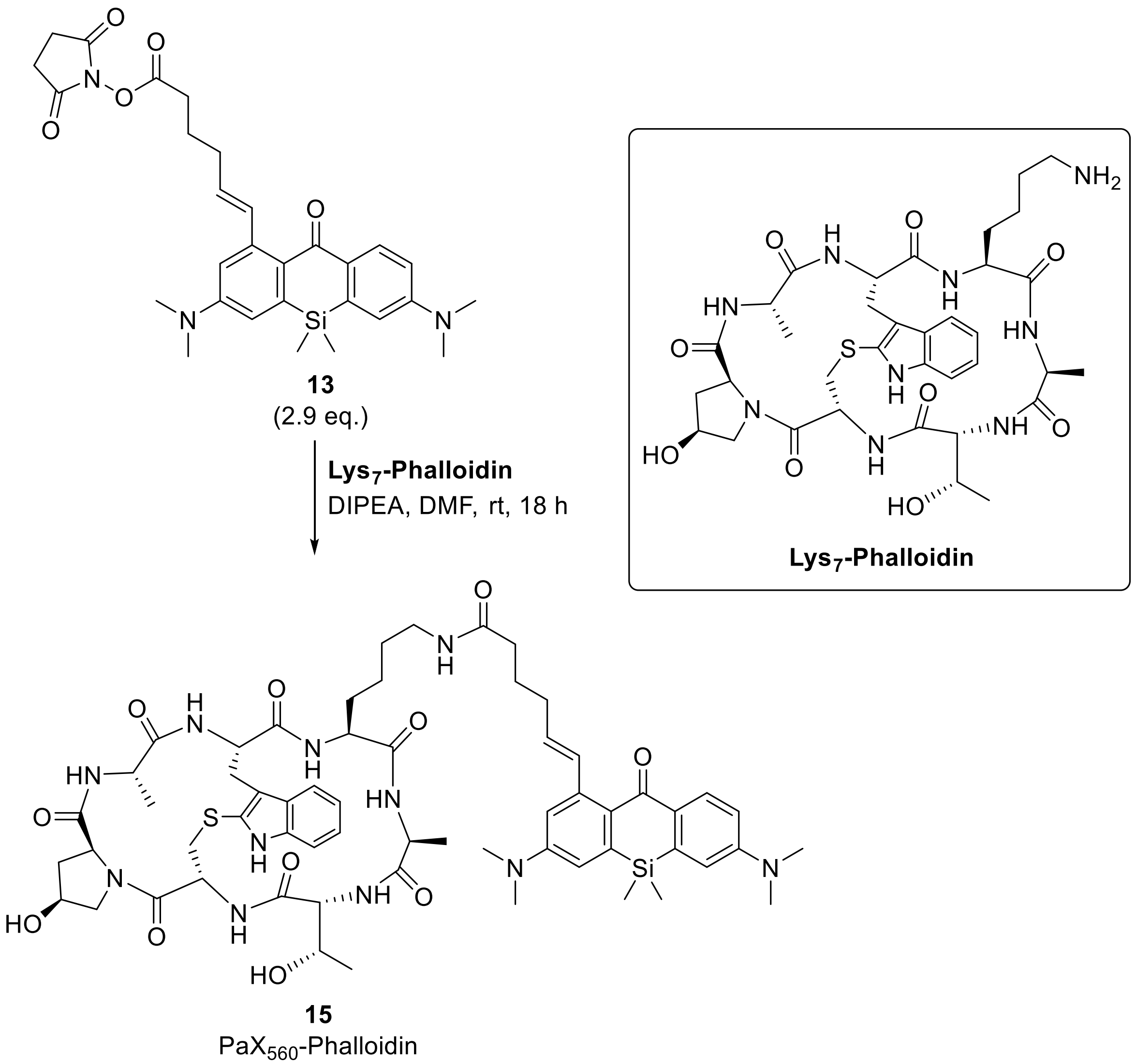


**Compound 15 (PaX$_{560}$-Phalloidin).** In an amber vial, compound **13** (2.0 mg, 3.8 µmol, 2.9 equiv) and **Lys$^{7}$-Phalloidin** (trifluoroacetate salt, Bachem Cat. Nr. H-7636; 1 mg, 1.3 µmol) were dissolved in DMF (200 µL). DIPEA (90 µL) was added, and the reaction mixture was at rt for 18 h. The solvents were removed in vacuo, and the product the product was isolated by preparative HPLC (Interchim Uptisphere Strategy PhC4 250×21.2 mm 5 µm, solvent flow rate 18 mL/min, gradient 40% to 85% A:B, A – acetonitrile + 0.1% (v/v) formic acid, B – water + 0.1% (v/v) formic acid) and freeze-dried from dioxane to give 1.5 mg (97%) of **15** as light pink solid.

HRMS (ESI) m/z: $[M+2H]^{2+}$ Calcd for $C_{60}H_{79}N_{11}O_{11}SSi$: 595.7798, found: 595.7786.

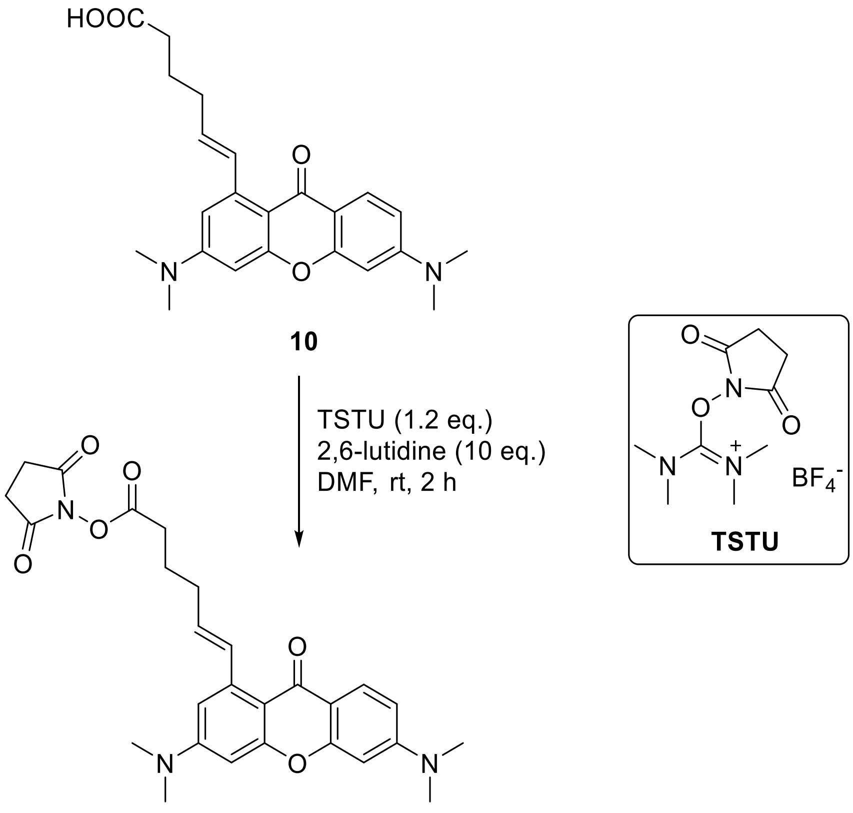


**16**
$PaX_{480}$-NHS

**Compound 16 ($PaX_{480}$-NHS).** In an amber vial, compound **10** (23 mg, 0.058 mmol) and TSTU (*N*,*N*,*N′*,*N′*-tetramethyl-*O*-(*N*-succinimidyl)uronium tetrafluoroborate; 35 mg, 0.12 mmol) were dissolved in DMF (500 µL) and 2,6-lutidine (62 mg, 0.58 mmol) was added and stirred for 2 h at rt. The volatiles were removed in vacuo. The product was isolated by flash chromatography on a Biotage Isolera system (12g Interchim SiHP 30 µm cartridge, gradient 20% to 100% ethyl acetate/hexane) and freeze-dried from 1,4-dioxane to give **16** as an off-white solid. Yield 24 mg (81%, remainder dioxane).

$^{1}$H NMR (400 MHz, $CDCl_3$): δ 8.07 (d, *J* = 9.0 Hz, 1H), 7.94 (d, *J* = 15.7 Hz, 1H), 6.67 (dd, *J* = 9.0, 2.5 Hz, 1H), 6.62 (dd, *J* = 2.6, 0.6 Hz, 1H), 6.44 (d, *J* = 2.6 Hz, 1H), 6.42 (d, *J* = 2.4 Hz, 1H), 5.99 (dt, *J* = 15.6, 6.8 Hz, 1H), 3.10 (s, 6H), 3.08 (s, 6H), 2.83 (s, 4H), 2.74 (dd, *J* = 8.1, 7.2 Hz, 2H), 2.48 – 2.39 (m, 2H), 2.02 (p, *J* = 7.6 Hz, 2H).
$^{13}$C NMR (101 MHz, $CDCl_3$): δ 176.7, 169.3, 168.8, 159.3, 157.4, 154.3, 153.1, 142.2, 133.4, 130.1, 127.8, 113.1, 109.5, 109.1, 107.9, 97.2, 96.6, 40.3, 40.2, 32.1, 30.7, 25.7, 24.5.
HRMS (ESI) m/z: $[M+H]^+$ Calcd for $C_{27}H_{29}N_3O_6$: 492.2129, found: 492.2130.

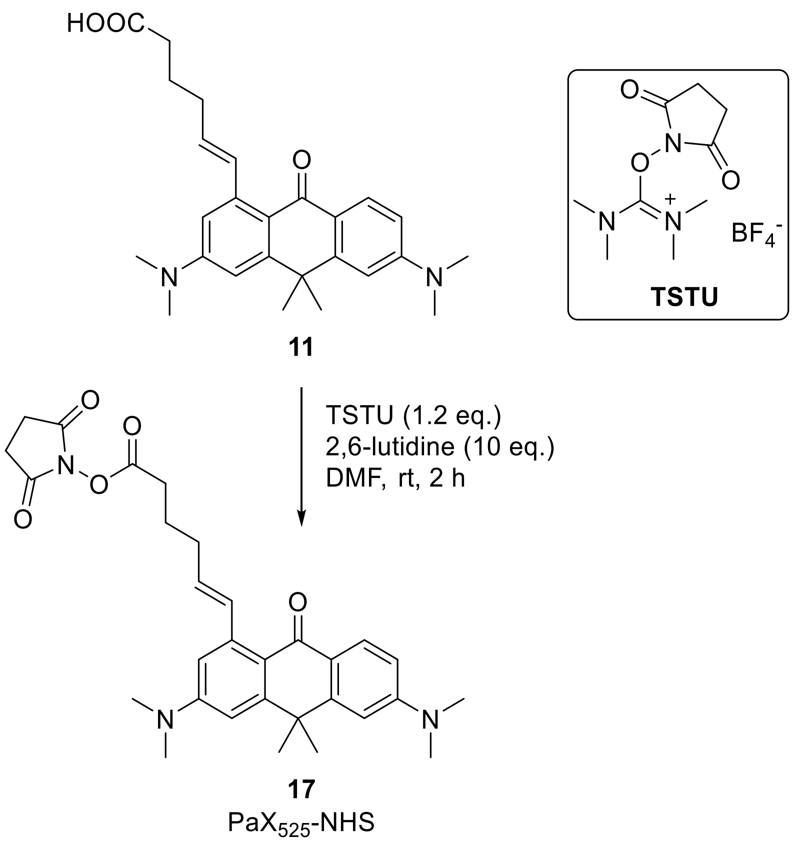

**Compound 17 ($PaX_{525}$-NHS).** In an amber vial, compound **11** (8.6 mg, 0.020 mmol) and TSTU (12 mg, 0.04 mmol) were dissolved in DMF (200 µL). 2,6-Lutidine (21 mg, 0.20 mmol) was added and the reaction mixture was stirred for 3 h at rt. The volatiles were removed *in vacuo*. The product was isolated by flash chromatography on a Biotage Isolera system (12g Interchim SiHP 30 µm cartridge, gradient 20% to 80% ethyl acetate/hexane) and freeze-dried from dioxane to yield 9.2 mg (89%) of **17** as a pale yellow solid.

HRMS (ESI) m/z: $[M+H]^+$ Calcd for $C_{30}H_{35}N_3O_5$: 518.2649, found: 518.2647.

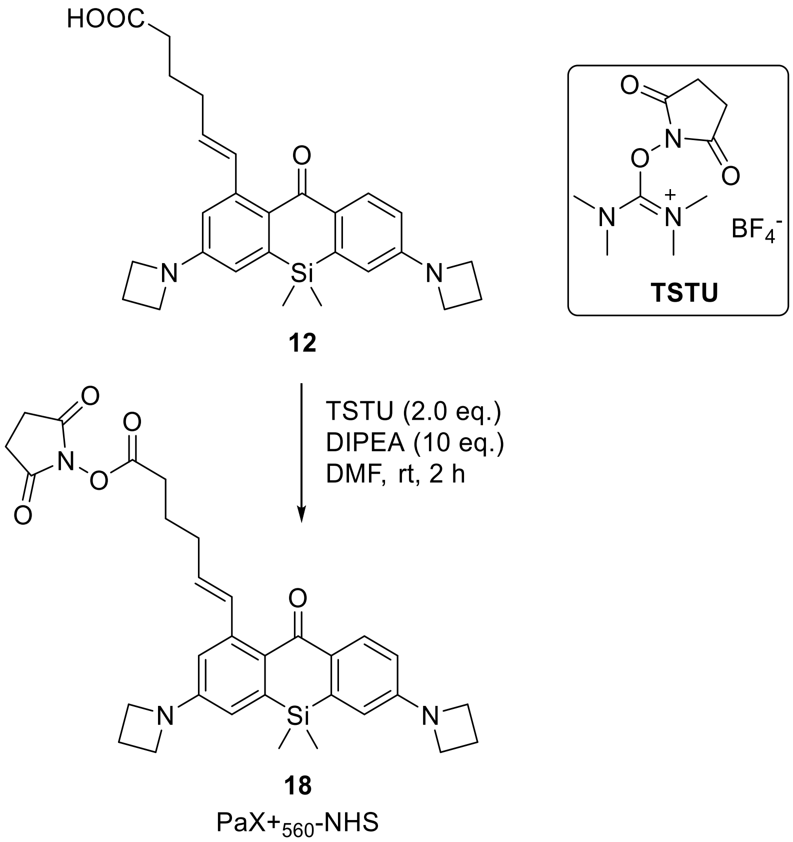

**Compound 18 ($PaX_{560+}$-NHS).** In an amber vial, compound **12** (11 mg, 0.023 mmol) and TSTU (14 mg, 0.046 mmol) were dissolved in DMF (200 µL). DIPEA (30 mg, 0.23 mmol) was added and the reaction mixture was stirred for 45 min at rt. The volatiles were removed *in vacuo*. The product was isolated by flash chromatography on a Biotage Isolera system (12g Interchim SiHP 30 µm cartridge, gradient 20% to 80% ethyl acetate/hexane) and freeze-dried from dioxane to yield 13 mg (98%) of **18** as a yellow solid.

$^{1}$H NMR (400 MHz, DMSO-$d_6$) δ 7.95 (d, *J* = 8.7 Hz, 1H), 7.09 (d, *J* = 15.6 Hz, 1H), 6.56 (d, *J* = 2.5 Hz, 1H), 6.54 (d, *J* = 2.5 Hz, 1H), 6.50 (dd, *J* = 8.7, 2.5 Hz, 1H), 6.42 (d, *J* = 2.5 Hz, 1H), 5.85 (dt, *J* = 15.5, 6.8 Hz, 1H), 3.98 (t, *J* = 7.4 Hz, 4H), 3.95 (t, *J* = 7.3 Hz, 4H), 2.86 – 2.78 (s, 4H), 2.81 (t, *J* = 7.3 Hz, 2H), 2.36 (p, *J* = 7.2 Hz, 4H), 2.28 (q, *J* = 7.1 Hz, 2H), 1.83 (p, *J* = 7.4 Hz, 2H), 0.40 (s, 6H).

$^{13}$C NMR (101 MHz, DMSO-$d_6$) δ 186.9, 170.3, 169.2, 152.3, 151.8, 142.2, 138.3, 134.3, 131.9, 130.4, 127.8, 113.4, 112.8, 112.0, 111.7, 51.5, 51.5, 31.3, 29.7, 25.5, 24.2, 16.2, -1.2.

HRMS (ESI) m/z: $[M+H]^+$ Calcd for $C_{31}H_{35}N_3O_5Si$: 558.2419, found: 558.2416.

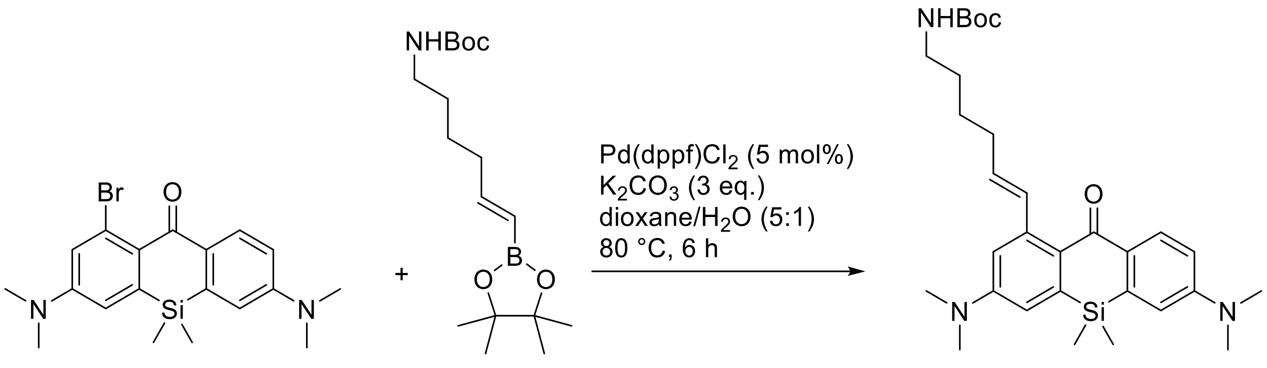


**Compound PaX$_{560}$-NHBoc.** In a 10 mL round-bottom flask, compound **11** (80 mg, 0.20 mmol), compound **S3** (88 mg, 0.26 mmol, 1.3 equiv, prepared according to literature procedure [18]), Pd(dppf)$Cl_2$·$CH_2Cl_2$ (8.2 mg, 0.01 mmol, 5 mol%) and $K_2CO_3$ (82 mg, 0.6 mmol, 3 equiv) were placed, dioxane (1.7 mL) and water (0.3 mL) were added, the mixture was degassed and stirred at 80 °C for 6 h. On cooling, the reaction mixture was diluted with ethyl acetate and poured into brine (50 mL), extracted with ethyl acetate (3 × 25 mL), and the combined extracts were dried over $Na_2SO_4$. The product was isolated by flash chromatography on Biotage Isolera system (12 g Interchim SiHP 30 µm cartridge, gradient 20% to 80% ethyl acetate/hexane) and freeze-dried from dioxane to give 101 mg (97%) of **PaX$_{560}$-NHBoc** as yellow solid.

$^1$H NMR (400 MHz, $CDCl_3$): δ 8.23 (d, *J* = 8.9 Hz, 1H), 7.25 (d, *J* = 15.5 Hz, 1H), 6.81 (dd, *J* = 9.0, 2.8 Hz, 1H), 6.78 – 6.73 (m, 3H), 5.88 (dt, *J* = 15.5, 6.8 Hz, 1H), 4.84 (br.s, 1H), 3.19 (q, *J* = 6.5 Hz, 2H), 3.09 (s, 6H), 3.07 (s, 6H), 2.36 – 2.27 (m, 2H), 1.68 – 1.53 (m, 4H), 1.44 (s, 9H), 0.45 (s, 6H).

$^{13}$C NMR (101 MHz, $CDCl_3$): δ 188.1, 156.3, 151.2, 150.8, 143.8, 141.3, 139.1, 134.8, 132.3, 131.4, 129.3, 128.4, 114.3, 113.9, 113.8, 113.4, 40.6, 40.2, 40.1, 32.6, 29.4, 28.6, 26.7, -0.8.

HRMS (ESI) m/z: $[M+H]^+$ Calcd for $C_{30}H_{43}N_3O_3Si$ 522.3146; Found 522.3143.

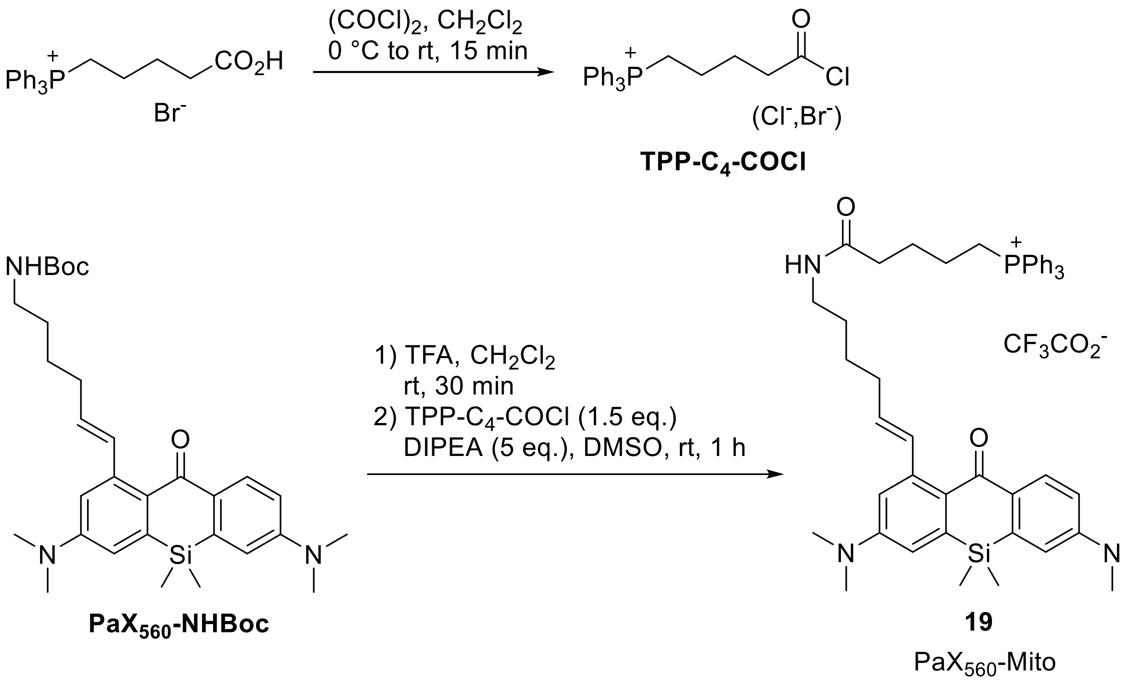


**Compound 19 (PaX$_{560}$-Mito).** A suspension of 4-(carboxybutyl)triphenylphosphonium bromide (44 mg, 0.1 mmol) in dry dichloromethane (1 mL) was cooled in ice-water bath, and oxalyl chloride (85 µL, 1.0 mmol, 10 equiv) was added. The mixture was allowed to warm up to rt and stirred for 15 min. The resulting clear light-yellow solution was evaporated to dryness and chased with dry dichloromethane (1 mL). The residue was dissolved in dry dichloromethane (1 mL) to give ~0.1 M solution of **TPP-C$_4$-COCl**.

Separately, in a 10 mL round-bottom flask, a solution of compound **PaX$_{560}$-NHBoc** (12 mg, 23 µmol) in dichloromethane (3 mL) and trifluoroacetic acid (300 µL) was stirred at rt for 30 min. The mixture was then diluted with toluene (2 mL), evaporated to dryness and chased twice with 3 mL toluene. Afterwards, the residue was dissolved in dry dichloromethane (1 mL), and DIPEA (30 µL) and the solution of **TPP-C$_4$-COCl** (350 µL of ~0.1 M in dichloromethane, ~35 µmol) were added. The reaction mixture was stirred at rt for 1 h, the solvents were evaporated and the product was isolated from the residue by preparative HPLC (12g Interchim Uptisphere Strategy PhC4 250×21.2 mm 5 µm, solvent flow rate 18 mL/min, gradient 40% to 90% A:B, A – acetonitrile + 0.1% (v/v) TFA, B – water + 0.1% (v/v) TFA) and freeze-dried from dioxane to give 20 mg (98%) of **19** (PaX$_{560}$-Mito) as light pink viscous oil (trifluoroacetate salt).

$^{1}$H NMR (400 MHz, DMSO-$d_6$): δ 7.97 (d, $J$ = 8.9 Hz, 1H), 7.92 – 7.85 (m, 3H), 7.82 – 7.71 (m, 12H), 7.16 (dt, $J$ = 15.5, 1.5 Hz, 1H), 6.87 (d, $J$ = 2.7 Hz, 1H), 6.85 (d, $J$ = 2.8 Hz, 1H), 6.82 (dd, $J$ = 9.0, 2.7 Hz, 1H), 6.71 (d, $J$ = 2.8 Hz, 1H), 5.87 (dt, $J$ = 15.5, 6.8 Hz, 1H), 3.62 – 3.51 (m, 2H), 3.05 (s, 6H), 3.03 (s, 6H), 2.21 – 2.07 (m, 4H), 1.71 (p, $J$ = 7.3 Hz, 2H), 1.58 – 1.46 (m, 2H), 1.45 – 1.39 (m, 4H), 0.43 (s, 6H).

$^{19}$F NMR (376 MHz, DMSO-$d_6$): δ -74.5.

$^{31}$P NMR (162 MHz, DMSO-$d_6$): δ 23.9.

$^{13}$C NMR (101 MHz, DMSO-$d_6$): δ 186.7, 171.3, 158.1 (q, $^{2}J_{C\text{-}F}$ = 35.1 Hz), 151.0, 150.5, 142.5, 140.8, 138.5, 134.92, 134.89, 133.64, 133.59, 133.5, 131.1, 130.5, 130.3, 130.2, 128.1 (q, $^{1}J_{C\text{-}F}$ = 213.7 Hz), 118.9, 118.1, 117.5, 114.6, 114.5, 114.0, 113.15, 112.7, 38.3, 34.4, 32.2, 30.4, 28.8, 26.31, 26.25, 26.1, 21.4, 21.3, 20.2, 19.7, -1.1 (C–P multiplets were not interpreted).

HRMS (ESI) m/z: $[M+H]^{2+}$ Calcd for $C_{48}H_{58}N_3O_2PSi$ 383.7012; Found 383.7008.

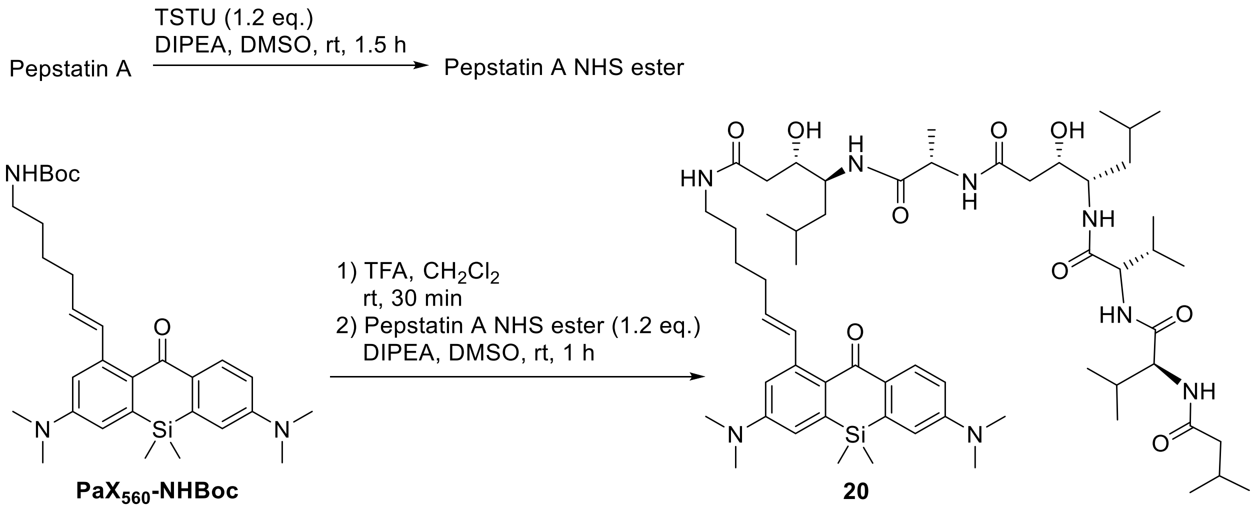


**PaX560-NHBoc** **20** PaX560-Lyso

**Compound 20 (PaX560-Lyso).** To a solution of Pepstatin A (19 mg, 27.6 µmol, 1.2 equiv) in the mixture of DIPEA (40 µL) and DMSO (800 µL), a solution of TSTU (*N*,*N*,*N′*,*N′*-tetramethyl-*O*-(*N*-succinimidyl)uronium tetrafluoroborate; 10 mg, 33 µmol, 1.2 equiv relative to Pepstatin A) in DMSO (100 µL) was added, and the reaction mixture was stirred at rt for 1.5 h to give the solution of Pepstatin A NHS ester in DMSO.

Separately, in a 10 mL round-bottom flask, a solution of compound **PaX560-NHBoc** (12 mg, 23 µmol) in dichloromethane (3 mL) and trifluoroacetic acid (300 µL) was stirred at rt for 30 min. The mixture was then diluted with toluene (2 mL), evaporated to dryness and chased twice with 3 mL toluene. Afterwards, the solution of Pepstatin A NHS ester in DMSO was added to the dry residue followed by additional DIPEA (50 µL). The reaction mixture was stirred at rt for 1 h, excess DIPEA was evaporated and the product was isolated from the crude mixture by preparative HPLC (Interchim Uptisphere Strategy PhC4 250×21.2 mm 5 µm, solvent flow rate 18 mL/min, gradient 40% to 90% A:B, A – acetonitrile + 0.1% (v/v) formic acid, B – water + 0.1% (v/v) formic acid) and freeze-dried from dioxane-water to give 13 mg (52%) of **20** (PaX560-Lyso) as light yellow solid.

$^1$H NMR (400 MHz, DMSO-*d*6): δ 8.00 – 7.89 (m, 2H), 7.85 – 7.71 (m, 2H), 7.47 (d, *J* = 8.8 Hz, 1H), 7.33 (d, *J* = 9.1 Hz, 1H), 7.17 (dt, *J* = 15.5, 1.4 Hz, 1H), 6.88 – 6.81 (m, 3H), 6.72 (d, *J* = 2.7 Hz, 1H), 5.88 (dt, *J* = 15.5, 6.8 Hz, 1H), 4.87 (d, *J* = 5.1 Hz, 1H), 4.83 (d, *J* = 4.9 Hz, 1H), 4.25 (p, *J* = 7.1 Hz, 1H), 4.18 (dd, *J* = 8.8, 7.3 Hz, 1H), 4.13 (dd, *J* = 8.9, 7.2 Hz, 1H), 3.88 – 3.73 (m, 4H), 3.15 – 3.04 (m, 1H), 3.05 (s, 6H), 3.03 (s, 6H), 2.19 (q, *J* = 6.8 Hz, 2H), 2.15 –

2.00 (m, 5H), 2.01 – 1.87 (m, 3H), 1.60 – 1.43 (m, 6H), 1.41 – 1.30 (m, 1H), 1.28 – 1.17 (m, 5H), 0.89 – 0.75 (m, 28H), 0.43 (s, 6H).

$^{13}$C NMR (101 MHz, DMSO-$d_6$): δ 186.7, 172.2, 171.6, 171.1, 170.8, 170.70, 170.65, 162.3, 151.0, 150.5, 142.5, 140.8, 138.4, 133.6, 131.1, 130.6, 129.2, 127.0, 114.4, 113.9, 113.1, 112.7, 69.2, 69.0, 58.0, 57.8, 50.7, 50.5, 48.3, 44.4, 38.6, 38.4, 35.8, 34.4, 32.3, 30.8, 30.4, 30.3, 30.1, 28.8, 26.4, 25.7, 24.2, 23.5, 23.3, 22.3, 21.9, 21.6, 21.1, 19.30, 19.25, 18.4, 18.3, 18.2, -1.1.

HRMS (ESI) m/z: $[M+2H]^{2+}$ Calcd for $C_{59}H_{96}N_8O_9Si$ 545.3608; Found 545.3599.

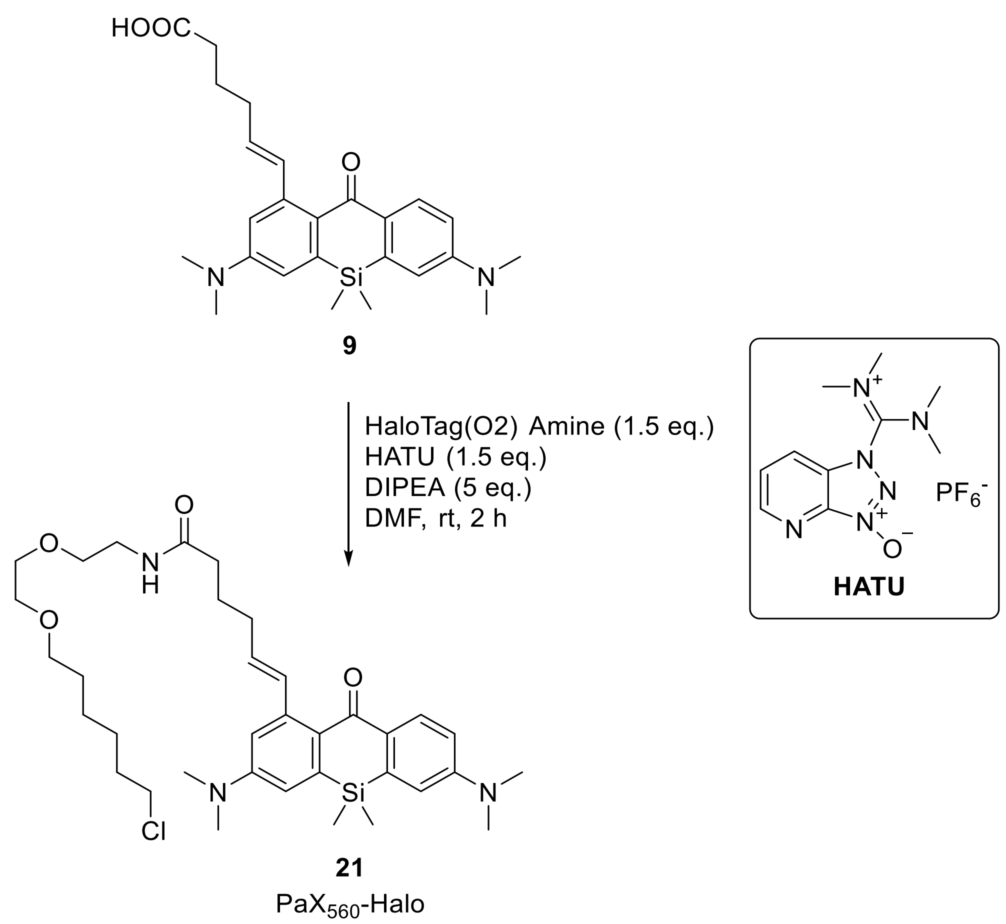


**Compound 21 (PaX$_{560}$-Halo).** In an amber vial, compound **9** (22 mg, 0.050 mmol), HaloTag(O2) Amine[19] (17 mg, 0.075 mmol, 1.5 equiv), and HATU (1-[bis(dimethylamino)methylene]-1*H*-1,2,3-triazolo[4,5-*b*]pyridinium 3-oxid hexafluorophosphate; 29 mg, 0.075 mmol, 1.5 equiv) were dissolved in DMF (200 µL). DIPEA (32 mg, 0.25 mmol, 5 equiv) was added and the mixture stirred for 2 h at rt. The volatiles were removed *in vacuo*. The product was isolated by flash chromatography on a Biotage Isolera system (12g Interchim SiHP 30 µm cartridge, 100% ethyl acetate) and freeze-dried from dioxane to yield **21** as a yellow solid. Yield 33 mg (84%, remainder dioxane).

$^1$H NMR (400 MHz, DMSO-*d*$_6$): δ 7.99 (d, *J* = 8.9 Hz, 1H), 7.92 (t, *J* = 5.6 Hz, 1H), 7.16 – 7.09 (m, 1H), 6.89 – 6.79 (m, 3H), 6.71 (d, *J* = 2.8 Hz, 1H), 5.84 (dt, *J* = 15.5, 6.9 Hz, 1H), 3.57 (t, *J* = 6.6 Hz, 2H), 3.48 – 3.44 (m, 2H), 3.44 – 3.39 (m, 4H), 3.31 – 3.27 (m, 4H), 3.24 (q, *J* = 5.8 Hz, 2H), 3.06 (s, 6H), 3.03 (s, 6H), 2.22 – 2.13 (m, 4H), 1.75 – 1.60 (m, 4H), 1.46 – 1.37 (m, 2H), 1.37 – 1.10 (m, 6H), 0.43 (s, 6H).

$^{13}C$ NMR (101 MHz, DMSO-$d_6$): δ 186.7, 172.2, 151.1, 150.6, 142.7, 140.9, 138.5, 134.4, 131.0, 130.6, 128.5, 126.9, 114.4, 113.9, 113.1, 112.9, 70.2, 69.6, 69.4, 69.2, 45.3, 38.6, 34.7, 32.0, 31.9, 29.0, 26.1, 25.1, 24.9, -1.1.

HRMS (ESI) m/z: $[M+H]^+$ Calcd for $C_{35}H_{52}ClN_3O_4Si$: 642.3488, found: 642.3487.

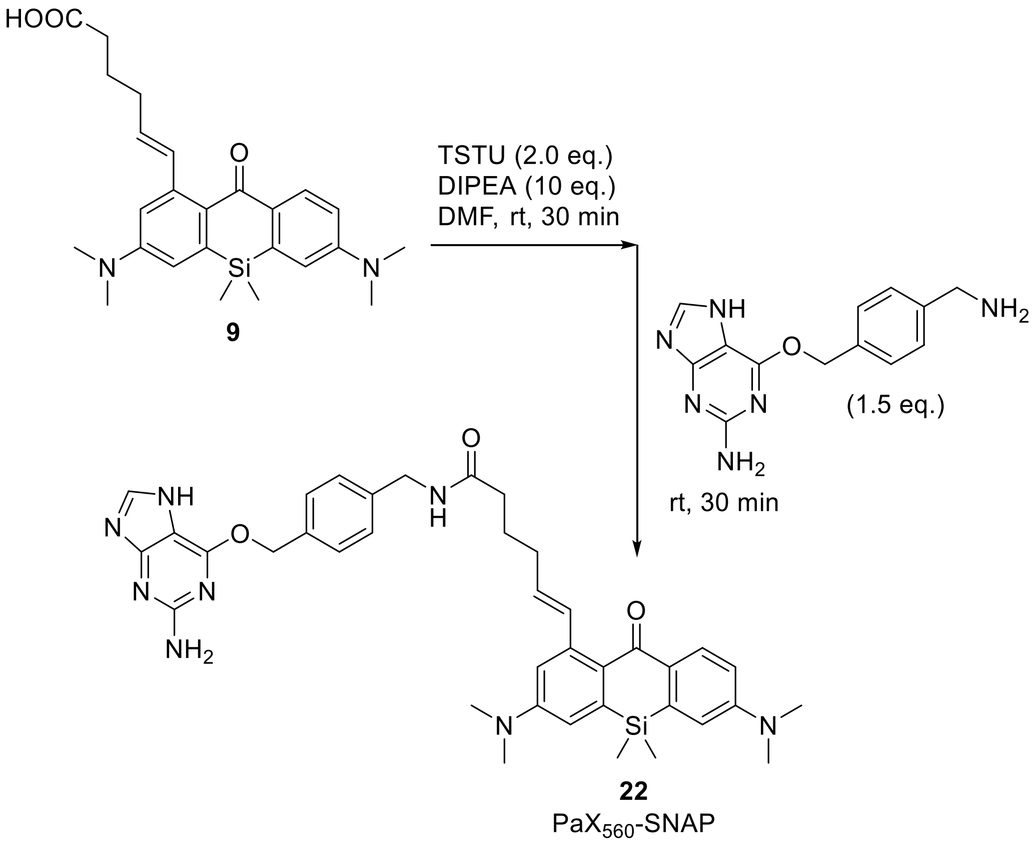


**Compound 22 (PaX$_{560}$-SNAP).** In an amber vial, compound **9** (10.4 mg, 0.024 mmol) and TSTU (14 mg, 0.048 mmol, 2 equiv) were dissolved in DMF (240 µL). DIPEA (31 mg, 0.24 mmol) was added, and the mixture was stirred for 30 min at rt. 6-((4-(Aminomethyl)benzyl)oxy)-7*H*-purin-2-amine[20] (20.0 mg, 0.036 mmol, 1.5 equiv) was then added, and the reaction mixture was stirred at rt for 30 min. The volatiles were removed *in vacuo*. The product was isolated by flash chromatography on a Biotage Isolera system (12g Interchim SiHP 30 µm cartridge, gradient 0% to 10% methanol/dichloromethane) and freeze-dried from dioxane to yield 15.1 mg (92%) of **22** as a yellow solid.

HRMS (ESI) m/z: $[M+H]^+$ Calcd for $C_{38}H_{44}N_8O_3Si$: 689.3378, found: 689.3378.

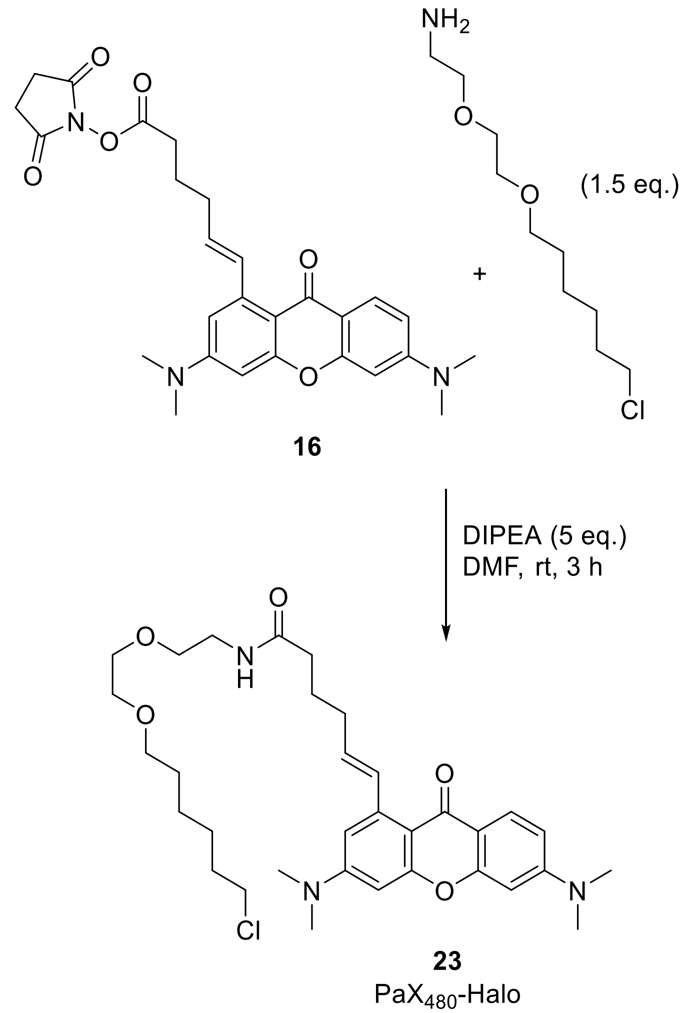


**Compound 23 (PaX480-Halo).** In an amber vial, compound **16** (9.9 mg, 0.02 mmol) and HaloTag(O2) Amine (6.7 mg, 0.03 mmol) were dissolved in DMF (100 µL). DIPEA (*N*,*N*-diisopropylethylamine; 12.9 mg, 0.1 mmol) was added and stirred for 3 hours at rt. The volatiles were removed *in vacuo*. The product was isolated by flash chromatography on a Biotage Isolera system (12g Interchim SiHP 30 µm cartridge, gradient 0% to 100% A/B, A: 100% ethyl acetate, B: 2% methanol-98% ethyl acetate) and freeze-dried from dioxane to yield 9.7 mg (81%) of **23** as an off-white solid.

$^1$H NMR (400 MHz, DMSO-$d_6$) δ 7.93 (t, *J* = 5.7 Hz, 1H), 7.84 (d, *J* = 9.0 Hz, 1H), 7.79 (d, *J* = 15.7 Hz, 1H), 6.76 (dd, *J* = 9.1, 2.4 Hz, 1H), 6.67 (d, *J* = 2.5 Hz, 1H), 6.48 (t, *J* = 2.8 Hz, 2H), 6.06 (dt, *J* = 15.6, 6.8 Hz, 1H), 3.58 (t, *J* = 6.6 Hz, 2H), 3.52 – 3.38 (m, 6H), 3.31 (t, *J* = 6.6 Hz, 2H), 3.22 (q, *J* = 5.8 Hz, 2H), 3.07 (s, 6H), 3.05 (s, 6H), 2.24 – 2.13 (m, 4H), 1.77 – 1.61 (m, 4H), 1.43 (dt, *J* = 14.2, 6.8 Hz, 2H), 1.39 – 1.18 (m, 4H).

$^{13}C$ NMR (101 MHz, DMSO-$d_6$) δ 174.9, 172.1, 158.5, 156.5, 154.1, 152.8, 141.2, 131.4, 131.4, 127.0, 111.7, 109.2, 107.9, 107.1, 96.6, 96.0, 70.2, 69.5, 69.4, 69.1, 45.3, 39.6, 38.5, 34.7, 32.0, 32.0, 29.0, 26.1, 24.9, 24.9.

HRMS (ESI) m/z: $[M+H]^+$ Calcd for $C_{33}H_{46}ClN_3O_5$: 600.3199, found: 600.3202.

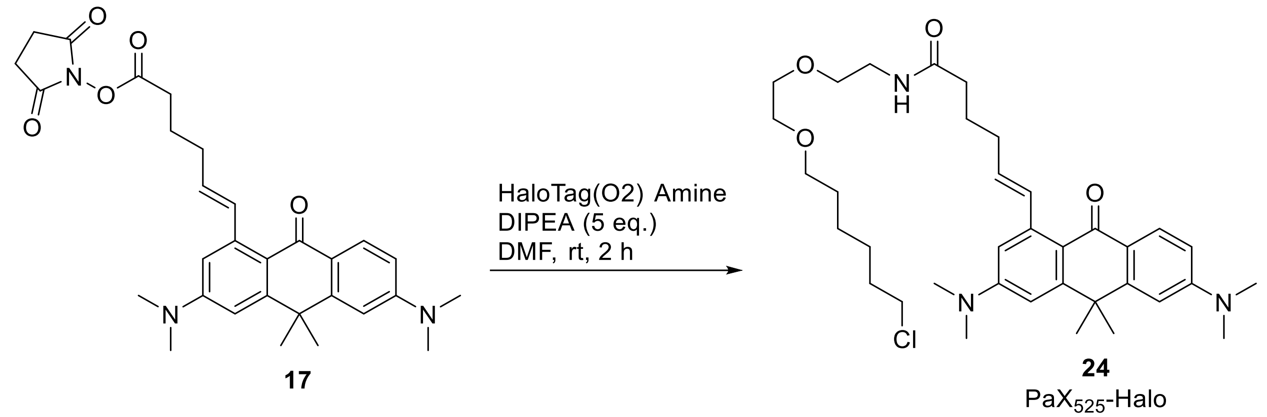

**Compound 24 ($PaX_{525}$-Halo).** To a solution of compound **17** (7.2 mg, 0.014 mmol) in DMF (100 µL) was added HaloTag(O2) Amine (4.7 mg, 0.021 mmol) and DIPEA (9.0 mg, 0.070 mmol), and the mixture stirred for 2 h at rt. The volatiles were removed *in vacuo*. The product was isolated by flash chromatography on a Biotage Isolera system (12g Interchim SiHP 30 µm cartridge, 100% ethyl acetate) and freeze-dried from dioxane to yield 5.2 mg (59%) of **17** as an orange oil.

HRMS (ESI) m/z: $[M+H]^+$ Calcd for $C_{36}H_{52}ClN_3O_4$: 626.3719, found: 626.3714.

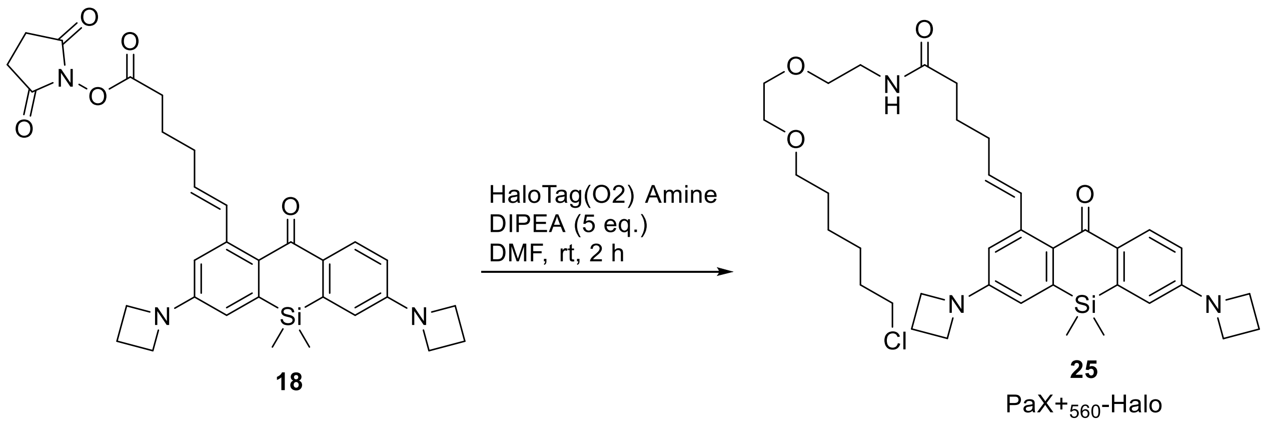

**Compound 25 (PaX+560-Halo).** To a solution of compound **18** (12 mg, 0.022 mmol) in DMF (100 µL) was added HaloTag(O2) Amine (6.8 mg, 0.030 mmol) and DIPEA (14 mg, 0.11 mmol), and the mixture stirred for 2 h at rt. The volatiles were removed *in vacuo*. The product was isolated by flash chromatography on a Biotage Isolera system (12g Interchim SiHP 30 µm cartridge, gradient 0% to 5% methanol/dichloromethane) and freeze-dried from dioxane to yield 7.7 mg (54%) of **25** as an yellow oil.

$^{1}$H NMR (400 MHz, DMSO-$d_6$) δ 7.95 (d, *J* = 8.7 Hz, 1H), 7.93 (t, *J* = 5.8 Hz, 1H), 7.06 (d, *J* = 15.5 Hz, 1H), 6.55 (dd, *J* = 10.4, 2.5 Hz, 2H), 6.50 (dd, *J* = 8.7, 2.5 Hz, 1H), 6.41 (d, *J* = 2.5 Hz, 1H), 5.81 (dt, *J* = 15.4, 6.9 Hz, 1H), 3.98 (t, J = 7.3 Hz, 4H), 3.95 (t, J = 7.3 Hz, 4H), 3.58 (t, *J* = 6.3 Hz, 2H), 3.47 (dd, *J* = 5.7, 3.3 Hz, 2H), 3.45 – 3.37 (m, 4H), 3.29 (t, *J* = 6.6 Hz, 2H), 3.23 (q, *J* = 5.8 Hz, 2H), 2.36 (p, *J* = 7.2 Hz, 4H), 2.16 (p, *J* = 7.2 Hz, 4H), 1.76 – 1.60 (m, 4H), 1.42 (p, *J* = 6.8 Hz, 2H), 1.38 – 1.16 (m, 4H), 0.40 (s, 6H).

$^{13}$C NMR (101 MHz, DMSO-$d_6$): δ 187.0, 172.2, 152.3, 151.8, 142.5, 140.6, 138.3, 133.9, 131.9, 130.4, 128.8, 127.7, 113.3, 112.8, 112.0, 111.6, 70.2, 69.6, 69.4, 69.2, 51.5, 51.5, 38.6, 34.7, 32.0, 29.1, 26.2, 25.1, 24.9, 16.2, -1.2.

HRMS (ESI) m/z: $[M+H]^+$ Calcd for $C_{37}H_{52}ClN_3O_4Si$: 666.3488, found: 666.3485.

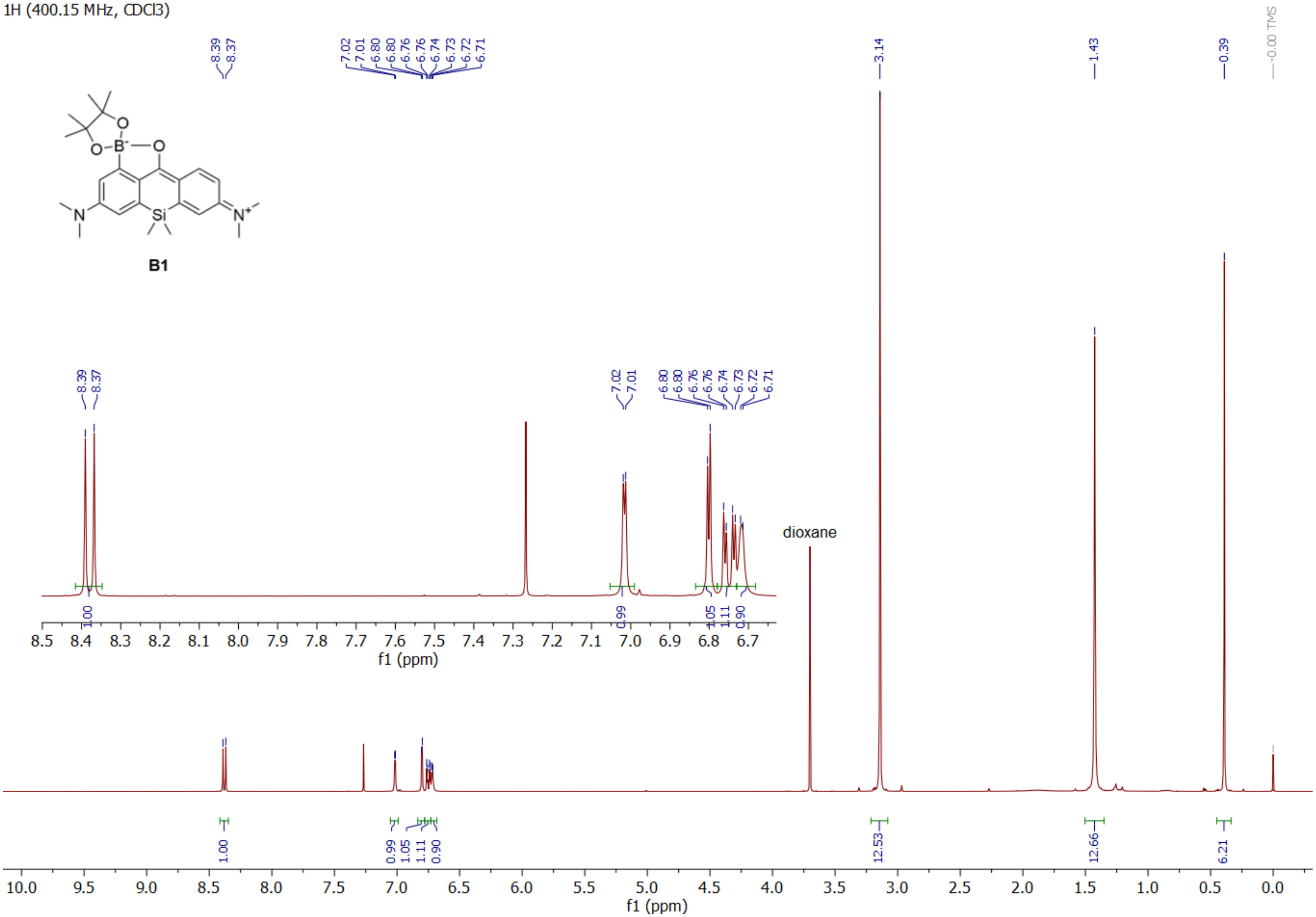
1H (400.15 MHz, CDCl3)
B1
8.39
8.37
7.02
7.01
6.80
6.80
6.76
6.76
6.74
6.73
6.72
6.71
3.14
1.43
0.39
-0.00 TMS
dioxane
1.00
0.99
1.05
1.11
0.90
12.53
12.66
6.21
f1 (ppm)

13C (100.63 MHz, CDCl3)

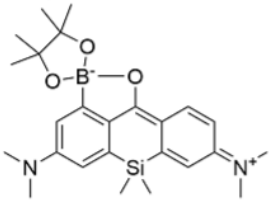

B1

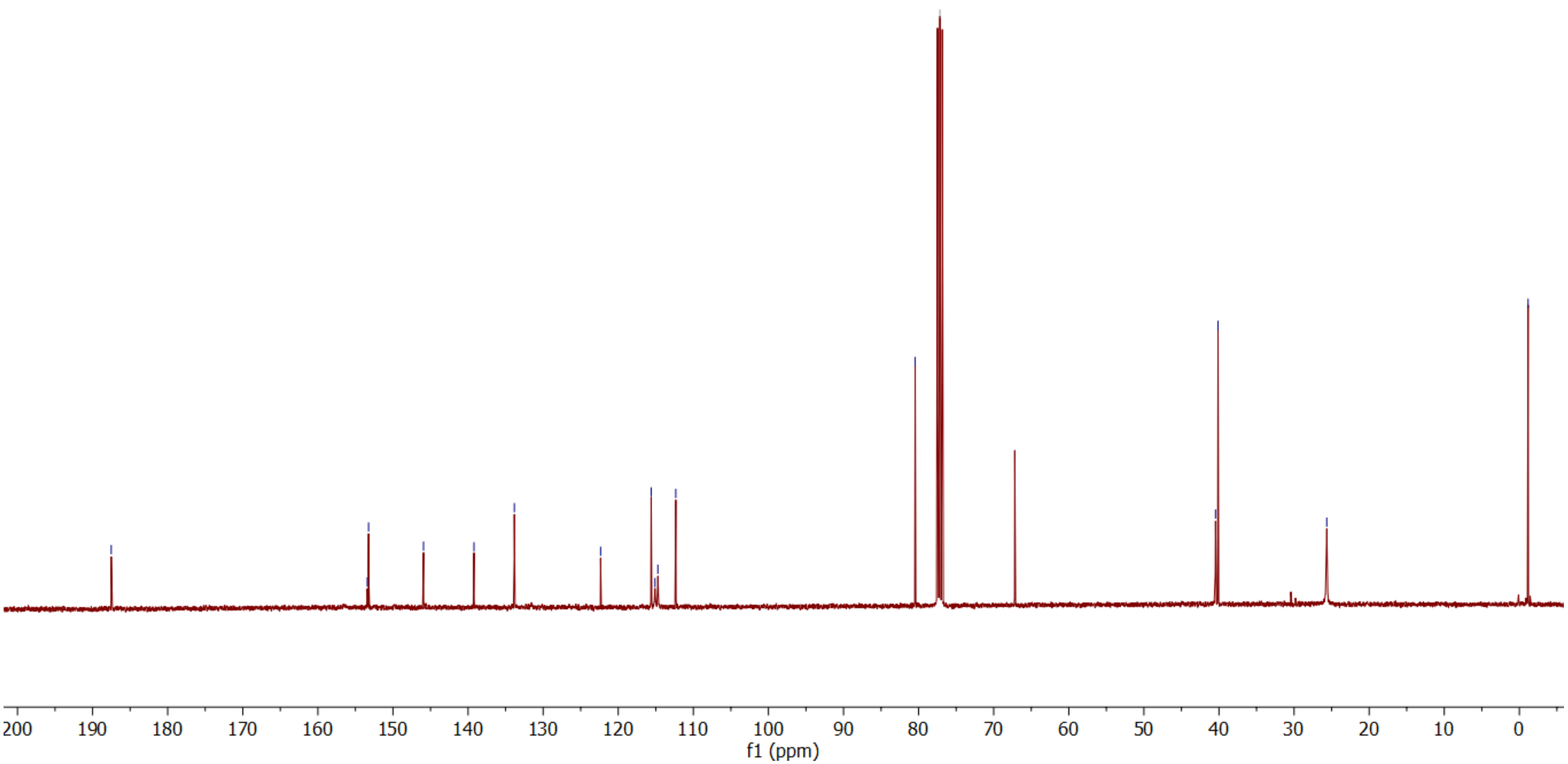

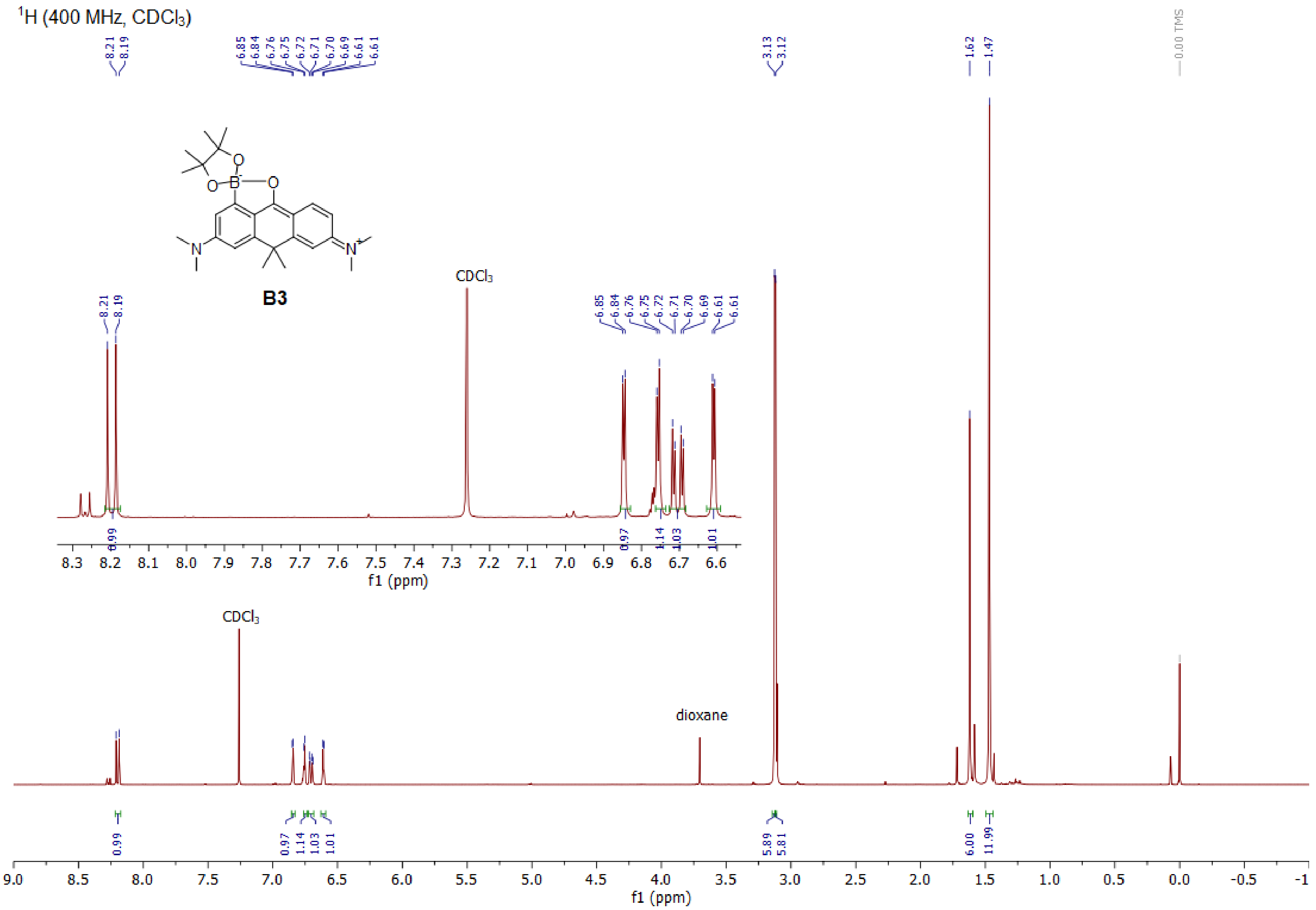

1H (400 MHz, CDCl3)
B3
CDCl3
CDCl3
dioxane
0.00 TMS
f1 (ppm)
f1 (ppm)

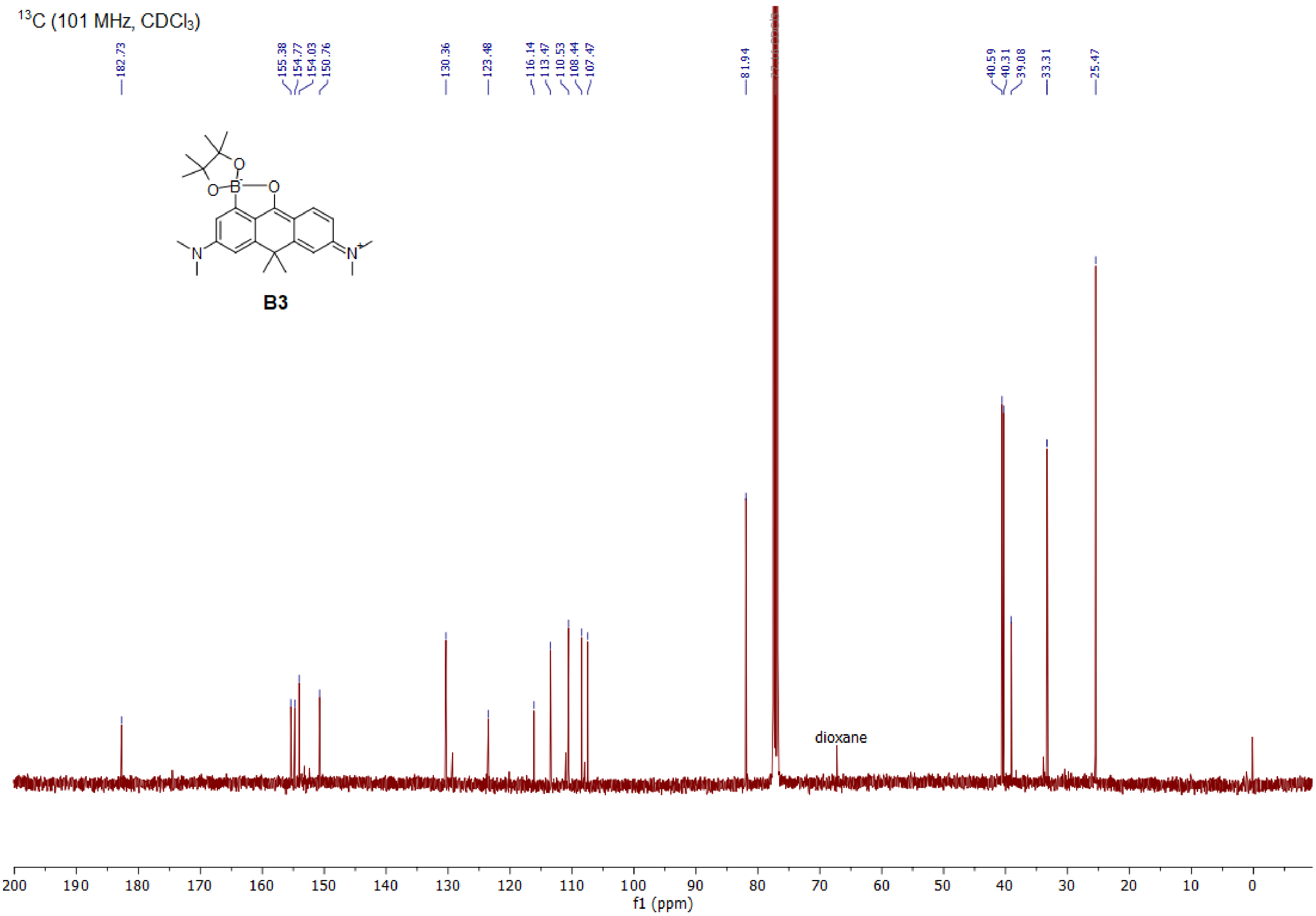

13C (101 MHz, CDCl3)
182.73
155.38
154.77
154.03
150.76
130.36
123.48
116.14
113.47
110.53
108.44
107.47
81.94
40.59
40.31
39.08
33.31
25.47
B3
dioxane
200 190 180 170 160 150 140 130 120 110 100 90 80 70 60 50 40 30 20 10 0
f1 (ppm)

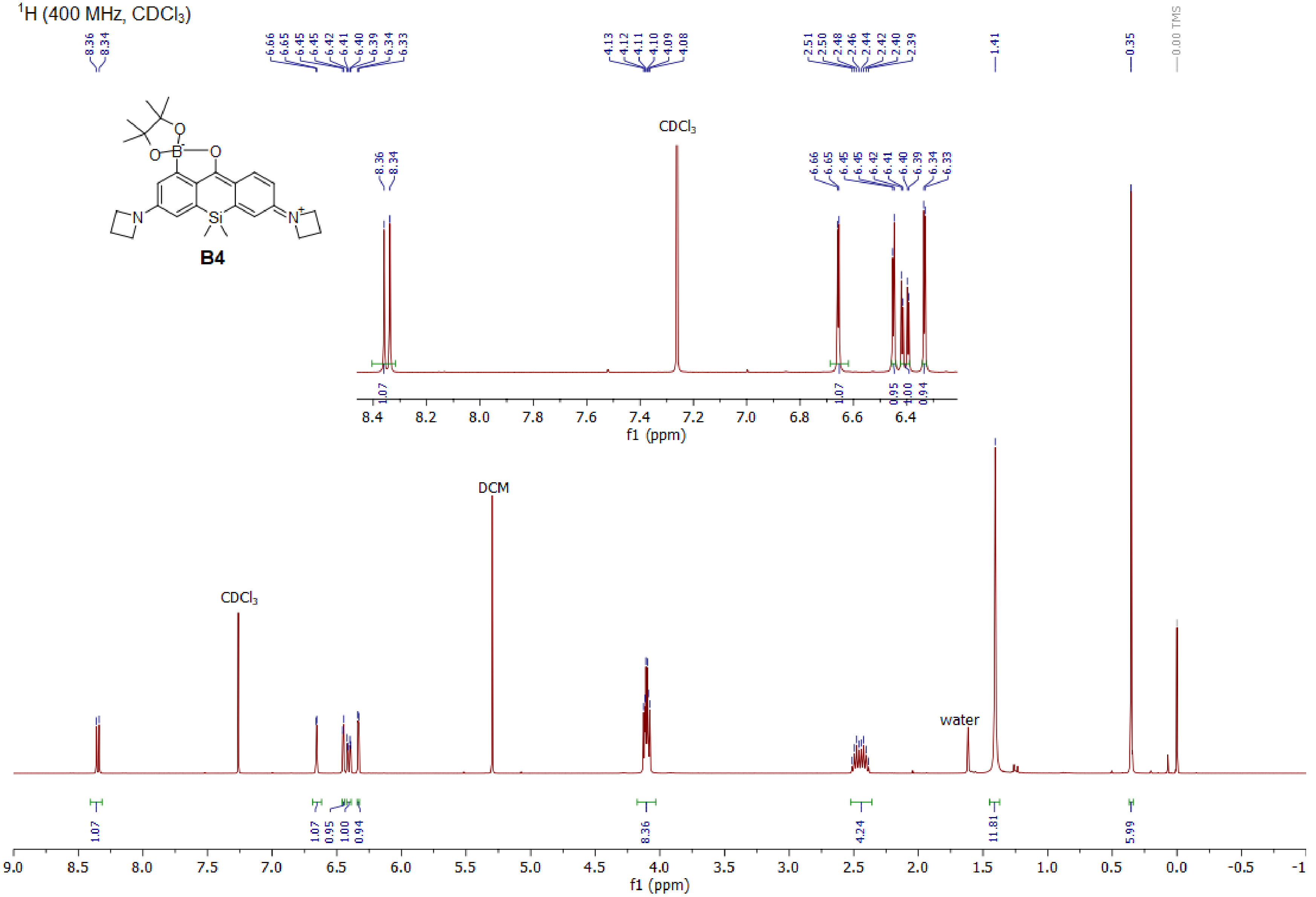

1H (400 MHz, CDCl3)
B4
8.36
8.34
6.66
6.65
6.45
6.45
6.42
6.41
6.40
6.39
6.34
6.33
4.13
4.12
4.11
4.10
4.09
4.08
2.51
2.50
2.48
2.46
2.44
2.42
2.40
2.39
1.41
0.35
0.00 TMS
CDCl3
DCM
water
1.07
1.07
0.95
1.00
0.94
8.36
4.24
11.81
5.99
f1 (ppm)

$^{13}C$ (101 MHz, $CDCl_3$)

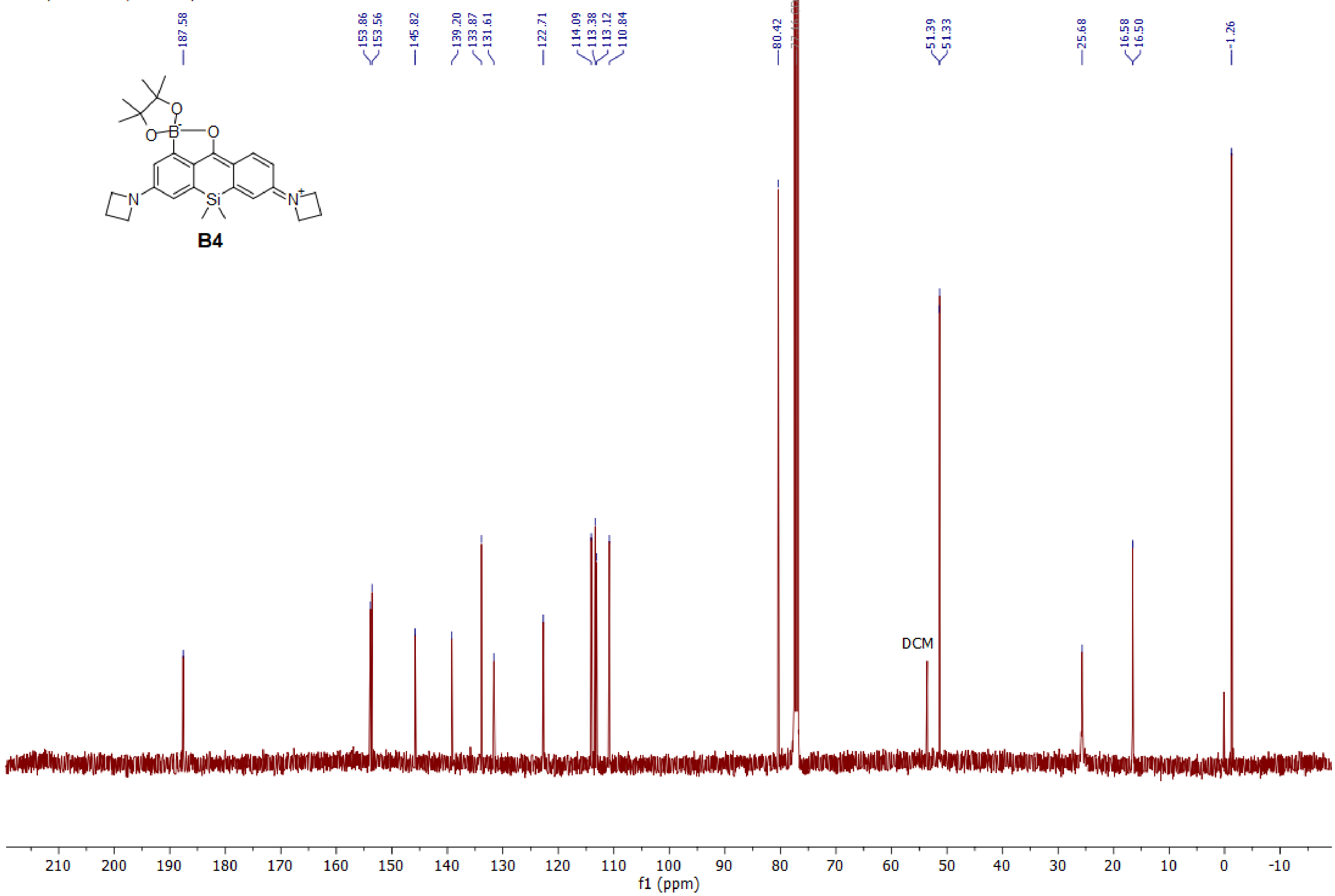

1H (400.15 MHz, CDCl3)

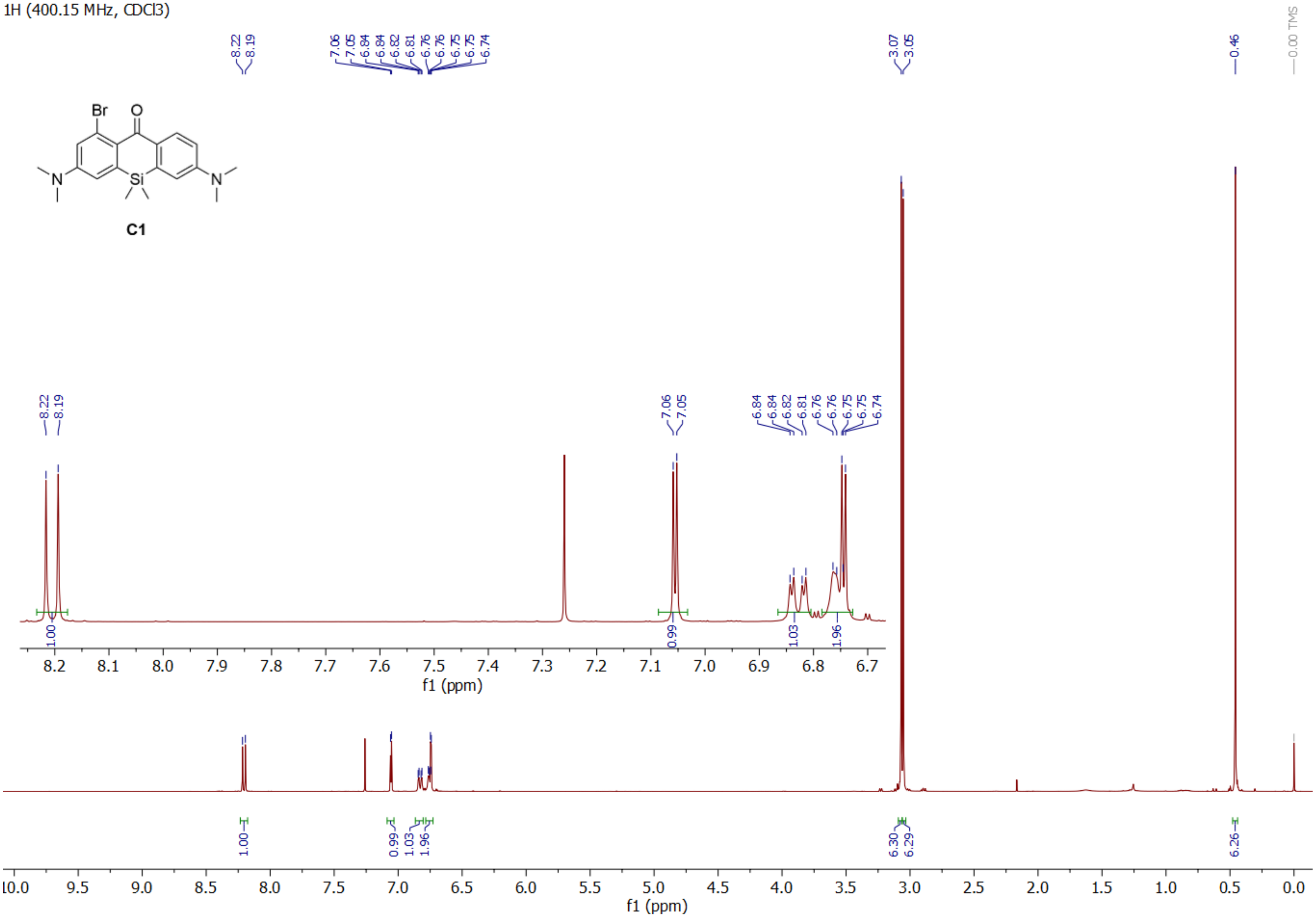

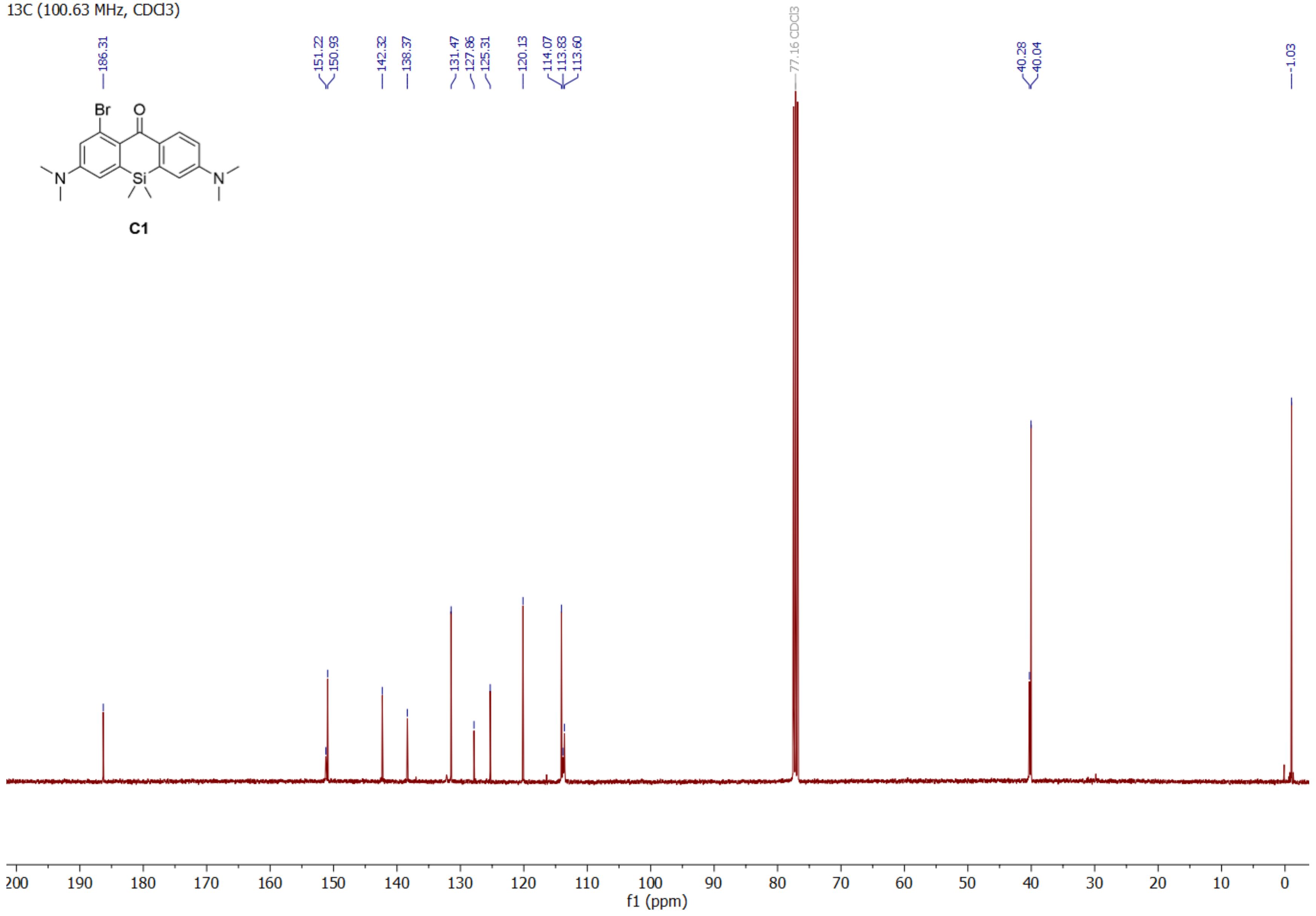
13C (100.63 MHz, CDCl3)
186.31
151.22
150.93
142.32
138.37
131.47
127.86
125.31
120.13
114.07
113.83
113.60
77.16 CDCl3
40.28
40.04
-1.03
Br
O
N
Si
N
C1
200
190
180
170
160
150
140
130
120
110
100
90
80
70
60
50
40
30
20
10
0
f1 (ppm)

1H (400.15 MHz, CDCl3)

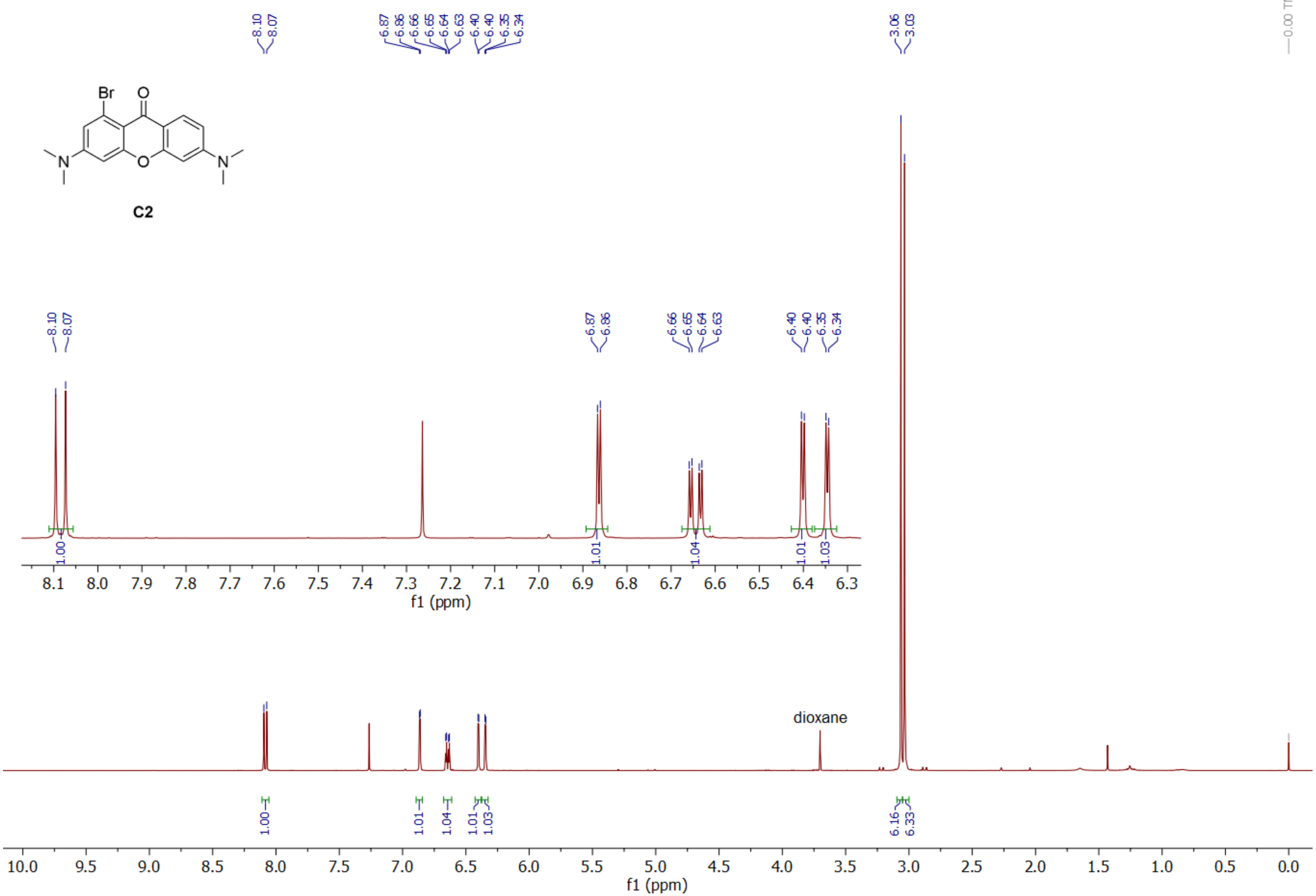

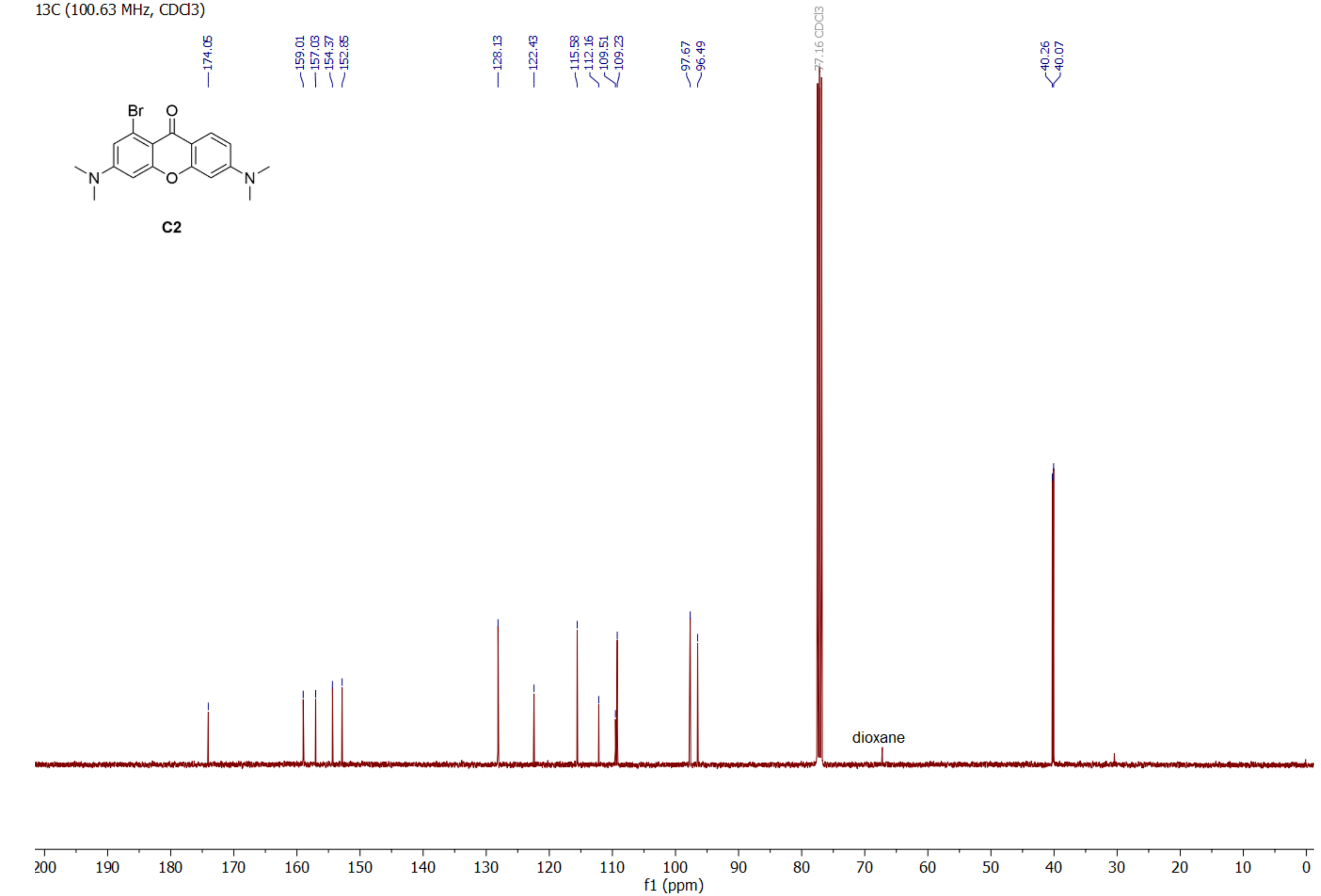
13C (100.63 MHz, CDCl3)
174.05
159.01
157.03
154.37
152.85
128.13
122.43
115.58
112.16
109.51
109.23
97.67
96.49
77.16 CDCl3
40.26
40.07
Br
O
N
O
N
C2
dioxane
200
190
180
170
160
150
140
130
120
110
100
90
80
70
60
50
40
30
20
10
0
f1 (ppm)

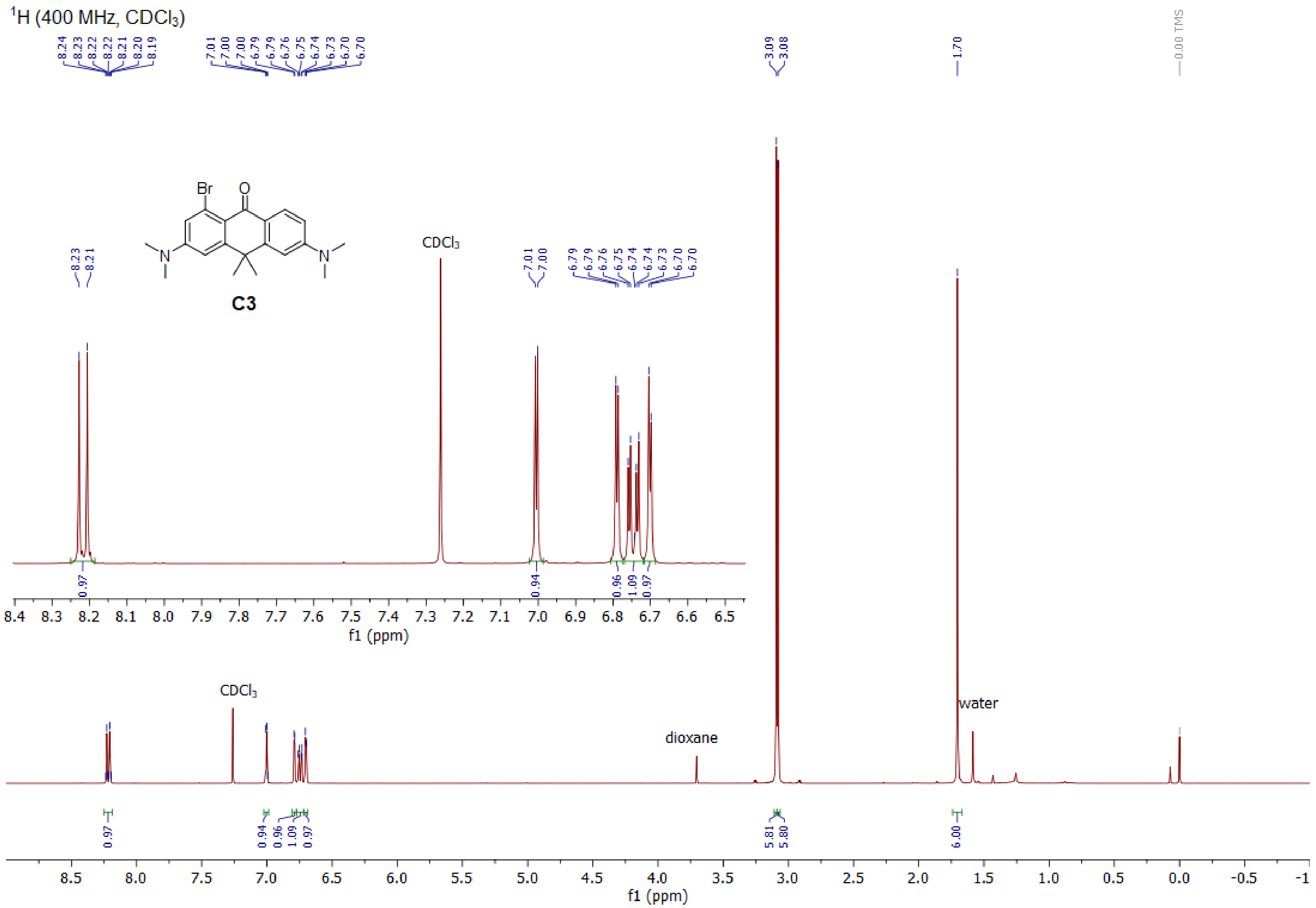

$^1$H (400 MHz, CDCl$_3$)
Br
O
N
N
C3
CDCl$_3$
dioxane
water
f1 (ppm)

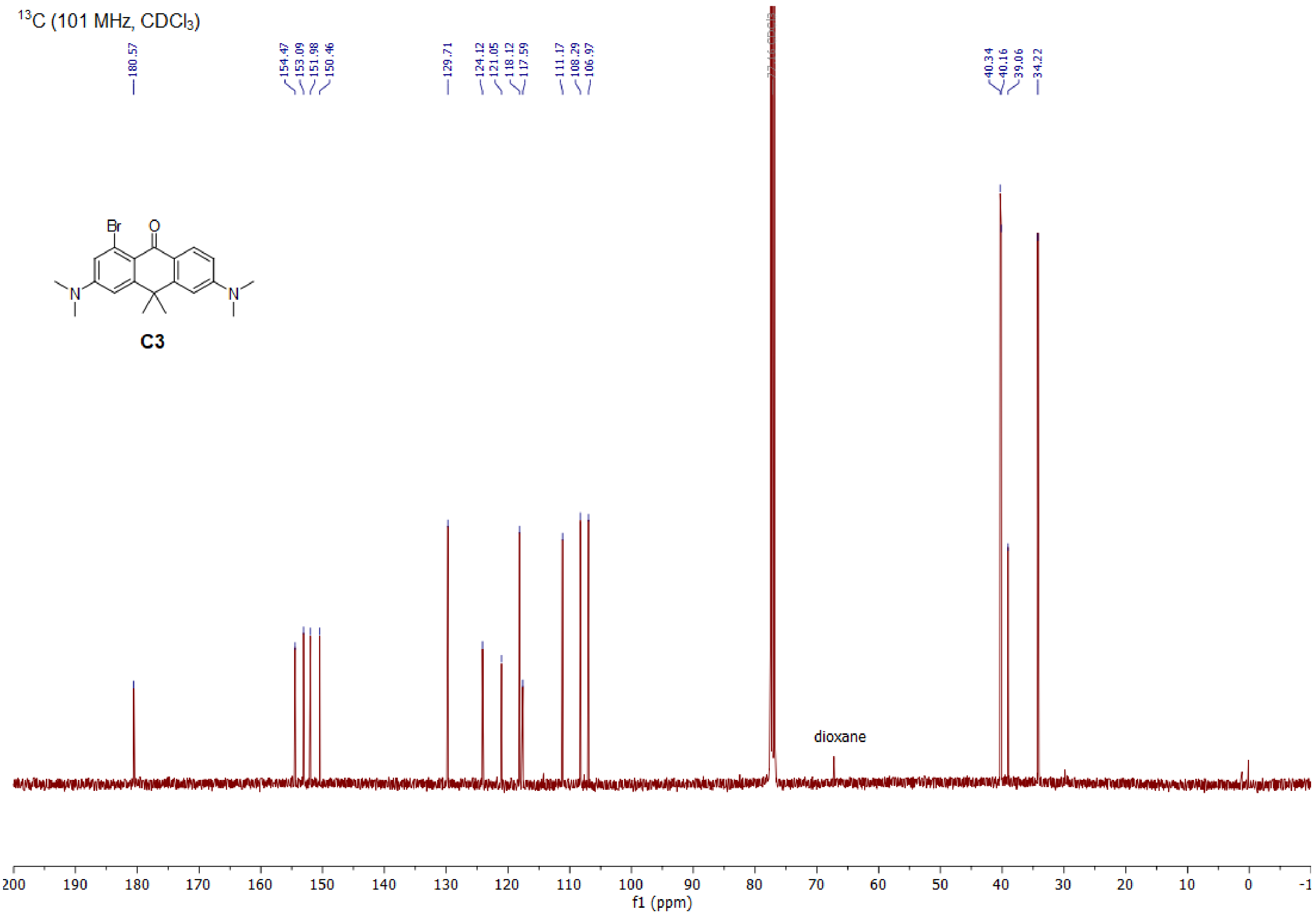
13C (101 MHz, CDCl3)
180.57
154.47
153.09
151.98
150.46
129.71
124.12
121.05
118.12
117.59
111.17
108.29
106.97
40.34
40.16
39.06
34.22
Br
O
N
N
C3
dioxane
200 190 180 170 160 150 140 130 120 110 100 90 80 70 60 50 40 30 20 10 0 -1
f1 (ppm)

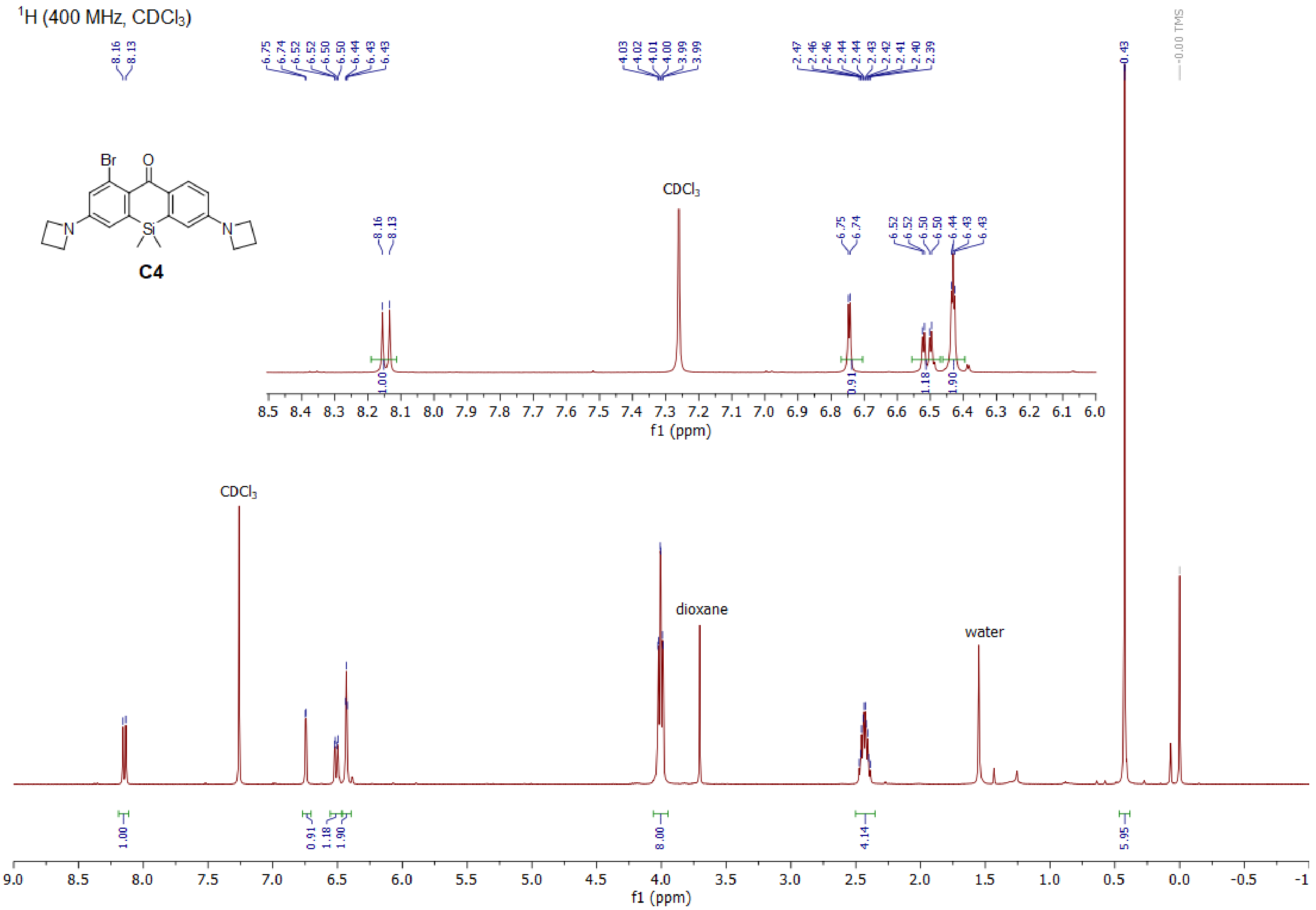

¹H (400 MHz, CDCl3)
C4
CDCl3
dioxane
water
0.00 TMS
f1 (ppm)

$^{13}C$ (101 MHz, $CDCl_3$)

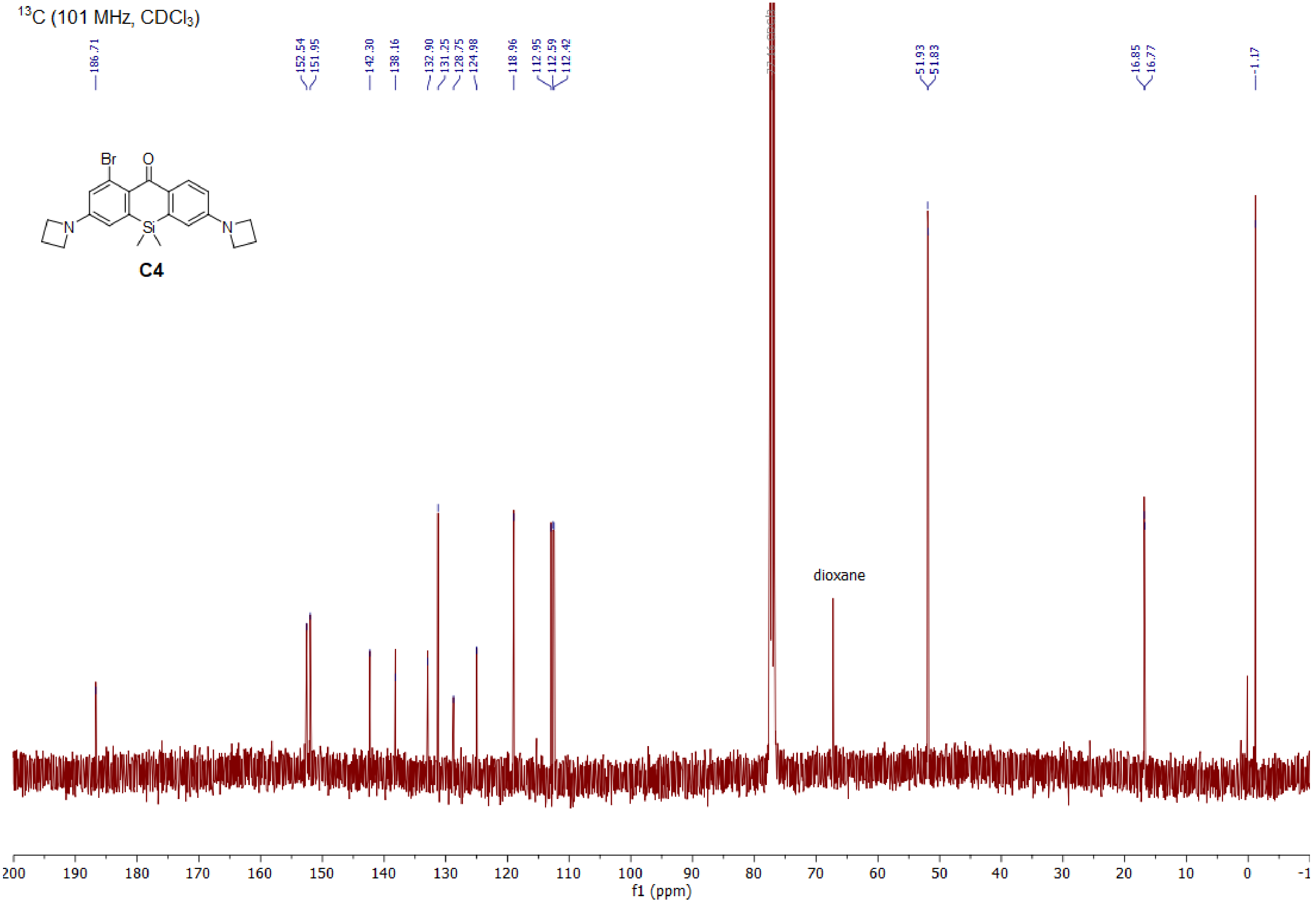

$^1$H (400 MHz, CDCl$_3$)

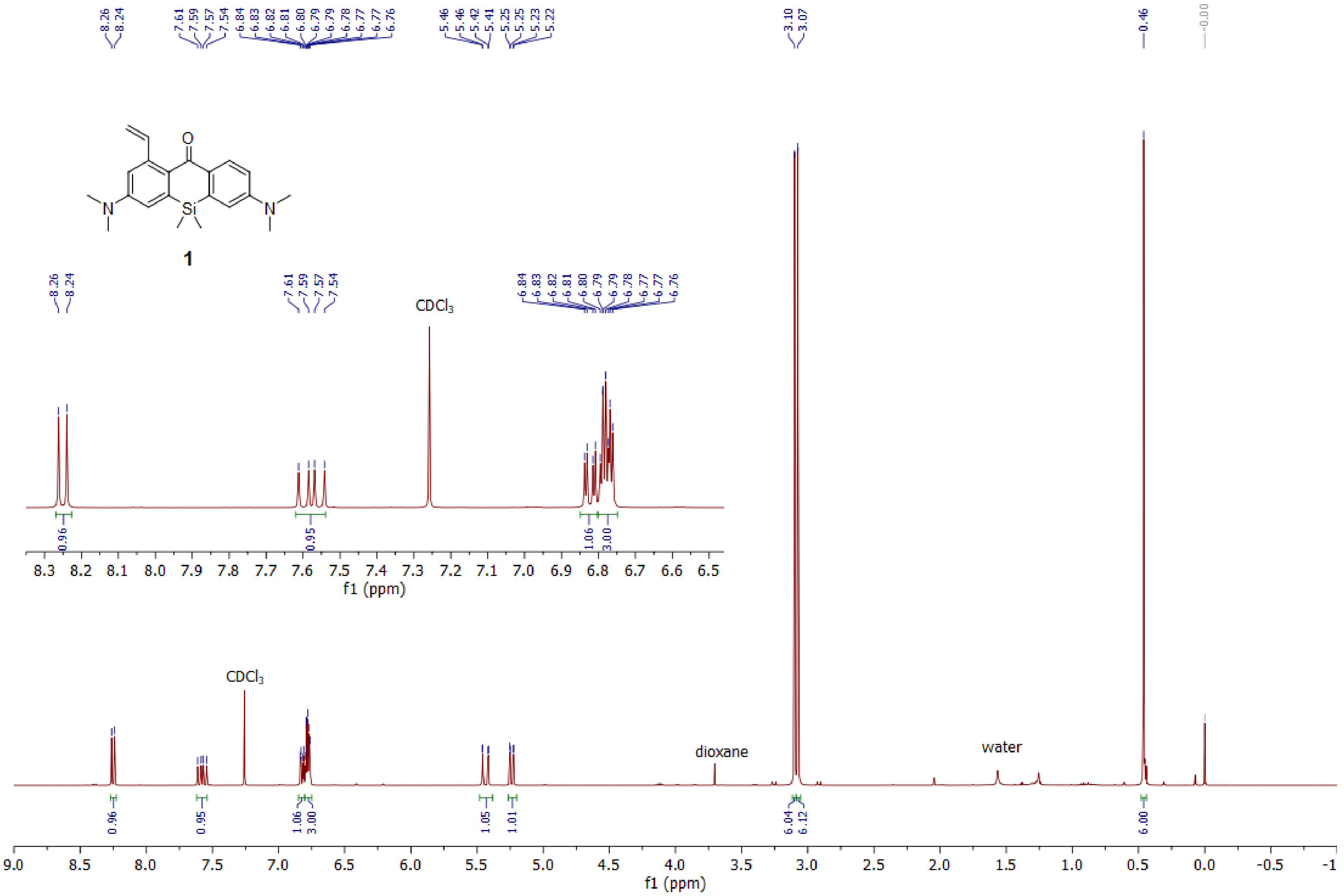

$^{13}C$ (101 MHz, $CDCl_3$)

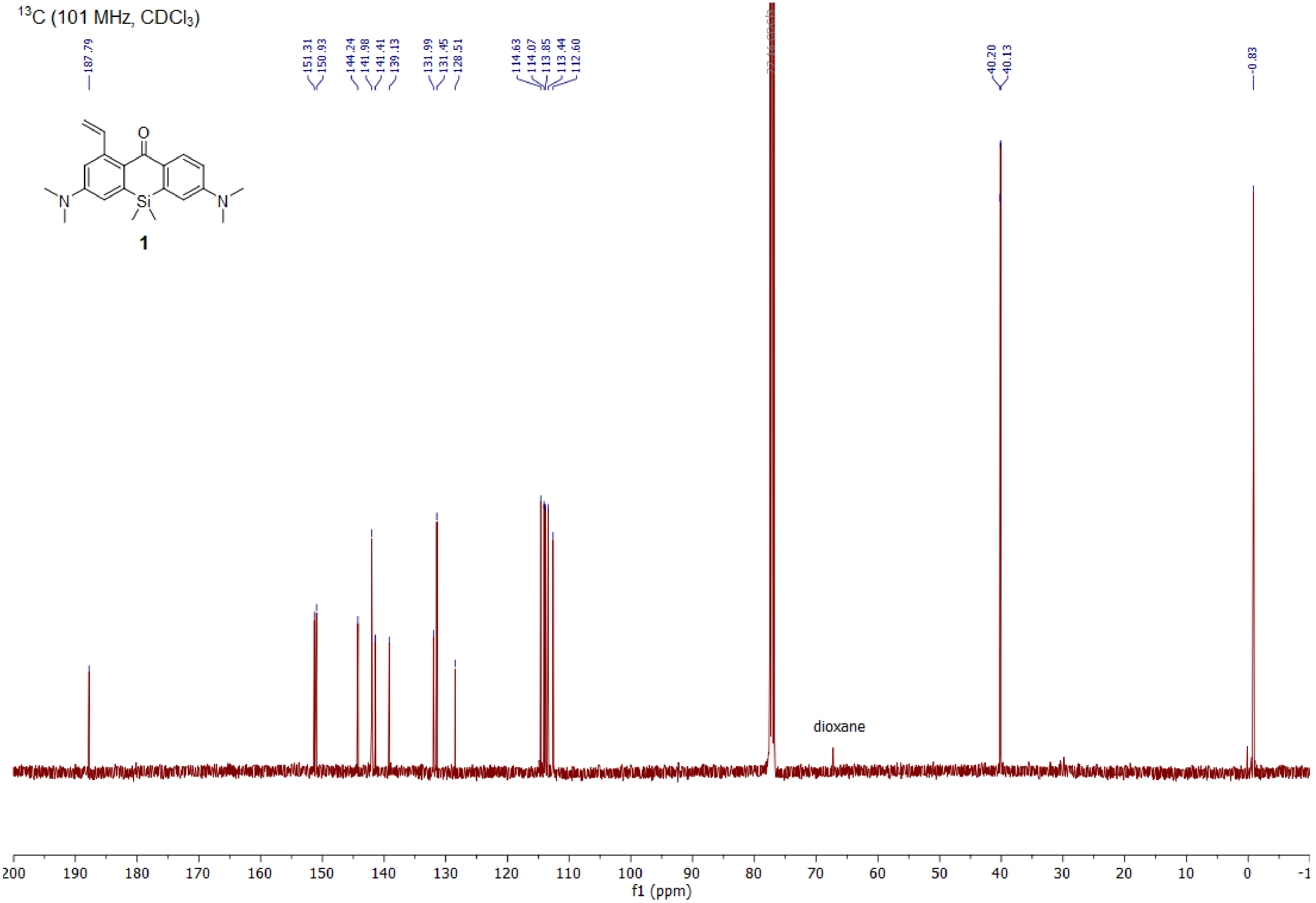

1H (400.15 MHz, CDCl3)

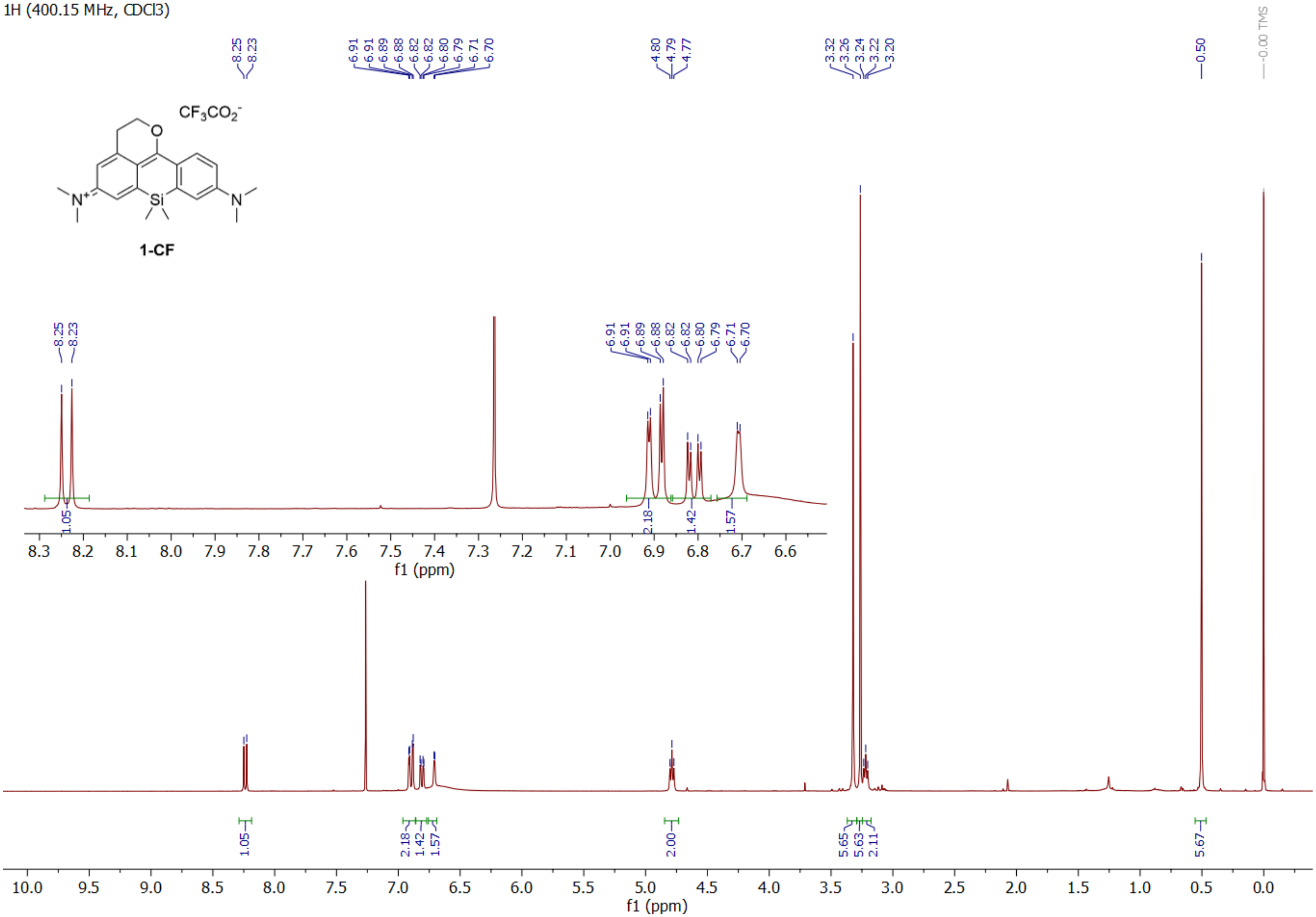

HSQC (400.15 MHz, CDCl3)

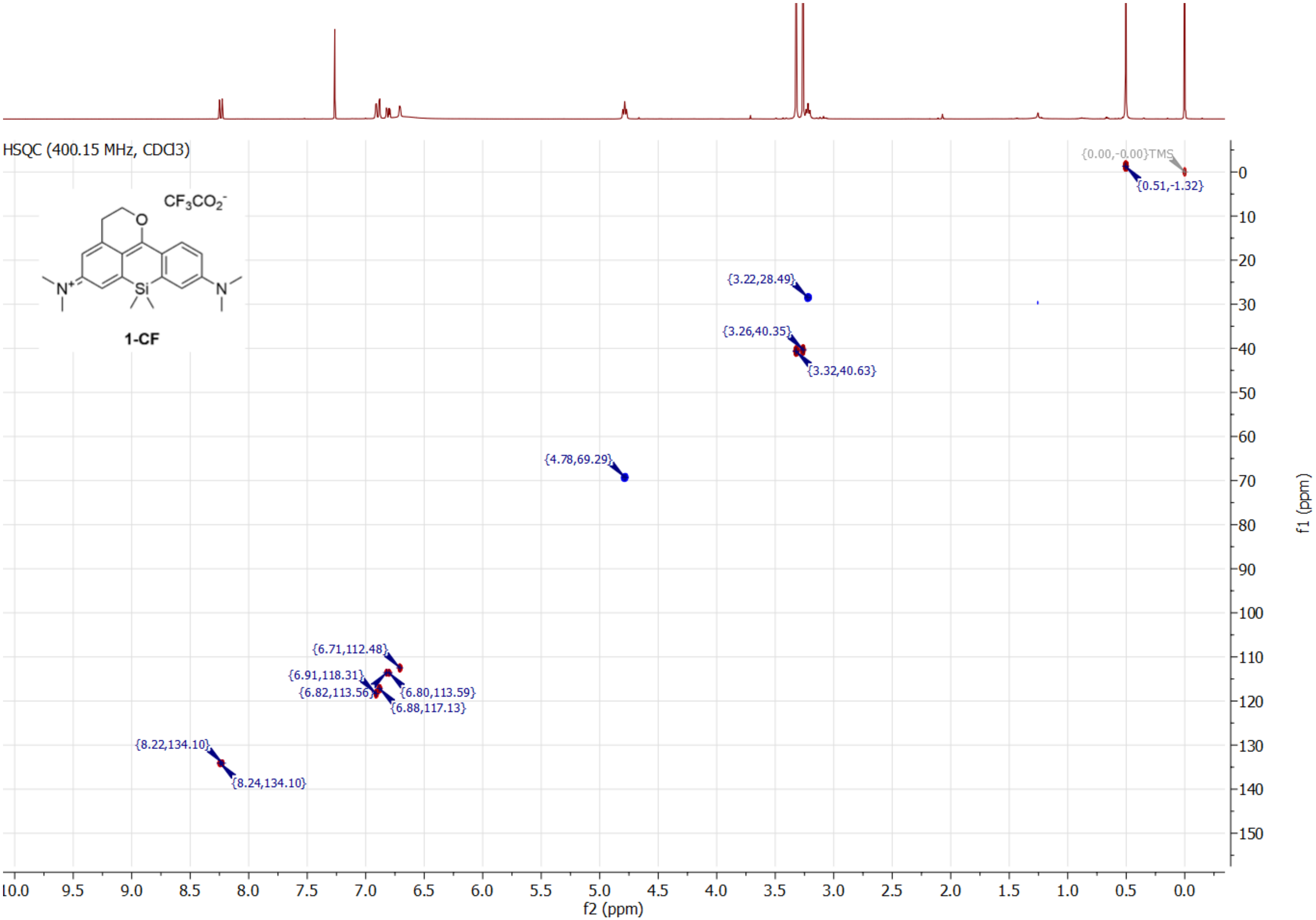

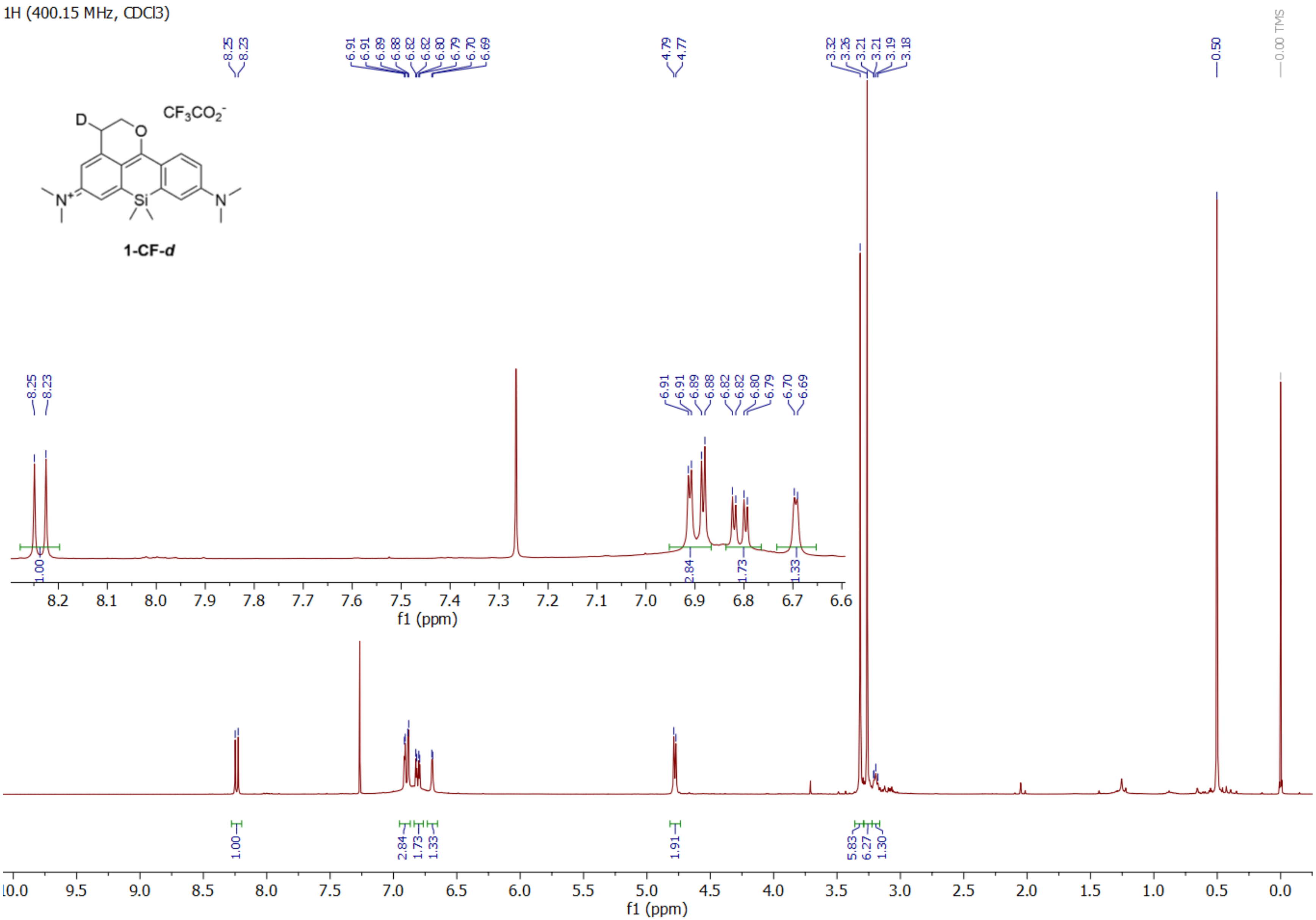

1H (400.15 MHz, CDCl3)
8.25
8.23
6.91
6.91
6.89
6.88
6.82
6.82
6.80
6.79
6.70
6.69
4.79
4.77
3.32
3.26
3.21
3.21
3.19
3.18
0.50
0.00 TMS
D
O
CF3CO2-
N+
Si
N
1-CF-d
1.00
2.84
1.73
1.33
1.91
5.83
6.27
1.30
f1 (ppm)

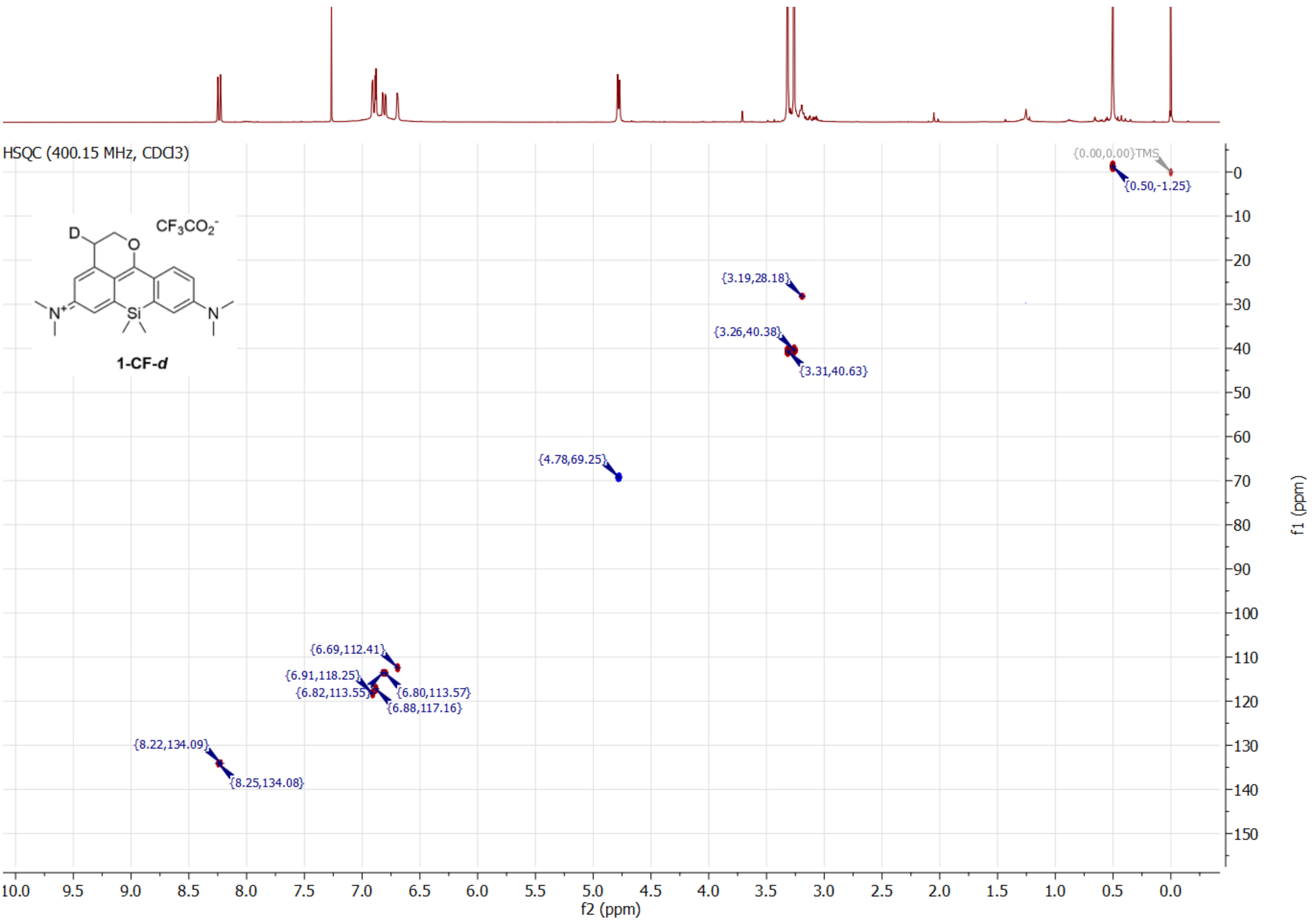
HSQC (400.15 MHz, CDCl3)
CF3CO2-
D
O
N+
Si
N
1-CF-d
{0.00,0.00}TMS
{0.50,-1.25}
{3.19,28.18}
{3.26,40.38}
{3.31,40.63}
{4.78,69.25}
{6.69,112.41}
{6.91,118.25}
{6.82,113.55}
{6.80,113.57}
{6.88,117.16}
{8.22,134.09}
{8.25,134.08}
f2 (ppm)
f1 (ppm)

<sup></sup>

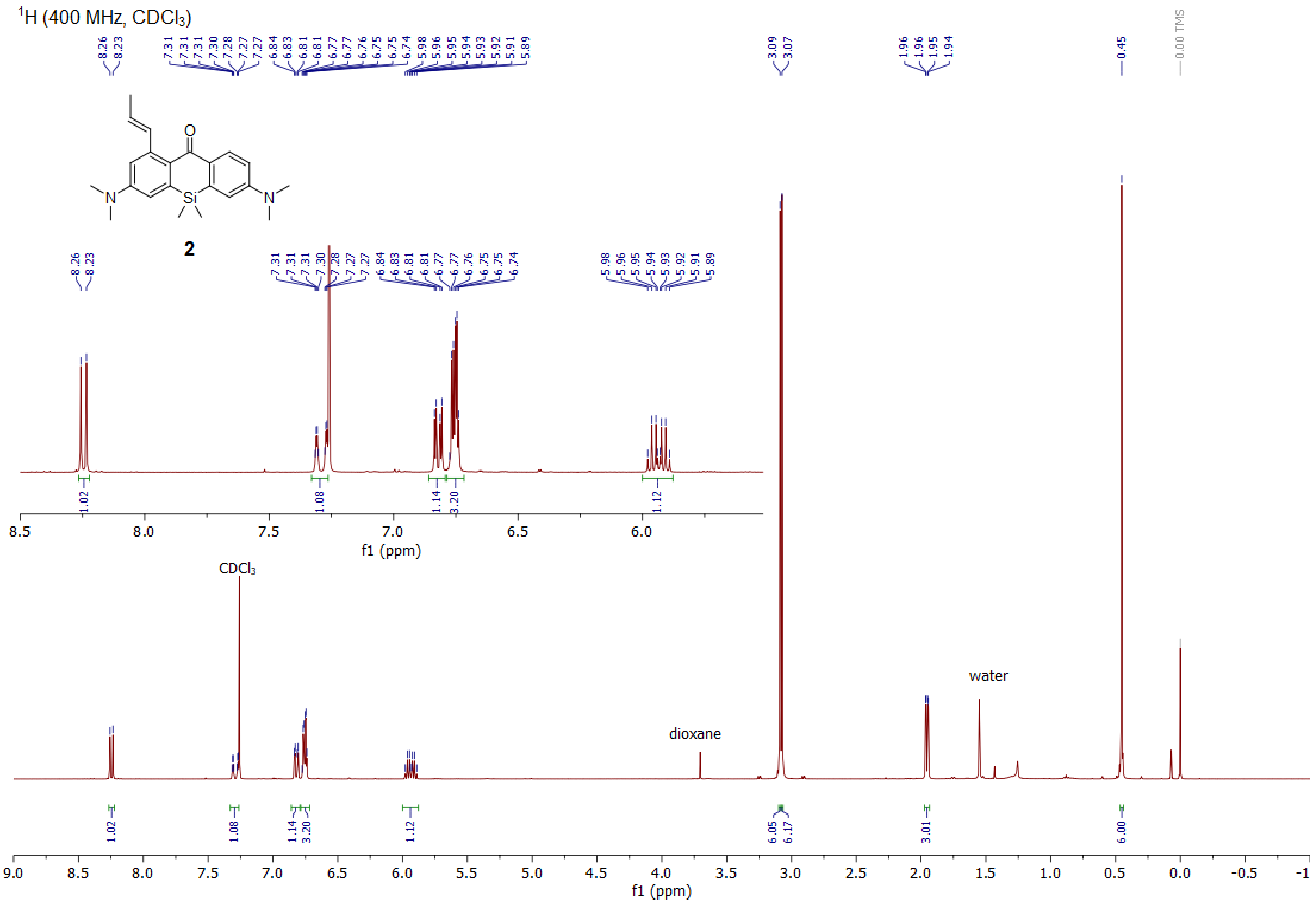

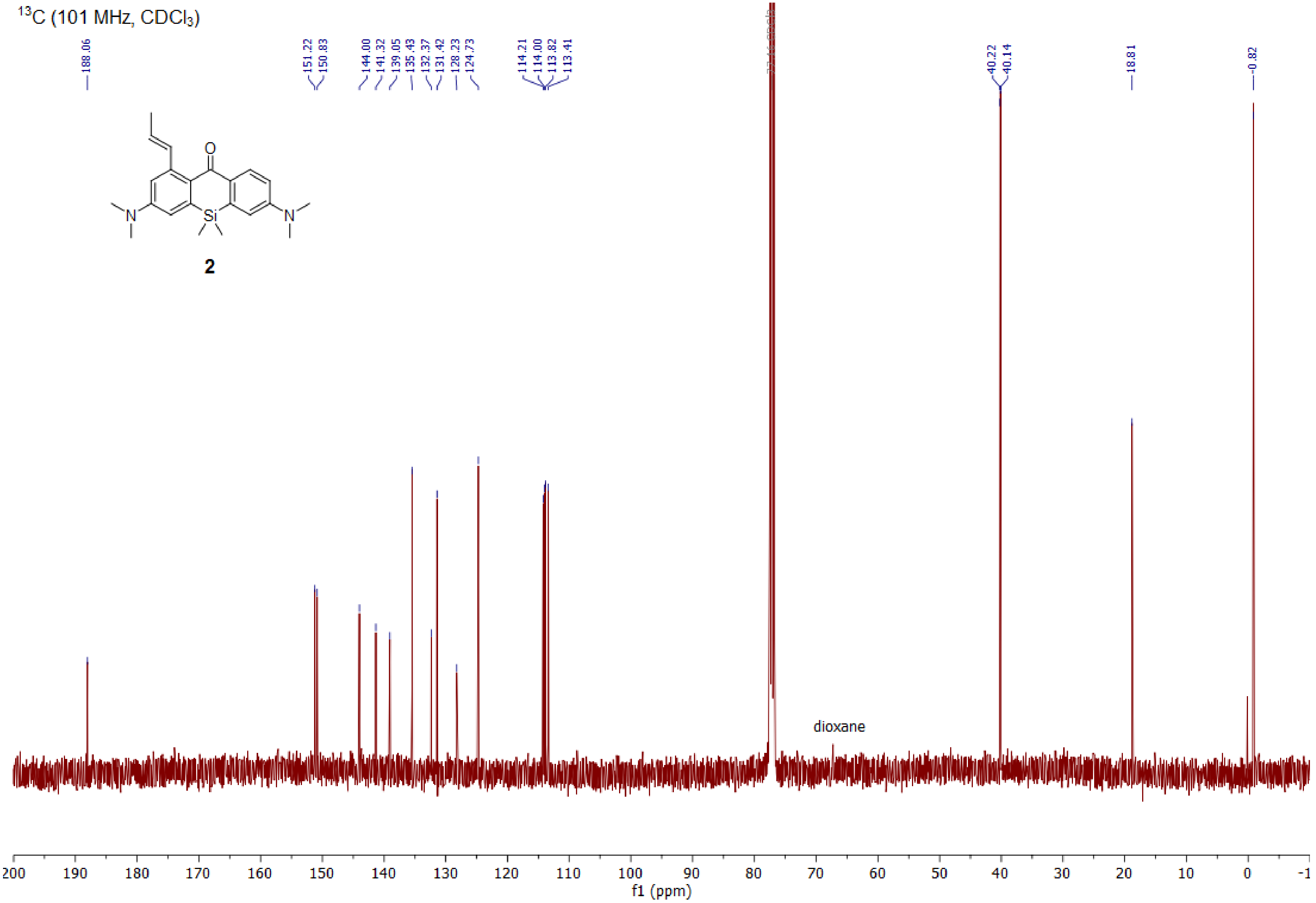

13C (101 MHz, CDCl3)
188.06
151.22
150.83
144.00
141.32
139.05
135.43
132.37
131.42
128.23
124.73
114.21
114.00
113.82
113.41
40.22
40.14
18.81
-0.82
O
N
Si
N
2
dioxane
200 190 180 170 160 150 140 130 120 110 100 90 80 70 60 50 40 30 20 10 0 -1
f1 (ppm)

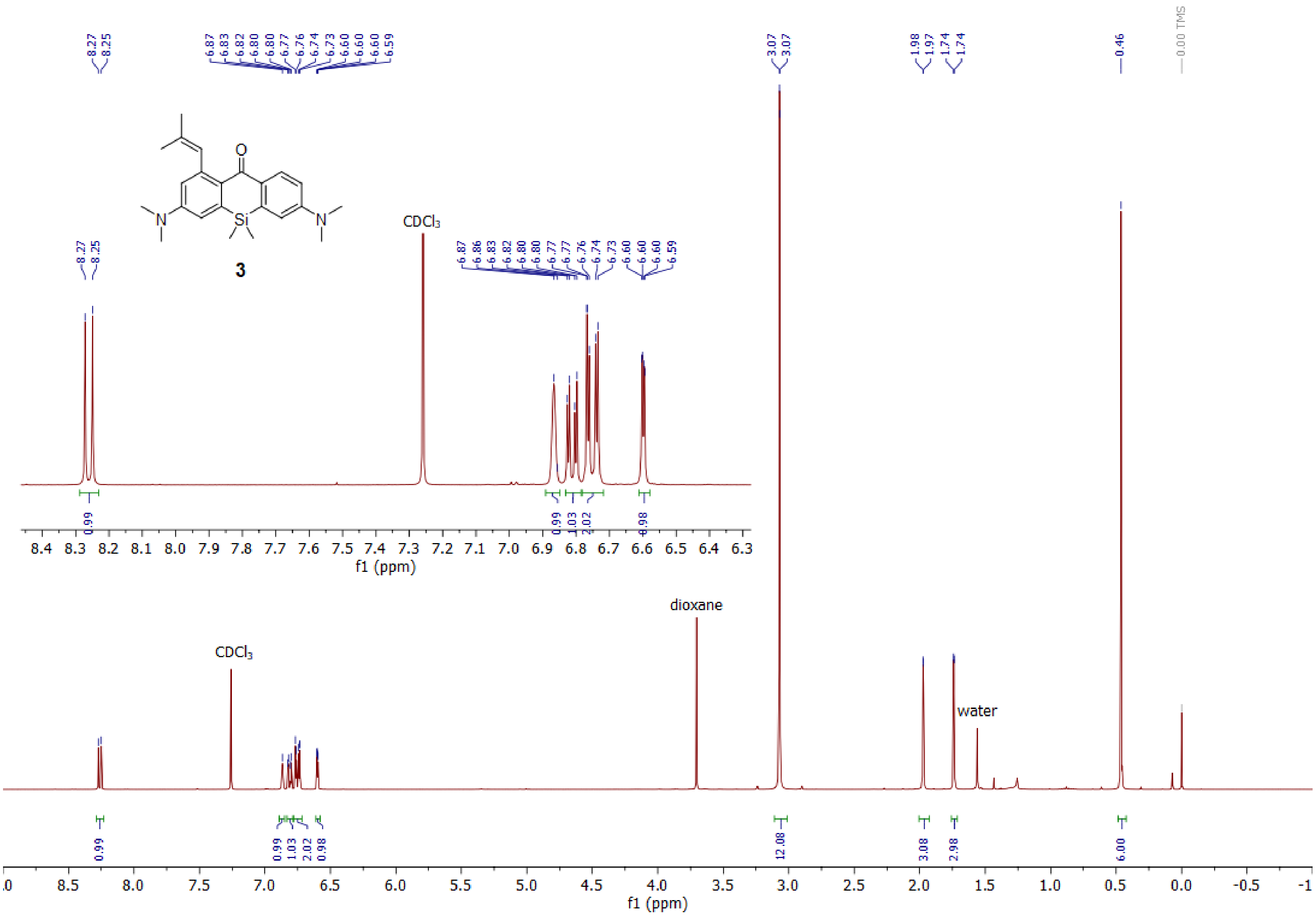
8.27
8.25
6.87
6.83
6.82
6.80
6.80
6.77
6.76
6.74
6.73
6.60
6.60
6.60
6.59
3.07
3.07
1.98
1.97
1.74
1.74
0.46
0.00 TMS
O
N
Si
N
3
CDCl3
6.86
6.77
0.99
1.03
2.02
0.98
f1 (ppm)
dioxane
water
12.08
3.08
2.98
6.00

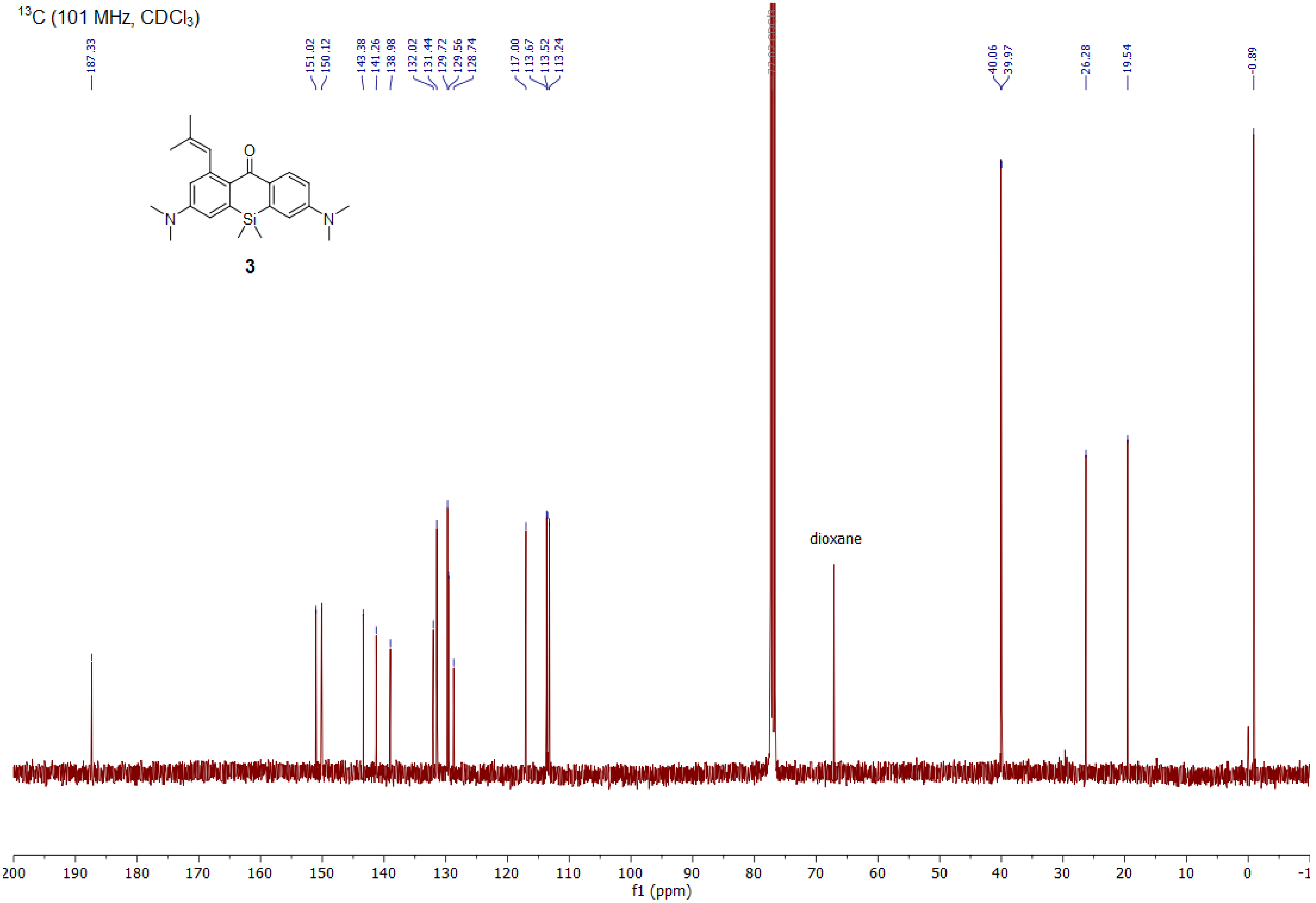
13C (101 MHz, CDCl3)
187.33
151.02
150.12
143.38
141.26
138.98
132.02
131.44
129.72
129.56
128.74
117.00
113.67
113.52
113.24
40.06
39.97
26.28
19.54
-0.89
O
N
Si
N
3
dioxane
200 190 180 170 160 150 140 130 120 110 100 90 80 70 60 50 40 30 20 10 0 -1
f1 (ppm)

[1]H (400 MHz, CDCl3)

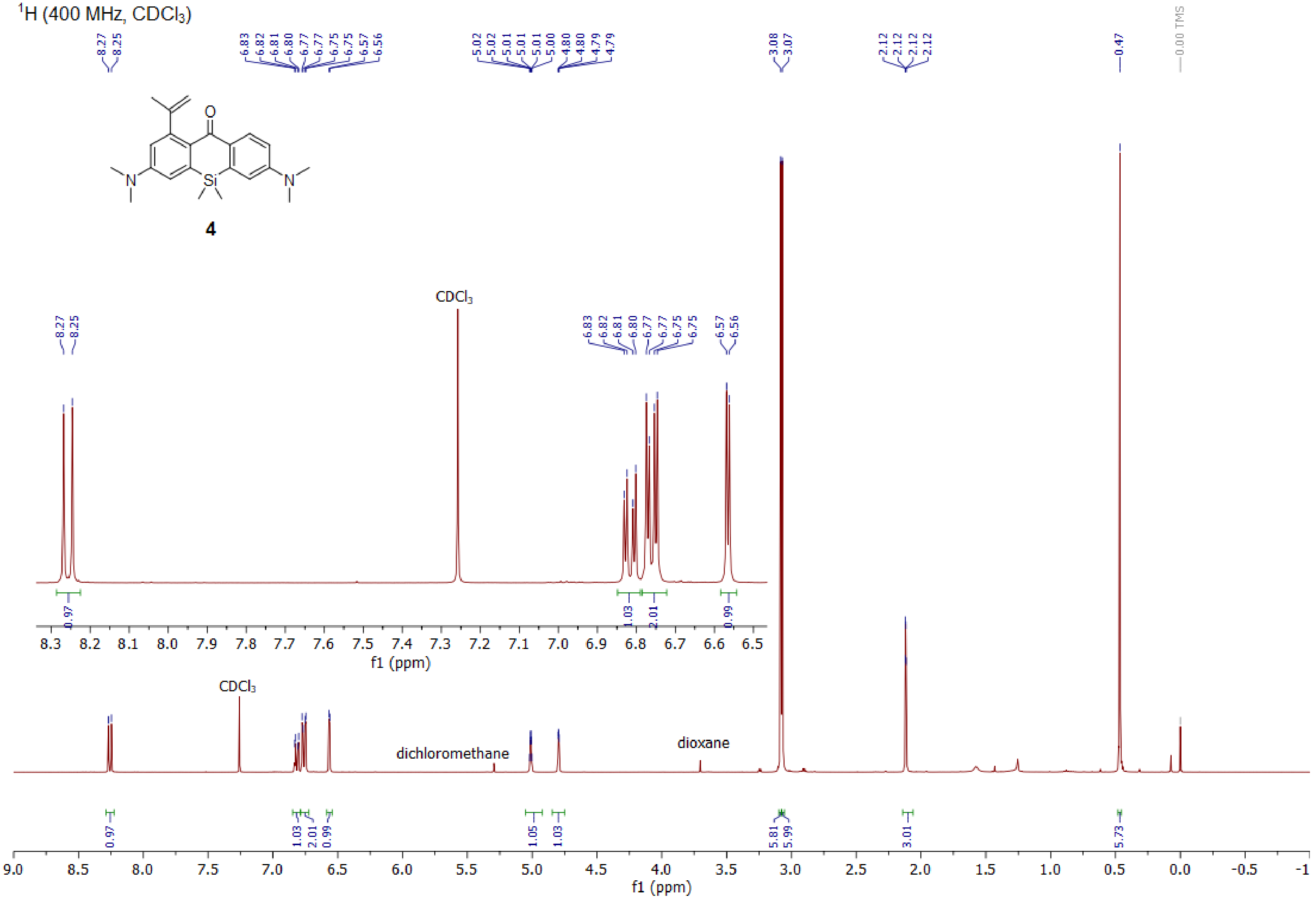

$^{1}H$ (400 MHz, $CDCl_3$)

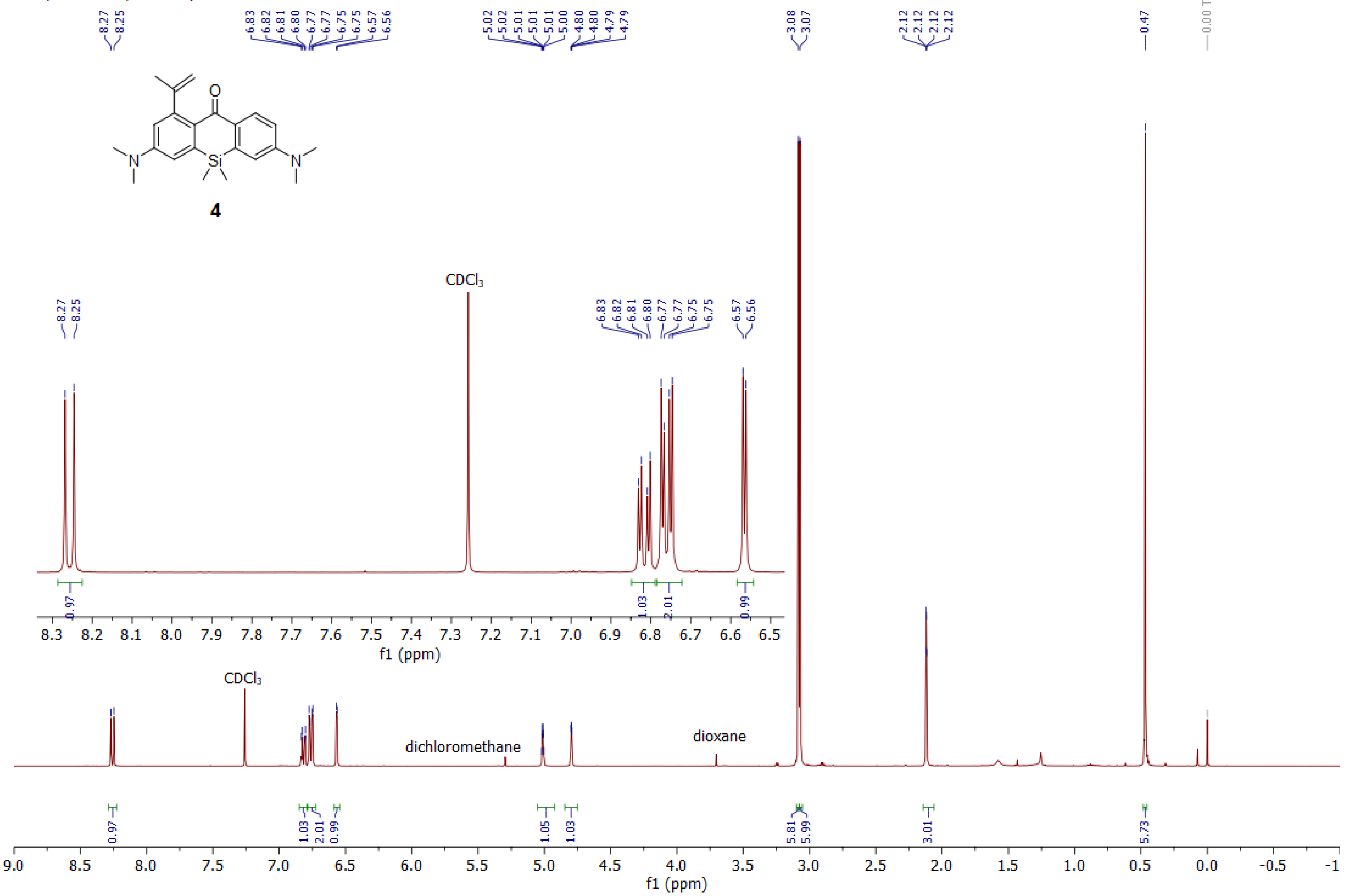

[1]H (400 MHz, $CDCl_3$)

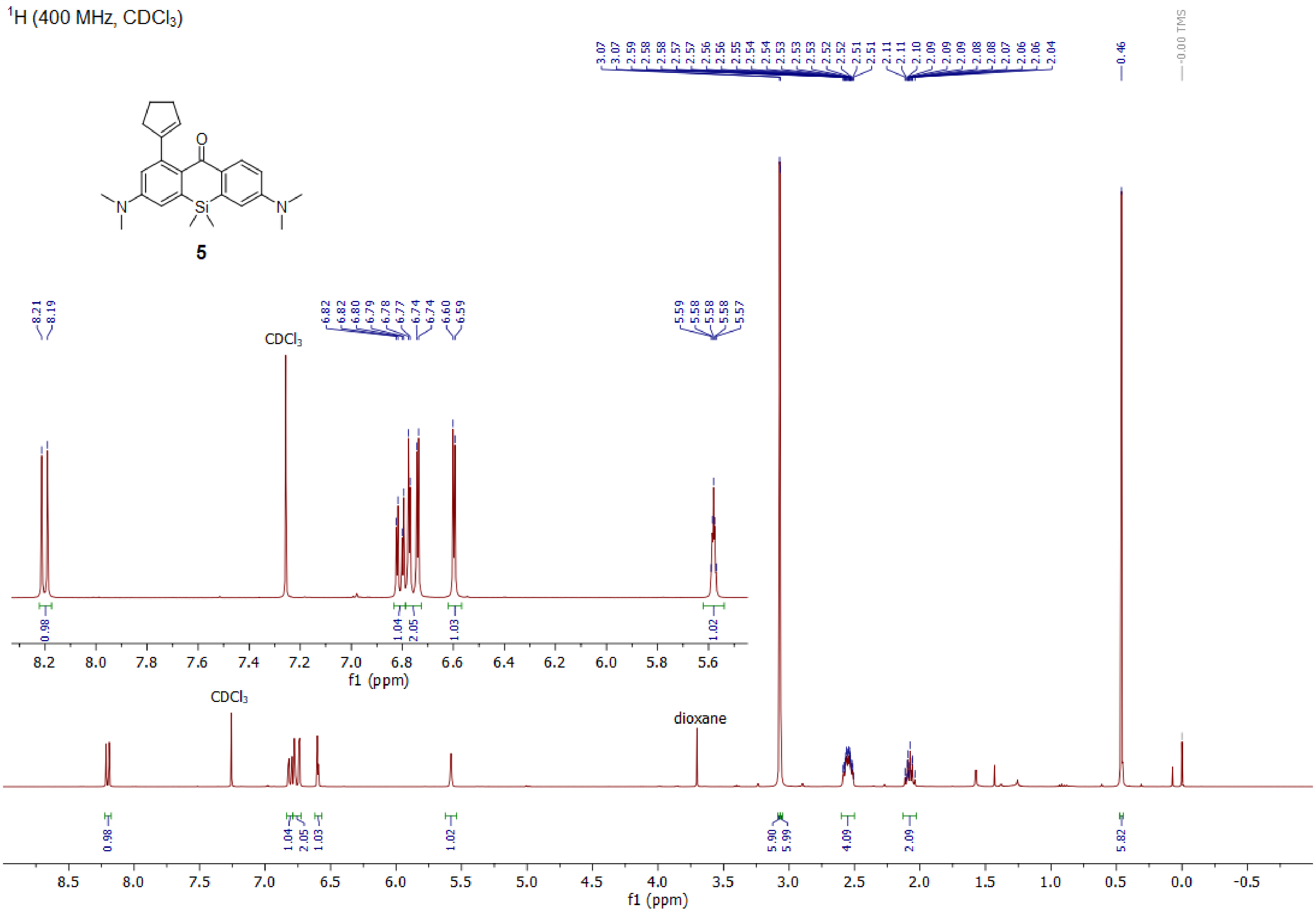

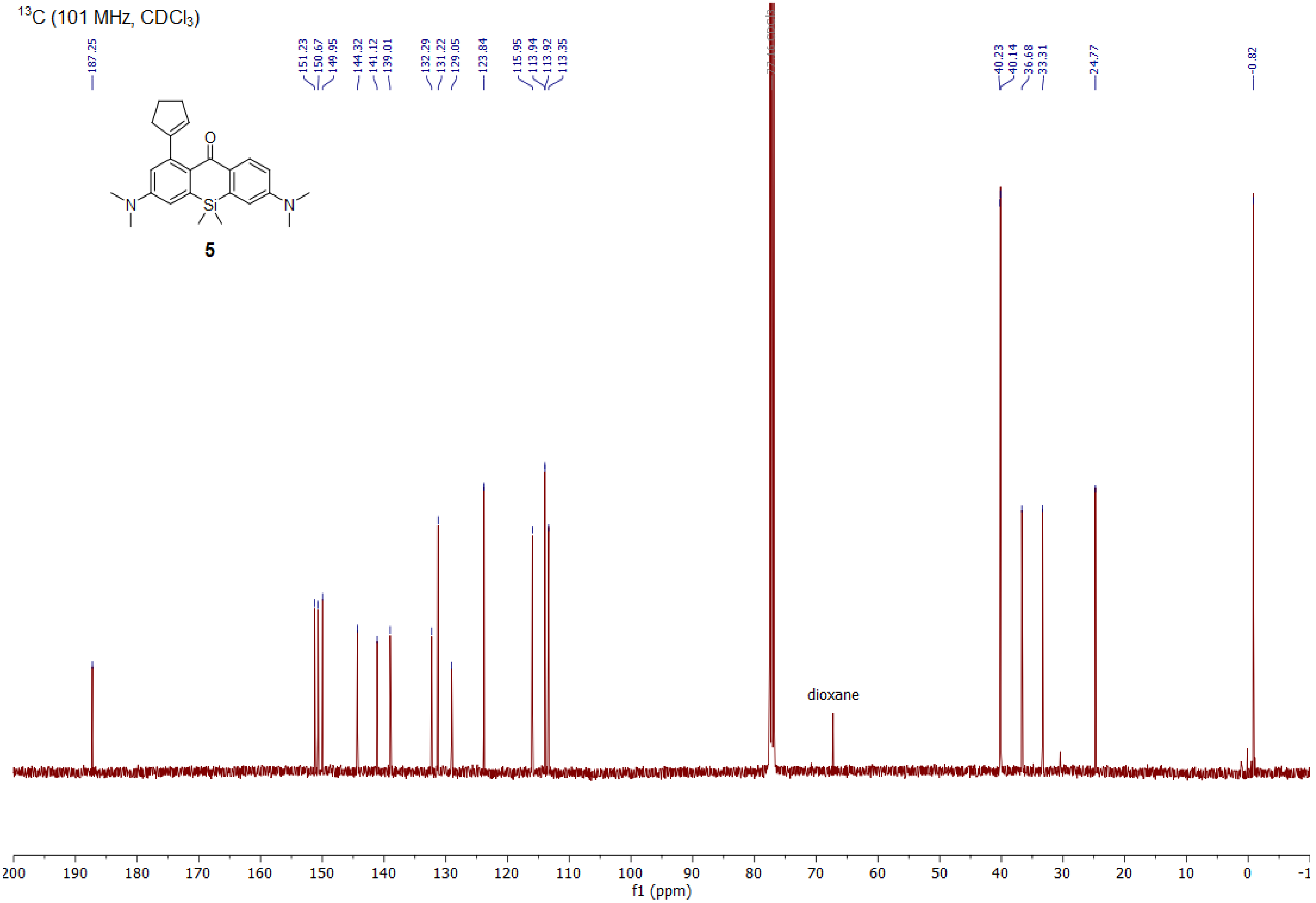

$^{13}$C (101 MHz, CDCl$_3$)
187.25
151.23
150.67
149.95
144.32
141.12
139.01
132.29
131.22
129.05
123.84
115.95
113.94
113.92
113.35
40.23
40.14
36.68
33.31
24.77
-0.82
O
N
Si
N
5
dioxane
200 190 180 170 160 150 140 130 120 110 100 90 80 70 60 50 40 30 20 10 0 -1
f1 (ppm)

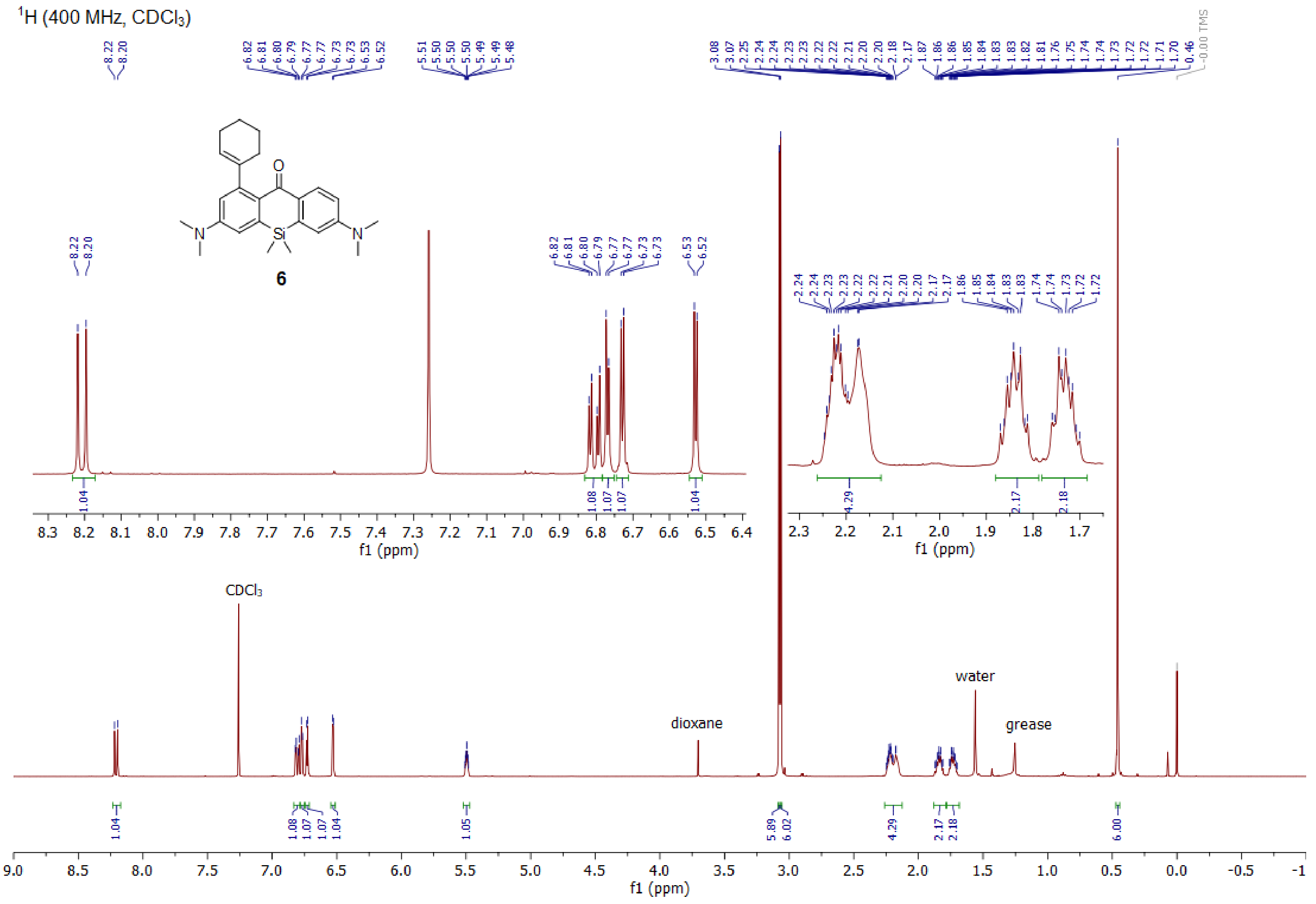

$^1$H (400 MHz, CDCl$_3$)
6
CDCl₃
dioxane
water
grease
f1 (ppm)

$^{13}C$ (101 MHz, $CDCl_3$)

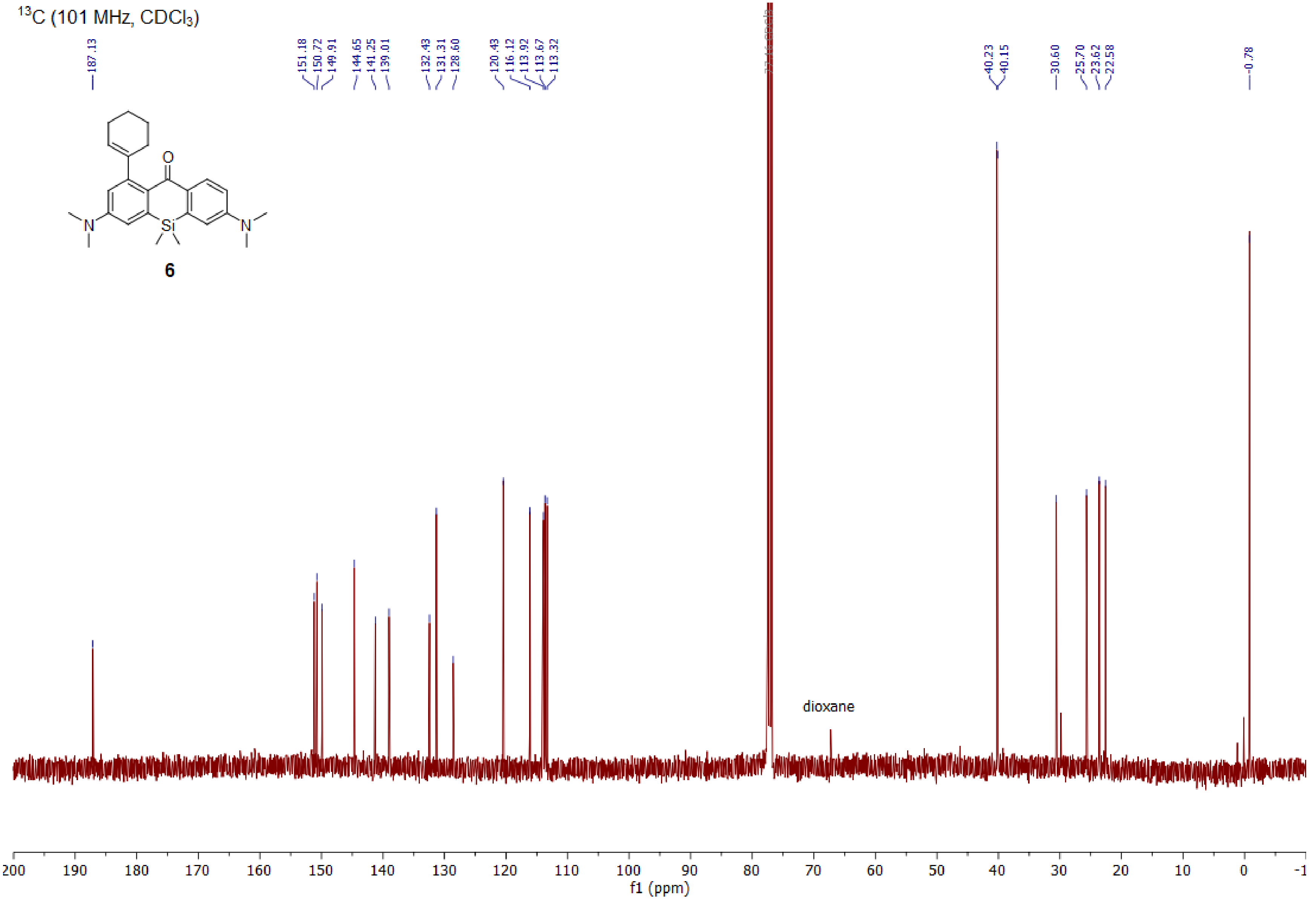

1H (400.15 MHz, CDCl3)

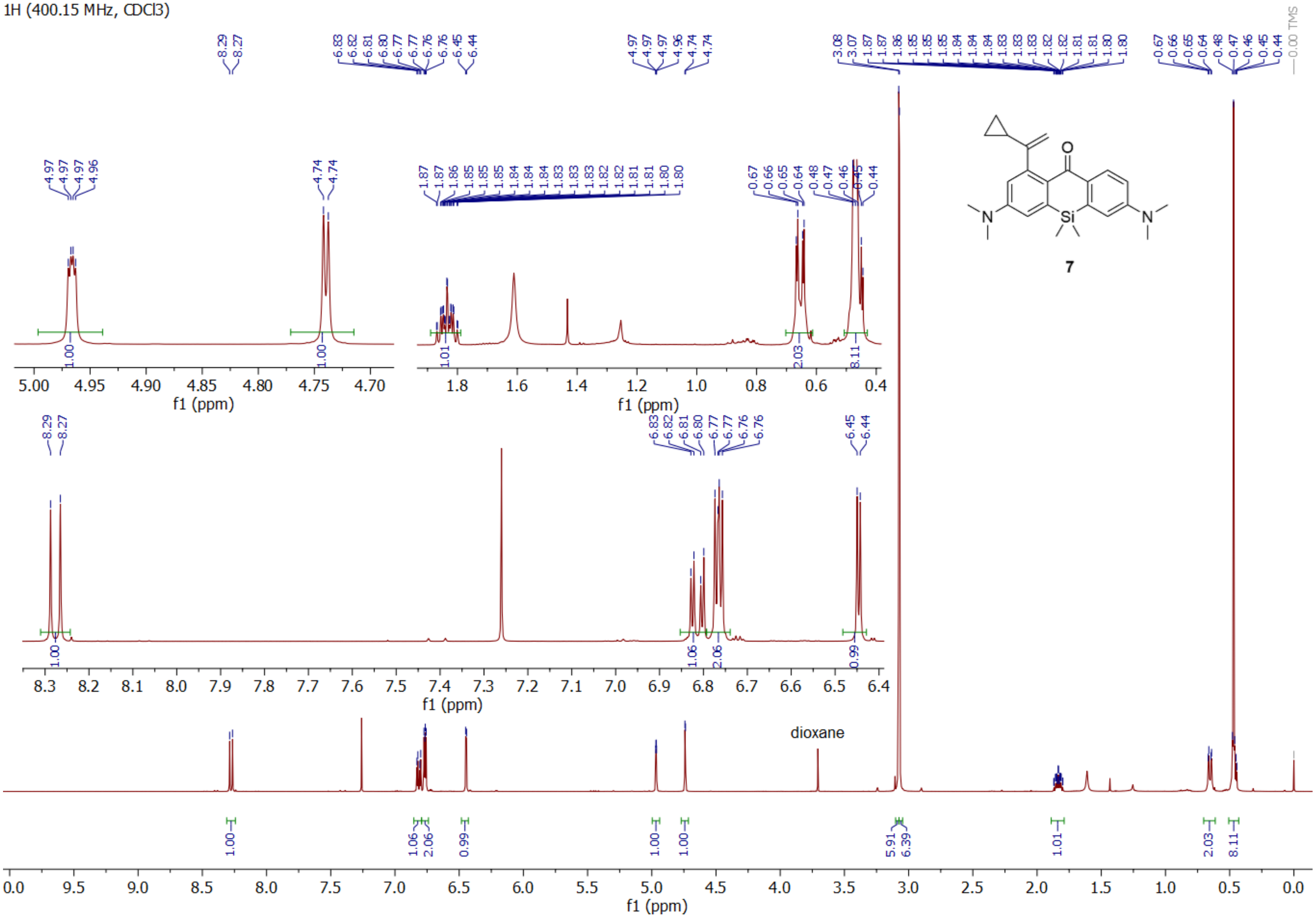

13C (100.63 MHz, CDCl3)

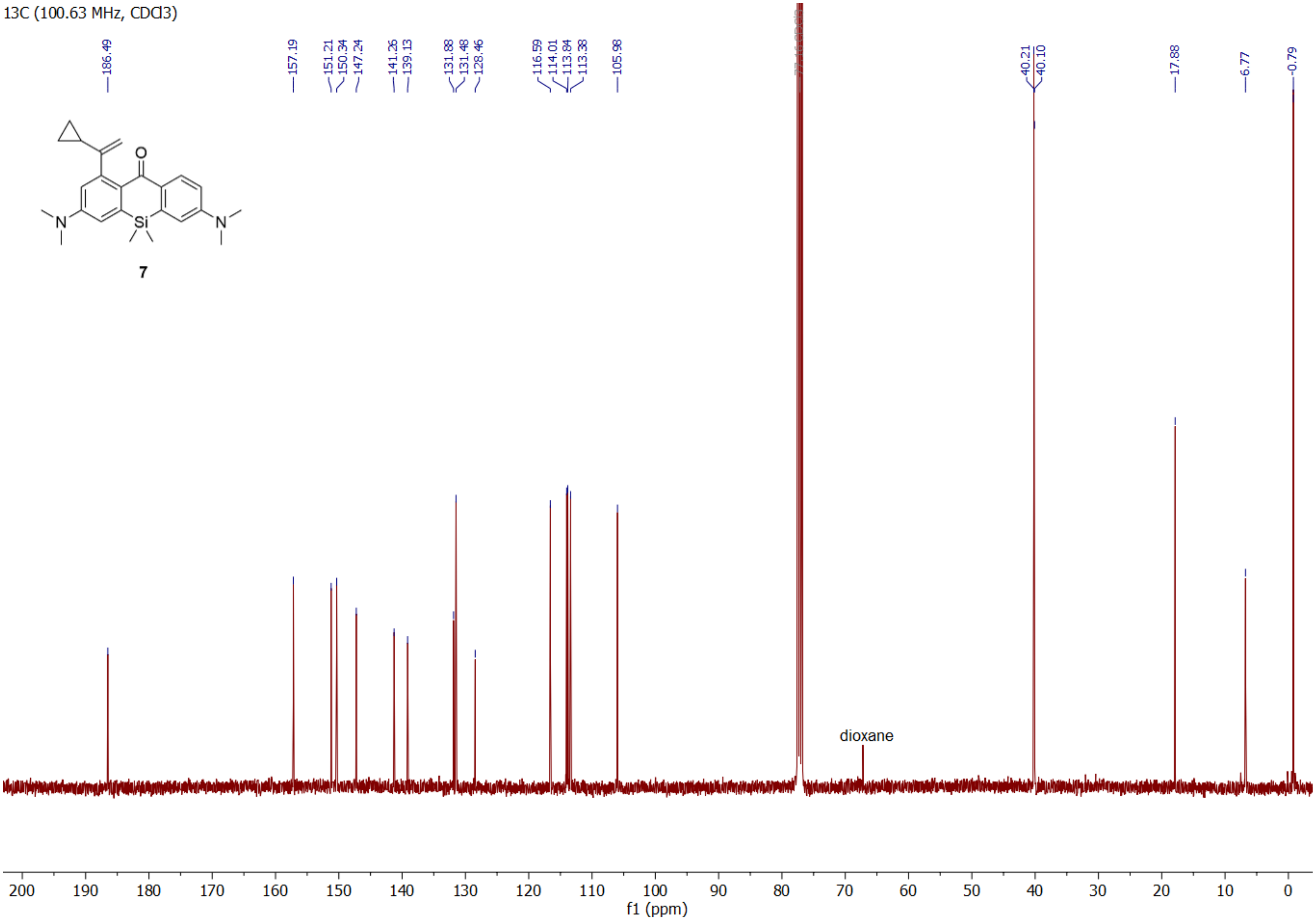

$^{1}H$ (400 MHz, $CDCl_3$)

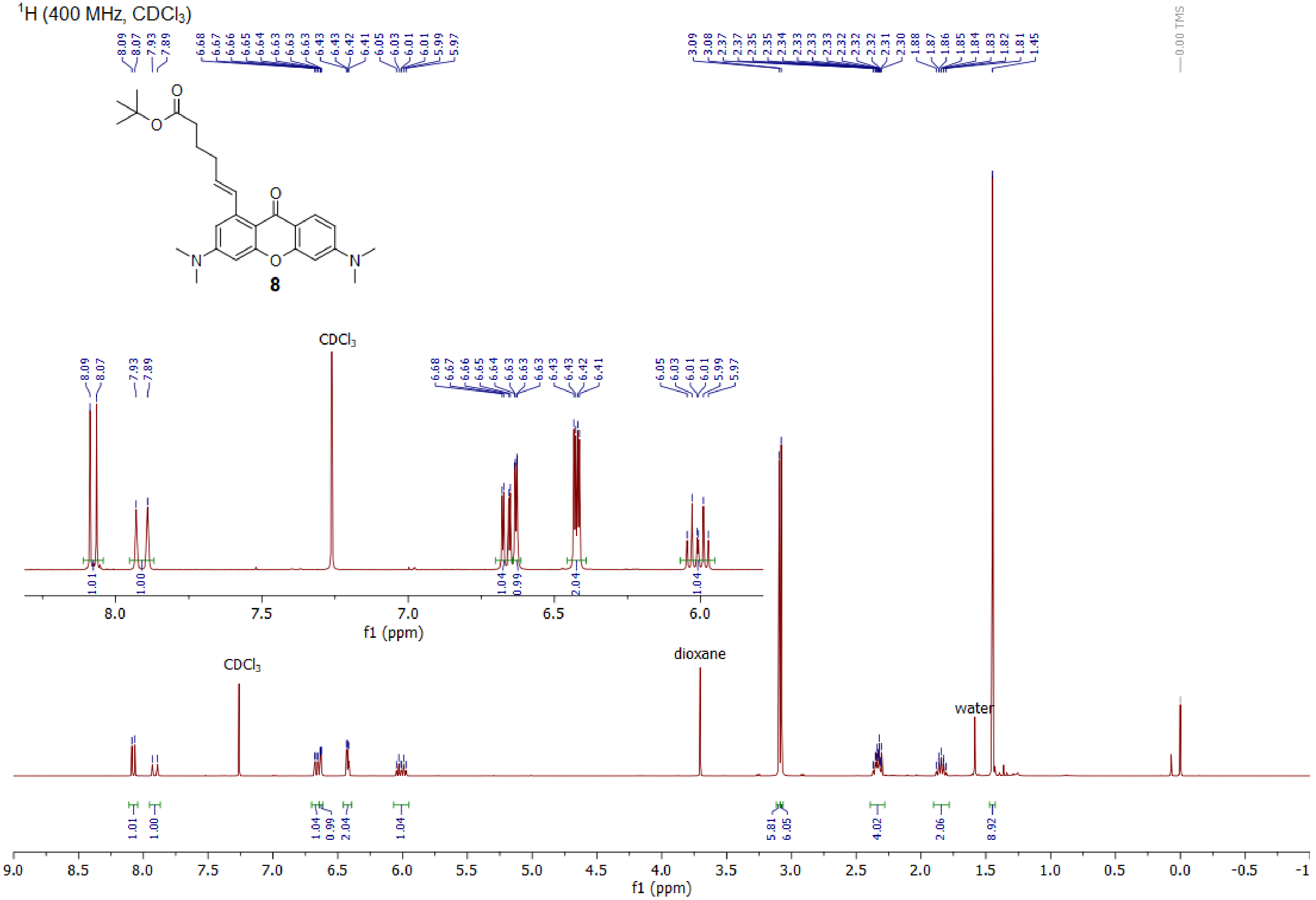

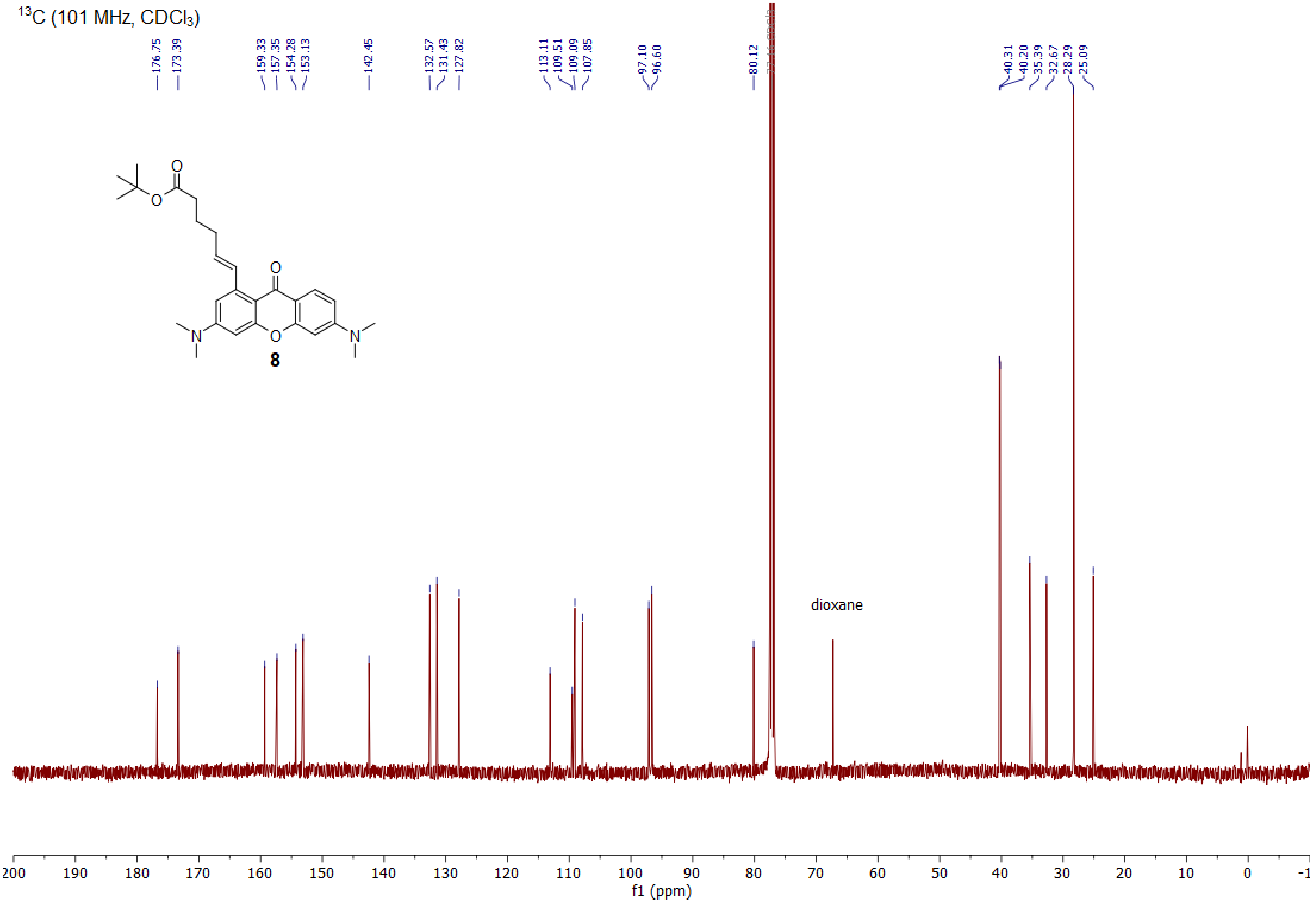

$^{13}C$ (101 MHz, $CDCl_3$)
176.75
173.39
159.33
157.35
154.28
153.13
142.45
132.57
131.43
127.82
113.11
109.51
109.09
107.85
97.10
96.60
80.12
40.31
40.20
35.39
32.67
28.29
25.09
O
O
O
O
N
N
8
dioxane
200
190
180
170
160
150
140
130
120
110
100
90
80
70
60
50
40
30
20
10
0
-1
f1 (ppm)

<sup></sup>

$^{1}$H (400 MHz, DMSO-$d_6$)

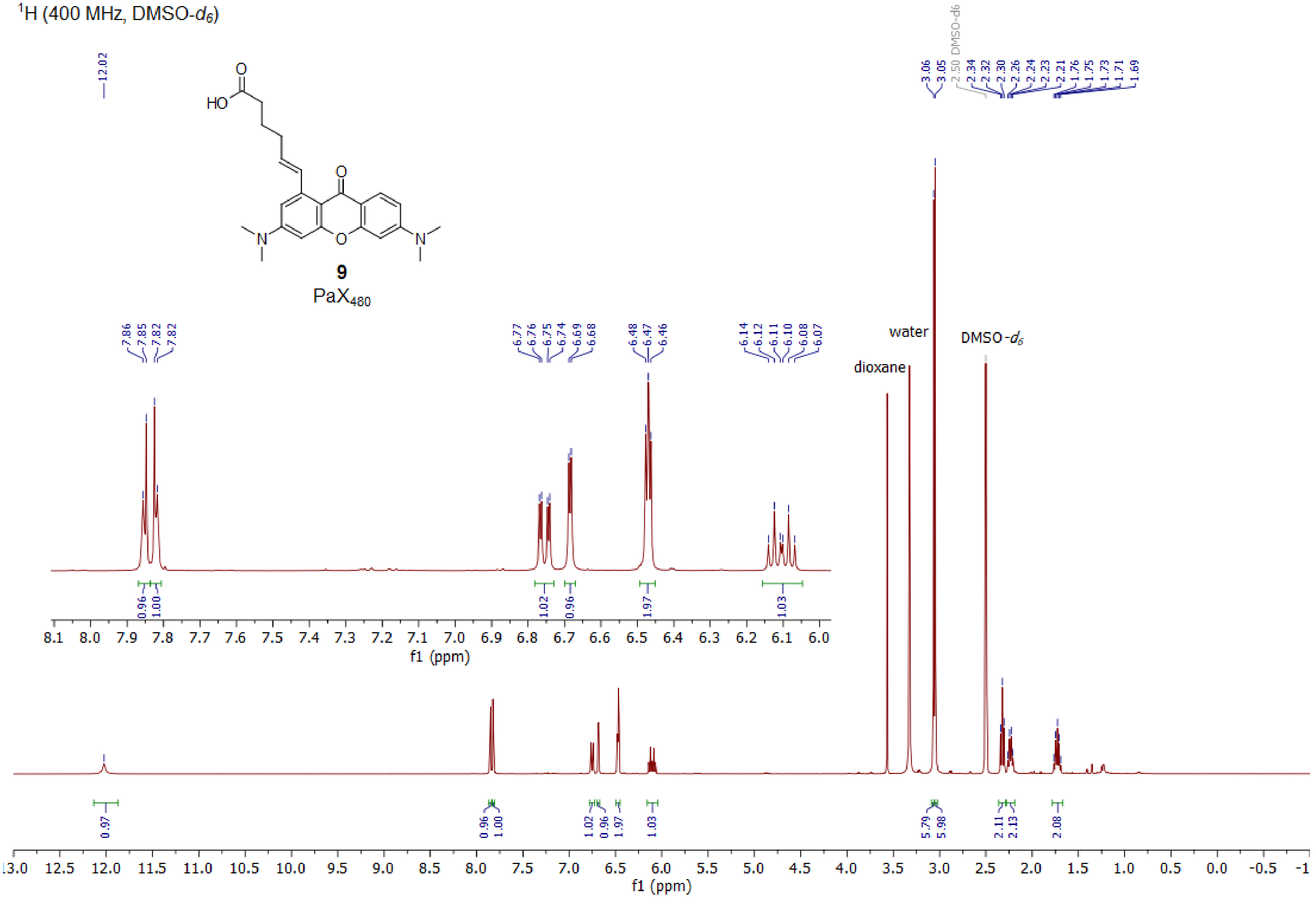

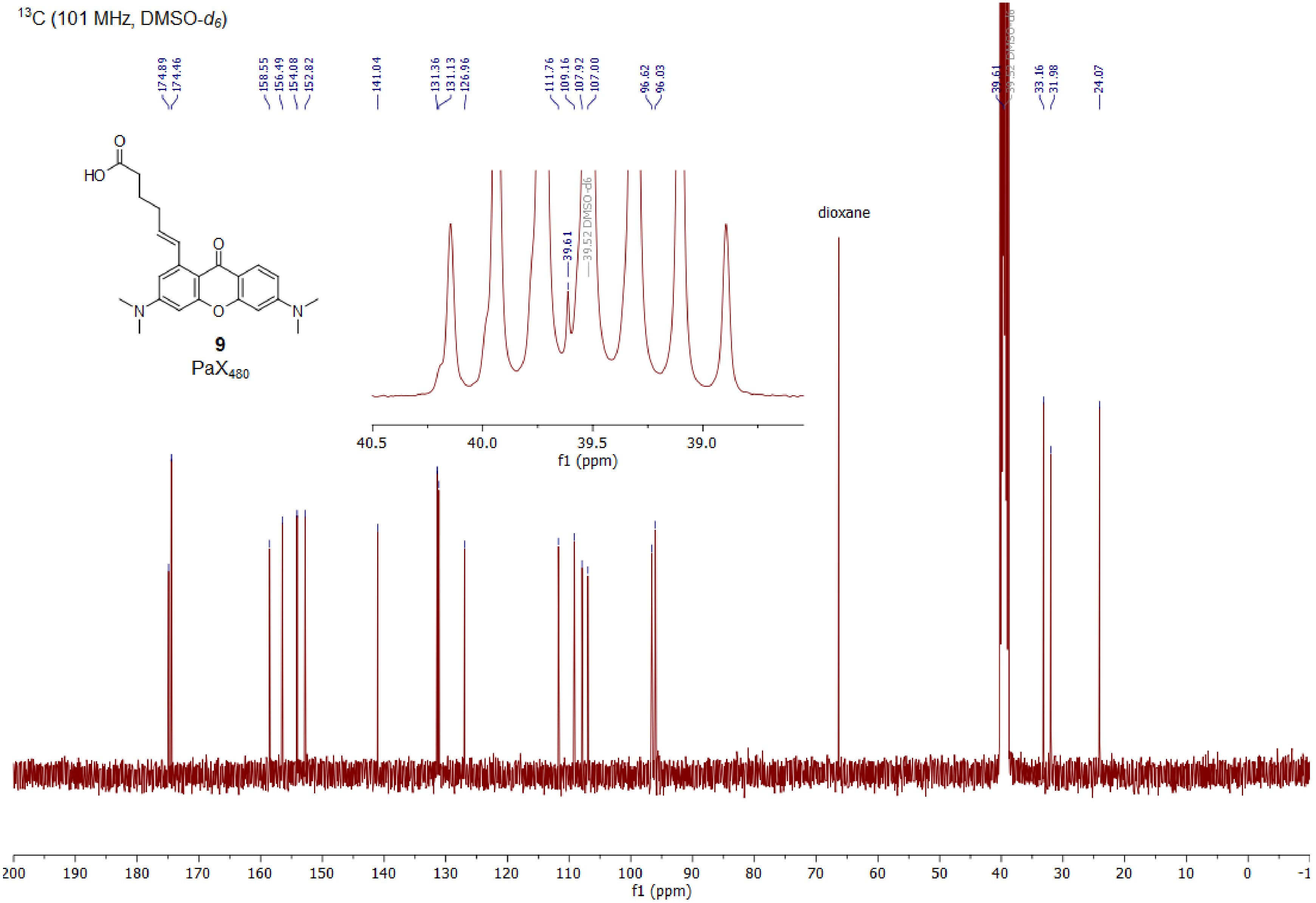

13C (101 MHz, DMSO-d6)
174.89
174.46
158.55
156.49
154.08
152.82
141.04
131.36
131.13
126.96
111.76
109.16
107.92
107.00
96.62
96.03
39.61
39.52 DMSO-d6
33.16
31.98
24.07
HO
O
N
9
PaX480
dioxane
40.5
40.0
39.5
39.0
f1 (ppm)
200
190
180
170
160
150
140
130
120
110
100
90
80
70
60
50
40
30
20
10
0
-1
f1 (ppm)

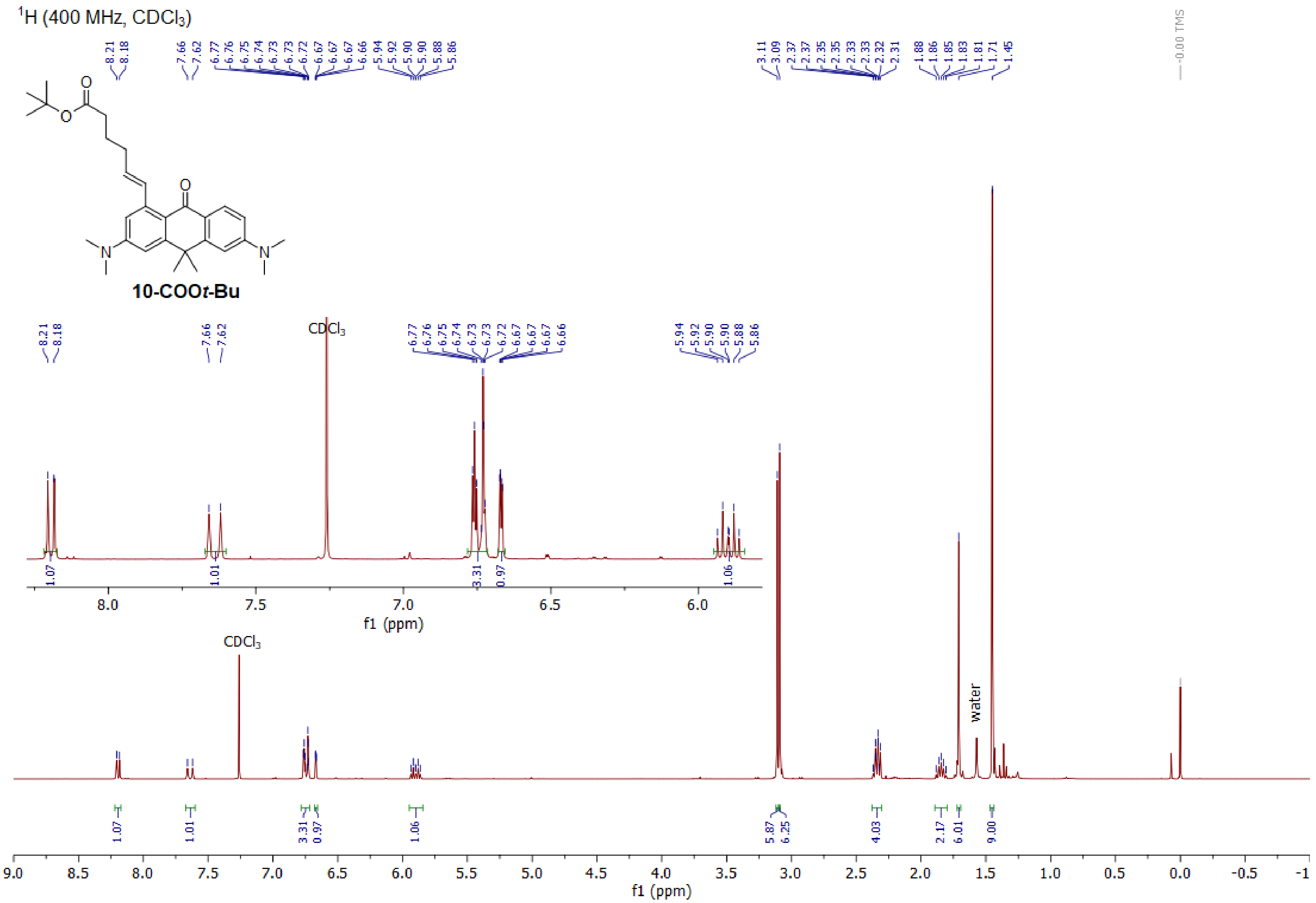

1H (400 MHz, CDCl3)
10-COOt-Bu
8.21
8.18
7.66
7.62
6.77
6.76
6.75
6.74
6.73
6.73
6.72
6.67
6.67
6.67
6.66
5.94
5.92
5.90
5.90
5.88
5.86
3.11
3.09
2.37
2.37
2.35
2.35
2.33
2.33
2.32
2.31
1.88
1.86
1.85
1.83
1.81
1.71
1.45
-0.00 TMS
CDCl3
water
1.07
1.01
3.31
0.97
1.06
5.87
6.25
4.03
2.17
6.01
9.00
f1 (ppm)

$^{13}C$ (101 MHz, $CDCl_3$)

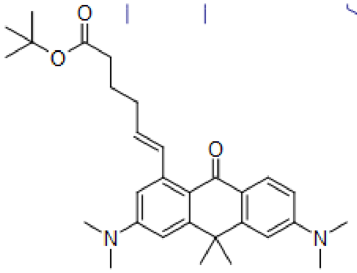

**10-COO*t*-Bu**

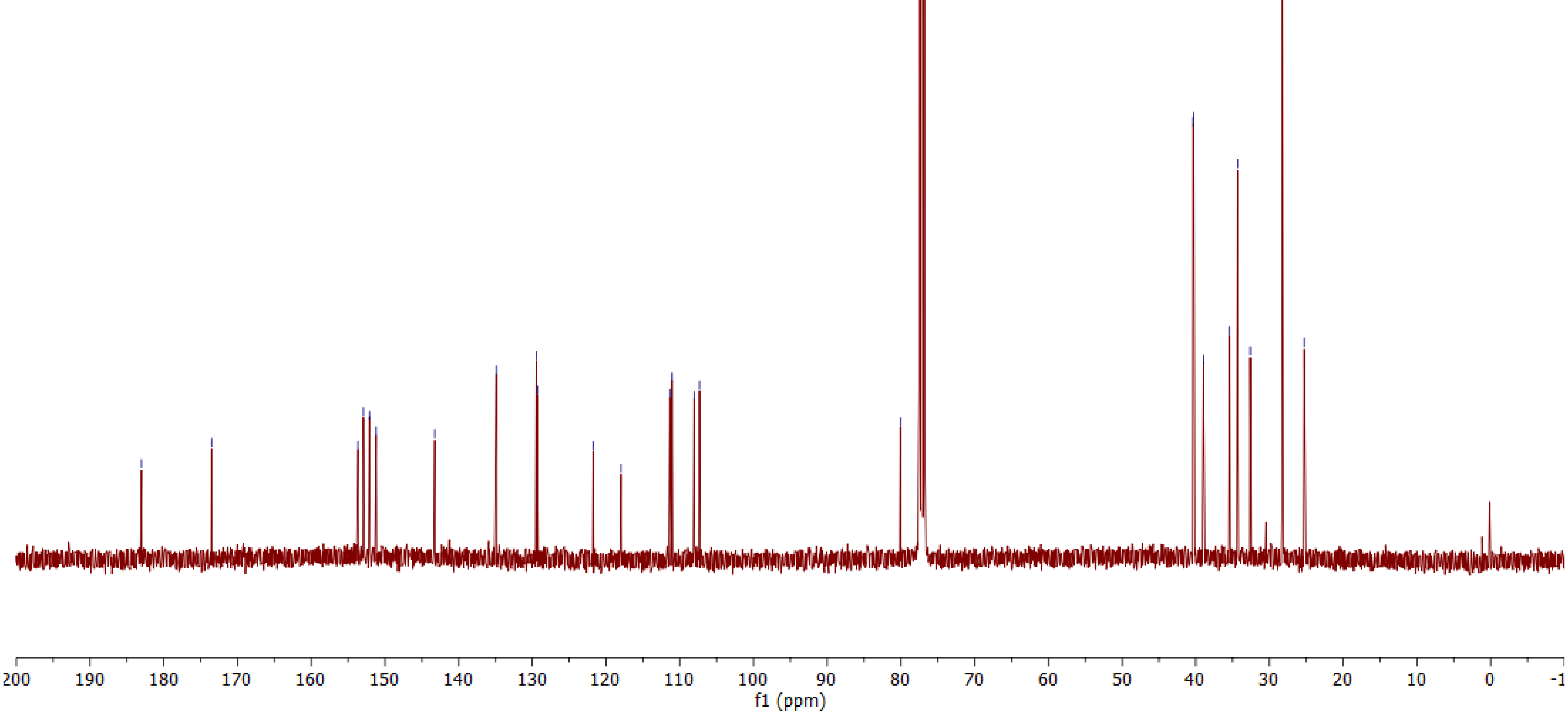

$^1$H (400 MHz, DMSO-$d_6$)

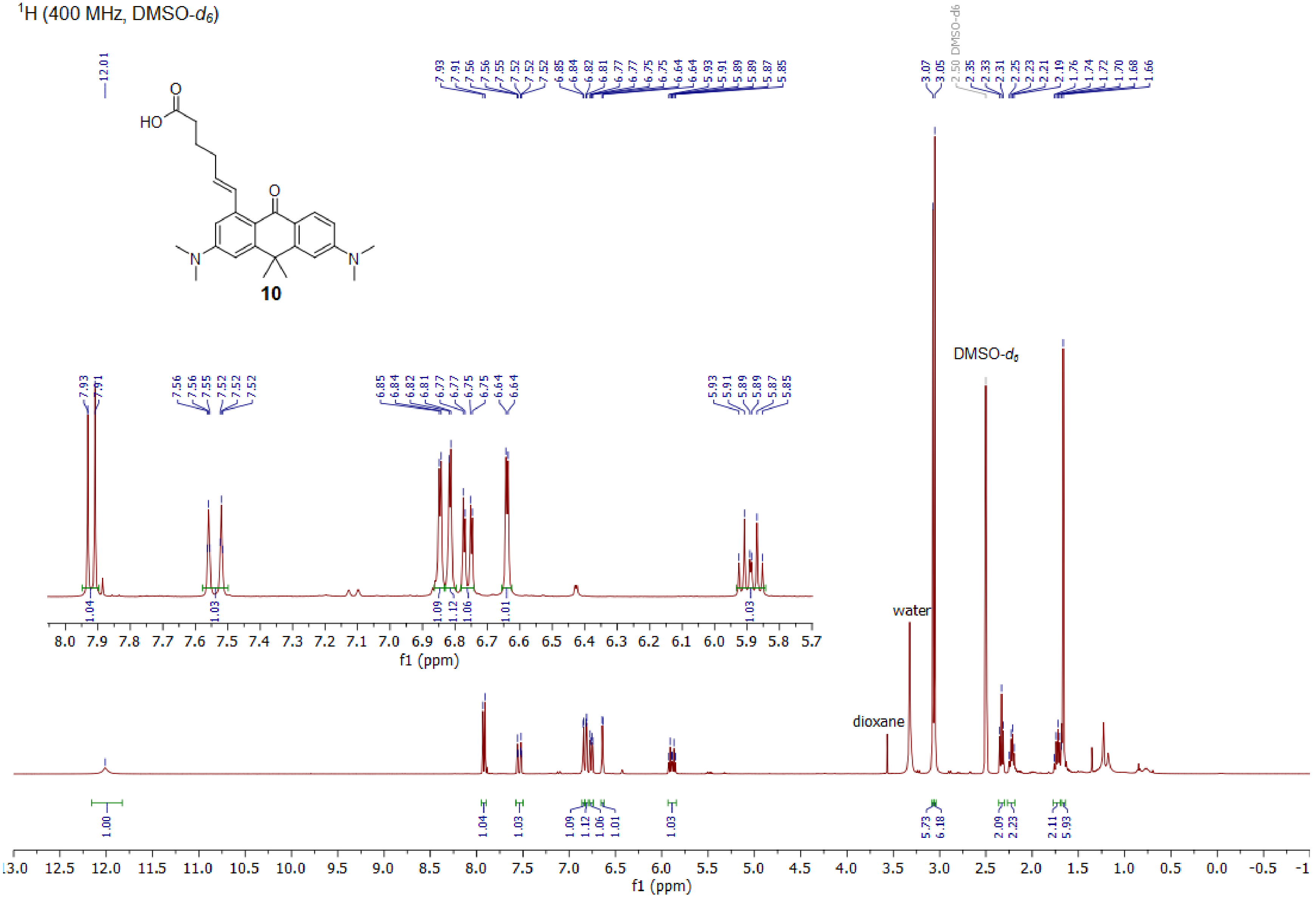

$^{13}$C (101 MHz, DMSO-$d_6$)

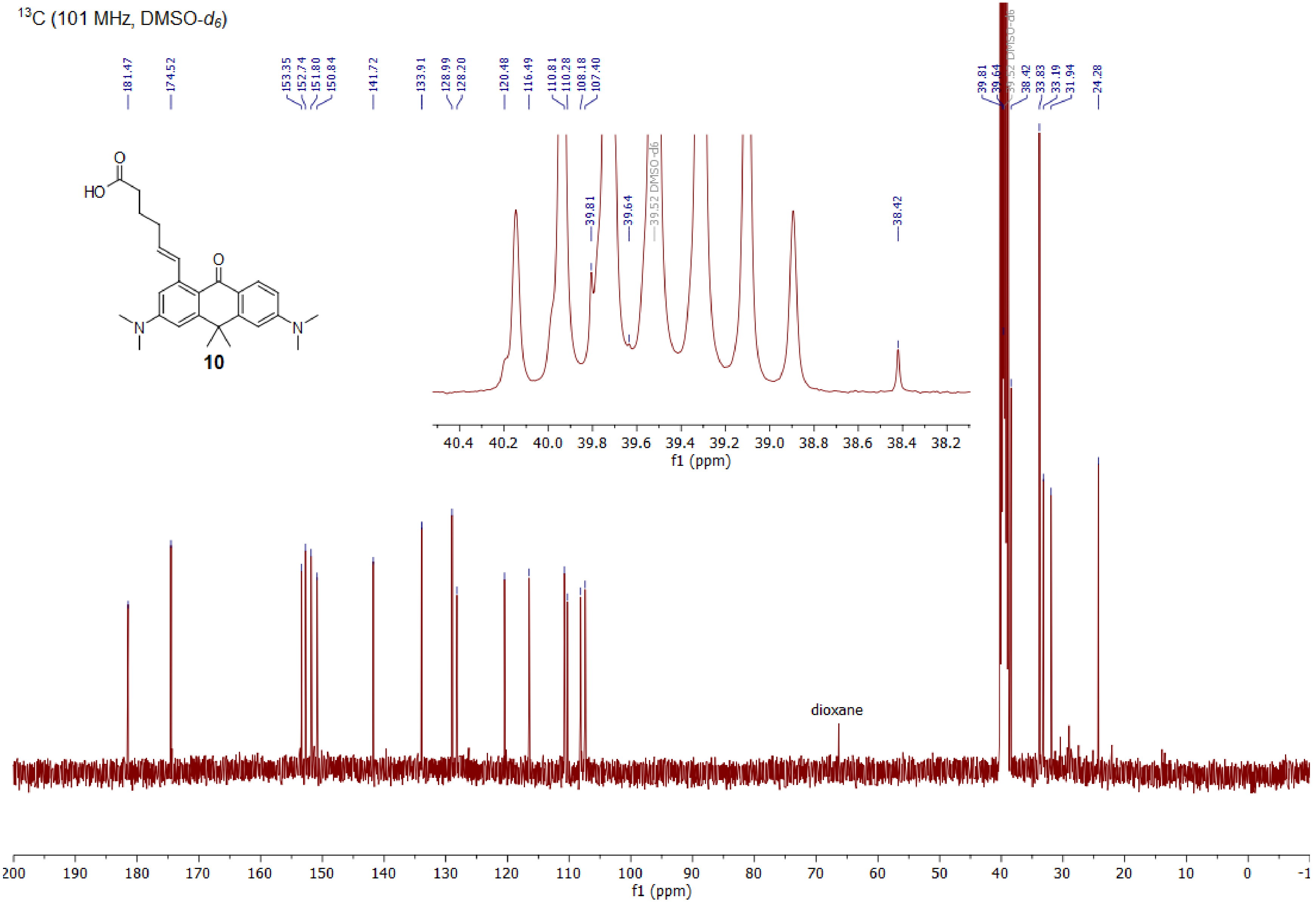

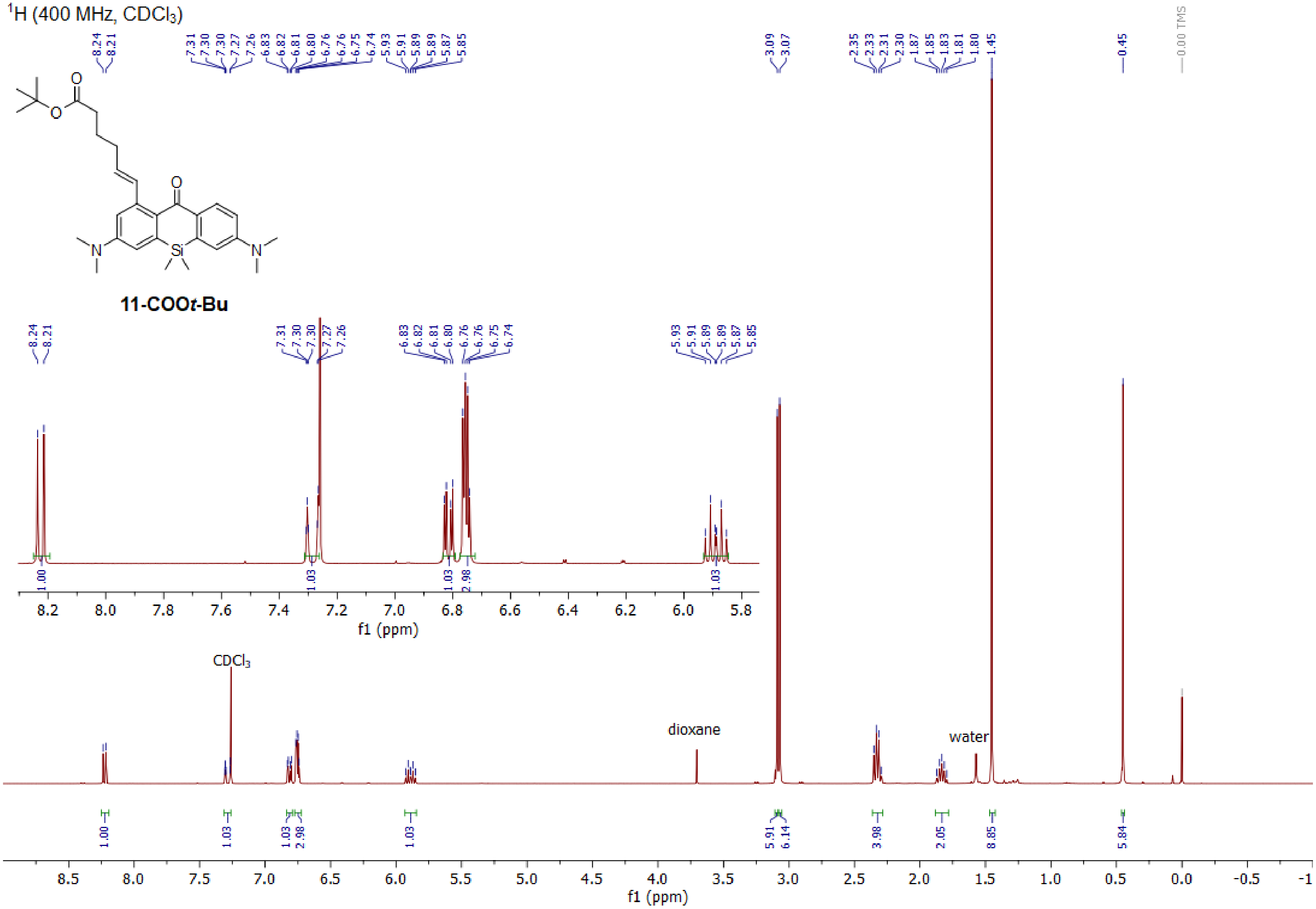

1H (400 MHz, CDCl3)
8.24
8.21
7.31
7.30
7.30
7.27
7.26
6.83
6.82
6.81
6.80
6.76
6.76
6.75
6.74
5.93
5.91
5.89
5.89
5.87
5.85
3.09
3.07
2.35
2.33
2.31
2.30
1.87
1.85
1.83
1.81
1.80
1.45
0.45
0.00 TMS
11-COOt-Bu
CDCl3
dioxane
water
1.00
1.03
1.03
2.98
1.03
5.91
6.14
3.98
2.05
8.85
5.84
f1 (ppm)

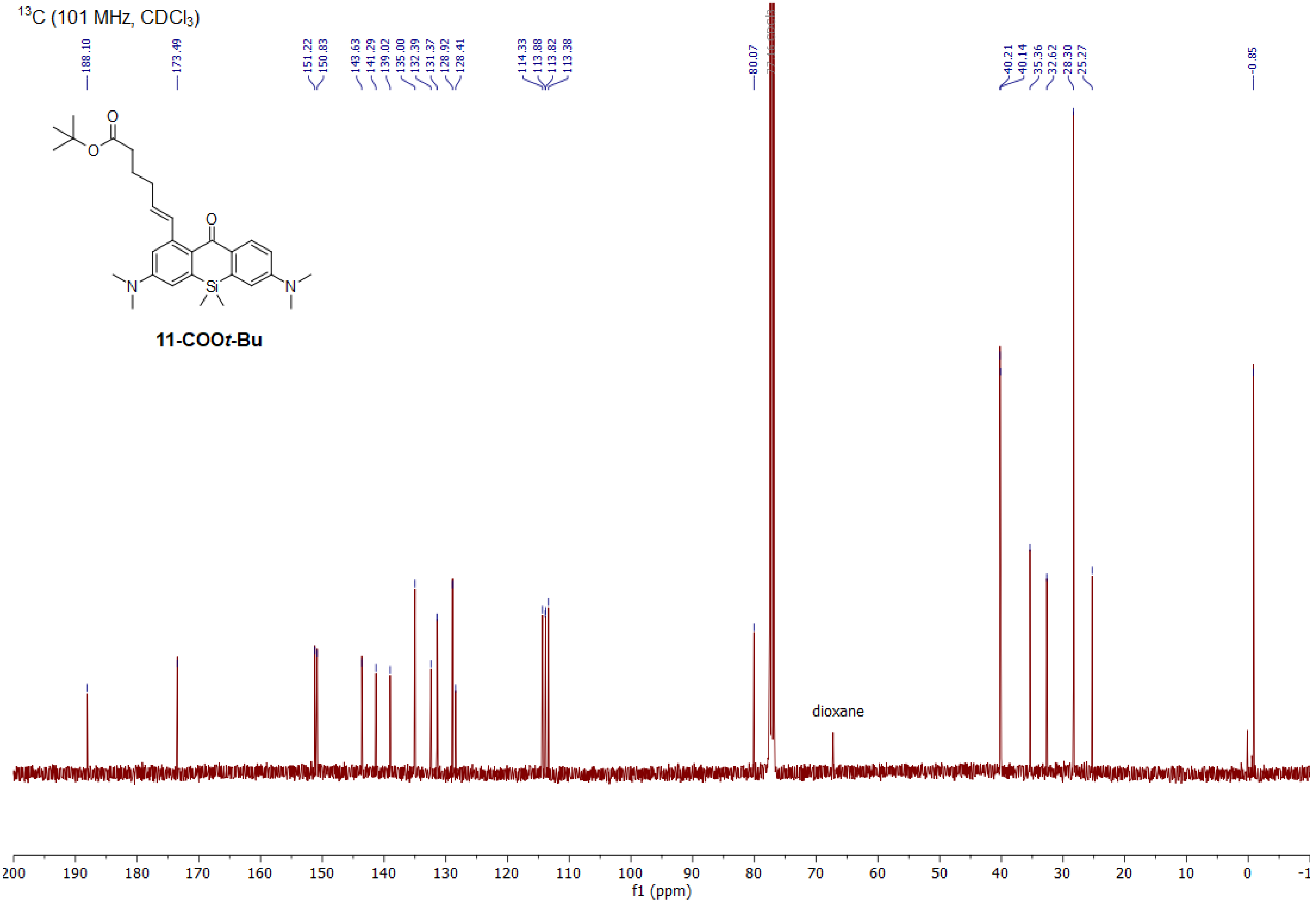

13C (101 MHz, CDCl3)
188.10
173.49
151.22
150.83
143.63
141.29
139.02
135.00
132.39
131.37
128.92
128.41
114.33
113.88
113.82
113.38
80.07
40.21
40.14
35.36
32.62
28.30
25.27
-0.85
11-COOt-Bu
dioxane
f1 (ppm)

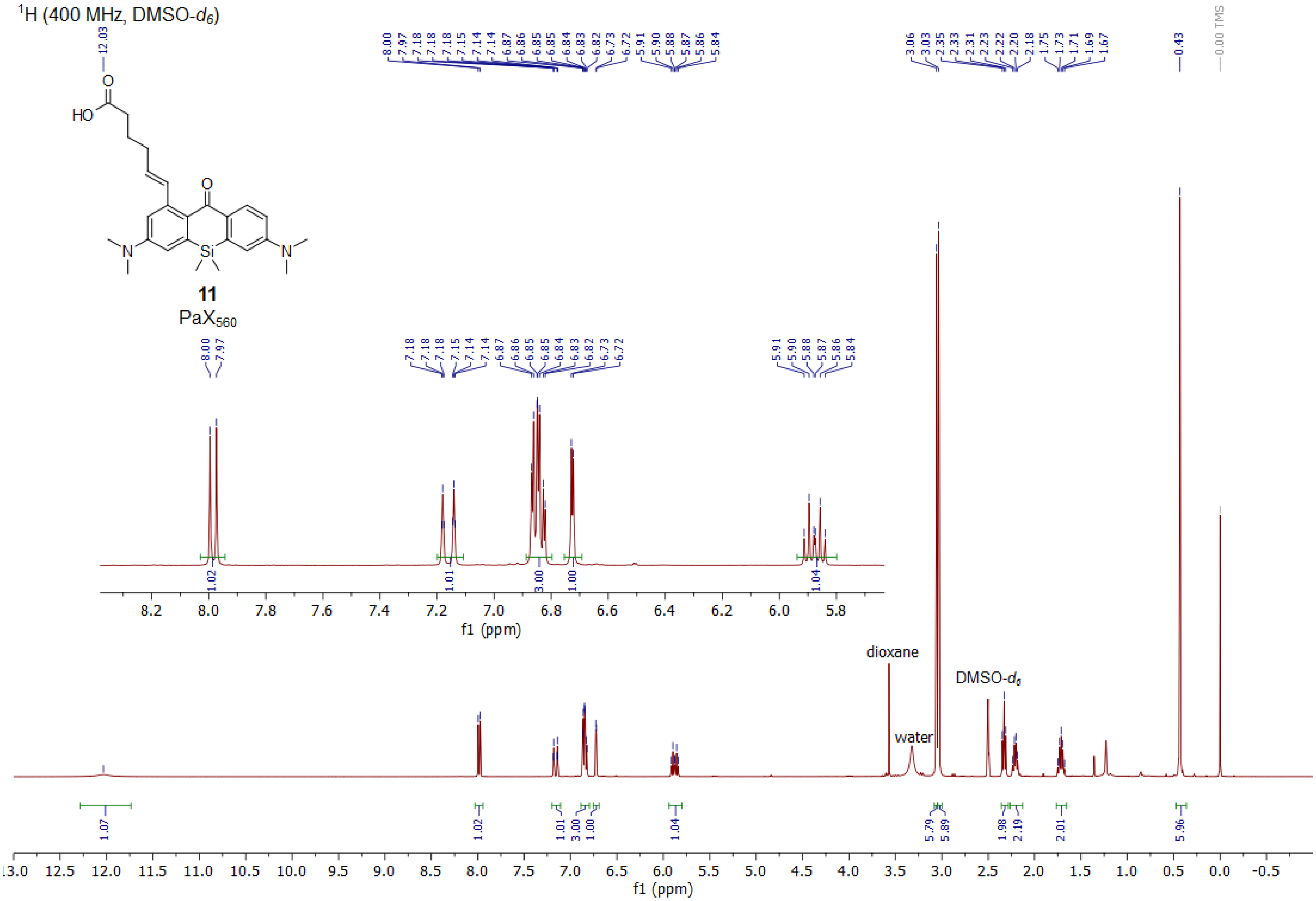

1H (400 MHz, DMSO-d6)
11
PaX560
dioxane
water
DMSO-d6
f1 (ppm)

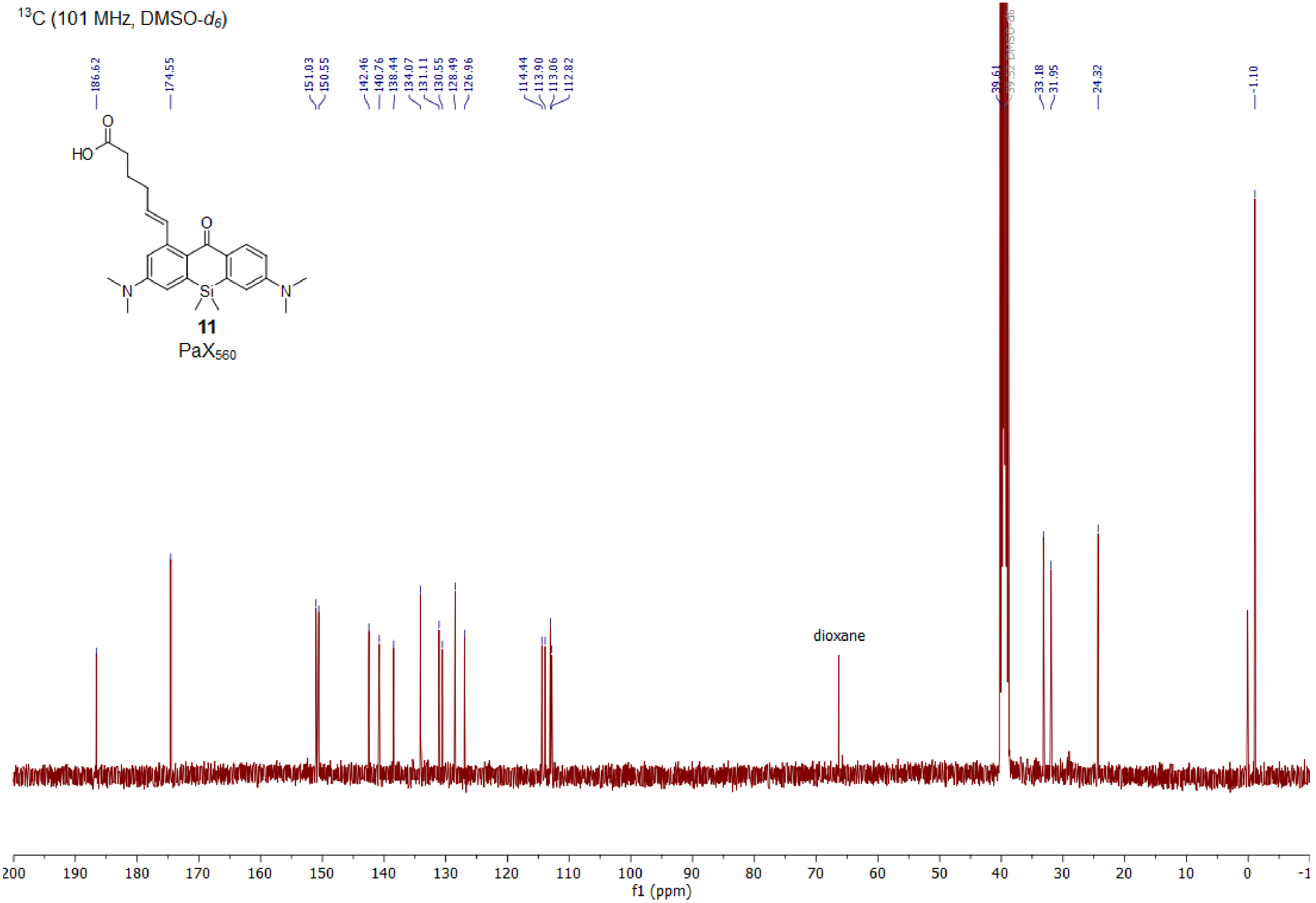

13C (101 MHz, DMSO-d6)
186.62
174.55
151.03
150.55
142.46
140.76
138.44
134.07
131.11
130.55
128.49
126.96
114.44
113.90
113.06
112.82
39.61
39.52 DMSO-d6
33.18
31.95
24.32
-1.10
HO
O
N
Si
11
PaX560
dioxane
200
190
180
170
160
150
140
130
120
110
100
90
80
70
60
50
40
30
20
10
0
-1
f1 (ppm)

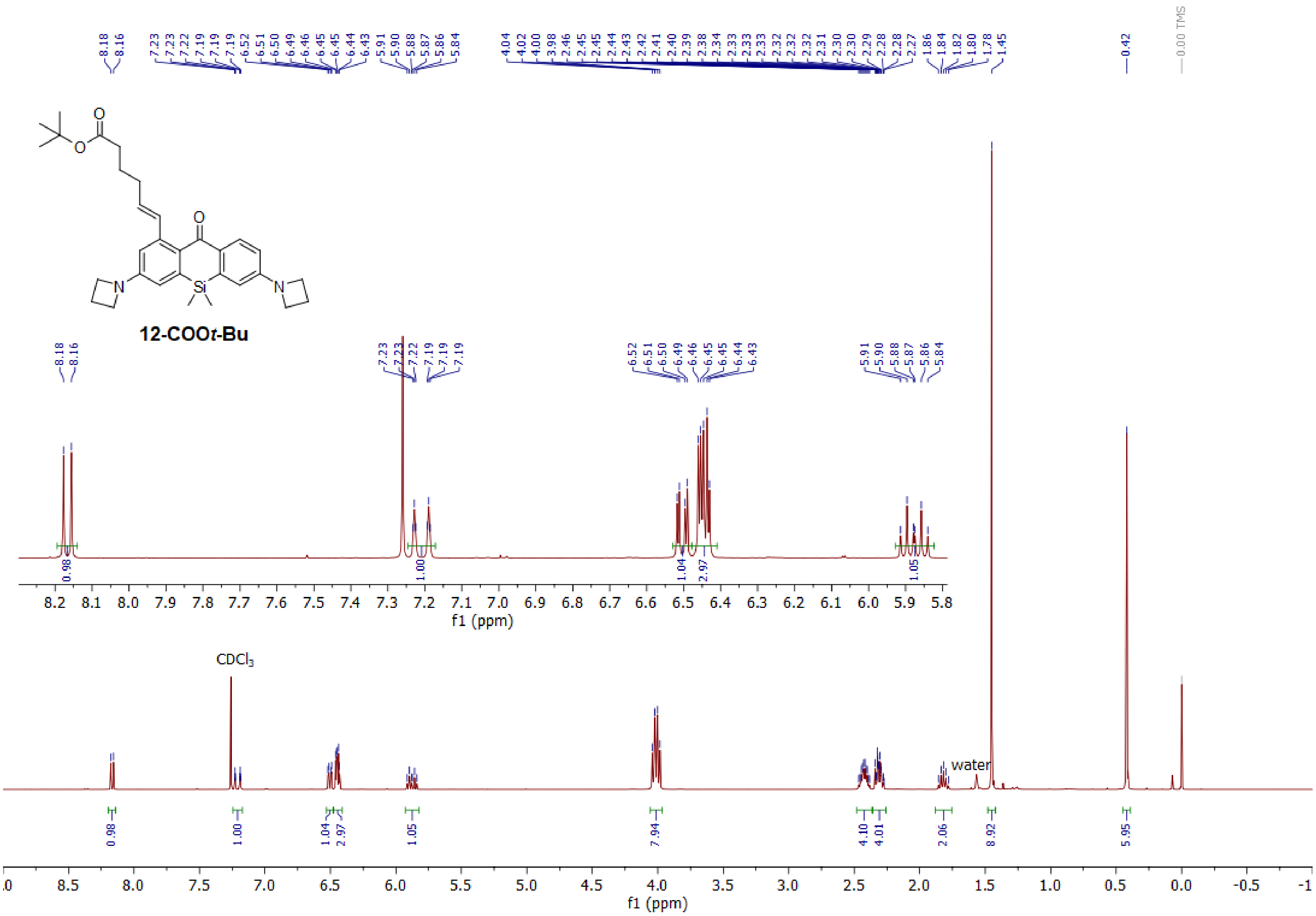
12-COO*t*-Bu
CDCl3
water
0.00 TMS
f1 (ppm)

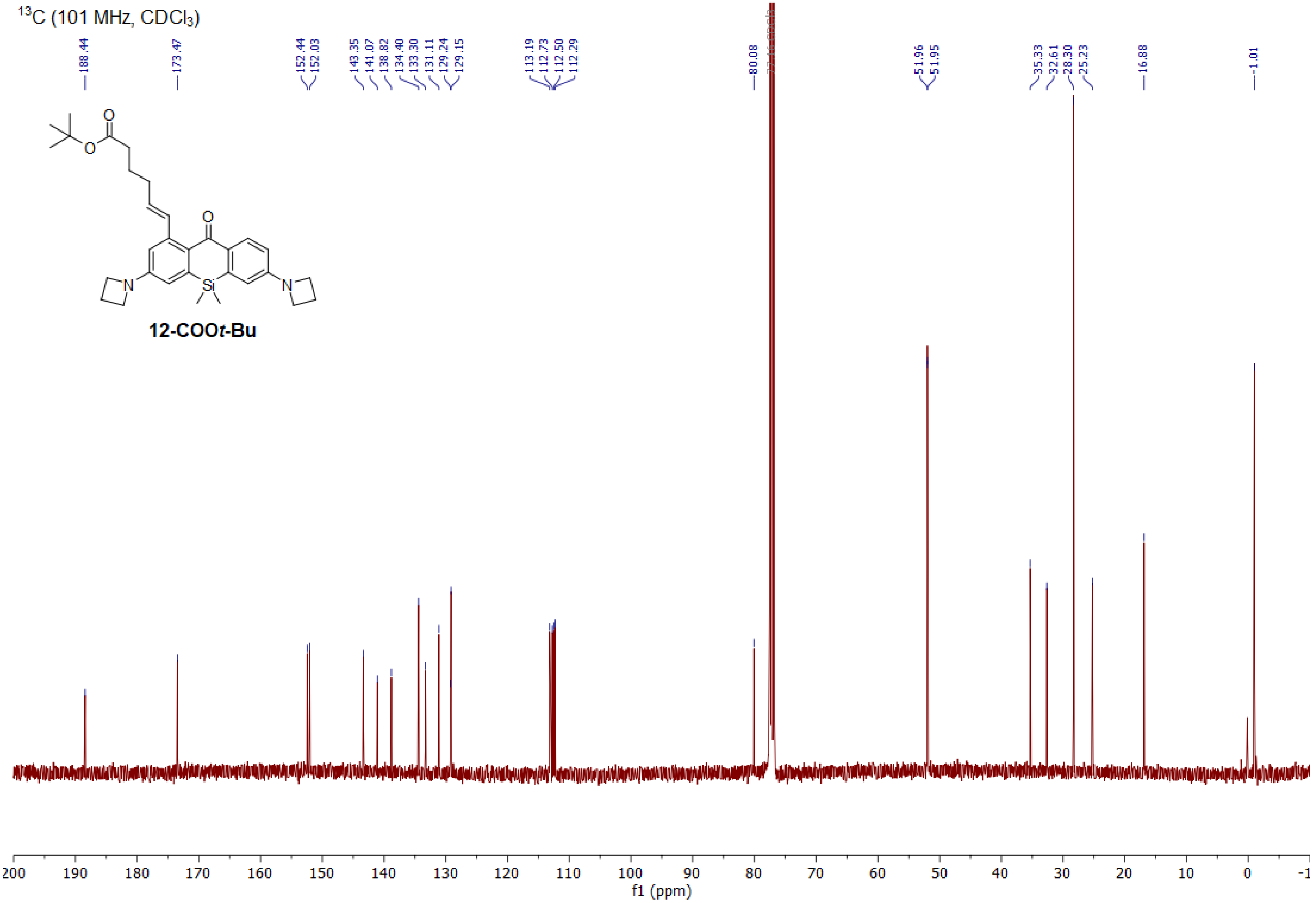

$^{13}C$ (101 MHz, $CDCl_3$)
188.44
173.47
152.44
152.03
143.35
141.07
138.82
134.40
133.30
131.11
129.24
129.15
113.19
112.73
112.50
112.29
80.08
51.96
51.95
35.33
32.61
28.30
25.23
16.88
-1.01
O
O
O
N
Si
N
12-COOt-Bu
200
190
180
170
160
150
140
130
120
110
100
90
80
70
60
50
40
30
20
10
0
-1
f1 (ppm)

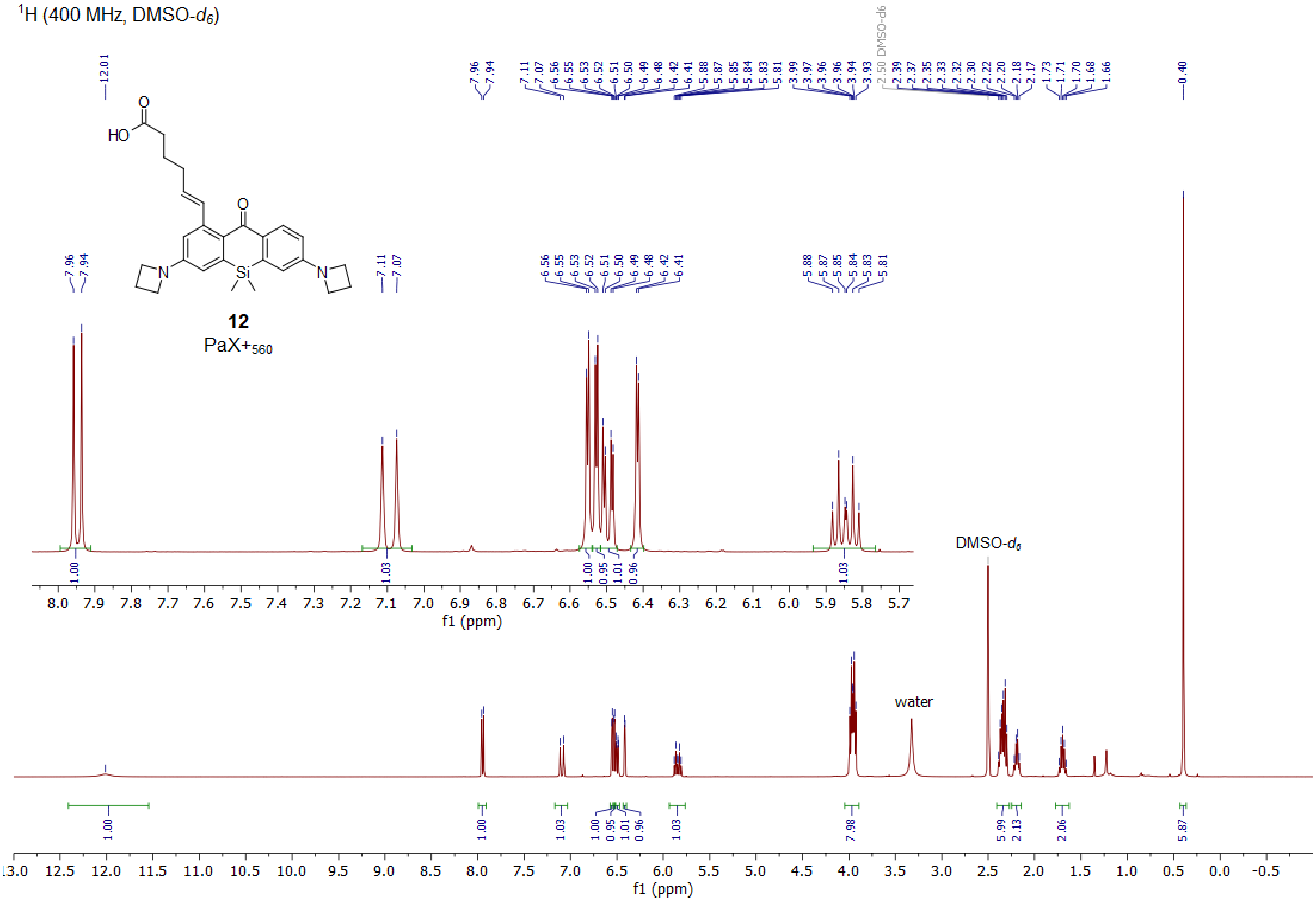

1H (400 MHz, DMSO-d6)
HO
O
O
N
Si
N
12
PaX+560
DMSO-d6
water
f1 (ppm)
f1 (ppm)

$^{13}C$ (101 MHz, DMSO-$d_6$)

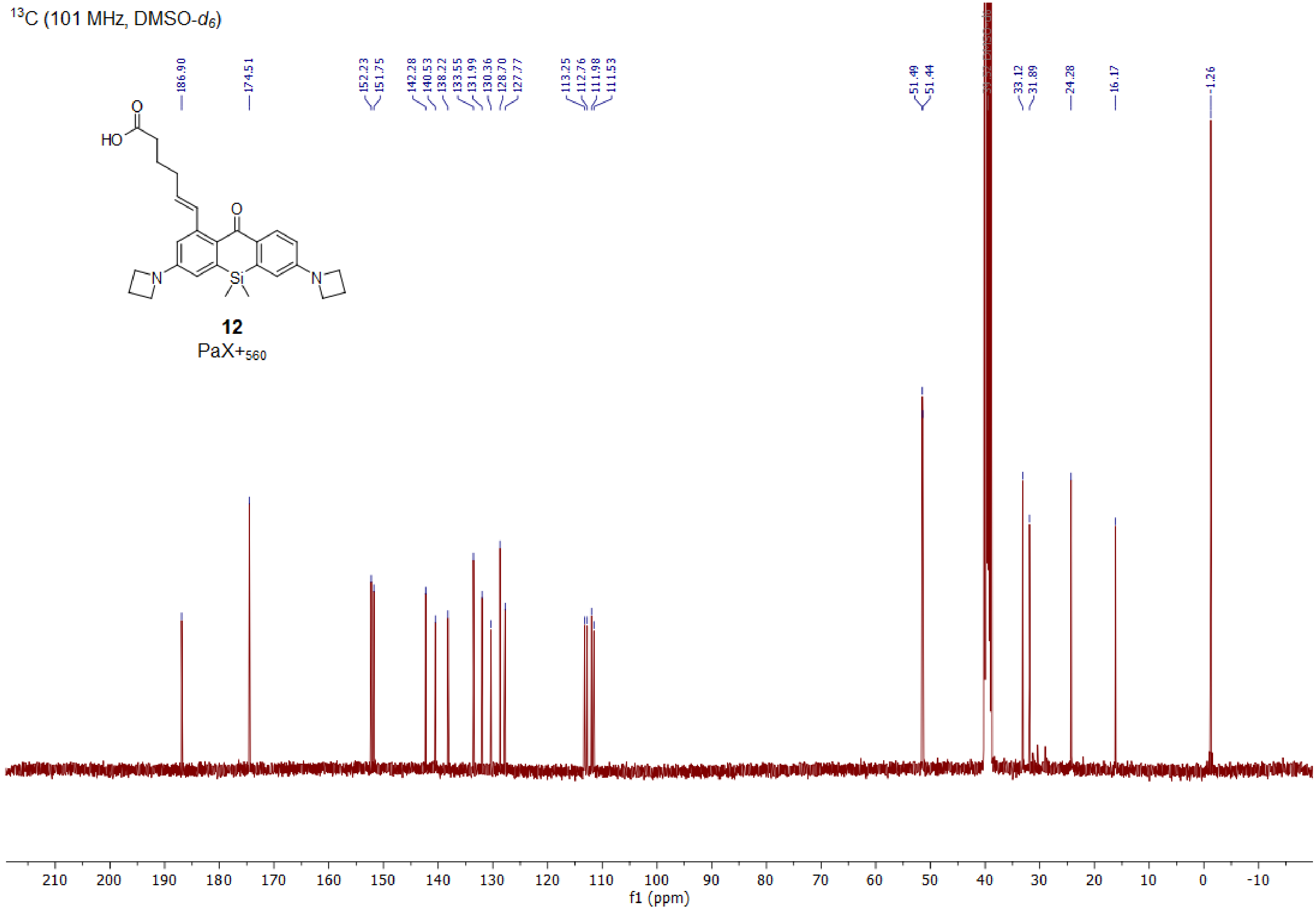

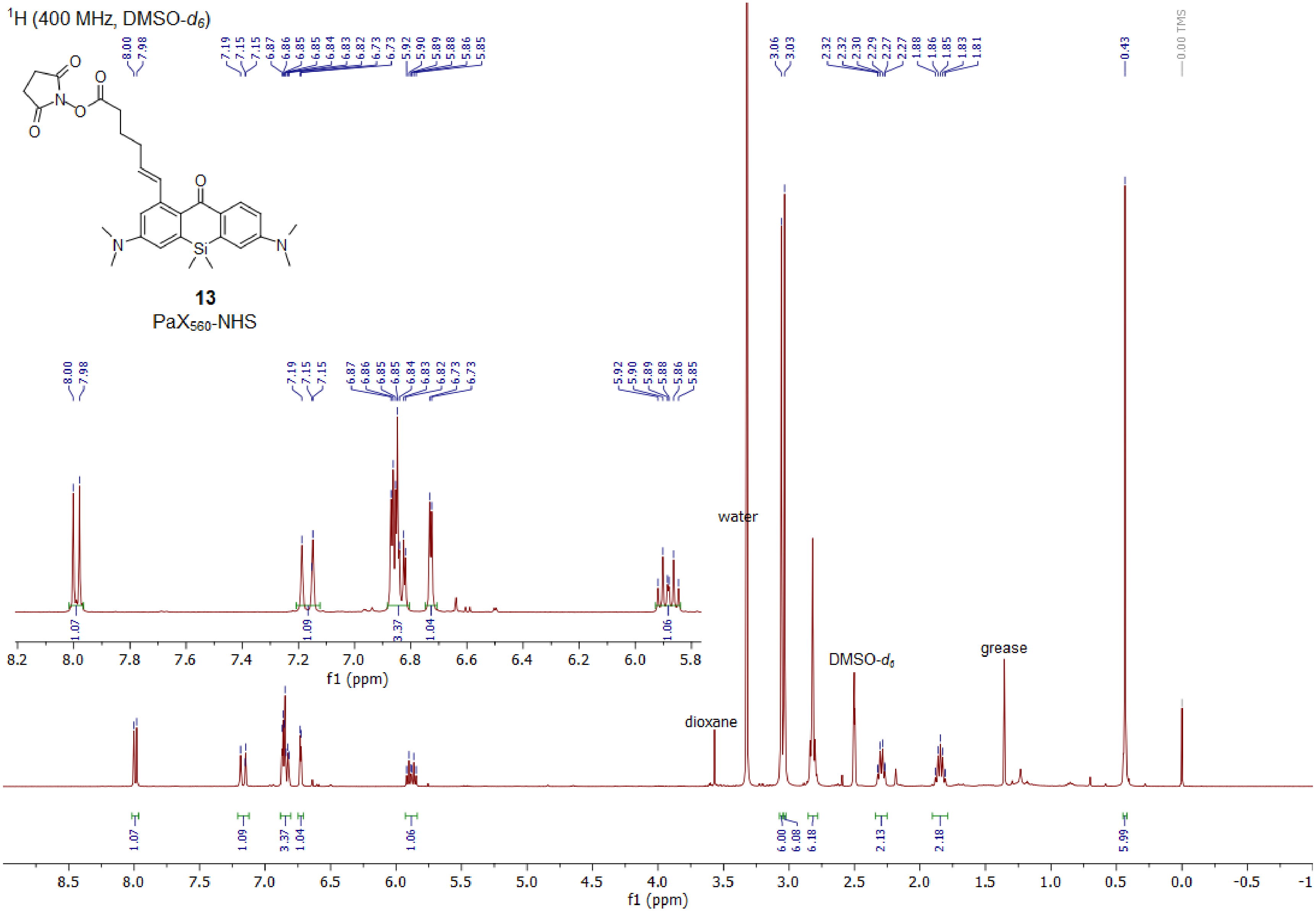

¹H (400 MHz, DMSO-d6)
13
PaX560-NHS
water
dioxane
DMSO-d6
grease
0.00 TMS
f1 (ppm)

[13]C (101 MHz, DMSO-$d_6$)

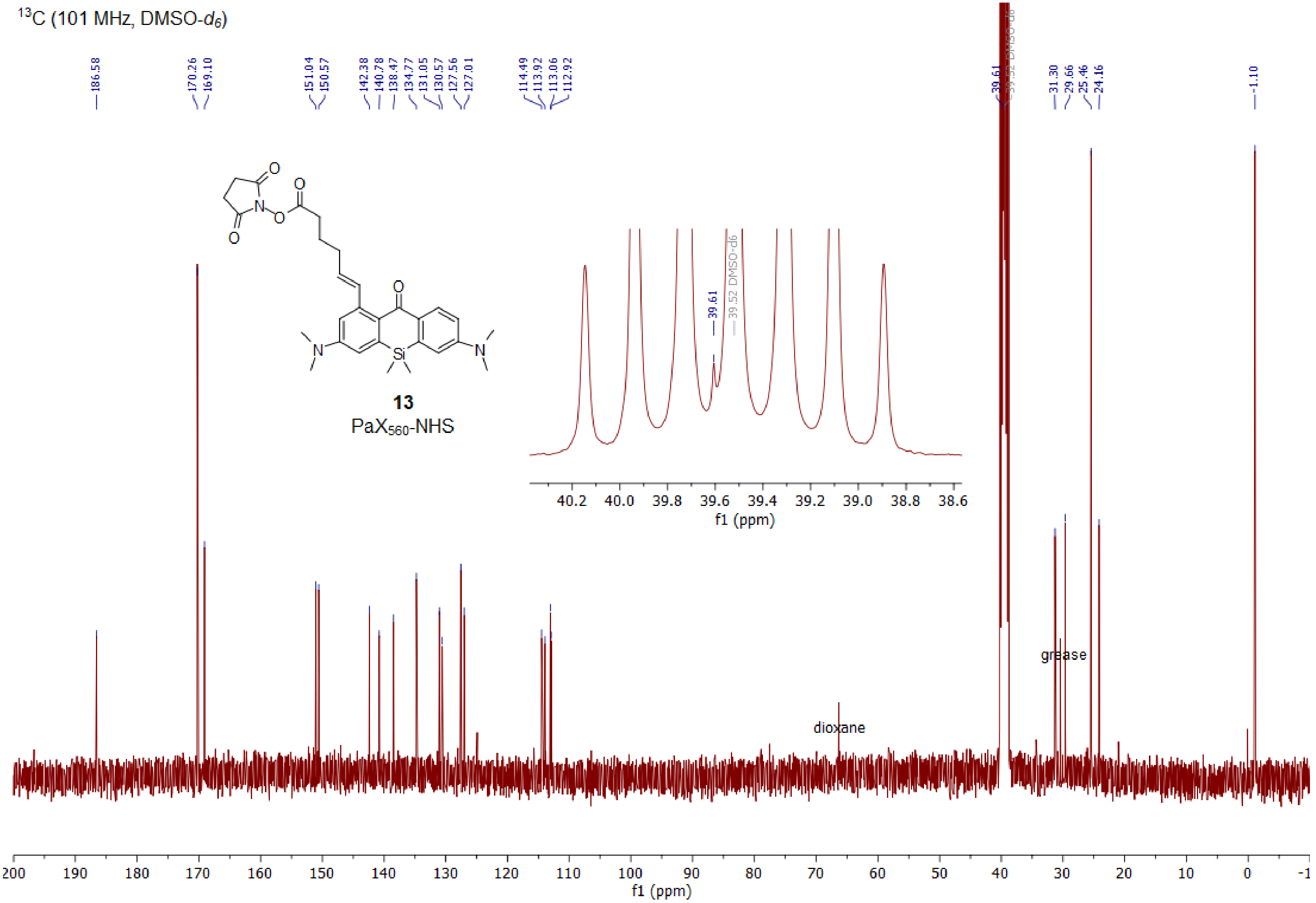

$^{1}H$ (400 MHz, $CDCl_3$)

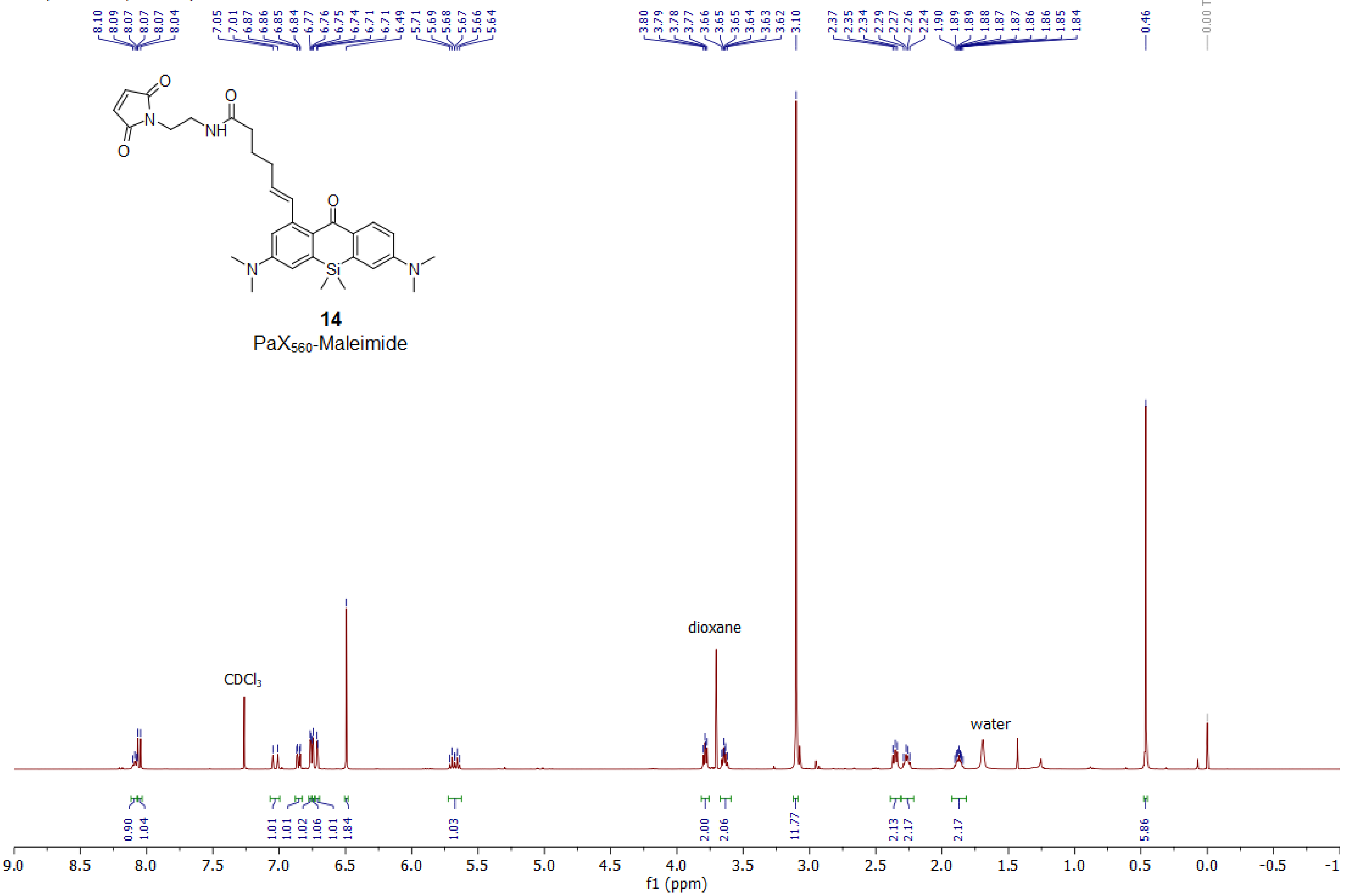

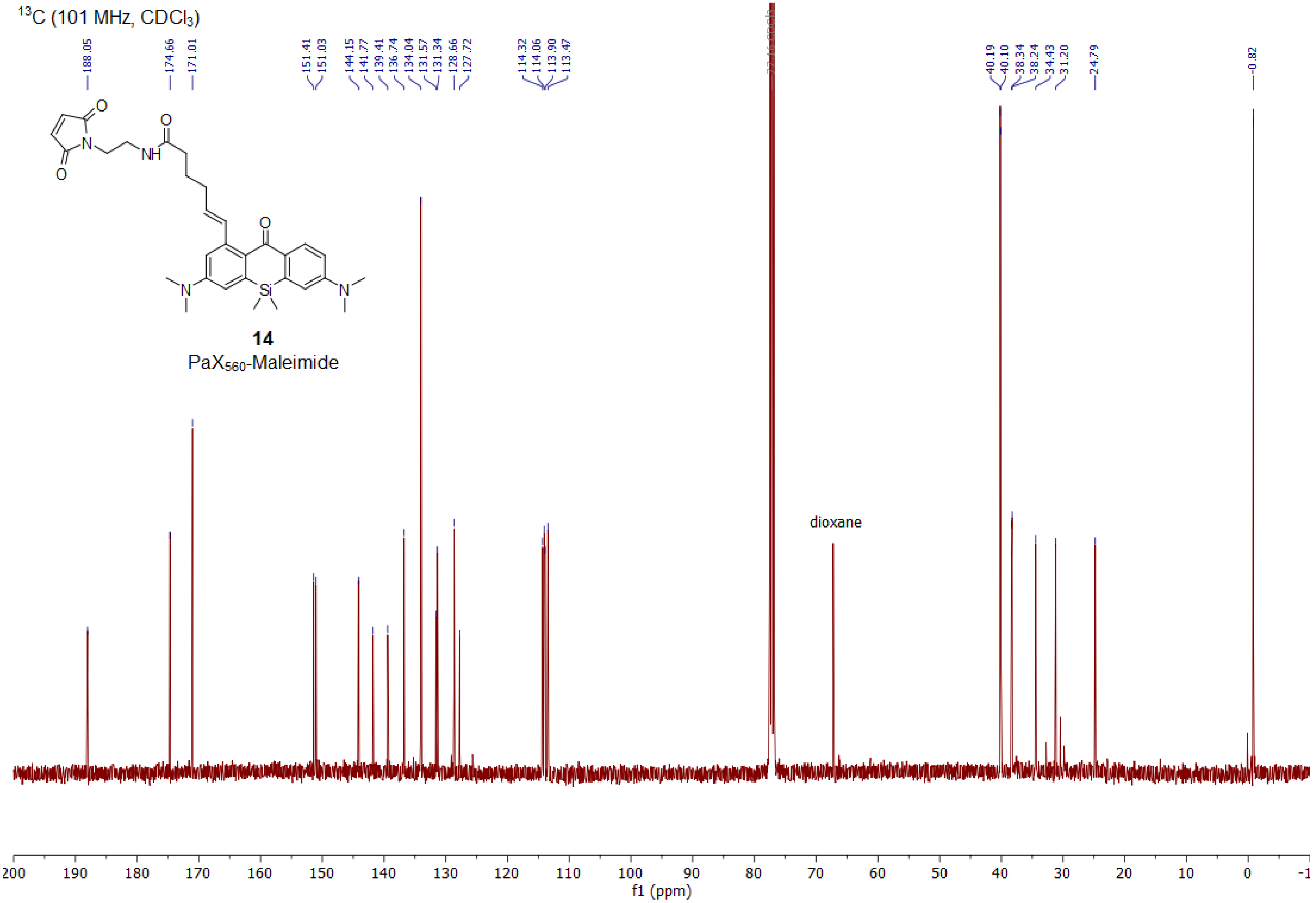

$^{13}C$ (101 MHz, $CDCl_3$)
188.05
174.66
171.01
151.41
151.03
144.15
141.77
139.41
136.74
134.04
131.57
131.34
128.66
127.72
114.32
114.06
113.90
113.47
40.19
40.10
38.34
38.24
34.43
31.20
24.79
-0.82
14
PaX560-Maleimide
dioxane
f1 (ppm)

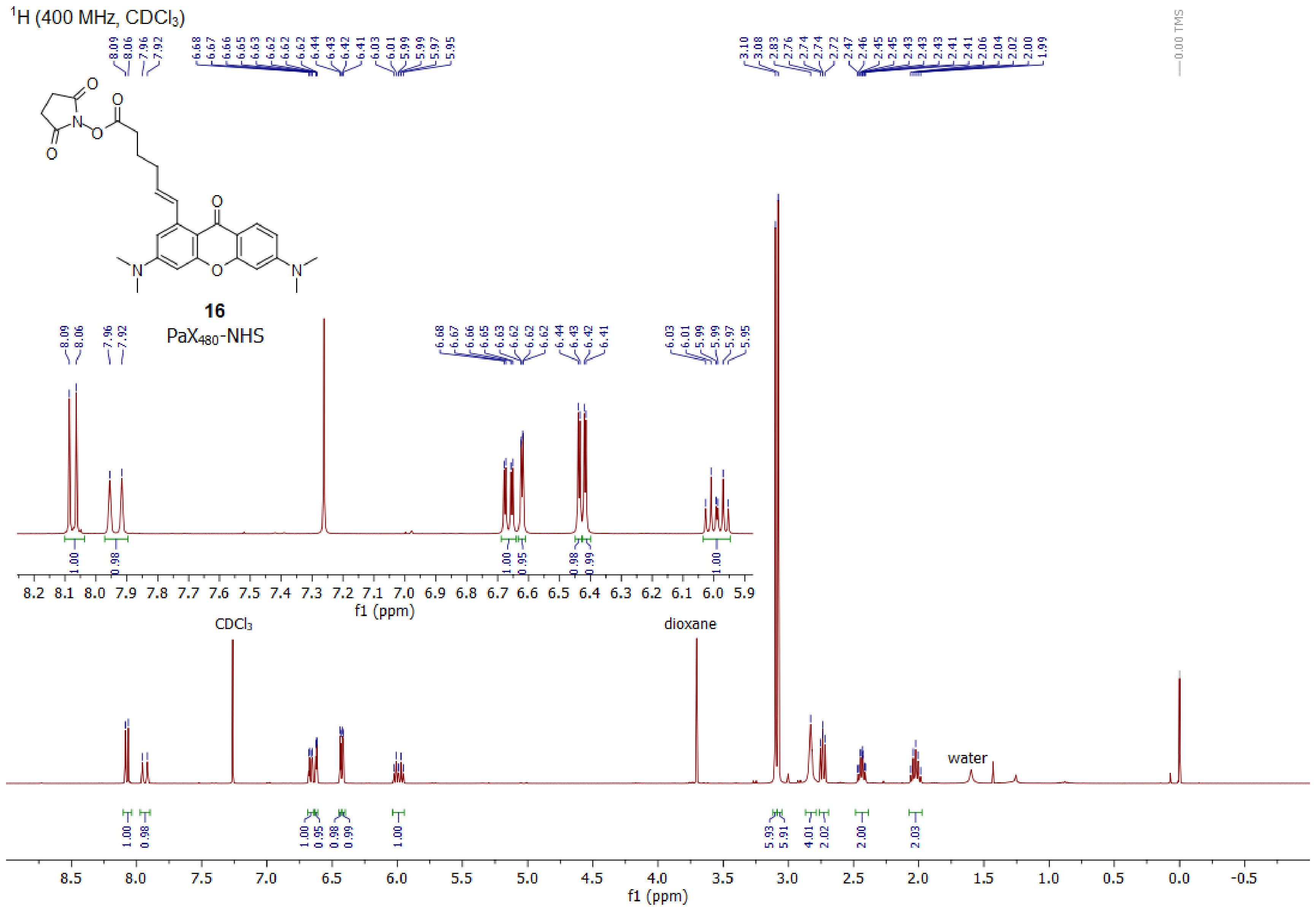

¹H (400 MHz, CDCl₃)
16
PaX480-NHS
CDCl3
dioxane
water
f1 (ppm)

$^{13}C$ (101 MHz, $CDCl_3$)

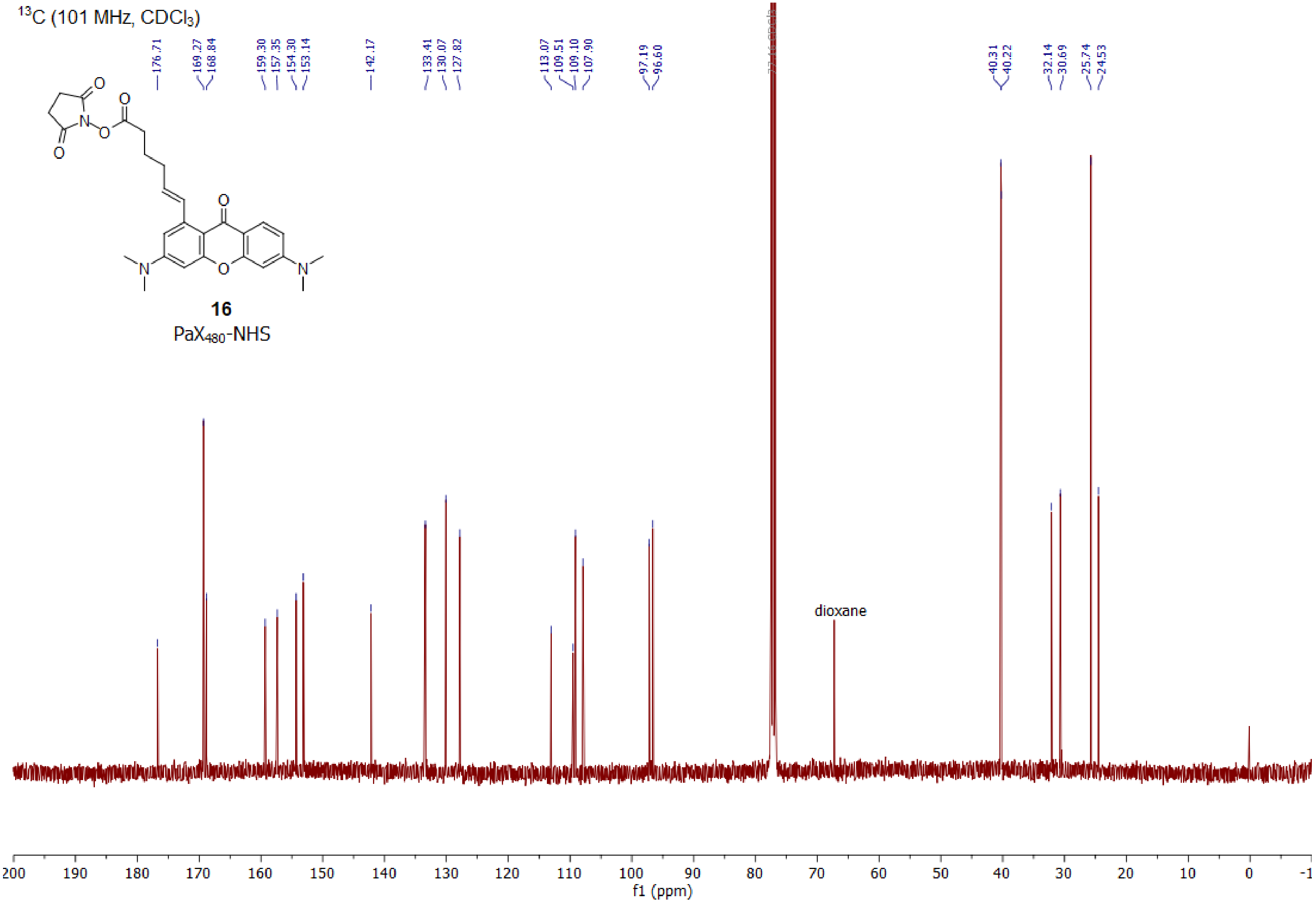

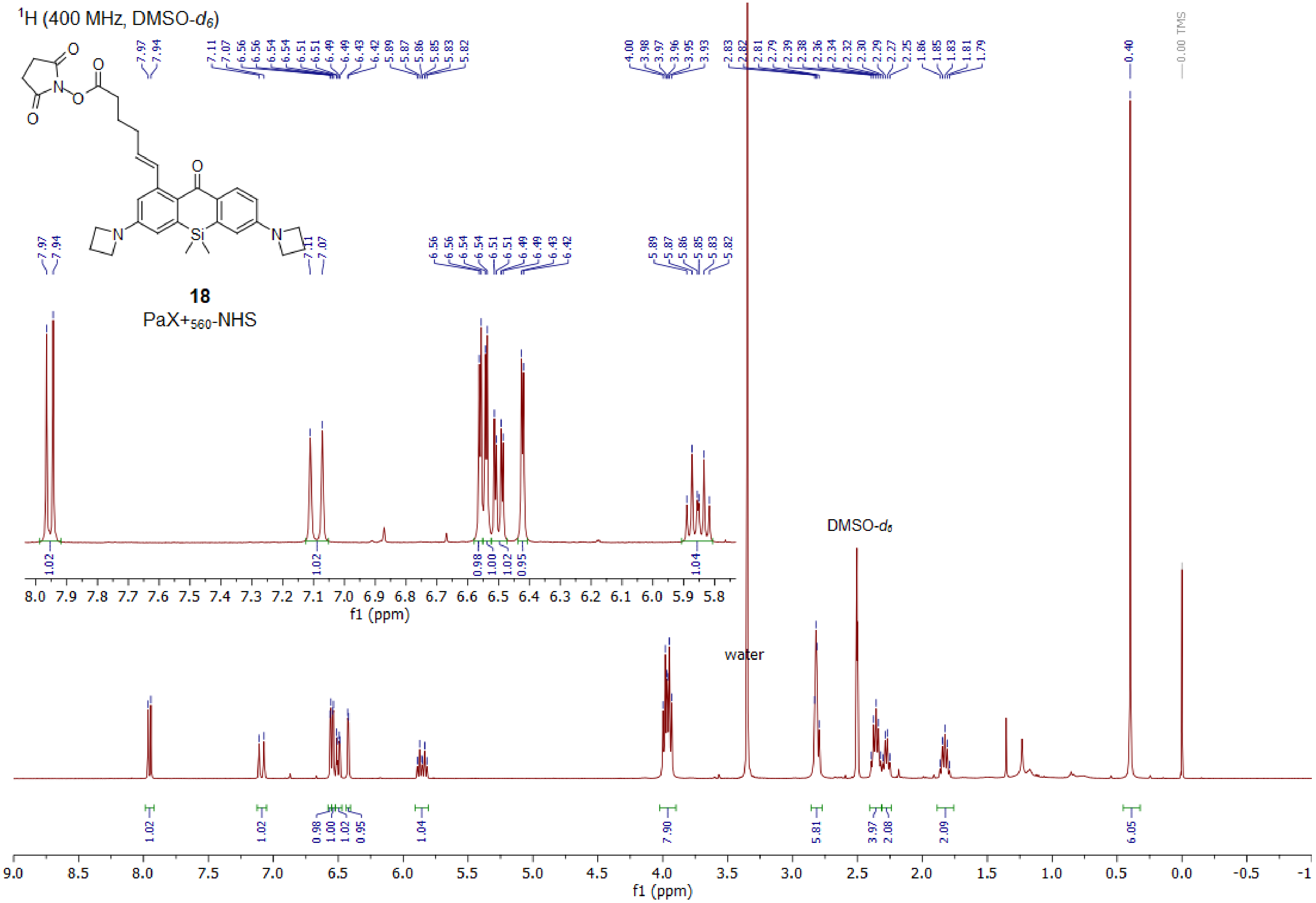

1H (400 MHz, DMSO-d6)
18
PaX+560-NHS
water
DMSO-d6
f1 (ppm)

$^{13}C$ (101 MHz, DMSO-$d_6$)

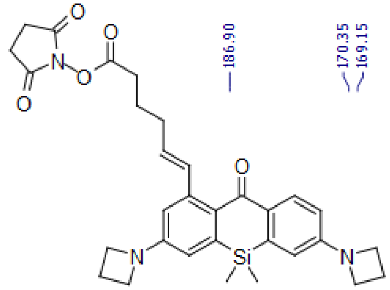

**18**

PaX+$_{560}$-NHS

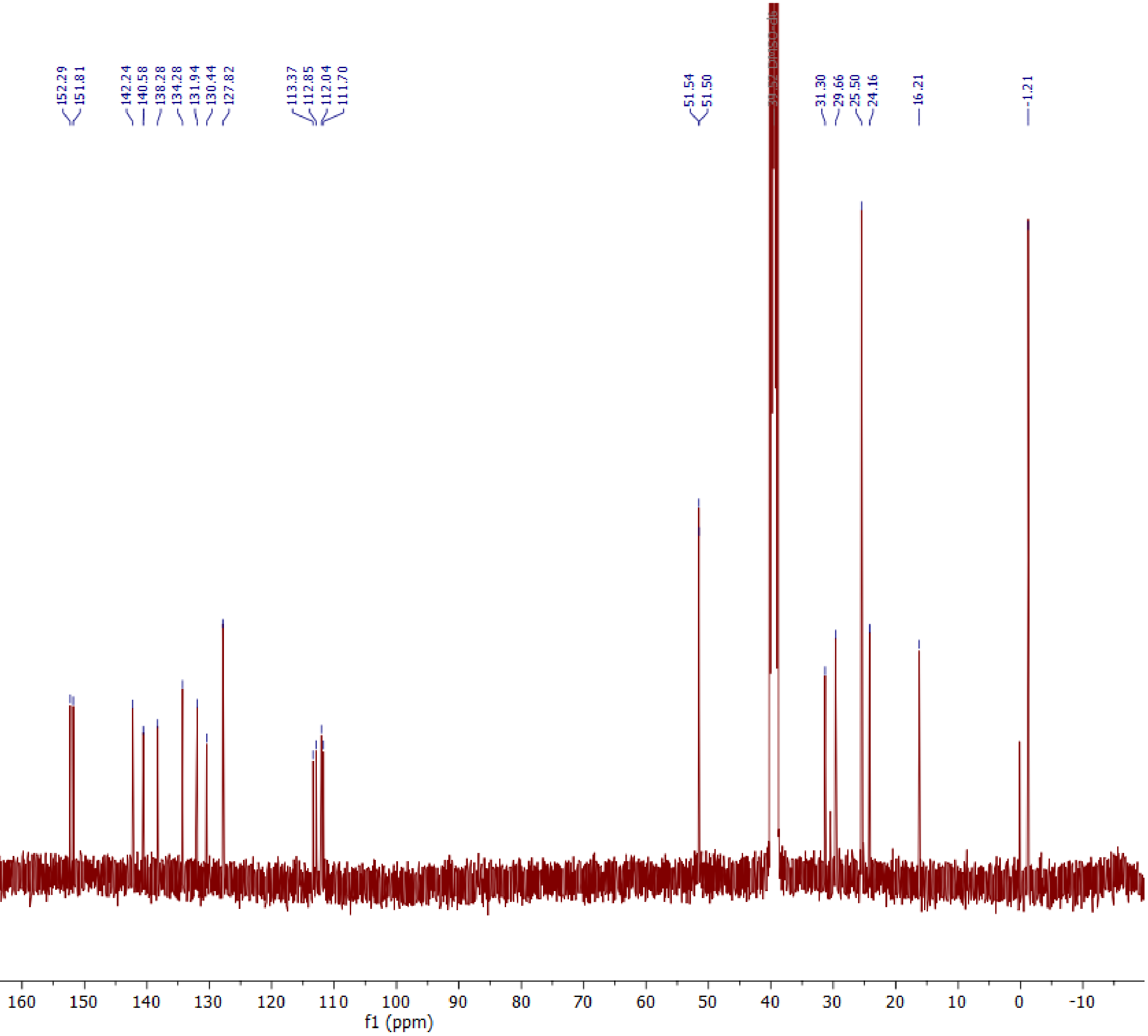

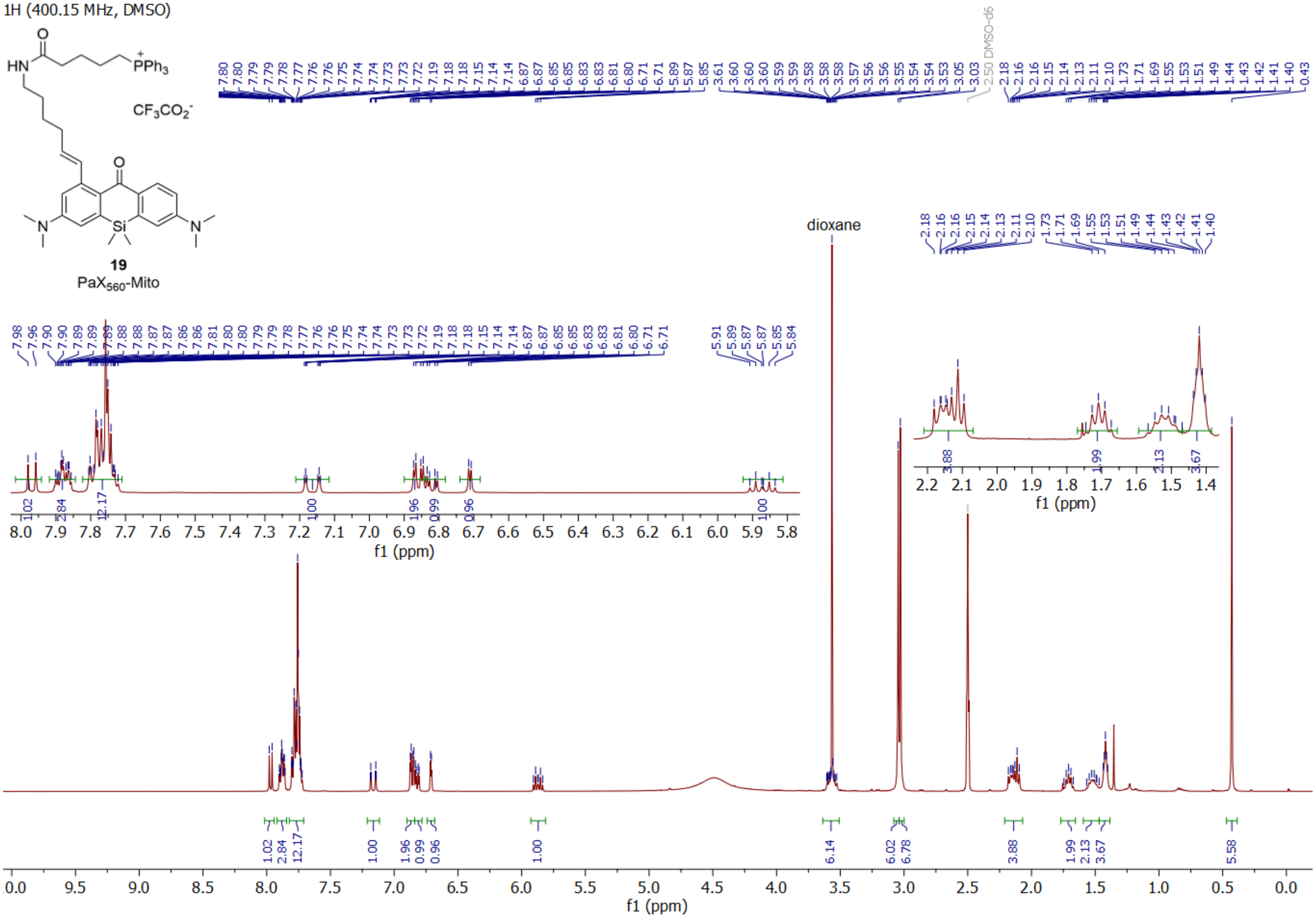
1H (400.15 MHz, DMSO)
HN
PPh3
CF3CO2-
N
Si
N
19
PaX560-Mito
dioxane
2.50 DMSO-d6
f1 (ppm)

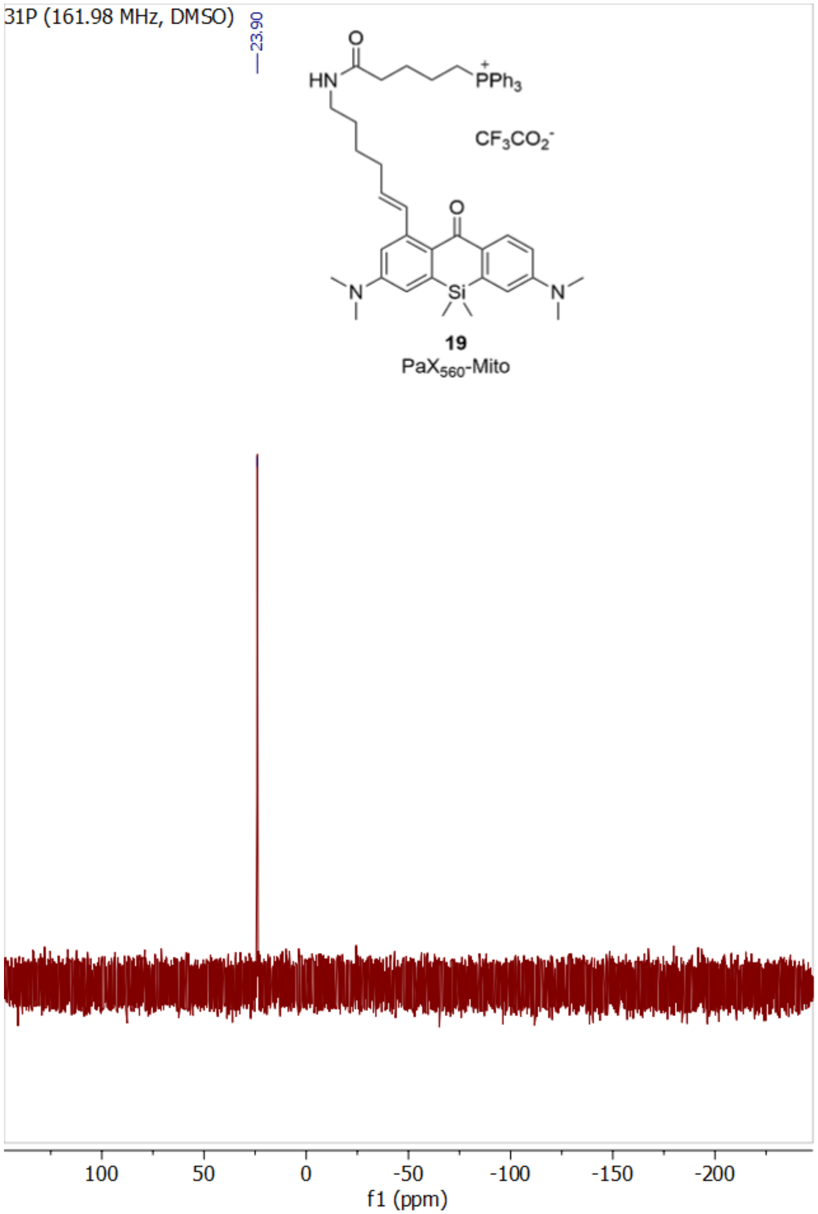
31P (161.98 MHz, DMSO)
23.90
HN
O
PPh3
CF3CO2-
O
N
Si
N
19
PaX560-Mito
100
50
0
-50
-100
-150
-200
f1 (ppm)

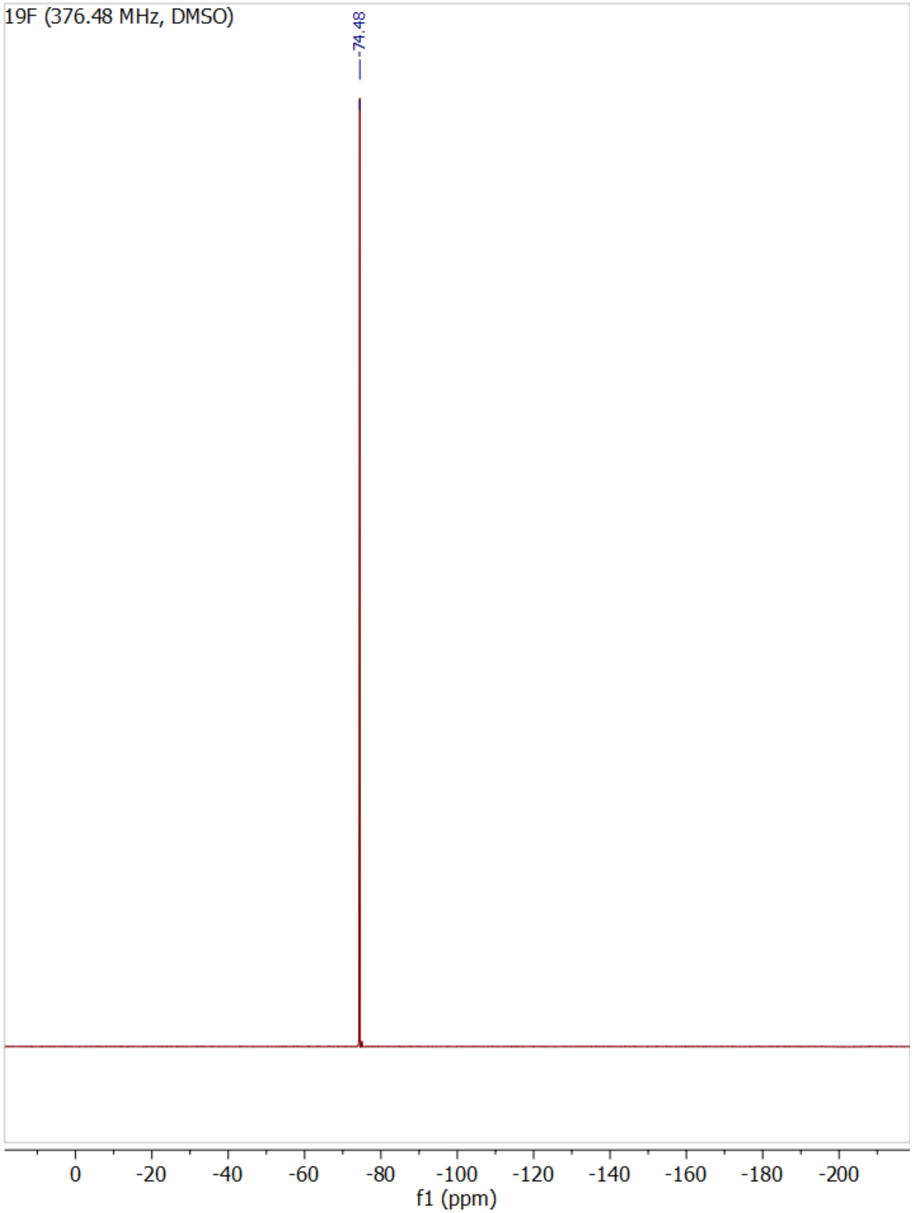
19F (376.48 MHz, DMSO)
-74.48
0
-20
-40
-60
-80
-100
-120
-140
-160
-180
-200
f1 (ppm)

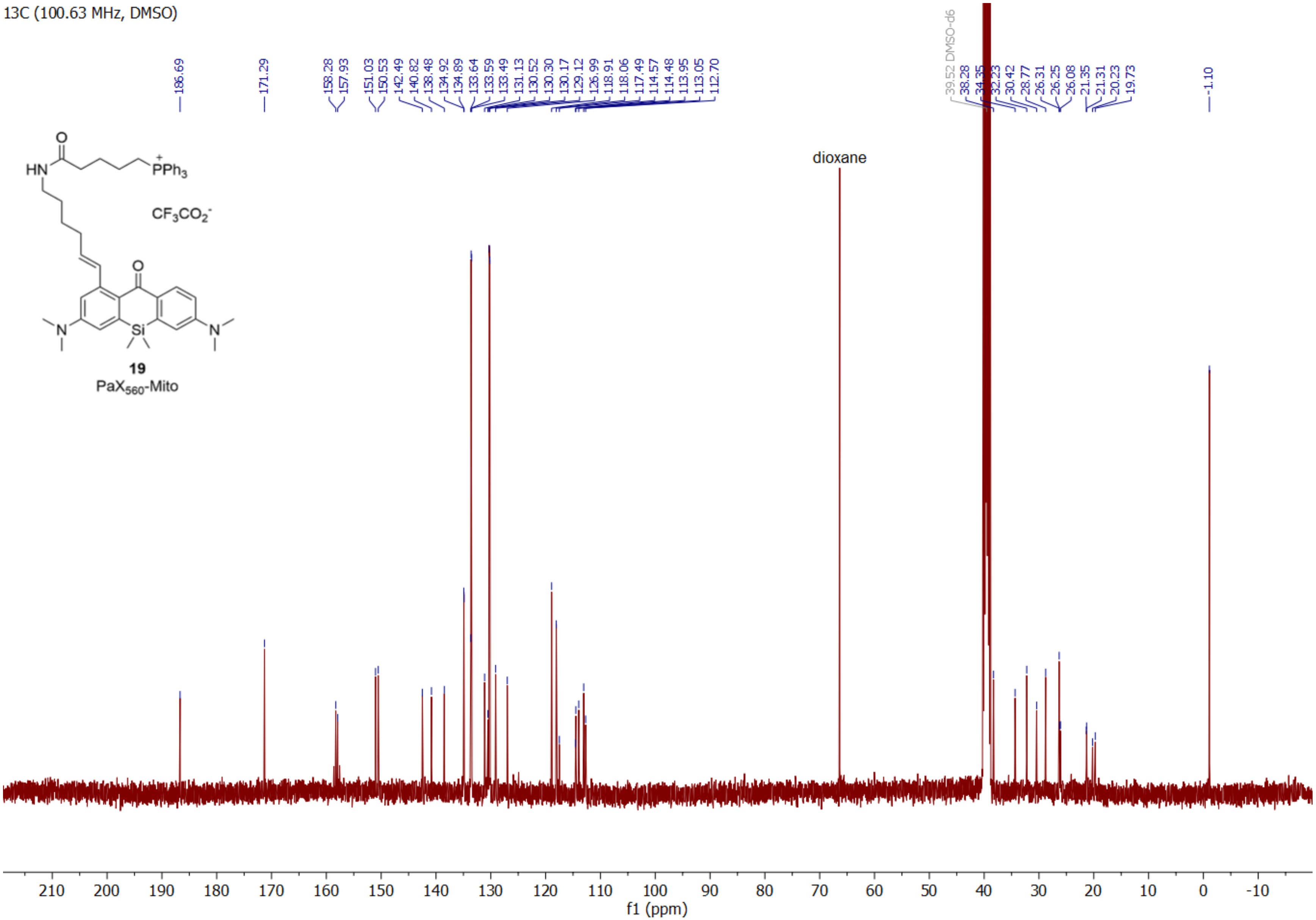

13C (100.63 MHz, DMSO)
186.69
171.29
158.28
157.93
151.03
150.53
142.49
140.82
138.48
134.92
134.89
133.64
133.59
133.49
131.13
130.52
130.30
130.17
129.12
126.99
118.91
118.06
117.49
114.57
114.48
113.95
113.05
112.70
39.52 DMSO-d6
38.28
34.35
32.23
30.42
28.77
26.31
26.25
26.08
21.35
21.31
20.23
19.73
-1.10
dioxane
HN
PPh3
CF3CO2-
N
Si
N
19
PaX560-Mito
f1 (ppm)

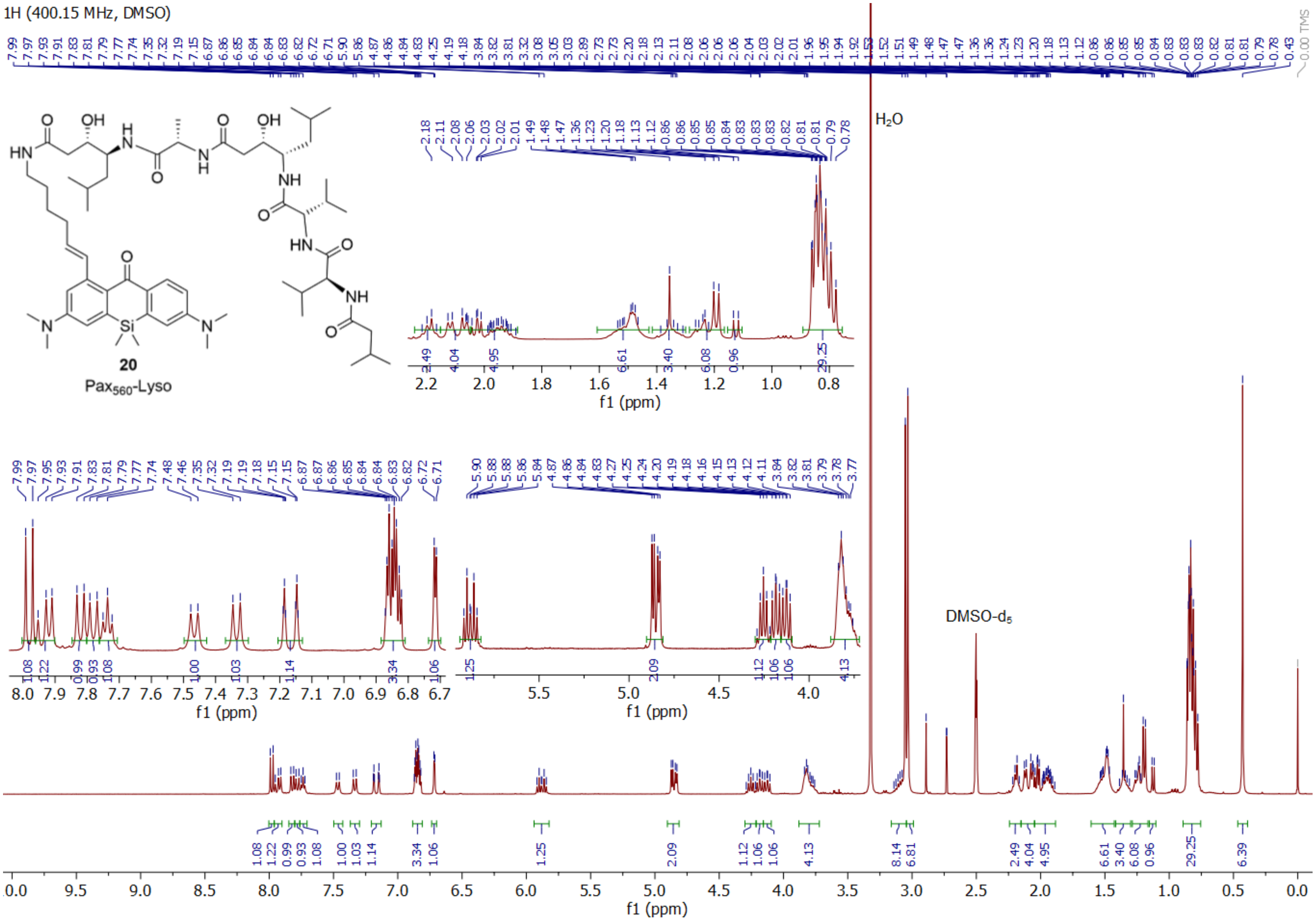
1H (400.15 MHz, DMSO)
20
Pax560-Lyso
H2O
DMSO-d5
f1 (ppm)

13C (100.63 MHz, DMSO)

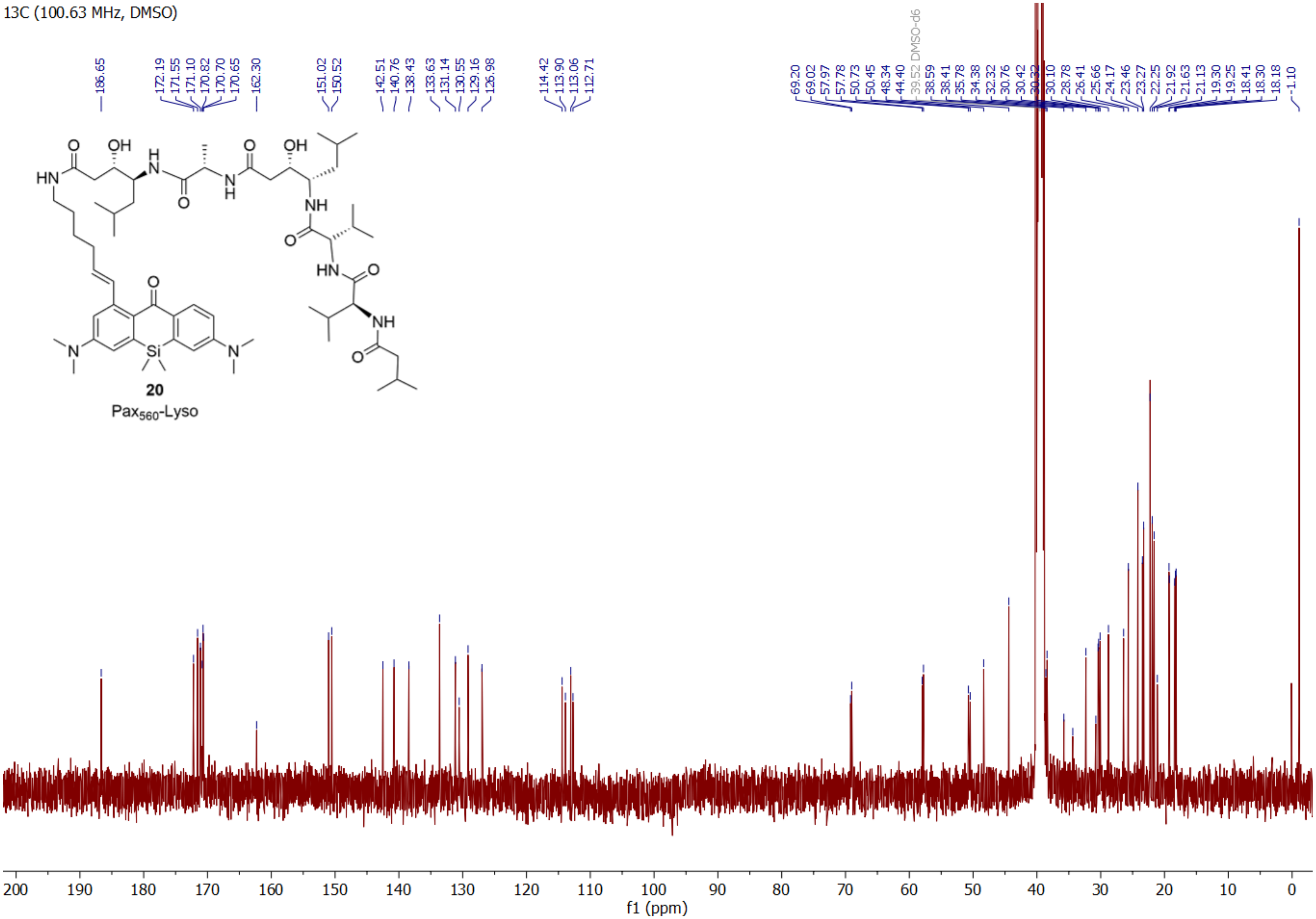

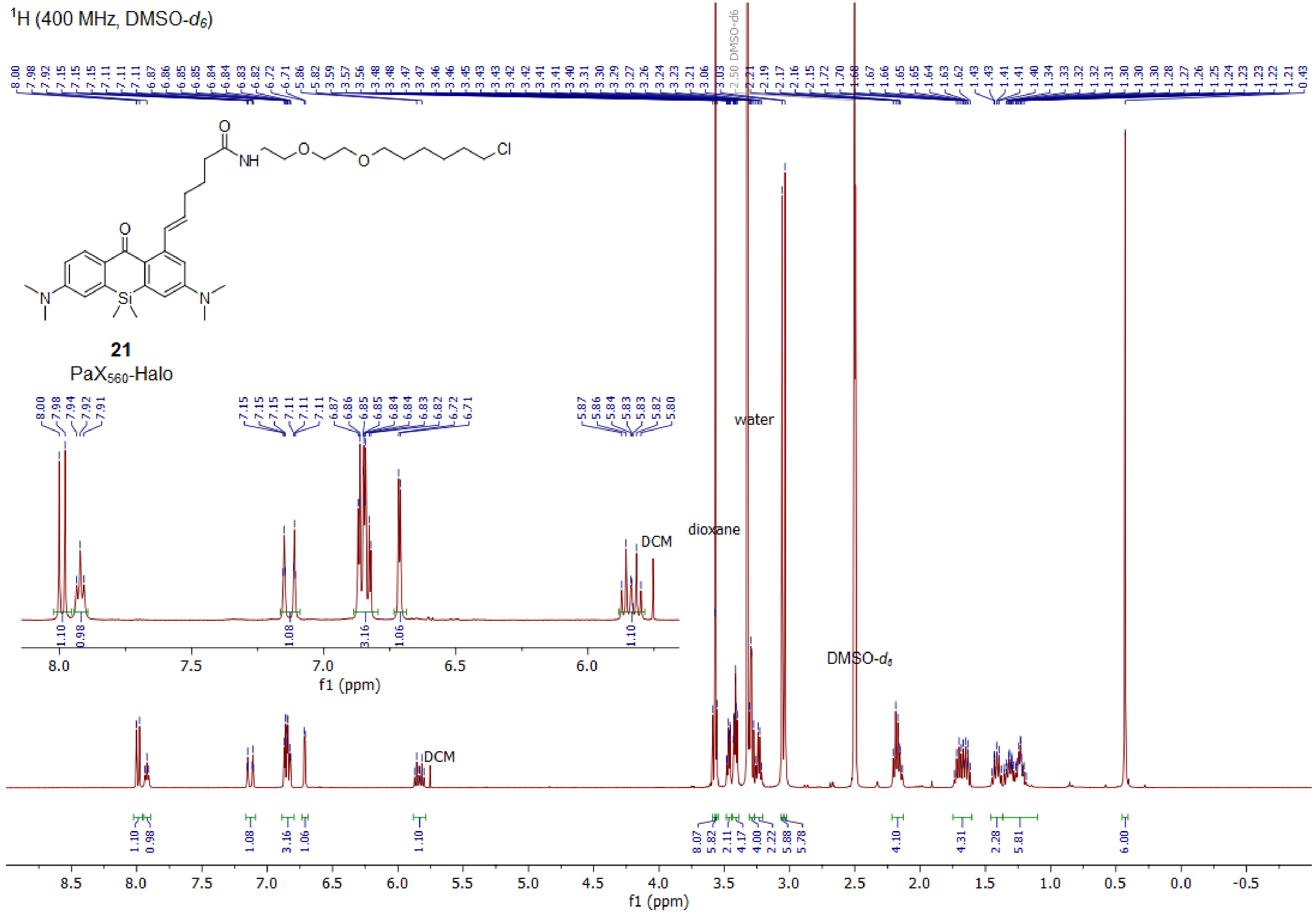

¹H (400 MHz, DMSO-d6)
21
PaX560-Halo
water
dioxane
DCM
DMSO-d6
f1 (ppm)

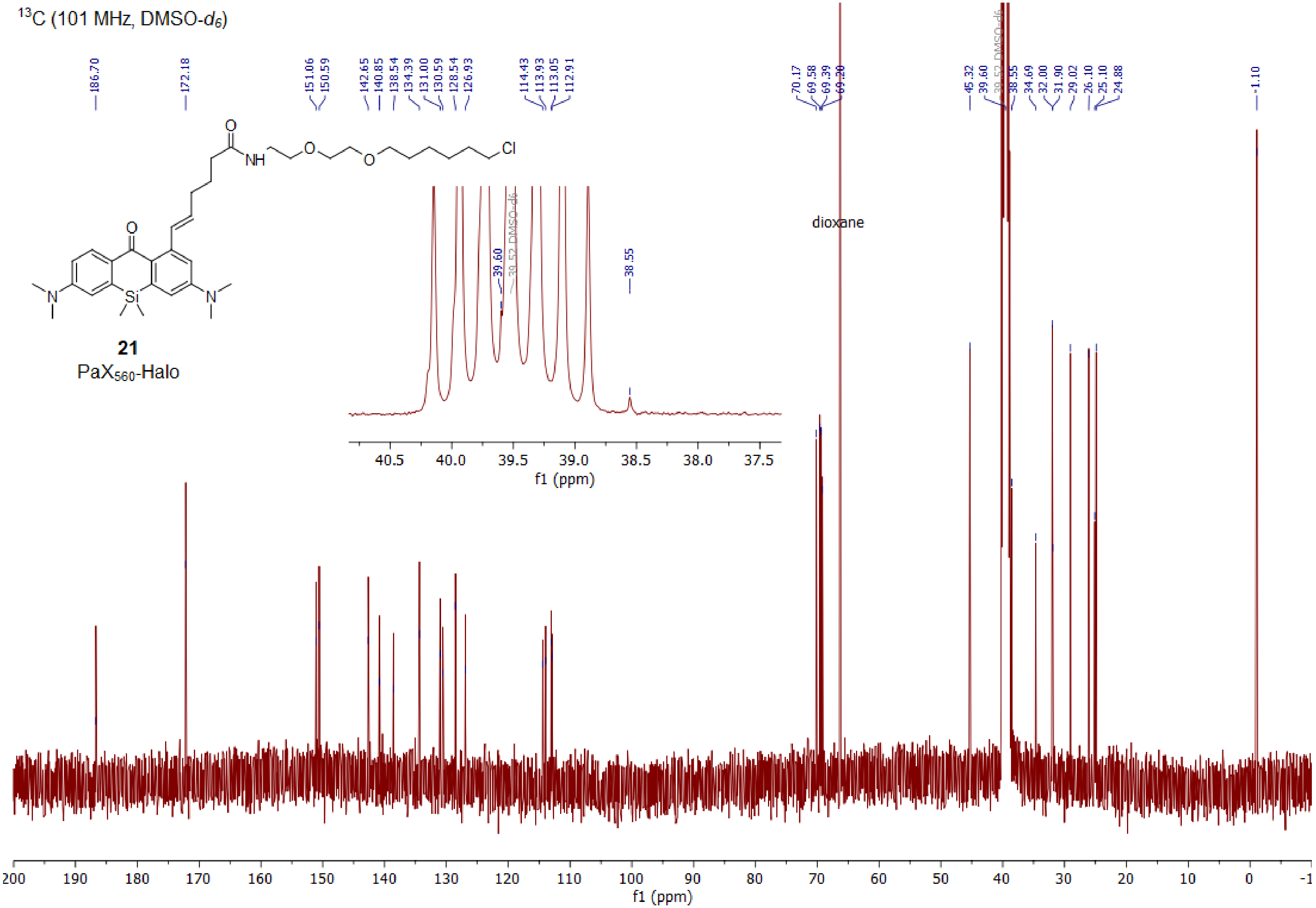

13C (101 MHz, DMSO-d6)
186.70
172.18
151.06
150.59
142.65
140.85
138.54
134.39
131.00
130.59
128.54
126.93
114.43
113.93
113.05
112.91
70.17
69.58
69.39
69.20
45.32
39.60
39.52 DMSO-d6
38.55
34.69
32.00
31.90
29.02
26.10
25.10
24.88
-1.10
dioxane
21
PaX560-Halo
f1 (ppm)

$^1H$ (400 MHz, DMSO-$d_6$)

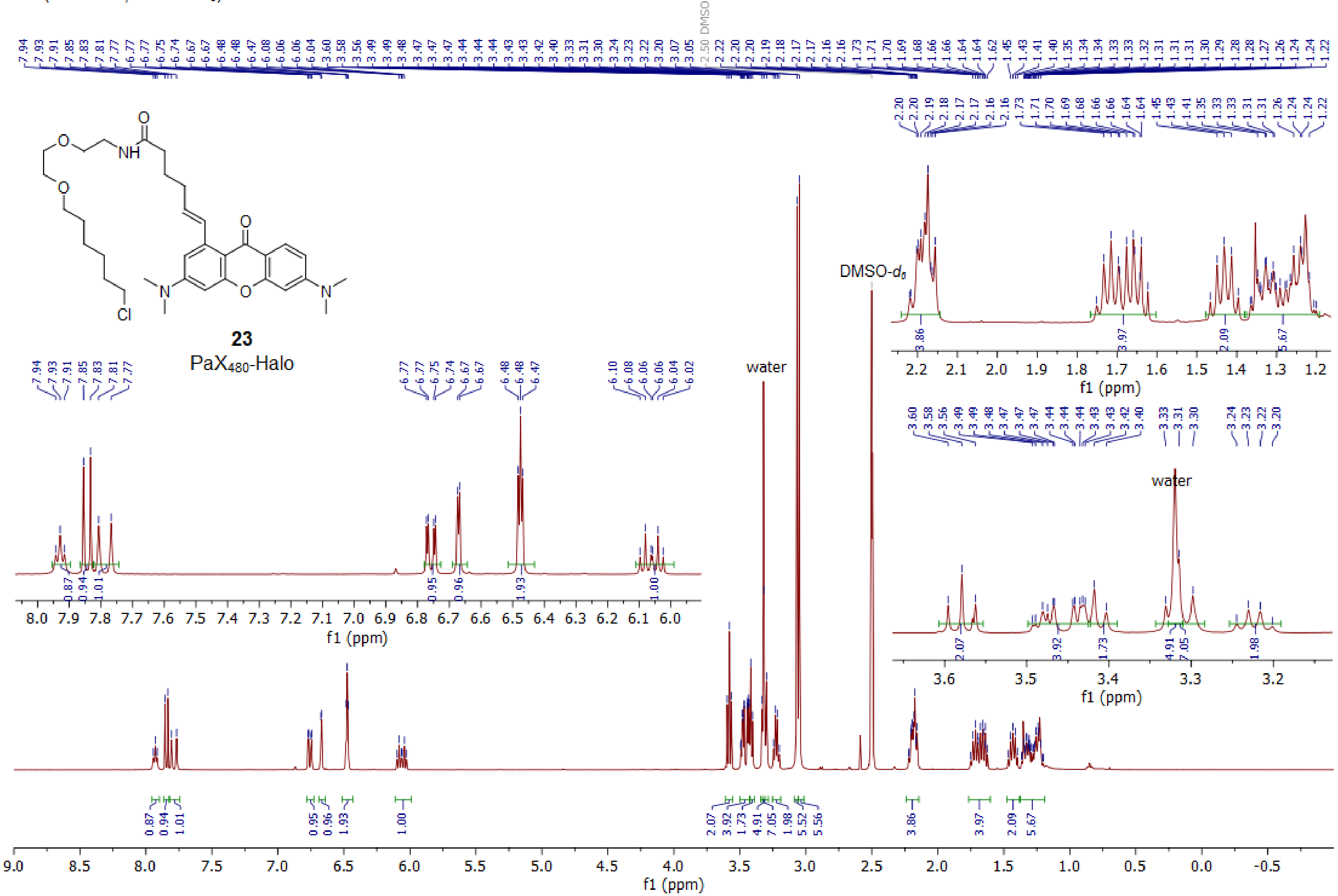

$^{13}C$ (101 MHz, DMSO-$d_6$)

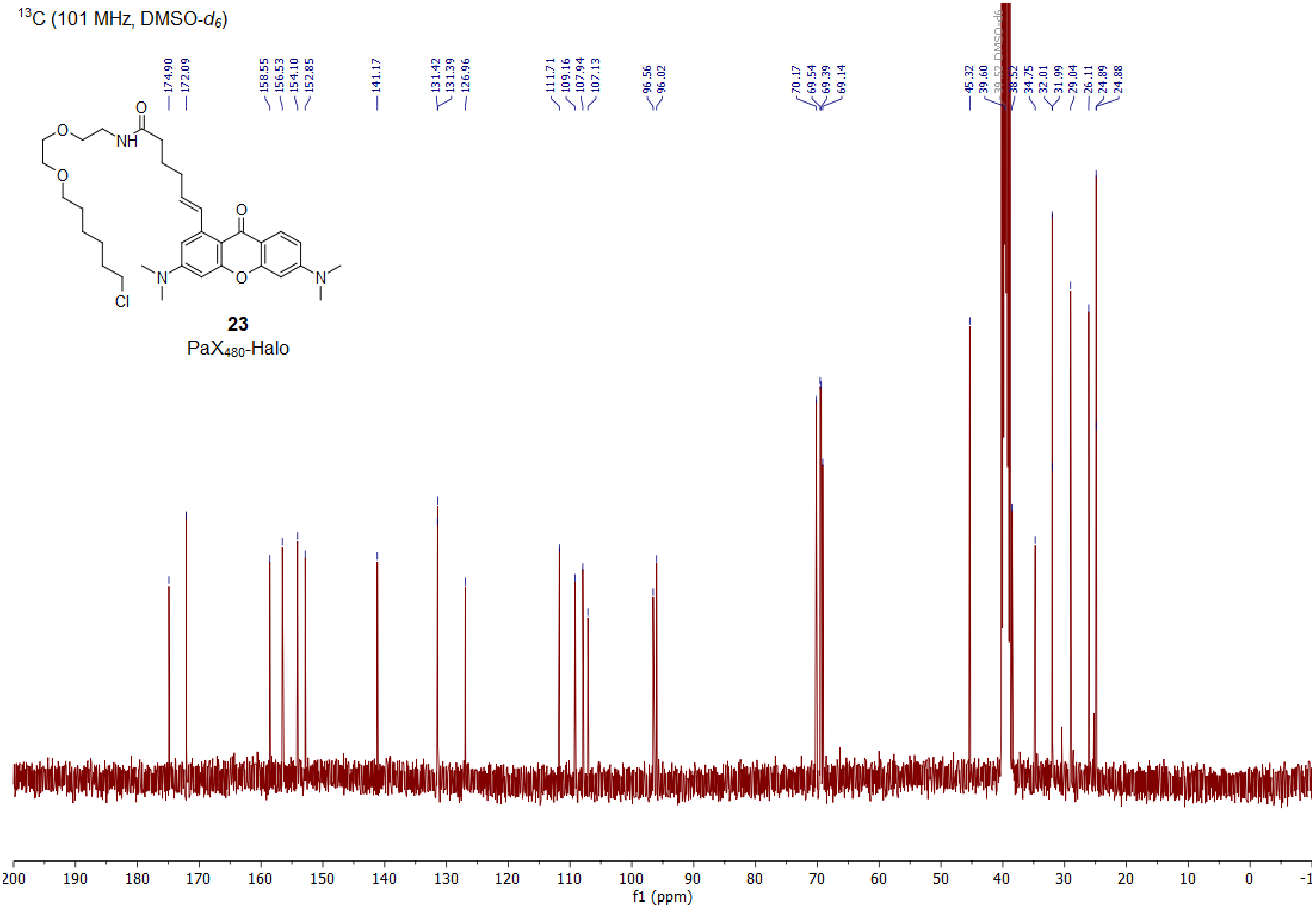

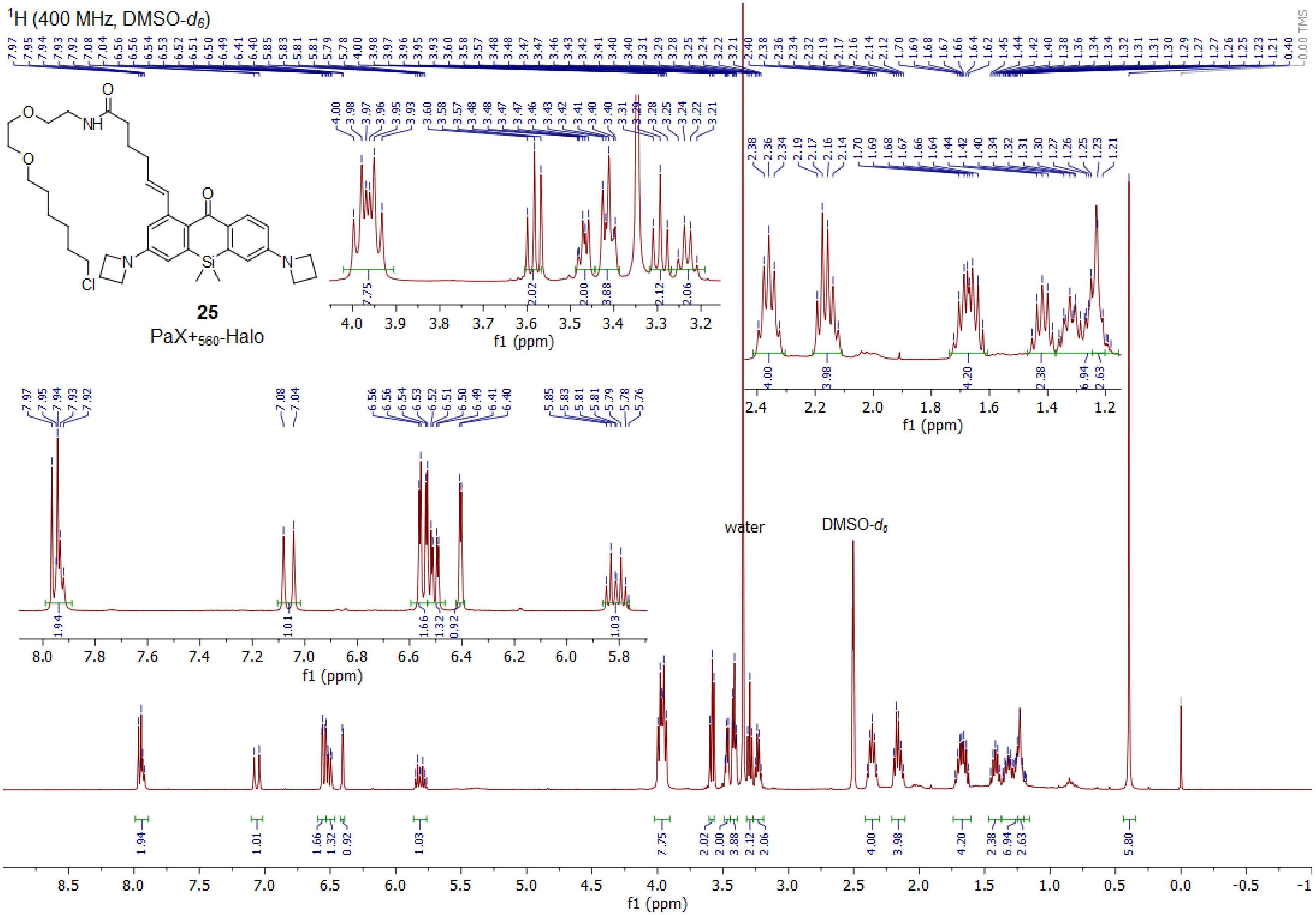
[1]H (400 MHz, DMSO-d6)
25
PaX+560-Halo
water
DMSO-d6
f1 (ppm)
0.00 TMS

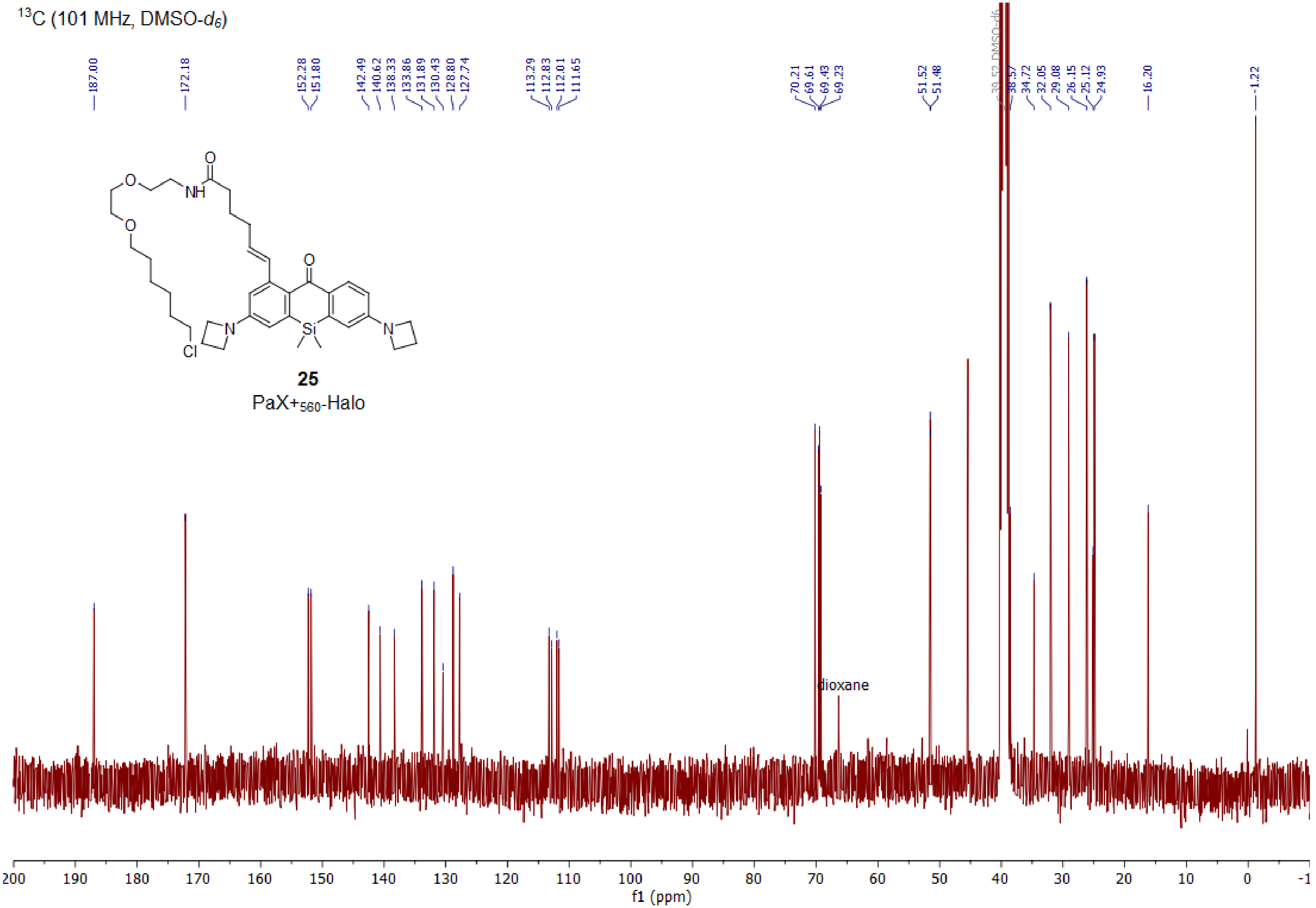

$^{13}C$ (101 MHz, DMSO-$d_6$)
187.00
172.18
152.28
151.80
142.49
140.62
138.33
133.86
131.89
130.43
128.80
127.74
113.29
112.83
112.01
111.65
70.21
69.61
69.43
69.23
51.52
51.48
39.52 DMSO-d6
38.57
34.72
32.05
29.08
26.15
25.12
24.93
16.20
-1.22
25
PaX+560-Halo
dioxane
200 190 180 170 160 150 140 130 120 110 100 90 80 70 60 50 40 30 20 10 0 -1
f1 (ppm)
