## Supplementary figures and images for "A general design of caging-group free photoactivatable fluorophores for live-cell nanoscopy"

### pwm_ledarray_v1_board.pdf

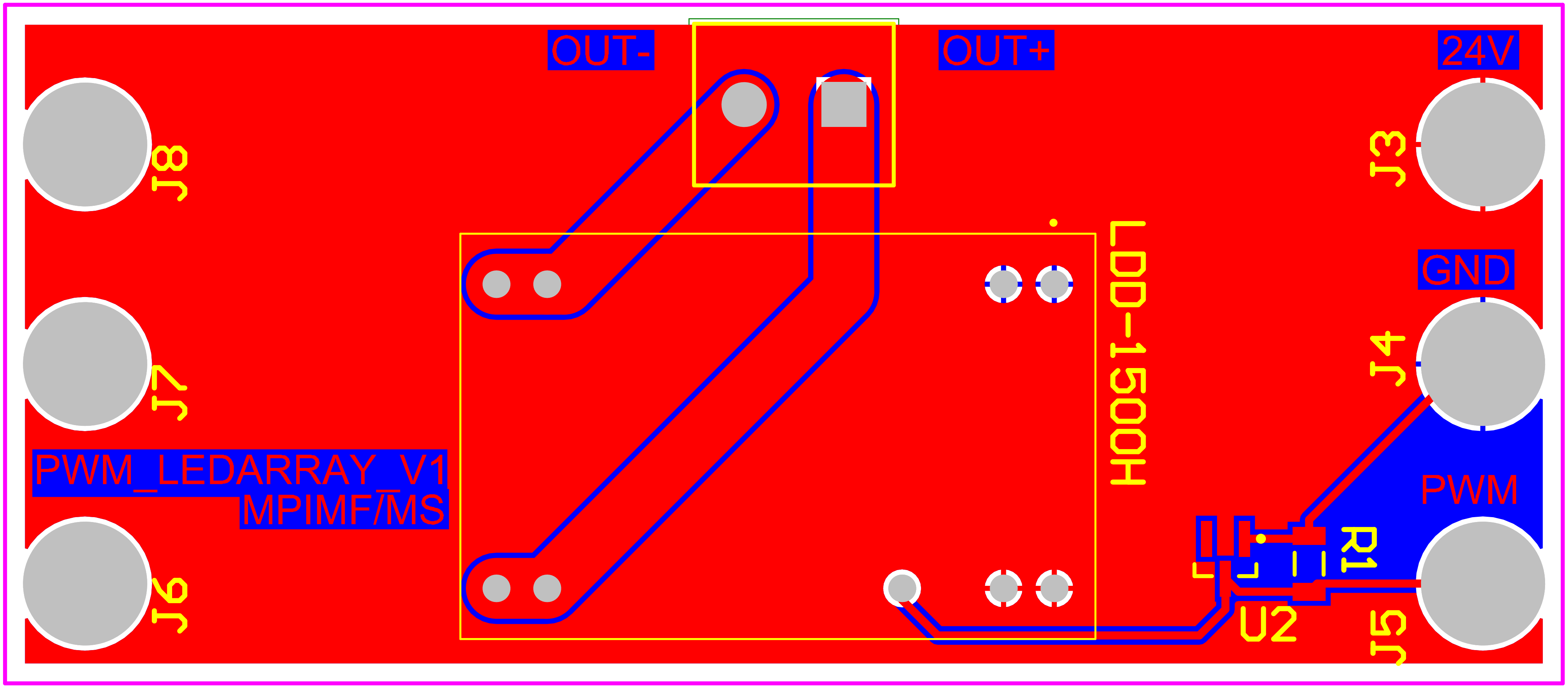

### pwm_ledarray_v1_schematic.pdf

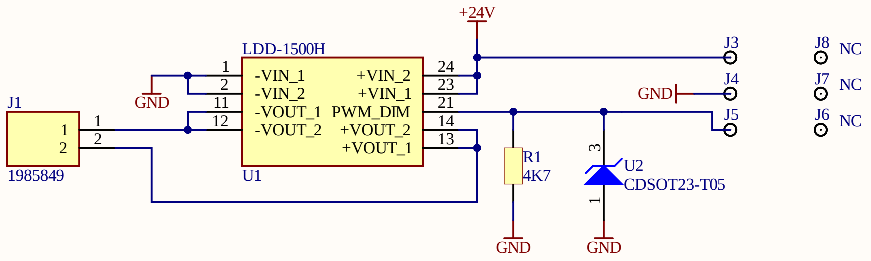


| Title | | |
|---|---|---|
| Size A4 | Number | Revision |
| Date: 8/31/2021 | | Sheet of |
| File: Z:\projects\..\leddriver.SchDoc | | Drawn By: |
